# Ligand-induced Conformational Plasticity of the CTLH E3 Ligase Receptor GID4

**DOI:** 10.1101/2025.07.01.662521

**Authors:** Daria Kotlarek, Katarzyna Dudek, Bartosz Woźniak, Martyna W. Pastok, Dmitrii Shishov, Sylvain Cottens, Michał Biśta, Emilia Krzywiecka, Karolina M. Górecka-Minakowska, Kinga Jurczak, Tomáš Drmota, Justyna Adamczyk, Szymon Faliński, Daria Gajewska, Marta Klejnot, Aleksandra Król, Monika Cuprych-Belter, Iwona Mames, Arnaud Mathieu, Aleksandra Podkówka, Ziemowit Pokładek, Kamil Przytulski, Alicja N. Skowron, Magdalena Sypień, Anna Szlachcic, Toshimitsu Takagi, Weronika Wanat, Igor H. Wierzbicki, Janusz Wiśniewski, Michał J. Walczak

## Abstract

The application of targeted protein degradation (TPD) is currently constrained by the limited availability of low-molecular-weight molecules that can recruit E3 ligases other than CRBN (Cereblon) or VHL (Von Hippel-Lindau ligase). In this study, we present the structure-based drug design (SBDD) of high-affinity ligands that engage E3 ligase GID4 (Glucose-induced degradation protein 4) in biophysical and cellular experiments. Through structural studies and molecular modeling, we identified three clusters of compounds that induce distinct conformations of GID4. We characterized potential exit vectors and used the most promising ligand as a building block to prepare bifunctional degraders in the form of proteolysis-targeting chimeras (PROTACs). Although ternary complex formation was successful *in vitro*, degradation of BRD4 was not observed, highlighting the need for further optimization of the degraders. Finally, we theoretically investigated the likelihood of the identified GID4 conformations participating in protein-protein interactions mediated by molecular glue mechanisms. We believe the expanded ligand diversity discovered in this study may pave the way for tuning the selectivity and efficacy of interactions involving GID4 and its neosubstrates.

**GRAPHICAL ABSTRACT:** 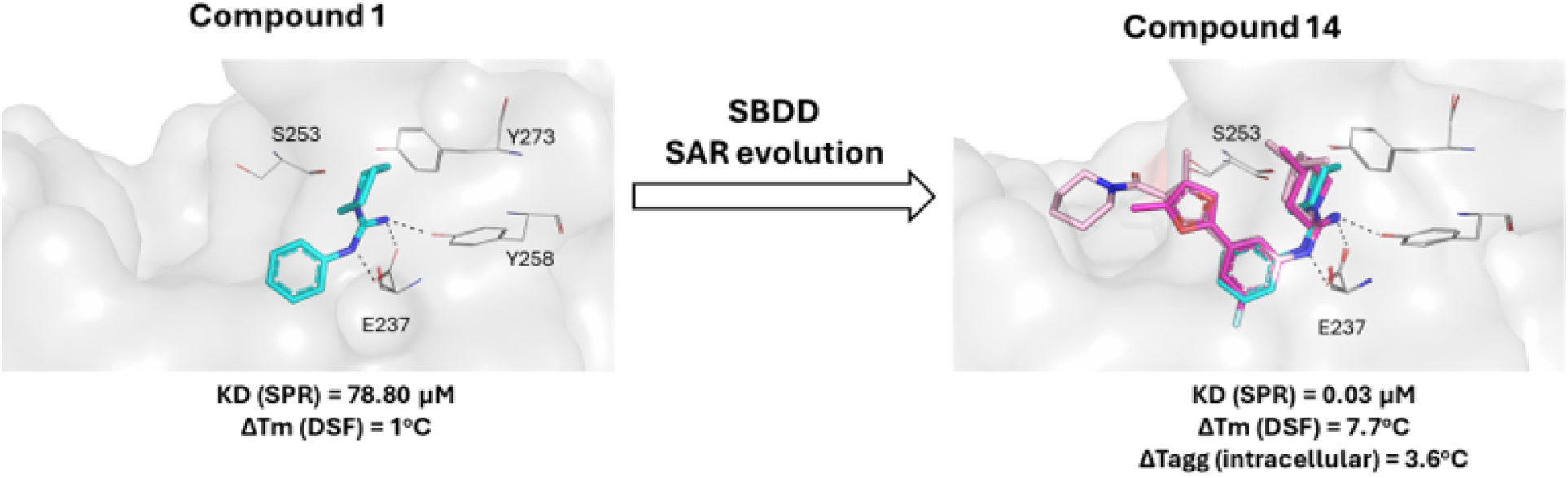

## INTRODUCTION

Targeted protein degradation (TPD) is emerging as a new modality in drug discovery and, as opposed to protein inhibitors, offers the pharmacology of events rather than pharmacology of occupancy.^[1,2]^ TPD relies on the utilization of the endogenous ubiquitin-proteasome system (UPS) machinery to degrade disease-related proteins. This mechanism is mediated either via molecular glues or modular molecules called proteolysis targeting chimeras (PROTACs). ^[3–5]^ Despite differences between these two modalities, both converge by bringing an E3 ligase complex and a protein of interest (POI) into proximity for subsequent ubiquitination and proteasomal degradation. TPD offers significant advantages over traditional inhibitor-based approaches, most notably the ability to target proteins without well-defined binding pockets - such as transcription factors and scaffolding proteins - which have historically been considered “undruggable”.^[6,7]^ Nonetheless, a substantial obstacle to the rapid progress of the technology is the limited number of validated E3 ligases, beyond the frequently employed “workhorses”, cereblon (CRBN) and von Hippel-Lindau (VHL).^[8,9]^ Therefore, recent efforts aim to extend the diversity of E3 ligases for TPD application, drawing from approximately 600 E3 ligases identified in humans. It is expected that the discovery of chemical handles to engage more E3 ligases will enable targeted protein degradation in a tissue-or disease-specific manner. Furthermore, a greater variety of available E3s would be beneficial for combination therapies, reducing side effects, and minimizing acquired resistance.

The C-terminal to LisH (CTLH) complex is an evolutionarily conserved, multi-subunit complex maintaining the E3 ligase activity. It is known by various subunit nomenclatures across different species.^[10–12]^ There is no exact match between GID and CTLH subunits across organisms, as their structures and substrate-control mechanisms have evolved divergently.^[13]^ GID function is well-understood in budding yeast, where it participates in the degradation of the gluconeogenic enzymes once the cell enters glucose-replete conditions.^[14,15]^ In higher eukaryotes, although CTLH E3 enzymes have not been definitively linked to the control of gluconeogenesis, they are hypothesized to mark glycolytic enzymes with ubiquitin for non-proteolytic regulation. This suggests a sophisticated regulatory mechanism in which these enzymes modulate carbohydrate metabolism, even if the precise connection to gluconeogenesis remains to be fully elucidated.^[16]^ In yeast, this metabolic switch depends on the CTLH substrate receptor, Glucose-Induced Degradation protein 4 homolog (GID4).^[17,18]^ This observation paved the way for the identification of the N-terminal proline-containing sequence (Pro/N-end) of the enzymes Fbp1, Icl1, and Mdh2 as the cognate recognition motif for GID4.^[19,20]^ The molecular details of GID4/CTLH binding to the Pro/N-end degron were further elucidated by crystal structures of human GID4 (hGID4) bound to N-terminal peptides.^[21]^ hGID4 adopts an eight-stranded β-barrel fold containing a deep central cavity. This central tunnel, surrounded by four flexible loops (L1-L4), forms the specific binding pocket for the N-terminus of the substrate peptide.^[22]^ Available crystal structures suggest a substantial degree of plasticity within the binding pocket, indicating the capacity to accommodate various endogenous proteins. This structural flexibility makes GID4 both an intriguing and challenging target for the development of small molecule binders. Indeed, the ability to accommodate various types of degron sequences inside the GID4 binding pocket was demonstrated by Dong *et al.* through mutation-scanning peptide arrays and phage display screening. These studies revealed the importance of hydrophobic residues at position 1 (Pro, Ile, Leu, Val, Phe, or Trp), typically followed by a small side-chain residue at position 2. While most amino acids are tolerated at position 3 - except for Gly, Asp, and Pro - the downstream sequence pattern is important for binding and affinity.^[23]^ This sequence promiscuity is facilitated by the pliability of the four hairpin loops (L1-L4), as evidenced by the crystal structures of hGID4. Taken together, these findings suggest that GID4 may engage a broader range of substrates beyond those containing the canonical Pro/N-degron motif.

The physiological substrates of hGID4 have recently been identified utilizing tools such as the GID4-specific chemical probe PFI-7 and methods including proximity-dependent biotinylation and degradomics.^[24]^ Notably, the mammalian counterparts of *S. cerevisiae’s* proteins Fbp1, Icl1, and Mdh2 lack the N-terminal-Pro (Nt-Pro) residue and are not targeted by the human GID4/CTLH complex. Recently, Yi *et al.* established a functional link between yeast and human GID4/CTLH by demonstrating the degradation of the metabolic enzyme, 3-hydroxy-3-methylglutaryl (HMG)-coenzyme A (CoA) synthase 1 (HMGCS1).^[25]^ HMGCS1 is recognized by GID4/CTLH via its Nt-Pro degron motif upon nutrient stress and mTORC1 inhibition. Furthermore, Bagci *et al.* demonstrated that the GID4/CTLH complex regulates cell migration by recognizing the proline degron and mediating the degradation of a GTPase activating enzyme ARHGAP11A.^[26]^ Further, Owens *et al.* linked the GID4/CTLH complex to proteins involved in RNA processing, transcription, splicing, and chromatin remodeling. Interestingly, although some interactors contain the Nt-Pro degron (*e.g*., DDX17, DDX21, DDX50) and compete with PFI-7 for GID4 binding, GID4-dependent degradation of these substrates was not observed. Furthermore, other interactors identified in that study lacked the specific degron sequence and were not degraded, suggesting potentially a broader functional repertoire for the GID4/CTLH complex beyond proteolysis.^[27]^ In this work, we present the development of high-affinity chemical probes for GID4, utilizing Fragment-Based Screening (FBS) followed by a structure-guided optimization approach. These efforts yielded a series of GID4 ligands that stabilize the protein in distinct conformations. We identified three conformational states based on the measured distances between the mobile hairpin loops (L1-L4) and compared these to previously reported GID4 crystal structures. Ultimately, our lead compound (**14**), exhibits an affinity of 23 nM in biophysical assays and demonstrates target engagement in a cellular environment. We additionally explored a suitable linker attachment point on compound 14 for the synthesis of bifunctional degraders. These findings deepen the understanding of GID4 binding pocket plasticity and provide a significant addition to the limited toolkit of available GID4 chemical probes.

## RESULTS

### Identification of Compound 1 - Fragment Hit of GID4

To develop GID4-selective ligands, we conducted biophysical screenings of a library comprising approximately 2200 fragments. Two versions of recombinant GID4 (amino acid region 116-300) were purified for this study: an untagged version, and an N-terminally Avi-tagged version. The untagged protein was employed in differential scanning fluorimetry (DSF), while the biotinylated Avi-tagged protein was utilized for surface plasmon resonance (SPR) experiments. To confirm the integrity and activity of the GID4 protein, DSF was performed in the presence of c=50 μM endogenous degron peptide, PGLWKS. The untagged GID4 protein exhibited a denaturation temperature of Tm = 61.5 °C with a sigmoidal two-state transition, increasing to Tm = 62.4 °C in the presence of the peptide, confirming the high stability of the folded protein. Immobilization of the biotinylated GID4 on the SPR chip yielded quality sensograms indicating saturable binding of the PGLWKS degron peptide. The measured affinity (K_D_) of 2.13 μM was consistent with previously reported literature data (Figure S1).^[21]^ Next, we performed the DSF screen at a constant fragment concentration (c=500 μM) that revealed two hits. Among these, compound **1** induced the largest thermal shift (ΔTm = +1.0°C), indicating binding and consequent protein stabilization. In parallel, SPR was used as an orthogonal screening method. Primary binding level and dose-response screens revealed 28 fragments with K_D_ ^GID4^ < 1 mM (hit rate ∼1.3%), including both hits identified by DSF. Subsequent validation confirmed compound **1**, which contains an N,N’-disubstituted guanidine moiety, as the most potent fragment with the K_D_^GID4^ of 78.80 μM as determined by a steady-state affinity model (Figure 1).

**Figure 1.**
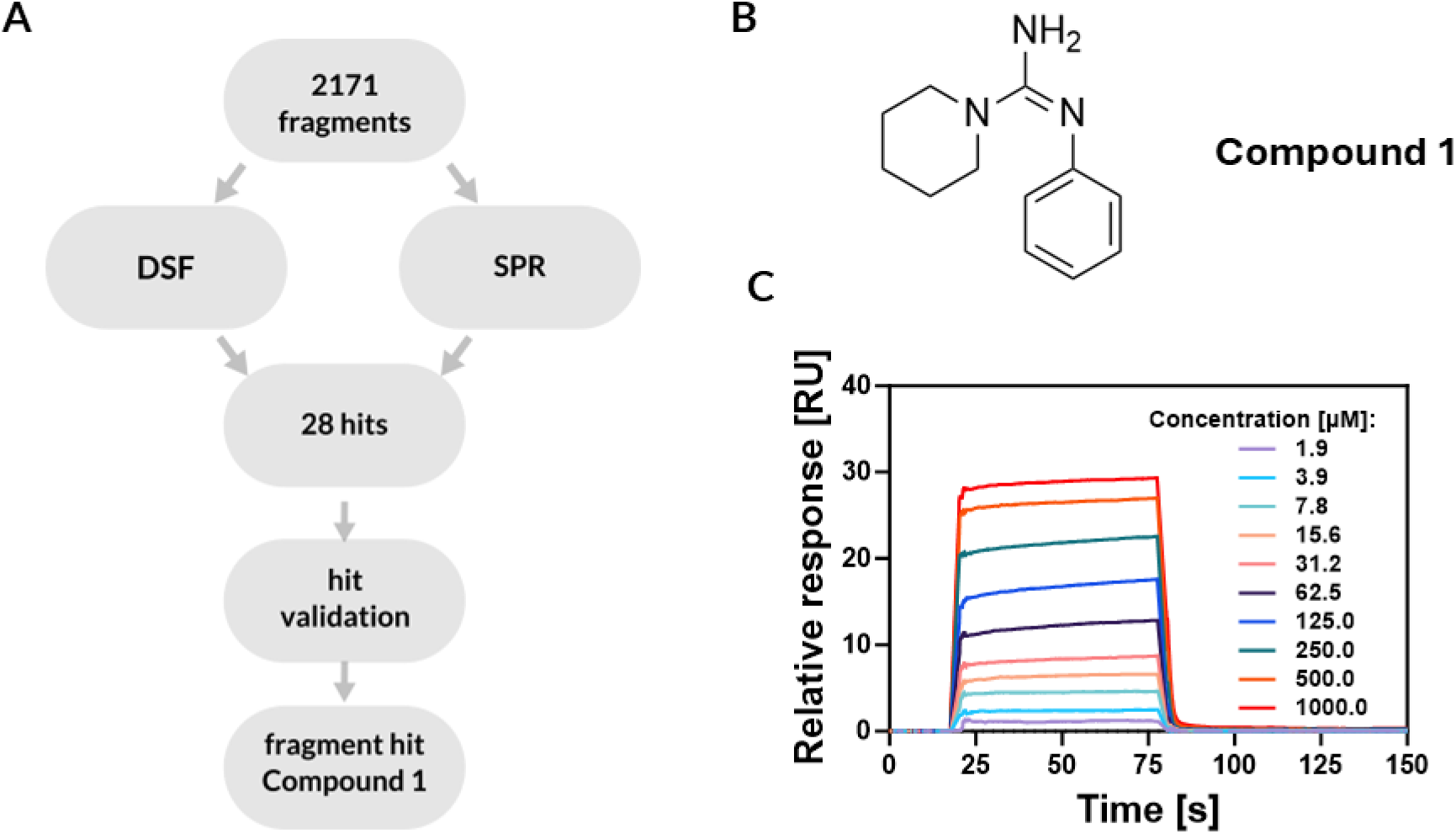
Identification of fragment hit - compound 1. **A)** Workflow of hit identification, **B)** Chemical structure of compound 1. **C)** Representative SPR sensograms demonstrating binding of compound 1 to the immobilized GID4. K_D_^GID4^ was determined by a steady-state affinity binding model (N=3).

Next, to understand the binding of compound **1**, we determined its co-crystal structure with GID4 (see Table S1 for X-ray diffraction data and model refinement statistics, PDB: 9QDX). As previously reported, GID4 adopts an antiparallel, eight β-strand barrel fold linked with an insert of three α-helices.^[21]^ One end of the β-barrel contains a deep hydrophobic binding pocket surrounded by four loops: L1 (134-138), L2 (163-169), L3 (240-250), and L4 (275-279). These loops define a degron-binding site which overlaps with the ligand site. A comparison of the L2 and L3 loop positions between the compound **1**-bound structure and the degron-bound structure (PDB: 6CD9) reveals that the loops in the degron-bound form adopt a more ‘closed-like’ conformation. This results in a more constricted binding pocket relative to the hit-bound structure (Figure S2F). Furthermore, loop 3 in the compound **1**-bound form appears more mobile, as the electron density for residues 246-248 is not well-defined, suggesting increased flexibility in this region upon ligand binding. The N, N’-disubstituted guanidine moiety of compound **1** mimics the binding of the N-terminal proline of the endogenous degron peptide, consistent with previously reported GID4 crystal structures (PDB: 6CDG, 6CD9, 6CDC). In the reported crystal structure with compound **1**, Glu237 forms hydrogen bonds with the guanidinium group. Notably, the Glu237Ala mutation has been reported to abolish interactions with the degron peptide. According to the COSMIC database, a Glu237Lys mutation has also been identified in breast carcinoma (https://cancer.sanger.ac.uk/cosmic).^[21,28]^ The guanidinium cation of compound **1** engages in a putative cation–π interaction with Tyr273, while simultaneously acting as a hydrogen bond donor to the phenolic oxygen of Tyr258. Although Tyr273 contains a polar hydroxyl group, its aromatic ring contributes to the protein interior by extending its phenyl group deeper into the hydrophobic core toward Leu171 and Ile161 – reaching further than the native degron peptide. This orientation likely facilitates the observed cation-π interaction between Tyr273 and compound **1**. Additionally, the guanidine moiety superimposes the pyrrolidine ring of the proline residue from the GID4 degron tetrapeptide PSRW (PDB: 6CD9). In our structure, the piperidine ring of compound **1** is positioned within weak hydrogen-bonding distance of Ser253 (∼3.05 Å) and Gln132 (∼3.08 Å). Glutamine 132 is the second key residue that stabilizes the N-terminal proline of the degron peptide via hydrogen bonds; its mutation to alanine significantly reduces this interaction.^[21]^ The GID4 binding pocket is predominantly hydrophobic, lined by the aromatic side chains of Tyr273, Phe254, as well as aliphatic residues: Ile249, Leu240, Leu171, Leu164, Ile161, Leu159 and Val141 (Figure 2B and Figure 2D). This hydrophobic character is essential for binding as alanine-substitutions of Tyr273 and neighboring hydrophobic residues (Phe254, Leu240, Leu171, Leu159, Val141, Ile161 and Leu164) disrupt this patch, resulting in 6-to 14-fold reduction in degron binding affinity.^[21]^

**Figure 2.**
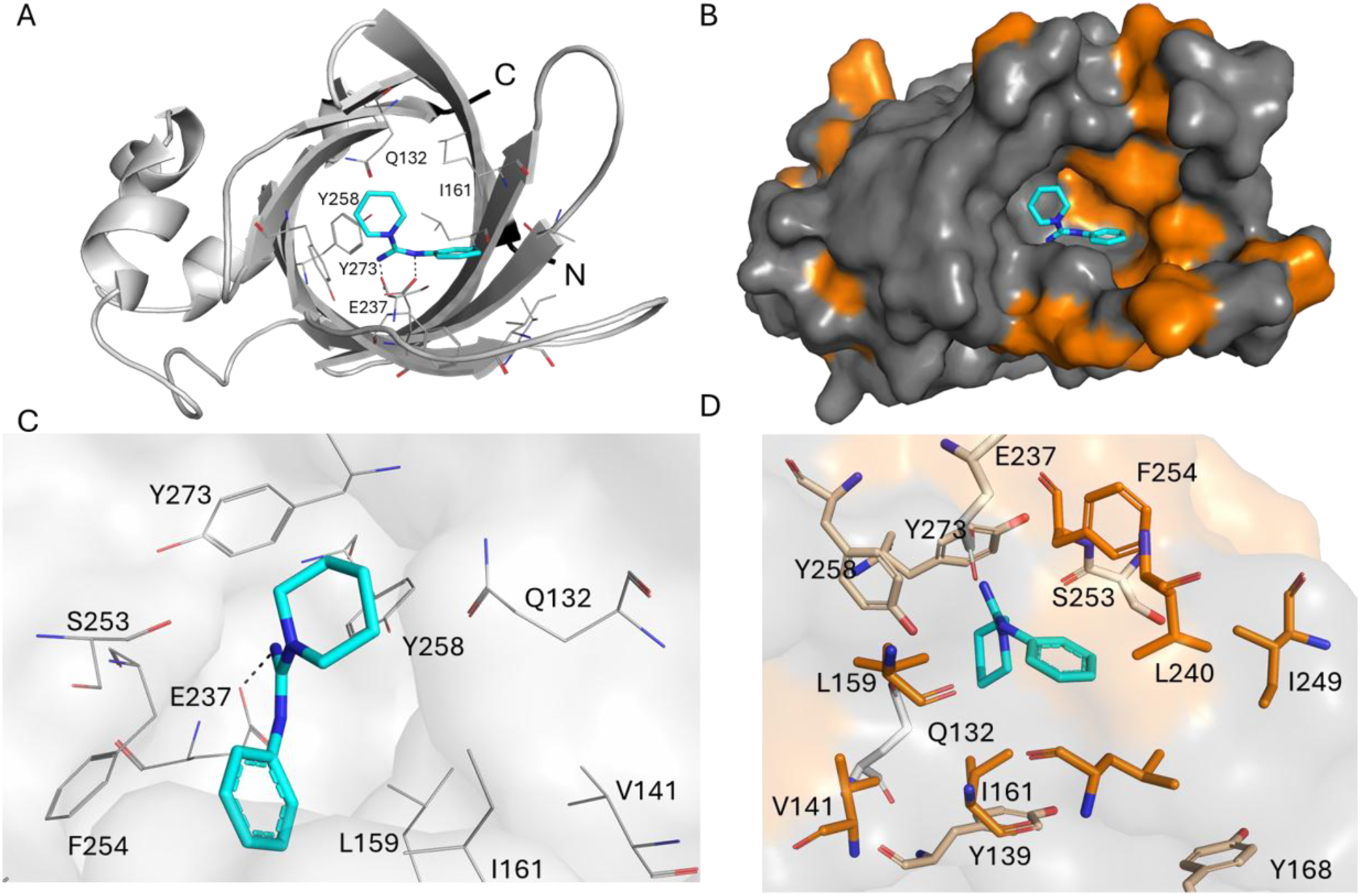
The crystal structure of GID4 complex with compound 1 (PDB: 9QDX). **A)** Top view of the GID4 binding pocket with compound 1 (turquoise) bound in the degron binding pocket, and GID4 represented as cartoon (grey). The N-and C-termini of the protein are labeled as N and C, respectively. **B)** The surface representation of GID4 with all hydrophobic residues colored orange (the rest of the amino acids are grey) with compound 1 (turquoise) bound in the GID4 binding pocket. The binding pocket has a hydrophobic character. **C)** Key interactions with Glu237, Ser253 and Tyr273 are labeled for reference. **D)** Close view of the hydrophobic pocket, where the phenyl moiety of compound 1 is surrounded by hydrophobic residues Ile161, Leu171, Leu240, and Phe254.

### Development of GID4 Ligand Series

Following the identification of the binding pocket and the key interactions of the initial fragment, we initiated the synthesis of a series of analogs to explore the structure-activity relationship (SAR) (Table 1). The resulting changes in affinity were evaluated using surface plasmon resonance (SPR) and differential scanning fluorimetry (DSF). The monohalogenated analogs (compounds **2** and **3**) validated the chemical series as a starting point for the hit expansion, with the 3-chloro-5-fluorophenyl derivative showing comparable potency to compound **3**. Exploration of the piperidine substitution pattern indicated that methyl substitutions improved affinity towards GID4 with the most prominent result at positions 2 and 4. In the next step, we replaced the 3-Cl group to find an additional potency boost. Screening of diverse scaffolds revealed a significant improvement in binding affinity for the 5-methyl furan-2-yl substituent, especially when combined with a 4-Me piperidine ring (compound **9**, Table 1).

**Table 1.**
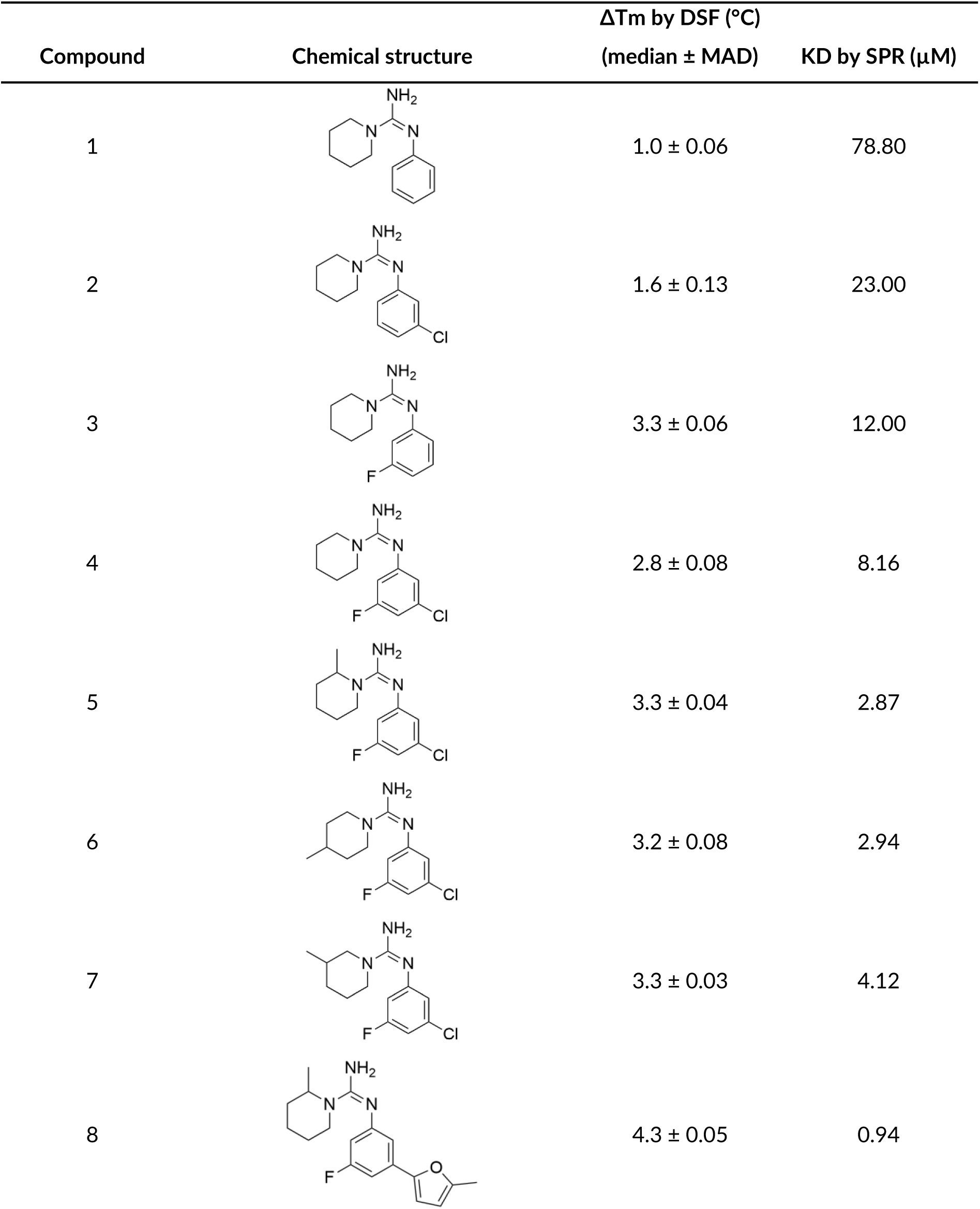
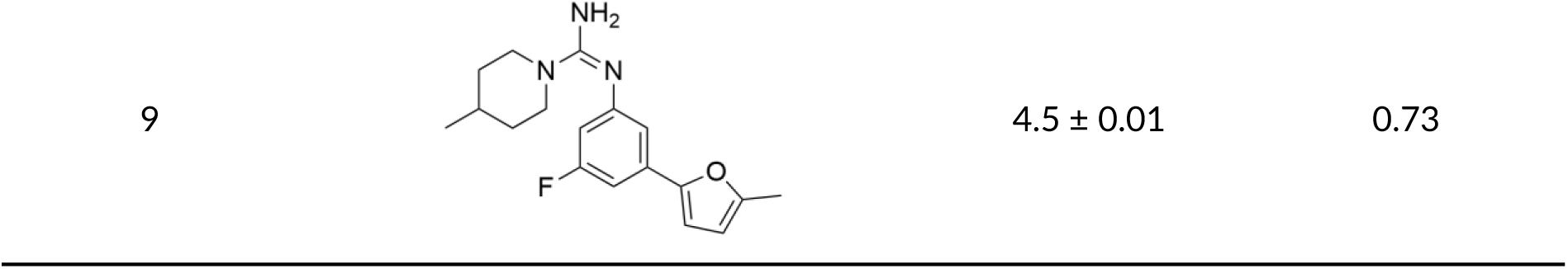
Binding to GID4 by analogs of compound 1.

| Compound | Chemical structure | $\Delta T_m$ by DSF ($^{\circ}\text{C}$) | |
| --- | --- | --- | --- |
| | | (median ± MAD) | KD by SPR ($\mu\text{M}$) |
| 1 | | $1.0 \pm 0.06$ | 78.80 |
| 2 | | $1.6 \pm 0.13$ | 23.00 |
| 3 | | $3.3 \pm 0.06$ | 12.00 |
| 4 | | $2.8 \pm 0.08$ | 8.16 |
| 5 | | $3.3 \pm 0.04$ | 2.87 |
| 6 | | $3.2 \pm 0.08$ | 2.94 |
| 7 | | $3.3 \pm 0.03$ | 4.12 |
| 8 | | $4.3 \pm 0.05$ | 0.94 |

To further elucidate the basis for the enhanced binding of compound **9**, we determined its crystal structure in complex with GID4 (Figure 3, PDB: 9QDY). Compound **9** occupies the GID4 degron-binding pocket in a manner equivalent to structure of compound **1**, with its guanidine moiety recapitulating the interaction with the side chain of Glu237. Moreover, the guanidinium cation also engages in a putative cation-π interaction with Tyr273, while acting as a hydrogen bond donor to the hydroxy group of Tyr258 as seen in compound **1** structure. However, compound **9** extends beyond the canonical peptide cavity to reach Leu171, Tyr168, and Thr173, whereas the peptidic degron occupies the pocket region involving residues: Gly251, His275, and Ser278. The region into which compound **9** extends is lined with hydrophobic residues, exhibiting a more pronounced hydrophobic character that engages the furan and piperidine moieties through hydrophobic contacts (Figure 3D). Resembling compound **1**, the piperidine ring of compound **9** resides in a surface groove formed by Tyr258, Tyr273, Tyr139, and Ile161, which is further lined with Gln132, Gln282, and Ser134. Notably, the piperidine ring is placed within a cone-shaped hydrophobic cavity between Tyr258 and Tyr273. The superior binding affinity of compound **9** relative to compound **1** is likely due to these expanded hydrophobic interactions that reach beyond the canonical degron-binding site, coupled with the stabilization provided by the conserved polar interaction with Glu237. Furthermore, in the compound **9**-bound structure of GID4, loops L2 and L3 adopt conformations similar to those observed in the compound **1**-bound state (Figure S2F). Comparably to compound **1**-bound form, residues 245-248 of loop 3 remain mobile as their electron density is not well-defined.

**Figure 3.**
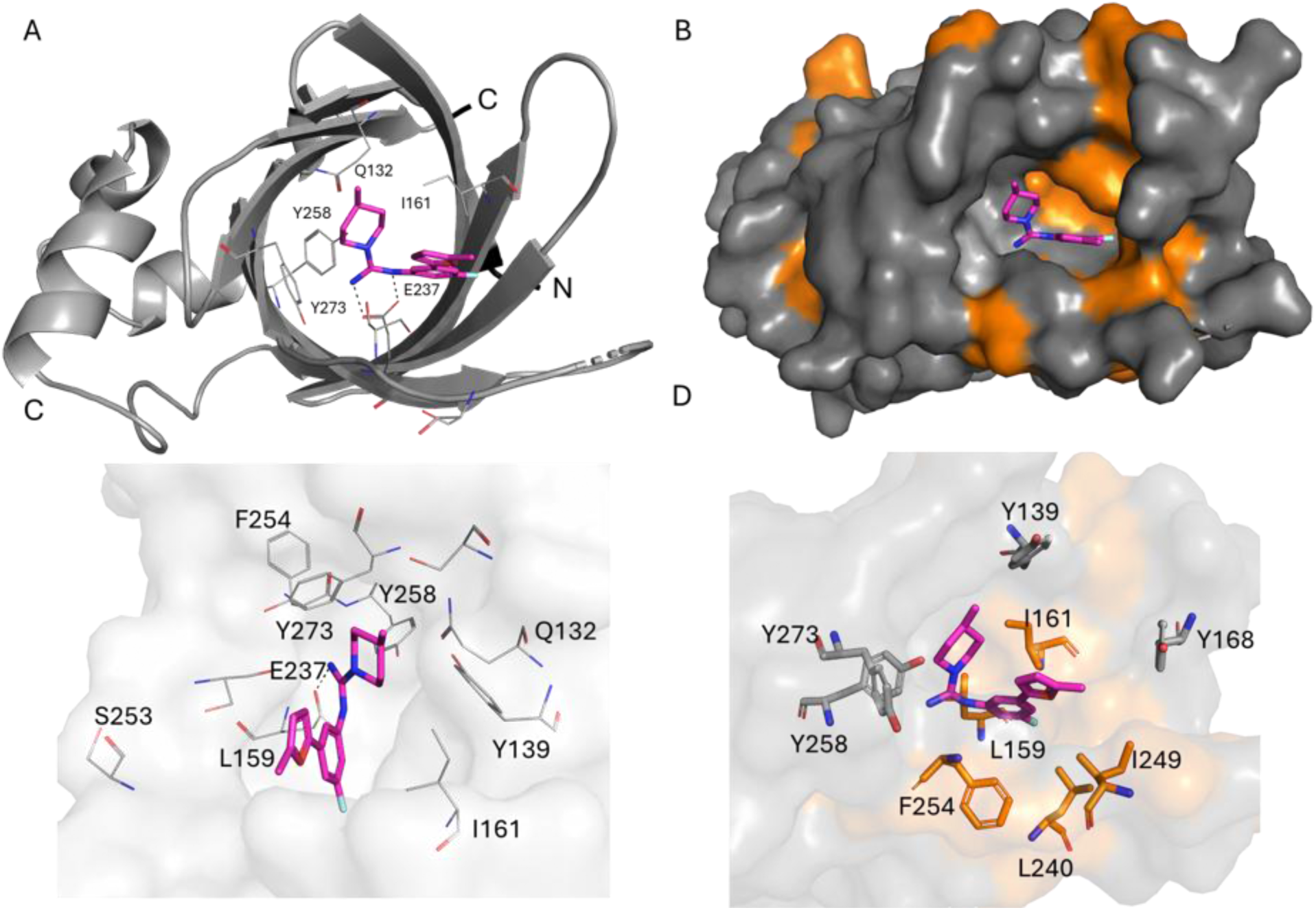
The crystal structure of GID4 protein in complex with compound 9 (PDB: 9QDY). **A)** Top view of the GID4 binding pocket with compound 9 (magenta) bound in the degron binding pocket, and GID4 represented as cartoon (grey). The N-and C-termini of the protein are labeled as N and C, respectively. **B)** The surface representation of GID4 with all hydrophobic residues colored orange (the rest of the amino acids are grey) with compound 9 (magenta) bound in the GID4 binding pocket. The binding pocket has a hydrophobic character. **C)** Key interactions with Glu237, Ser253, and Tyr273 are labeled for reference. **D)** Close view of the hydrophobic pocket, where the phenyl and furan moieties of compound 9 are surrounded by hydrophobic residues Ile161, Leu171, Leu240, and Phe254.

Encouraged by these observations, we continued the optimization of compound **9**. The introduction of another methyl group at the position 4 of the piperidine ring led to an improved affinity of compound **10** (K_D_^GID4^=0.21 µM). However, a notable metabolic liability of this-series is the known instability associated with furan rings.^[29]^ Consequently, our efforts were focused on the substitution of the furan moiety to identify analogs with enhanced potency and improved metabolic stability. To this end, we synthesized a series of amide derivatives at the 4-position of the furan ring (Table 2). Among these, compound **12,** bearing a dimethyl amide group emerged as a potent GID4 ligand with a 4-fold improvement in potency. In contrast, secondary amides - which possess an additional hydrogen bond donor - displayed inconsistent results: compound **11** retained the affinity of compound **10**, while compound **13** showed a significant loss in activity. As a final step in the optimization process, we examined various tertiary amides to assess the influence of ring size and heteroatom type on the binding mode. From these efforts, we found that the piperidine derivative, compound **14**, provided a 2-fold potency boost over compound **12**, whereas the pyrrolidine analog (compound **17)** resulted in a nearly 2-fold loss in activity. Moreover, replacing the carbon at position 4 of the piperidine ring with heteroatoms (O or N) reduced activity towards GID4. Notably, the lead compound in this series (compound **14**, K_D_ ^GID4^=0.03 µM) represents a > 2500-fold increase in potency, relative to the initial hit, compound **1** (K_D_ ^GID4^=78.80 μM).

**Table 2.**
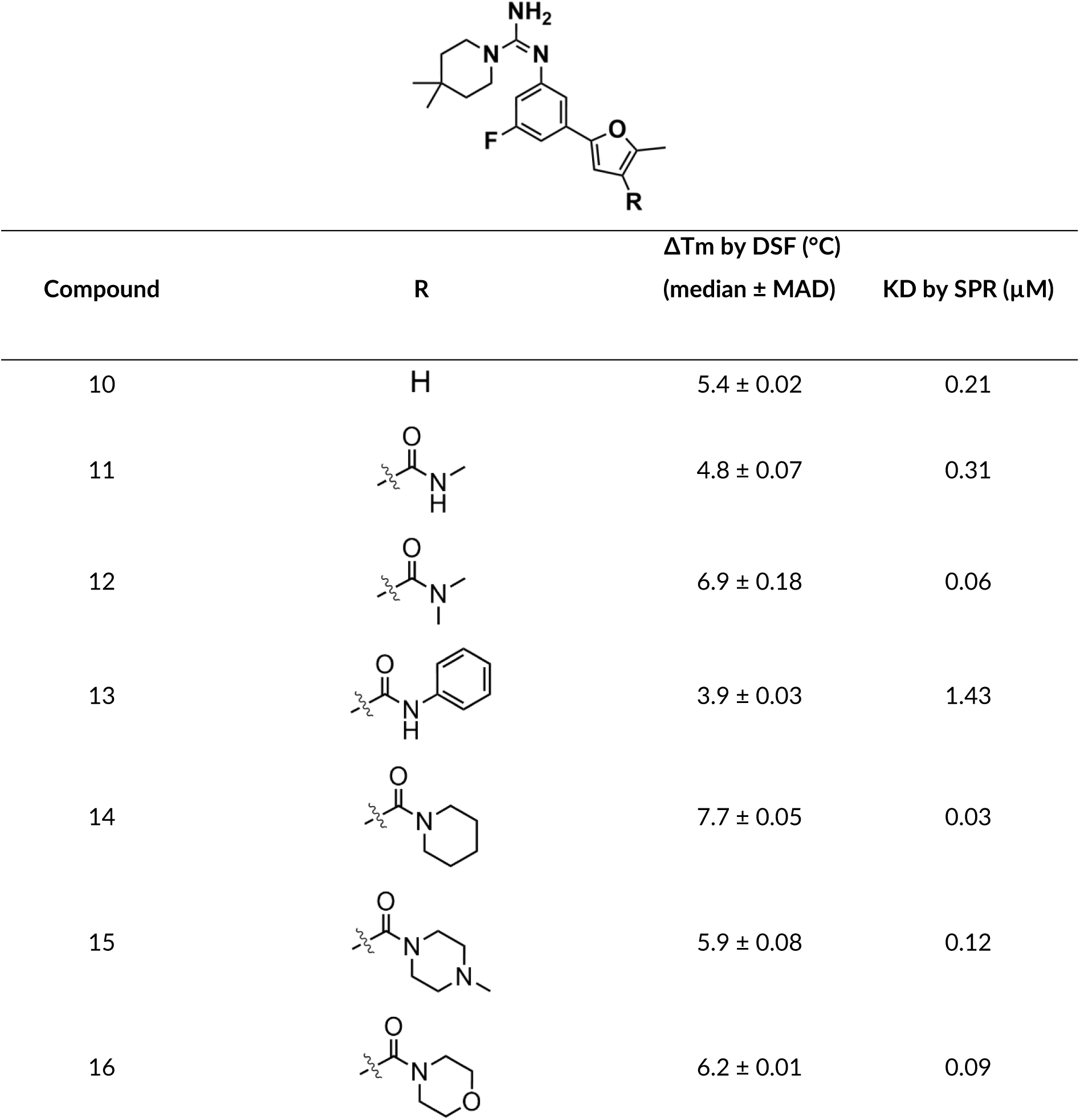
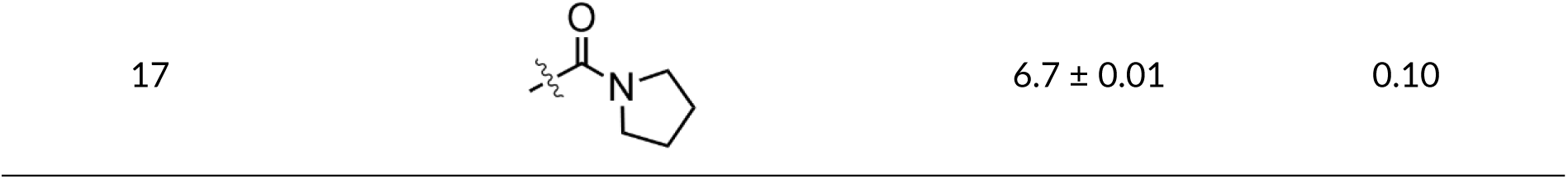
SAR of substituents at C4 of the furan ring.

| Compound | R | $\Delta T_m$ by DSF ($^{\circ}\text{C}$) | |
| --- | --- | --- | --- |
| | | (median ± MAD) | $K_D$ by SPR ($\mu\text{M}$) |
| 10 | H | $5.4 \pm 0.02$ | 0.21 |
| 11 | | $4.8 \pm 0.07$ | 0.31 |
| 12 | | $6.9 \pm 0.18$ | 0.06 |
| 13 | | $3.9 \pm 0.03$ | 1.43 |
| 14 | | $7.7 \pm 0.05$ | 0.03 |
| 15 | | $5.9 \pm 0.08$ | 0.12 |
| 16 | | $6.2 \pm 0.01$ | 0.09 |

Finally, to explain the structural basis for the significant potency enhancement, we determined the crystal structure of GID4 in complex with compound **14** (PDB: 9QDZ). The guanidine moiety of compound **14** interacts with the side chain of Glu237, resembling the binding pose observed for compound **1** and compound **9**. Similar to compound **9**, compound **14** extends beyond the canonical peptide cavity; its furan and piperidine ring moieties are oriented toward Leu171, Tyr168, and Thr173 (Figure 4), effectively exploiting a region of the binding pocket not occupied by the N-degron. Furthermore, compound **14** interacts with an even bigger surface of the binding pocket compared to compound **9**, making more contacts with the amino acid side chains and reaching towards loop 3 and Tyr168. The guanidinium cation of compound **14** is also additionally positioned proximal to Tyr273, consistent with a potential cation-π interaction, and donates a hydrogen bond to the phenolic oxygen of Tyr258. Moreover, in the compound **14**-bound state, the piperidine side chain attached to the guanidine moiety - which superimposes the second residue of the degron peptide-bound form - extends toward Ser253. The loops L2 and L3 adopt conformations comparable to those observed in the compound-bound GID4 structure (Figure S2F). The superior affinity of compound **14** relative to compounds **9** and **1** is likely attributed to a combination of an additional interaction with Ser253, more extensive hydrophobic contacts within the pocket, and stabilization provided by the polar interaction with Glu237.

**Figure 4.**
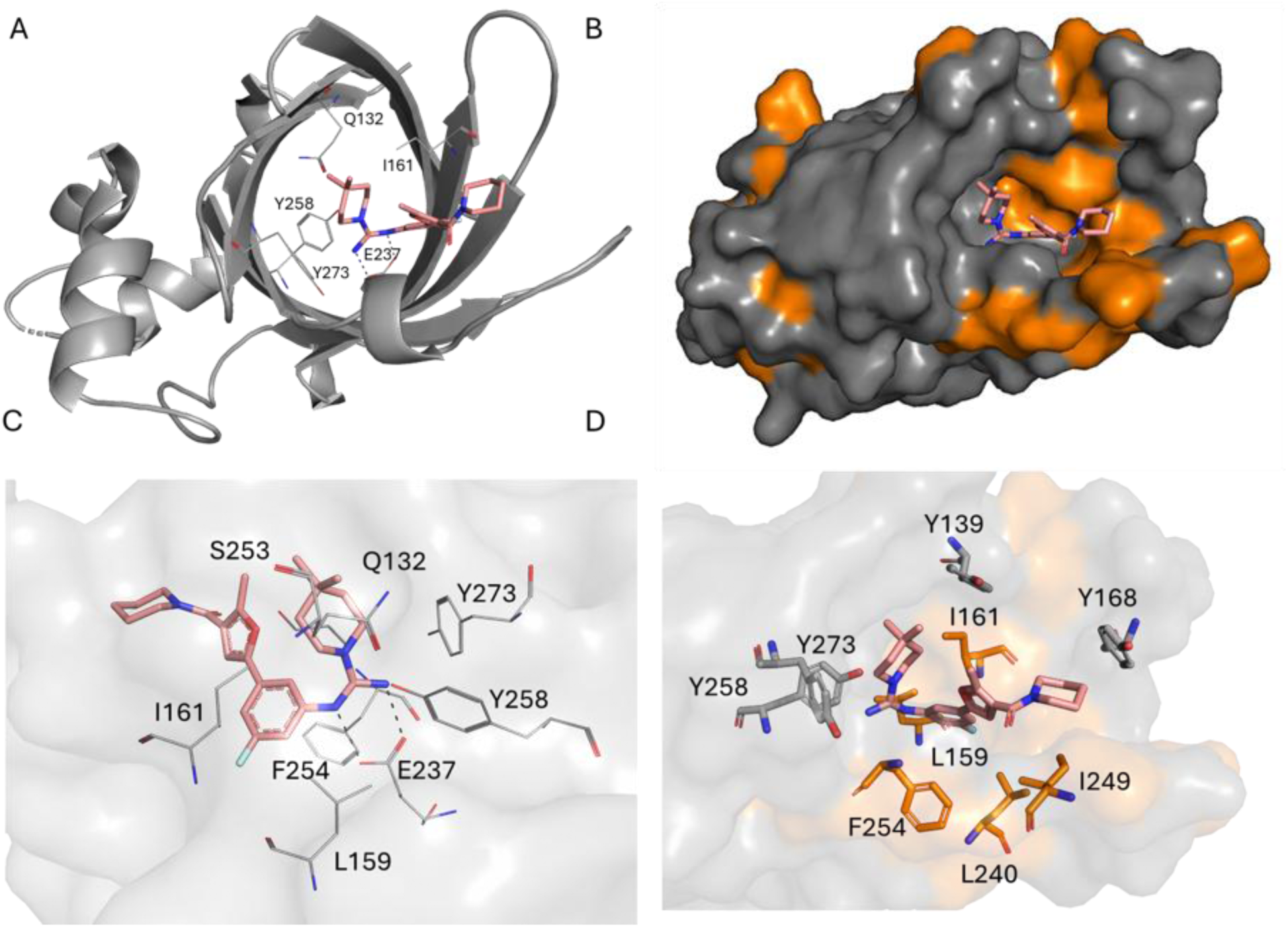
The crystal structure of GID4 protein in complex with compound 14 (PDB: 9QDZ). **A)** Top view of the GID4 binding pocket with compound 14 (salmon) bound in the degron binding pocket, and GID4 represented as cartoon (grey). The N-and C-termini of the protein are labeled as N and C, respectively. **B)** The surface representation of GID4 with all hydrophobic residues colored orange (the rest of the amino acids are grey) with compound 14 (salmon) bound in the GID4 binding pocket. The binding pocket has a hydrophobic character. **C)** Key interactions with Glu237, Ser253 and Tyr273 are labeled for reference. **D)** Close view of the hydrophobic pocket, where the phenyl and furan moieties of compound 14 are surrounded by hydrophobic residues Ile161, Leu171, Leu240, and Phe254.

### Identification of the Exit Vector in GID4 Ligands

Our next objective was to identify a suitable linker attachment point on compound **14** for the synthesis of bifunctional degraders. We explored two potential exit vectors from the piperidine ring (positions 3 and 4) as presented in Table 3. The overall binding affinity of tested compounds **18-23** was determined to be weaker than that of compound **14** with a slight preference for the 3-substituted derivatives (represented by racemic mixtures). Three different substitution patterns *-* methoxy, 2-(2-methoxyethoxy) ethyl, and 1-(4-methylpiperazin-1-yl) ethan-1-one – yielded affinities in the range of K_D_^GID4^=0.05 to 0.14 μM, as measured by SPR. To determine if the linker extensions established additional interactions with the protein, we solved the co-crystal structures of GID4 in complex with compounds **18**, **21,** and **33** (PDB: 9QZG, 9QZI and 9QZH respectively). Unfortunately, in all three structures, the portions of the ligands extending beyond the furan moieties were not supported by electron density (Figures S4-S6), indicating high mobility and orientation toward the solvent-exposed opening of the binding pocket. For compounds **21** and **33**, the linker atoms were modeled with zero occupancy to avoid over-interpreting the structural data. Consequently, the exact trajectories of these linker extensions could not be determined and should be regarded as tentative (Figure S7 and S8). Additionally, the GID4/compound **33** complexes crystallized with four molecules in the asymmetric unit (in contrast to the single chain in other structures), with ligand present in only two of the chains (chain C and chain D). Nevertheless, all tested compounds remained active and proved their utility as building blocks for the bifunctional degraders.

**Table 3.**
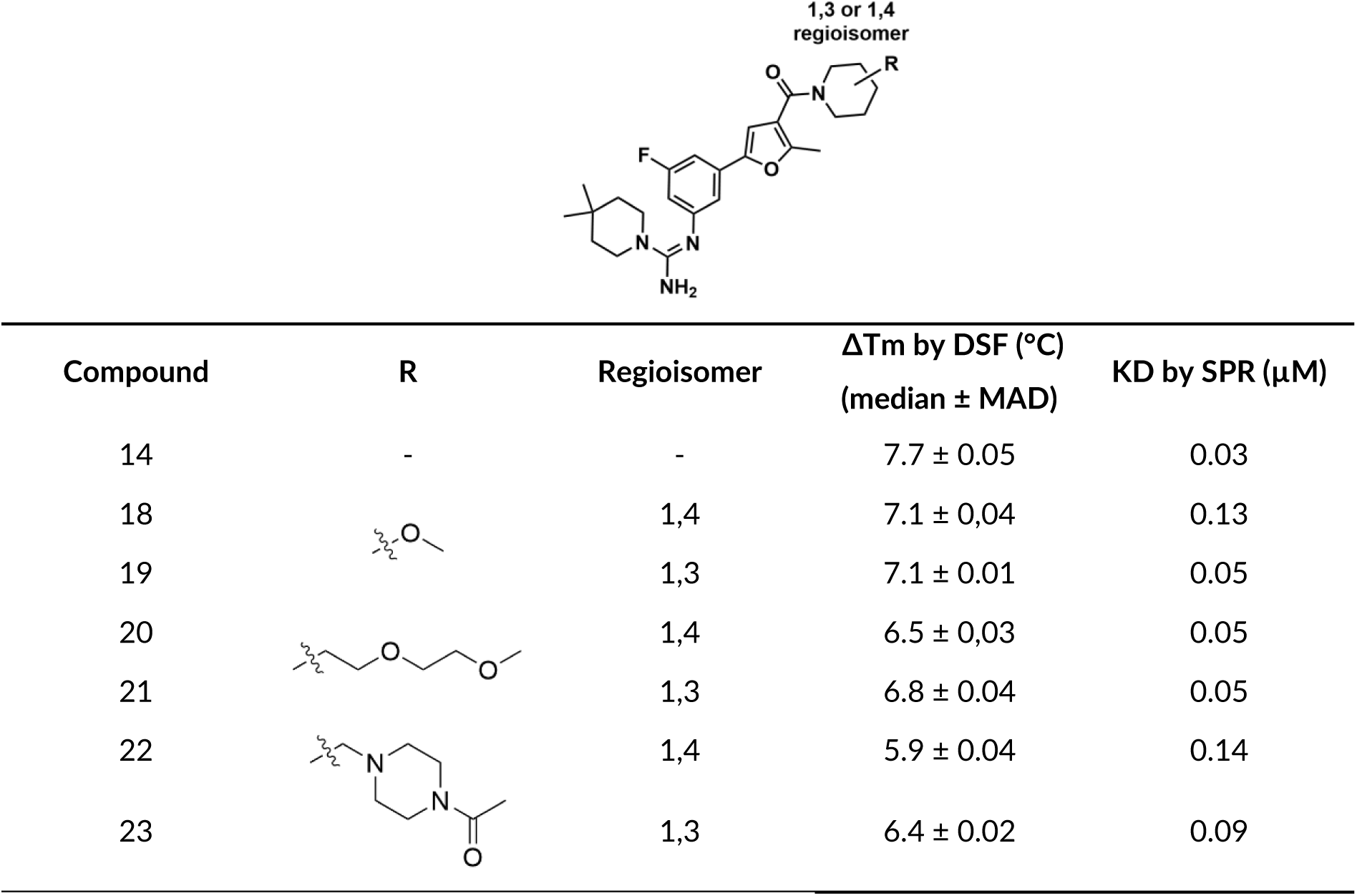
Further modifications of the piperidine moiety in compound 14 and assessment of potential exit vector for bifunctional degraders.

| Compound | R | Regioisomer | $\Delta T_m$ by DSF ($^{\circ}\text{C}$)<br>(median ± MAD) | KD by SPR ($\mu\text{M}$) |
| --- | --- | --- | --- | --- |
| 14 | - | - | $7.7 \pm 0.05$ | 0.03 |
| 18 | | 1,4 | $7.1 \pm 0.04$ | 0.13 |
| 19 | | 1,3 | $7.1 \pm 0.01$ | 0.05 |
| 20 | | 1,4 | $6.5 \pm 0.03$ | 0.05 |
| 21 | | 1,3 | $6.8 \pm 0.04$ | 0.05 |
| 22 | | 1,4 | $5.9 \pm 0.04$ | 0.14 |
| 23 | | 1,3 | $6.4 \pm 0.02$ | 0.09 |

### Intracellular GID4 Engagement Assay

Next, we investigated the cellular activity of the developed GID4 binders. For this purpose, we generated a HEK293T cell line, stably expressing full-length GID4 fused to an N-terminal HiBiT tag. Cells were incubated with either a DMSO vehicle or ligands for 1h, and the luminescence readout was performed while ramping the temperature from 40.4 to 63.5°C. The temperature of aggregation (T_agg_) was defined as the temperature at which 50% of the maximal signal was observed. For the DMSO-treated reference sample, the T_agg(DMSO)_ was established at 54.5 °C. Ligand binding typically stabilizes the protein, resulting in a positive shift in T_agg_ (ΔT_agg_), thereby validating target engagement and suggesting cell membrane permeability. Representative denaturation curves are shown in Figure 5A. While the initial hit compound **1,** did not stabilize the protein in cells, a gradual increase in T_agg_ was observed for compounds **2**, **9**, and **14**. Compound **14** exhibited one of the most significant shifts (ΔT_agg_ = + 3.6 °C), reflecting efficient intracellular target engagement. To confirm the on-target effect of compound **14**, we synthesized a negative control, which maintained similar physicochemical properties but was incapable of interacting with GID4. In this compound, the amine group required for the critical interaction with Glu237 was protected with a ter*t*-butyloxycarbonyl group, which precluded the GID4 binding. As expected, no intracellular target engagement was observed for this modified analog. Figure 5B illustrates the correlation between the thermal stabilization (ΔT_m_) obtained via *in vitro* DSF and the ΔT_agg_ values from the cellular experiment. This strong correlation demonstrates that the higher *in vitro* stability, the more effectively it predicts efficient target engagement in a cellular environment. A few outliers were observed from this trend (Figure 5B, purple squares), which may indicate suboptimal cell membrane permeability – potentially linked to elevated lipophilicity (cLogP) - or compound instability within the cellular medium due to the furan ring. Importantly, all tested analogs with the additional linker fragments (compounds **18-23**) retained activity in the target engagement assay. Higher thermal shift values (ΔT_agg_) were observed for flexible polyethylene glycol (PEG) linkers (compounds **20** and **21**) compared to the more rigid substitutions (compounds **22** and **23**).

**Figure 5.**
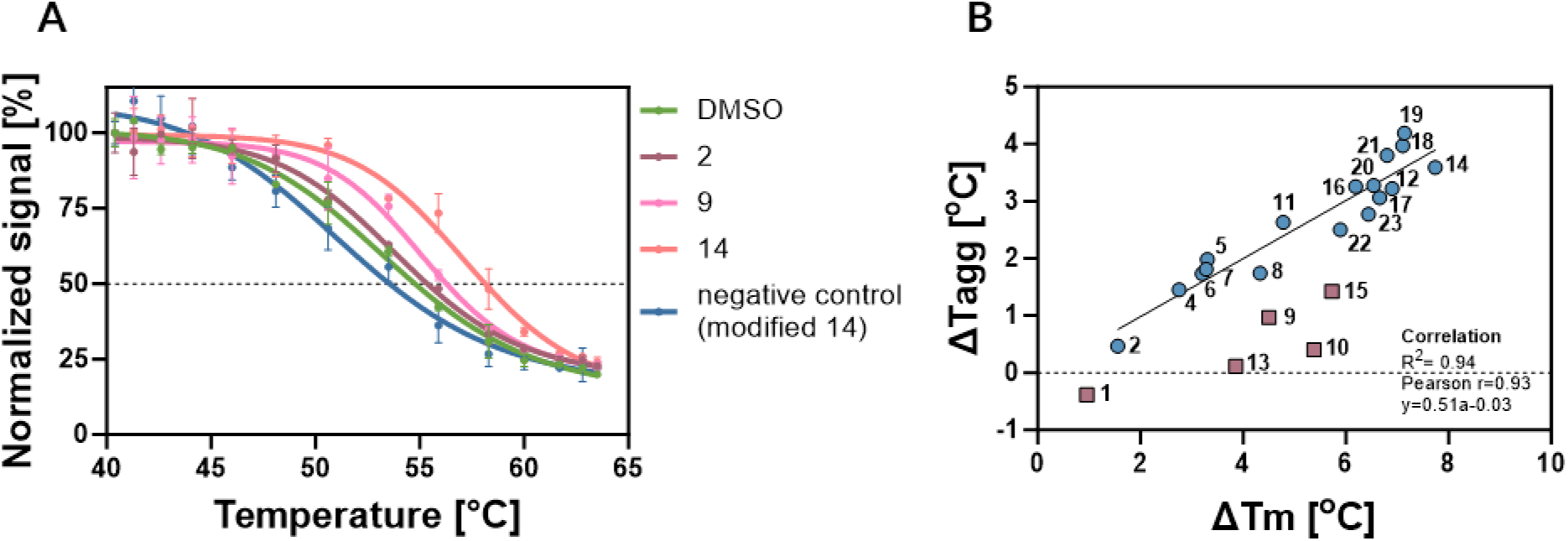
Intracellular target engagement assay. **A)** Denaturation curves obtained from cells treated with selected GID4 binders (compound **2, 9, 14**, and negative control of compound 14). Data presented here are the mean ± s.d. of two independent biological replicates, run in duplicates. **B)** Correlation between ΔT_m_ (DSF) and ΔT_agg_ (intracellular GID4 engagement assay) values obtained for GID4 binders. Compounds are indicated by numbers. Outliers are marked as purple squares and were not included in the calculation of the Pearson correlation coefficient (r).

### Synthesis of GID4-BRD4 Bifunctional Degraders

Following the confirmation of cellular target engagement, we assessed the ability of GID4 to induce targeted protein degradation by synthesizing a panel of GID4-BRD4 bifunctional compounds. We utilized our lead ligand **14** together with a well-known BRD4 ligand JQ1, exploring a range of standard linkers with varying exit vectors and lengths, including rigid alkyl, and flexible PEG (2x, or 4xPEG) (Figure 6A). Compound **26**, featuring the inactive (–)-JQ1 enantiomer, was synthesized as a negative control. We assessed binary interactions with GID4 and BRD4, as well as the ternary complex formation by SPR and HTRF (Figure 6 and Table S2). All bifunctional compounds, except for **26,** retained high affinity for BRD4 (K_D_ ^BRD4^ =0.006-0.02 µM). However, GID4 binding was attenuated relative to the parent GID4 ligand (K_D_ ^GID4^ =0.32-0.63 µM for bifunctional compounds vs. K_D_^GID4^ =0.03 µM for compound **14**). Crucially, all degraders equipped with the active JQ1 warhead mediated ternary complex formation between GID4 and BRD4 (Figure 6 B-C). We further utilized NanoBRET assay to confirm BRD4 engagement, in both intact and lysed HEK293T cells (Table S2). While all compounds (excluding the negative control **26)** engaged BRD4, we observed significantly higher pEC_50_ values in lytic mode (EC_50_=0.05-0.08 µM) compared to live-cell mode (EC_50_=0.40-2.51 µM). This discrepancy indicates potential permeability limitations or active efflux of the bifunctional molecules. Despite successful BRD4 engagement, none of the degraders stabilized GID4 in the intracellular engagement assay, resulting in negative ΔT_agg_ values (Table S2). This was consistent across all compounds, independent of linker length, composition, or exit vector. Consequently, no degradation of BRD4 was observed (Figures S11–S13), highlighting that while ternary complex formation is achievable *in vitro*, the physicochemical properties of these GID4-based PROTACs currently limit their intracellular efficacy.

**Figure 6.**
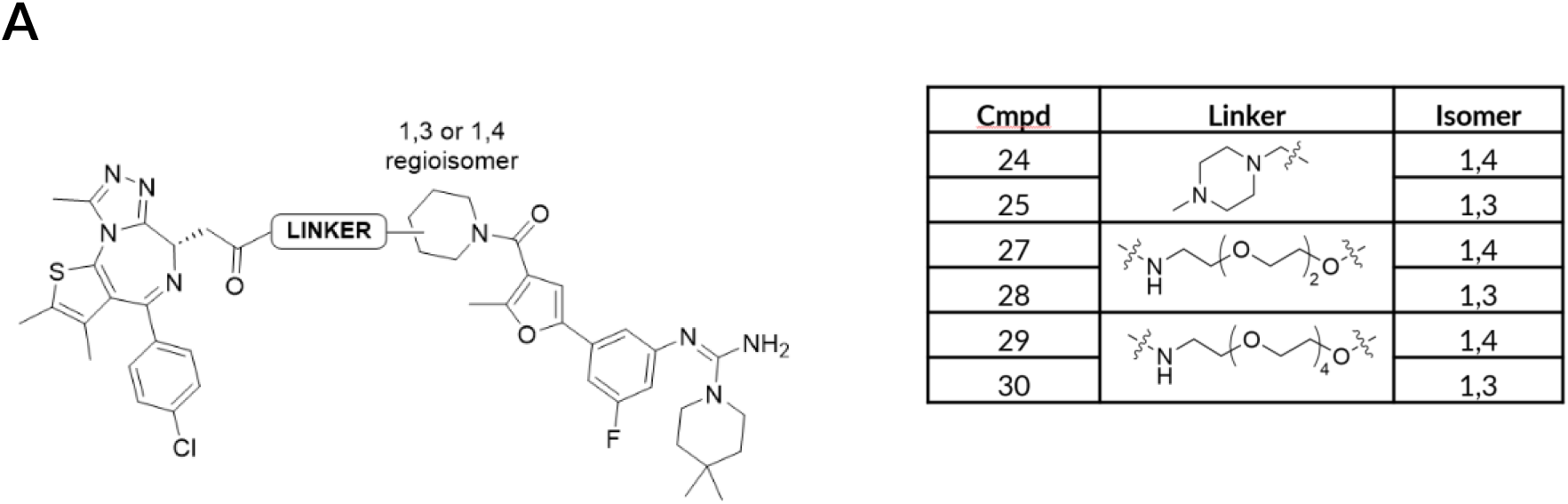
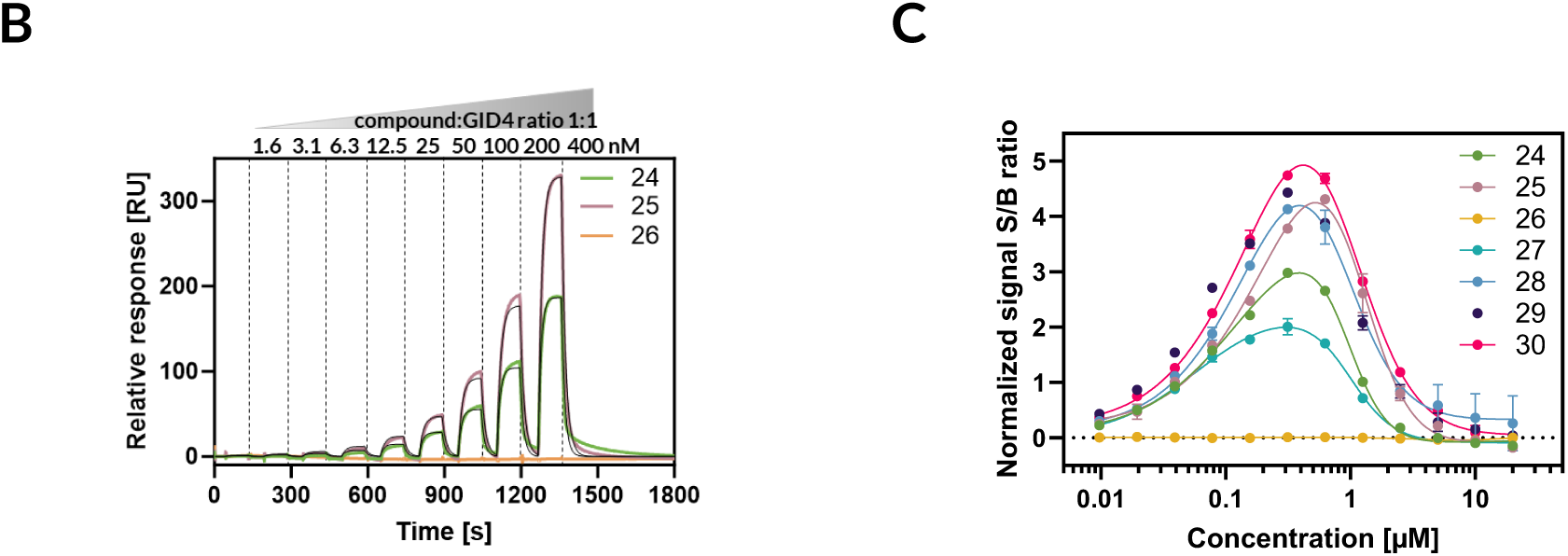
Synthetized GID4-based bifunctional degraders mediate ternary complex formation between GID4 and BRD4. **A)** Structures of synthesized bifunctional degraders and corresponding linkers. **B)** Examples of SPR sensograms of ternary complexes between immobilized BRD4/bifunctional degrader/GID4 for selected compounds. **C)** HTRF assay as an orthogonal method for confirmation of ternary complex formation. Only the negative control 26 does not mediate the formation of the BRD4/GID4 complex.

### Observed Variabilities of GID4 Loops

During the ligand optimization process, we resolved 49 high-resolution X-ray structures of GID4-ligand complexes. Intrigued by the reported conformational flexibility of GID4 we leveraged our X-ray data alongside existing GID4 structures to explore the behavior of the flexible loops (L1-L4) flanking the ligand/peptide binding pocket.^[21,30,31]^ Analysis of these regions revealed significant structural variability, particularly within loops L2-L3, recapitulating the conformational heterogeneity observed between the apo-and holo-states previously described by Dong *et al.*^[21]^ To investigate these variations, we performed clustering to delineate distinct structural patterns. By calculating all pairwise distances between the loop residues, we aimed to capture the intricate movements in L1-L4. To simplify the analysis and facilitate further interpretation, we employed principal component analysis (PCA) to reduce dimensionality. From the PCA results, we extracted the most critical components, PC0 and PC1, which represented the primary sources of structural variability among the ensembles. Interestingly, two key distances, d_0_ and d_1_, emerged from PC0 and PC1: namely d_0_ (T165-S250; L2-L3) and d_1_ (H244-E279; L3-L4). These distances were then utilized as representative measures of structural variability for subsequent analysis and visualization. By focusing on these specific coordinates, we effectively captured the essential features of the loops orientations while minimizing data complexity. Clustering based on distances d_0_ and d_1_ yielded three distinct conformational clusters among the conformers, designated C1, C2, and C3 as illustrated in Figure 7 (see Table S3 for a full list of ligands and clusters association included in this study).

**Figure 7.**
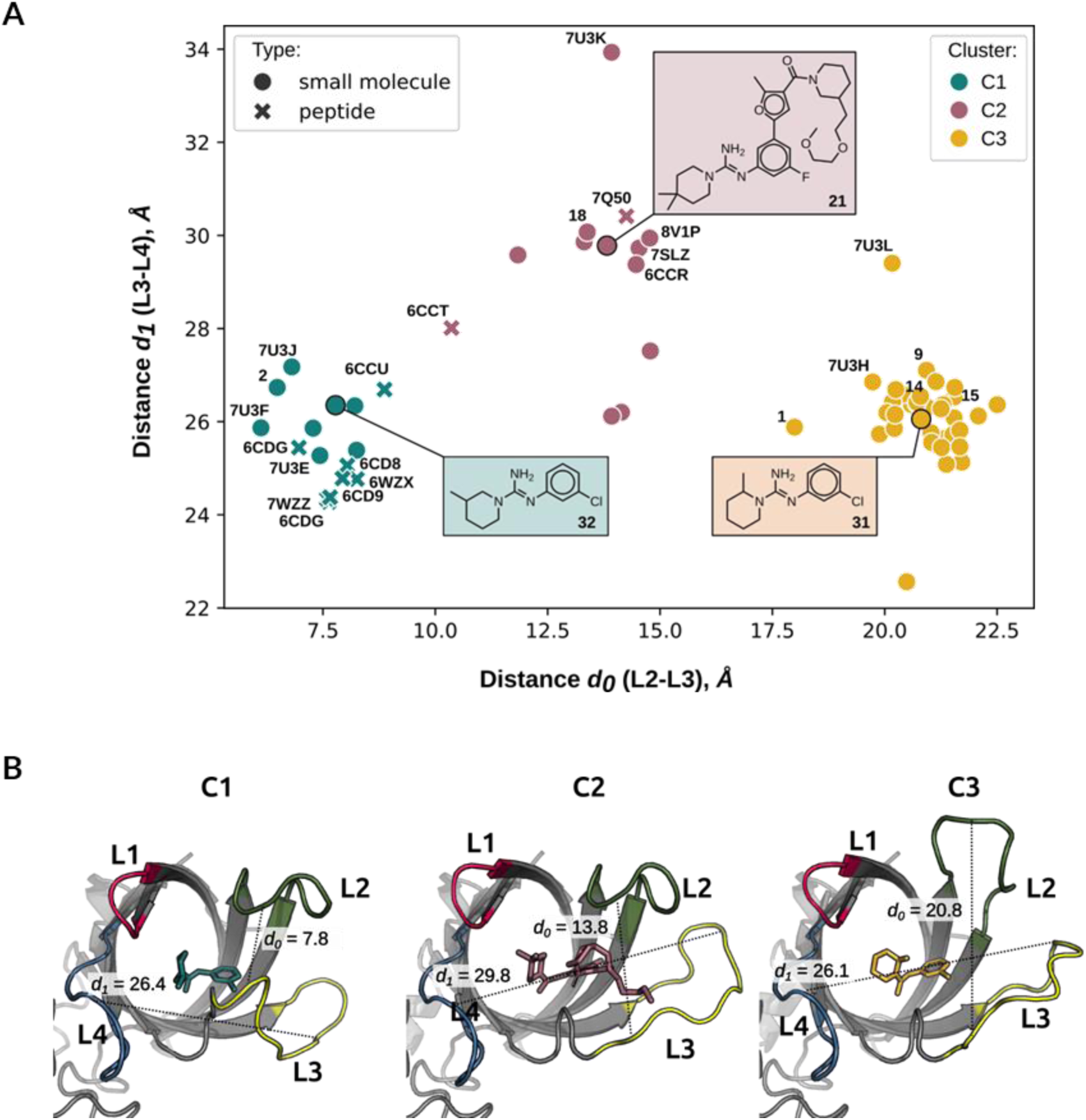
Conformational flexibility of GID4. **A)** GID4 structural variability and ligand-induced states. The plot in the distance coordinates between residues d_0_ (L2-L3; T165-S250) on the x-axis and d_1_ (L3-L4; H244-E279) on the y-axis. Each point in the plot corresponds to the individual protein structure (in-house data or PDB data). The color of the points indicates the cluster assignment (C1, C2, C3). Representatives of each cluster are marked with black edge colors and are presented in B. **B)** Representatives of protein-ligand complexes from each cluster: C1 - closed conformation with compound 32, C2 - semi-open conformation with compound 21, C3 - open conformation with compound 31.

The distinctive structural feature of the clusters was the degree of ‘openness’ of the binding pocket, with a significantly visible difference in the orientation of L2 and L3 (which is also aligned with distances d_0_, d_1_) (Figure 7). Each conformer, derived from a complex with a specific binder, allowed for the examination of structural features associated with its respective cluster. Cluster C1 is characterized by a fully closed loop conformations of L2 and L3 (Figure 7B). Notably, almost all peptide binders belonged to cluster C1, with the only exceptions being the PTLV peptide (PDB: 6CCT), and the 7Q50 - FDVSWFM peptide (PDB: 7Q50).[21,29] In addition, cluster C1 accommodates small molecule ligands (typically ≤20 heavy atoms). The absence of bulky substituents in these ligands appears to promote a closed conformation of loops L2 and L3. In contrast, cluster C2 is characterized by a semi-open state, distinguished by a distinct orientation of L3 (example in Figure 7B). Our analysis of these loop rearrangements, within protein-ligand complexes, suggests that the positioning of aromatic substituents plays a role in semi-open structures of cluster C2. Specifically, the presence of aromatic groups in proximity to L3 may cause rearrangement of the loop. This observation aligns with previous research by Chrustowicz *et al*., who noted the ability of peptide binders, featuring a unique large N-terminal residue (phenylalanine), to induce diverse loop arrangements, *e.g.*, peptide FDVSWFM (PDB: 7Q50).^[30]^ It is worth noting, that while the peptide PTLV (PDB: 6CCT) also resides within cluster C2, it lacks a bulky N-terminal residue; however, its position at the cluster boundary may be an artifact of its lower resolution, which leaves some L3 loop residues unresolved. Interestingly, these findings extended beyond our internal dataset of LMW binders, as small molecules from publicly available structures, *e.g.*, 7SLZ and 8V1P, also feature aromatic groups at analogous positions to the aforementioned N-terminal Phe residue.^[24]^ Among the analyzed structures, only PDB: 7U3K (compound 89 from Chana *et al.*) represents a significant exception, as it features an aliphatic chain occupying the position typically reserved for aromatic groups.^[31]^ This structural divergence is further highlighted by its distance from the cluster centroid (d_0_=13.9, d_1_=33.9, see Figure 7A), underscoring the unique conformational landscape of this specific ligand interaction. Interestingly, the structures in cluster C3 exhibit an open conformation, attributed to movements of both L2 and L3 (Figure 7B). Despite exhaustive efforts, no clear patterns have emerged to explain the fully open conformation observed in this cluster. Unlike clusters C1 and C2, there appears to be a lack of discernible structural elements or binding motifs that universally characterize this group. This intriguing observation suggests that the structural dynamics within this cluster may be driven by a complex interplay of factors that are not yet fully understood. Of note, our lead ligand, compound **14,** and the benchmark compound PFI-7 (PDB: 7SLZ) stabilize GID4 in distinct conformational states: C3 (open) and C2 (semi-open), respectively. This structural differentiation highlights the plasticity of the GID4 binding pocket and demonstrates that high-affinity engagement can be achieved through multiple conformational mechanisms.

## DISCUSSION

Our study has identified compound **14** as a potent GID4 binder, demonstrating robust intracellular target engagement. When compared to the recently revealed chemical probe PFI-7, both ligands exhibit similar affinity for GID4 and comparable molecular properties. However, considering these ligands as potential GID4 warheads for bifunctional degraders, their high molecular weight and lipophilicity place them in a less bioavailable space – closer to VHL ligands than to the compact and highly bioavailable CRBN ligands.^[32]^ Consistent with this, our BRD4 NanoBRET assay revealed higher pEC_50_ values in lytic mode compared to live-cell mode (Table S2), indicating that poor cell membrane permeability or active efflux may limit the intracellular efficacy of the GID4-based bifunctional compounds presented here. Nevertheless, systematic optimization of linker architecture and composition can often mitigate the unfavorable physicochemical properties of an E3 ligase warhead, as demonstrated by recent work with PFI-7. Bifunctional degraders, based on PFI-7 warhead have been shown to successfully induce the proteasomal degradation of BRD4 both *in vitro* and *in vivo*.^[33]^ The authors, however, underline the importance of the linker length as well as the cell-type-specific activity of developed degraders. Introducing additional intramolecular interactions within the degrader molecule may also help to overcome suboptimal molecular properties of a ligand by promoting folding that masks polar regions in a nonpolar environment. Such molecular’chameleons’ can mask polar regions in nonpolar environments - thereby reducing the effective polar surface area (PSA) - to facilitate passive cell membrane permeability.^[34]^ A similar strategy was shown for a VHL-based SMARCA2 degrader, where linker optimization reduced active efflux and achieved acceptable oral bioavailability of F=22%.^[35]^ In the case of compound **14**, despite the confirmed intracellular engagement of BRD4 and the successful mediation of a ternary complex *in vitro* (Table S2), our attempts to recruit GID4 using the bifunctional molecules was unsuccessful. This is likely explained by the >10-fold reduction in GID4 affinity observed for the bifunctional compounds relative to compound **14** alone. We hypothesize that the combination of reduced affinity and poor membrane permeability was sufficient to achieve a lack of efficient engagement of GID4 inside the cell, precluding the degradation of BRD4. This outcome underscores the necessity for further refinement of the chemical structures of GID4-based bifunctional compounds. Future efforts must focus on enhancing cellular permeability - potentially through chameleonic design - and optimizing the linker-warhead interface to maintain the high GID4 affinity. A limitation of our current assessment of degradation potential is the focus on a single target (BRD4) within a single cell line (HEK293), as well as an absence of an assay to evaluate ternary complex formation in a live-cell context. Incorporating such assay in future work could provide valuable insights for optimizing bifunctional compounds with respect to exit vector orientation, linker length and composition, as well as for selecting compatible degradation targets. Further research is required to better understand the mechanisms of GID4-mediated degradation and the regulation of the GID4/CTLH complex. A prominent hallmark of mammalian E3 ligase complexes is the use of molecular mimicry to modulate the access of *bona fide* degrons to the binding pocket.^[25,36]^ A growing body of evidence suggests that GID4 has multiple interaction partners, including those lacking the canonical Pro-N degron motif.^[26,27]^ This fact may be supported by the observed substantial mobility of the GID4 loops—particularly L2–L3—which allows the pocket to accommodate diverse interactors. It has been previously demonstrated that the flexible regions of proteins play an important role in facilitating protein-protein interactions, enabling the recognition of multiple targets at a single molecular surface and that an intrinsic disorder can serve as the structural basis for protein promiscuity.^[37,38]^ Our study significantly expands the known repertoire of ligands capable of inducing distinct GID4 loop conformations. Prior to this work, only a single example of an open-loop structure (cluster C3, PDB: 7UH3), had been reported.^[31]^ In contrast, our investigation has yielded over 30 additional structures exhibiting similar open-loop conformations. In particular, our most potent binder, compound **14**, also belongs to cluster C3, whereas the benchmark compound PFI-7 induces a semi-open loop configuration (C2 cluster).

Interestingly, similar conformational flexibility has been observed for CRBN, where molecular glues inducing a closed conformation are significantly more efficacious in driving target degradation. Inspired by this paradigm, we theoretically examined the potential of GID4 to function through a second modality in TPD: the molecular glue. To date, only a few E3 ligases with molecular glue abilities are known (*e.g.*, CRBN or DCAF15). To measure the probability of C1, C2 and C3 GID4 conformations to support molecular glue-mediated ternary complexes, we plotted pocket volume against the calculated protein-protein interface (PPI) using a small but comparative dataset of structures comprising popular ligases in targeted protein degradation and proteins known to interact with molecular glues (Figure S8).^[39]^ This analysis revealed a distinct grouping of proteins with known molecular glues. For example, CRBN features a mid-size binding pocket and a high PPI area. Moreover, the closed form of CRBN (CRBN^C^) exhibits a higher PPI value than the open form (CRBN^O^). This correlation aligns with findings by Watson *et al* and Yan *et al.*, which suggest that the closed form is essential for the stabilization of ternary complexes and subsequent protein degradation. ^[40,41]^ Conversely, ligases characterized by low PPI and large binding pockets are challenging for the development of molecular glues. While VHL is generally considered suboptimal for molecular glue strategies, a unique case of a molecular glue-induced complex between VHL and CDO1 has been reported.^[42]^ Our exploration of the GID4 centroids (C1, C2 and C3) suggests that the open form of GID4 (cluster C3) is structurally the most similar to ligases capable of forming molecular glue-mediated complexes. Overall, elucidating the structural features and conformational complexity of the GID4 may provide a roadmap for the future design of novel GID4-based molecular glues.

## CONCLUSIONS

This work validates the synergy of fragment-based drug discovery (FBDD) and rigorous structural analysis as a powerful paradigm for targeting E3 ligases. By employing standardized biophysical methods – including SPR and DSF - for hit identification and validation and iterative structure-based optimization, we successfully transformed a modest fragment into a potent GID4 lead (compound **14**). This compound features validated exit vectors and demonstrates robust activity in the intracellular target engagement assay. In addition, our comprehensive analysis of GID4 structural plasticity has revealed a distinct conformational landscape. The distinctive clusterization of the ligand-GID4 complexes might be helpful in better understanding the structure-activity relationship and design of new ligands to achieve desired properties and therapeutic outcomes. Collectively, these findings provide a robust foundation for further exploration into the therapeutic potential of GID4 in targeted protein degradation and molecular glue development.

## MATERIALS AND METHODS

### Recombinant Protein Expression and Purification

#### GID4 Expression and Purification

Human Glucose-induced degradation protein 4 homolog (Gid4, UniProt ID: Q8IVV7) residues 124-289 were cloned into a vector to generate a N-terminal 6xHis-SUMO-tag for crystallography studies, residues 116-300 were cloned into a vector to generate a N-terminal 6xHis-SUMO-tag for DSF studies and residues 116-300 were cloned into a vector to generate a N-terminal Strep-SUMO-3C-6xHis-AVI-tag for SPR experiments. All constructs were verified by sequencing. Protein was expressed by transforming plasmid into chemically competent recombinant *E. coli* BL21(DE3) strain. *E coli* was cultured in LB medium supplemented with 50 µg/mL kanamycin in shaker flasks at 37°C and 160 rpm. Once an optical density at 600 nm (OD_600_) reached 0.5-0.6, protein expression was induced by an addition of 0.1 mM IPTG. Cultures were subsequently incubated overnight at 16°C at 90 rpm before harvesting by centrifugation at 6000 *g* for 20 minutes at 4°C. Cell pellets were stored in –80 °C until purification. To purify protein, pellets were thawed and resuspended in 150 mL of lysis buffer (50 mM Tris/HCl pH 8.0 at 4°C, 300 mM NaCl, 5% glycerol) supplemented with 10 mM β-ME (2-Mercaptoethanol), 1 mM PMSF (Phenylmethanesulfonyl fluoride), 0.5 mM EDTA, cOmplete™protease inhibitors cocktail tablet (Roche) and Pierce^TM^ Universal Nuclease. Samples were subsequently lysed using EmulsiFlex cell disruptor by passing twice through it to maximize cell disintegration at 20,000-25,000 Bar. Afterwards, 0.1% of polyethyleneimine was added. The resulting lysate was centrifuged at 30,000 *g* at 4°C for 30 minutes. Clarified lysates were incubated for 1 hour either with 10 mL of Ni Sepharose 6 FF resin equilibrated in 50 mM Tris/HCl pH 8.0 at 4°C, 300 mM NaCl, 5% glycerol, 10 mM β-ME and 5 mM imidazole for His-tag proteins or with Streptactin®XT 4Flow®High Capacity (IBA Lifesciences) resin equilibrated in 50 mM Tris/HCl pH 8.0 at 4°C, 300 mM NaCl, 10% glycerol, 10 mM β-ME for Strep-tag protein. After incubation the resin was loaded into an empty gravity flow column, the unbound fraction was collected and subsequently the resin was washed with either 10 CV of 50 mM Tris/HCl pH 8.0 at 4°C, 300 mM NaCl, 5% glycerol, 10 mM BME followed by 20 mM imidazole for His-tag proteins or 10 CV of 50 mM Tris/HCl pH 8.0 at 4°C, 300 mM NaCl, 10% glycerol, 10 mM BME for Strep-tag protein. Finally, bound protein was eluted using 30 ml of 50 mM Tris/HCl pH 8.0 at 4°C, 300 mM NaCl, 5% glycerol, 10 mM BME and 300 mM imidazole for His-tag proteins. For Strep-tag protein, the bound fraction was eluted using 30 ml of 50 mM Tris/HCl pH 8.0 at 4°C, 300 mM NaCl, 10% glycerol, 10 mM BME and 5 mM biotin. Protein was cleaved overnight at 4°C using 1:100 (Protease:POI) either Ulp-1 SUMO protease (for crystallography and DSF experiments) or with HRV-3C protease for SPR experiments. Proteins with His-tag in the same time were dialysed to buffer containing 50 mM Tris/HCl pH 8.0 at 4°C, 300 mM NaCl and 10 mM β-ME. Cleaved proteins were further purified by reverse IMAC using HisTrap HP column (Cytvia), where flow through containing cleaved protein was collected, followed by size exclusion chromatography using HiLoad Superdex75 26/600 (Cytvia) equilibrated in 20 mM Tris/HCl pH 7.5 at 4°C, 100 mM NaCl, 0.5 mM TCEP for crystallography or in PBS supplemented with 1 mM DTT for DSF experiments. When needed for SPR experiments sample after cleavage was biotinylated using 0.05 mg/mL BirA enzyme in the buffer supplemented with 150 µM biotin, 5 mM MgCl_2_ and 2 mM ATP. Biotinylation experiments were performed overnight at 4°C. Following biotinylation, protein was subjected to size exclusion chromatography equilibrated in PBS supplemented with 1 mM DTT. Selected fractions were analyzed by SDS-PAGE. Protein purity was evaluated by Coomassie-stained SDS-PAGE. Fractions corresponding to pure GID4 protein were pooled and flash frozen using liquid nitrogen and stored at –80°C.

#### BRD4 Expression and Purification

Human wild type Bromodomain-containing protein 4 (BRD4, UniProt ID: O60885) residues range 444-168 were cloned in a vector producing a N-terminal 6xHisTag and TEV protease cleavage site, and C-terminal AviTag. The construct was verified by sequencing. Protein was expressed in *E. coli* BL21(DE3) in Luria Bertani medium (LB) supplemented with 50 µg/mL kanamycin at 37°C. Once OD_600_ reached 0.5, the temperature was reduced to 16°C, and protein production was induced using 0.1 mM IPTG and incubated overnight. Cells were harvested and cell pellets were stored in –80°C until purification. To purify protein cell pellets were thawed and resuspended in lysis buffer containing 50 mM Tris/HCl pH 8.0 at 4°C, 150 mM NaCl, 5 mM imidazole and supplemented with 1 mM DTT, 0.5 mM EDTA, cOmplete™protease inhibitors cocktail tablet (Roche) and 5U/µL nuclease. Samples were subsequently lysed using EmulsiFlex cell disruptor by passing twice through it to maximize cell disintegration at 20,000-25,000 Bar. Afterwards, 0.1% of polyethyleneimine was added. The resulting lysate was centrifuged at 30,000 *g* at 4°C for 30 minutes. The clarified lysate was filtered and loaded to HisTrap HP affinity column (GE Healthcare) followed by wash with excess buffer and elution with buffer containing 500 mM imidazole. To biotinylate, pooled IMAC fractions containing protein were desalted to buffer 50 mM Tris/HCl pH 8.0 at 4°C, 150 mM NaCl, 1 mM DTT and incubated with BirA biotin ligase at ratio of 1:50 in a buffer supplemented with 5 mM MgCl_2_, 211 µM biotin and 2.64 mM ATP at 4°C overnight. Finally, the protein was subjected to size exclusion chromatography (SEC) using HiLoad 26/600 Superdex 75 pg (GE Healthcare) equilibrated in a buffer containing: 50 mM Tris/HCl pH 8.0 at 4°C, 300 mM NaCl, 1 mM DTT. Fractions containing protein were combined, aliquoted, and snap frozen in liquid nitrogen. Samples were stored at-80 °C until further use.

#### Crystallization and X-Ray Structure Determination

Crystallization of GID4 protein was performed using the sitting drop vapour diffusion method. Protein was thawed and concentrated to ∼4 mg/mL using Amicon Ultra-0.5 Centrifugal Filter Unit with 10 kDa molecular weight cut-off. To co-crystallize protein GID4 with ligands of choice, ligands were prepared in 100% DMSO as 20 mM stocks prior to incubation for 1 hour in ratio 1:20 (v/v; ligand to protein). Subsequently protein/ligand complexes were mixed 1:1 with precipitant solution (conditions either: 0.1 M HEPES pH 7.0-7.5, 0.1 M lithium sulfate, 10-25% PEG4000 or 0.1 M TRIS pH 8.0-8.5, 0.1 M lithium sulfate, 10-25% PEG4000 or 0.1 M Na HEPES pH 7.5; 2.5-10% 2-propanol and 6-20% PEG4000 or 0.1 M HEPES pH 7.0-7.5, 10-25% PEG3350 or 0.1 M TRIS pH 8.0-8.5, 10-25% PEG3350. Plates were incubated at 19°C and were inspected regularly. Crystals appeared within 24 hours in the form of thin needles. Prior to data collection, crystals were harvested in 30% glycerol prepared in GID4 precipitant solution and flash-cooled in liquid nitrogen. X-Ray diffraction data were recorded either at Deutsches Elektronen-Synchrotron DESY (Hamburg, Germany) or at Swiss Light Source (Villigen, Switzerland). Structural data were determined using the CCP4 suite of programs.^[43]^ X-Ray data were first processed at beamline using XDS.^[44]^ The structures were determined using molecular replacement using PHASER and Protein Data Bank (PDB) code 6CD9 or own structures as a search model.^[45]^ Refinement was carried out using REFMAC with model building performed and compound descriptions generated using COOT and AceDRG, respectively.^[46–48]^ Figures were prepared using PyMOL.

#### Differential Scanning Fluorimetry

The effect of the compounds on stabilization of the GID4 recombinant protein (amino acid region 116-300) was investigated with the Differential Scanning Fluorimetry (DSF) assay. First, the GID4 protein was thawed and centrifuged at 18,000 *rcf* at 4°C for 5 min (Centrifuge 5430R, Eppendorf). The protein concentration in the supernatant was measured (BioSpectrometer Basic, Eppendorf) and used to prepare a master mix containing 2.5 μM of GID4 in the assay buffer PBS, pH=7.4, 1 mM DTT. The master mix solution was pipetted into a 384-well PCR plate (4titude®, FrameStar®). The plates were spun down at 1000 *rcf* for 10 sec at room temperature (Centrifuge 5804R, Eppendorf). Subsequently, SYPRO™ Orange Protein Gel Stain (cat. no. S6650, Thermo Scientific) and tested compounds were added to the plate using Echo® 555 liquid handler (Labcyte Inc.). Fragments were tested at the c=500 μM and other compounds at c=100 μM in the final reaction mixture of V=10 μL per well. SYPRO™ Orange was used at the final dilution of 7.5x, and the DMSO content was adjusted to 2%. The plate was spun down at 1000 *rcf* at 22°C for 1 min (Centrifuge 5804R, Eppendorf), shaken by Vibroturbulator (Union Scientific) at level 10 for 1 min, spun down again, and incubated in the darkness for 15 min at room temperature. The readout was done using ViiA™ 7 Real-Time PCR System (Applied Biosystems) with a ramp rate of 0.1°C/s and the temperature range from 25°C to 95°C. The ΔTm and SSMD values were determined by the dedicated in-house script.^[49]]^

#### Determinantion of KD^GID4^ by Surface Plasmon Resonance

The molecular interactions were tested by surface plasmon resonance using Biacore 8K instrument (Cytiva). The immobilization was performed on the SA series S Biosensor chip (Cytiva) in all channels (ch1-8) based on a standard capture protocol with a flow rate of 10 µL/min. The surface of flow cell 2 was conditioned by three injections of regeneration solution (1M NaCl in 50mM NaOH). Subsequently, a solution of a biotinylated GID4 (amino acid region 116-300) at concentration of c=2.5 µg/ml in the immobilization buffer (PBS pH 7.4, 0.05% P-20, 1mM DTT) was injected over the sensor surface for 300 s. The immobilization levels were assessed and oscillated at about 2500 RU in each channel. Subsequently, both flow cells were blocked with 50 µM biotin-PEO3 (Biotium) for 90 s at 10 µL/min. All parts of the flow system except for the sensor chip were washed with 50% isopropanol in 1M NaCl and 50 mM NaOH. For the dose-response experiments, the stock solutions of 20 or 100 mM compounds were loaded onto a 384-well LDV plate (Labcyte Inc.). Compounds were transferred to the 384-well SPR destination plate (781280, Greiner) by the Echo® 555 liquid handler (Labcyte Inc.) to create 2-fold dilution dose-response curves. For fragments, the curves consisted of 6 points with concentrations ranging from c=31.25 to 1000 µM, and for other compounds, the curves consisted of 8 points with concentrations ranging from c=0.19 to 100 µM. Each well of the destination plate, containing 1010 nL of the compound backfilled with DMSO, was topped up with 99 µL of sample buffer (PBS pH 7.4, 0.05% P-20, 1 mM DTT, 1% DMSO) using a MultiFlo FX Dispenser (BioTek). The prepared plates were then centrifuged at 1000 *rcf* for 1 min at room temperature using the 5804R centrifuge (Eppendorf). A dedicated analysis method was configured in the Biacore 8K Control Software (Cytiva), which included an association time of t=60s, a dissociation time of t=120 s, and a flow rate of 30 µL/min. The method also incorporated positive, negative and a carry-over controls and solvent correction consisting of 8 points. The results were analyzed using the dedicated method in Biacore Insight Evaluation Software (Cytiva). In multi-cycle kinetics experiments, steady-state affinity for fragments was determined based on analyte binding early stability point (15 seconds from the beginning of injection), while for other compounds, it was determined based on analyte binding late stability point (5 seconds before the end of injection). In single-cycle kinetics experiments, the equilibrium dissociation constant was determined by using a 1:1 binding model.

#### Determinantion of KD^BRD4^ by Surface Plasmon Resonance

The BRD4 was immobilized on a SA series S Biosensor chip (Cytiva). Firstly, the surface of flow cell 2 was conditioned by three injections of regeneration solution (1M NaCl in 50mM NaOH). Subsequently, the c=5 µg/ml solution of biotinylated BRD4 BD1 (amino acid region 44-168) in a running buffer (PBS pH 7.4, 0.05% P-20, 1mM DTT, 2% DMSO) was injected over the sensor surface for 300 s at a flow rate of 10 µL/min. The immobilization levels were assessed and oscillated at about 1400 RU in each channel. Subsequently, both flow cells were blocked with 50 µM biotin-PEO3 (Biotium) for 120 s at 30 µL/min. All parts of the flow system except for the sensor chip were washed with 50% isopropanol in 1M NaCl and 50 mM NaOH to remove nonspecifically bound biotin species from the microfluidics. Compounds were tested in the single-cycle kinetics mode using contact time 90 s, dissociation time 1800s and flow rate of 10 µL/min. The method contained startup, positive, negative and carry-over controls and solvent correction consisting of 4 points. Injections of running buffer (PBS pH 7.4, 0.05% P-20, 1mM DTT, 2% DMSO) were used as a reference. Compounds were injected at 9 concentrations per cycle in the range from c=0.0015 to 10 µM (3-fold dilution). K_D_ ^BRD4^ was determined in Biacore Insight Evaluation Software (Cytiva) by using 1:1 binding model.

#### Investigation of Ternary Complex Formation by Surface Plasmon Resonance

For investigation of ternary complex formation, the BRD4 protein was immobilized as described above. A dose-response experiment was then performed by using a single-cycle kinetics mode with the following parameters: contact time 90 s, dissociation time 1800s and flow rate 10 µL/min. The method contained solvent correction consisting of 4 points and a wash of the flow system with 50% DMSO to remove potential insolubilities. A reference cycle consisted of 9 injections of the untagged GID4 (amino acid region 116-300) in the range of concentrations of c=1.56-400 nM (2-fold dilution). The reference cycle was followed by the sample cycle where the same range of GID4 concentrations was mixed with the investigated compound in the 1:1 ratio. Each pair of the reference-sample cycles was separated by injection of running buffer serving as blank (PBS pH 7.4, 0.05% P-20, 1mM DTT, 2% DMSO). The data analysis was performed in Biacore Insight Evaluation Software (Cytiva). The sensorgrams from the sample cycle were blank-and reference-subtracted and fitted by using 1:1 binding model to determine the k_off_ and K ^ternary^. The half-life of the ternary complex was given by t_1/2_ = 0.693/k_off_. Cooperativity factor α was calculated as K_D_ ^BRD4^/ K_D_ ^ternary^.

#### Homogeneous Time Resolved Fluorescence

The effect of the bifunctional compounds on the mediation of a ternary complex between GID4 (amino acid region 116-300; 6His-tag Avi-tag) and BRD4 (BD1) (amino acid region 44-168; biotinylated) was investigated using the Homogeneous Time-Resolved Fluorescence assay (HTRF). Firstly, protein solutions were centrifuged at 18000 *rcf* at 4°C for 5 min (Centrifuge 5430R, Eppendorf). The concentration of each protein was measured 3 times (BioSpectrometer Basic, Eppendorf) and the median of these measurements was used to prepare protein solutions. A mix of the GID4 and BRD4 proteins at the concentration of c=100 nM each, 52.8 nM acceptor mAb anti-6His-d2 (61HISDLA, Cisbio) and the 12 nM donor Eu cryptate-labeled streptavidin (610SAKLA, Cisbio) were prepared in PPI Europium detection buffer supplemented with 1 mM DTT. The prepared mix was dispensed at V=10 μl per well on a 384-well low volume white microplate (Greiner, 784075). Subsequently, tested compounds were added to the plate in the range of concentrations from c=0.0097 to 20 µM (2-fold dilution, 12 points) by using Echo® 555 liquid handler (Labcyte Inc.). DMSO was backfilled to all wells, resulting in a final DMSO content of 0.5%. Wells containing only reaction mix and DMSO served as a background. The plate was sealed with a transparent film, and it was shaken by Vibroturbulator (Union Scientific) at level 3 for 30 sec at room temperature. Then, the plate was spun down at 1000 *rcf* for 60 sec at room temperature (Centrifuge 5804R, Eppendorf) to make sure the liquids were at the bottom of the wells. The plate was incubated at room temperature, and the readout was performed after 60 min using PHERAstar FSX Microplate Reader (BMG Labtech) in the time-resolved fluorescence dual emission mode (filters: TR 337 665 620). The data were analyzed using GraphPad Prism 9 and the dedicated script developed in-house to determine the EC_50_ values.

### Chemical Synthesis

#### General Methods

Solvents and reagents were of reagent grade and were used as supplied by the manufacturer. Organic extracts were routinely dried over anhydrous Na_2_SO_4_. Concentration refers to rotary evaporation under reduced pressure. Chromatography refers to flash chromatography using disposable Interchim PF-SiHP columns (4 to 120 g silica, 50 μm particle size) on a Interchim PuriFlash 5.020 or XS 520 Plus PuriFlash automatic purification system. Preparative HPLC was performed on a Waters AutoPurification system fitted with a YMC Triart C18 (250 x 20 mm, 5µ); alternatively Thermo Fisher Scientific Ultimate 3000 system fitted with Hypersil GOLD C18 (250 x 21.2 mm, 12µ) was used; the mobile phase consisted of a mixture of solvents: water (0.1% formic acid or 20 mmol/L NH_4_HCO_3_) and acetonitrile (w/wo 0.1% formic acid). Liquid chromatography-mass spectrometry (LC-MS) analyses were obtained with a Shimadzu model LCMS-2020 mass spectrometer utilizing ES-API ionization fitted with a phenomenex (Kinetex XB-C18, 2.6 μm particle size, 2.1*50 mm dimensions) reverse-phase column at 40 degrees Celsius. UPLC conditions: mobile phase A = 0.1% formic acid in water, mobile phase B = 0.1% formic acid in acetonitrile; flow rate = 0.5 mL/min; compounds were eluted with a gradient of 5% B/A to 95% B/A for 4 minutes. Alternatively, the LC-MS data were obtained with Water Acquity H Class UPLC attached with Waters SQD 2 mass spectrometer utilizing ES-API ionization fitted with a Waters Acquity UPLC BEH C8 (2.1 x 50 mm, 1.7 μm particle size) reverse-phase column at 45 degrees Celsius. UPLC conditions: mobile phase A = 0.05% formic acid in water, mobile phase B = 0.05% formic acid in acetonitrile; flow rate = 0.8 mL/min; compounds were eluted with a gradient of 5% B/A to 95% B/A for 4 minutes. The purity of all the final compounds was shown to be >95% by UPLC−MS or UPLC. ^1^H NMR spectra were obtained with a Bruker Advance 400 MHz or 500 MHz NMR instrument. Unless otherwise indicated, all protons were reported in Methanol-d4 or Dimethyl Sulfoxide-d6 solvent as parts per million (ppm) with respect to residual MeOD (3.31 ppm) or DMSO-d6 (2.50 ppm).

#### Cell Culture

HEK293T cells (DSMZ, cat. ACC 635, RRID: CVCL_0045) were grown in DMEM high glucose with pyruvate (Gibco, cat. 11995065) supplemented with 10% fetal bovine serum (Gibco, cat. 10500-064), 1% Penicillin-Streptomycin (cat. L0022-100, Biowest).

#### Intracellular Target Engagement Assay - GID4

The DNA fragment encoding full length human Glucose-induced degradation protein 4 homolog in fusion with N-terminal HiBiT tag was cloned into a piggyBac vector [HiBit-GID4(FL)] using HiFi DNA assembly reaction. Such vector was used to transfect HEK293T cells. Stable expression in pools of cells was achieved using antibiotic selection (Puromycin, 2 μg/ml). Briefly, stably transfected cells, in serum-free cell culture medium (DMEM GlutaMAX), were seeded into 384-well PCR plates (2×10^3^ cells / 10 μl / well) (Bio-Rad, HSP3865) using the multichannel Finnpipette (Thermo Fisher Scientific). Tested compounds (16 μM final tested concentration, 25 nl) or DMSO vehicle control (0.25 %) were subsequently added using Echo Acoustic Liquid Handler (E550, Labcyte) and incubated for 1h at 37 °C, 5% CO_2_. The plate was sealed and heated at a temperature gradient (40.4 – 63.5°C) for 3.5 minutes using the pre-heated thermal cycler (C1000 Touch with 384-well module, Bio-Rad) and then cooled to 25 °C. PheraSTAR microplate reader (BMG Labtech) and Nano-Glo^®^ HiBiT Lytic Detection system (10 μl, PROMEGA) were used for the signal readout (RLU values, Luminescence). T_agg_ for GID4 in the presence of DSMO was determined at 54.5 °C.

#### NanoBRET – in Cell BRD4 Target Engagement Assay

Commercially available, HEK293 HiBiT-BRD4 cells stably expressing the LgBiT protein (HiBiT-BRD4 KI HEK293(LgBiT) CPM, cat. CS302312, Promega), and NanoBRET® Intracellular TE BET BRD Tracer (cat. N234A, Promega), were used. The assay was run in 384-well plate format (cat. 3574, Corning). Cells were prepared (3.4×10^3^ cells/ml) in OptiMEM without phenol red (cat. 11058-021, Gibco) and they were mixed with the BET Tracer (0.1 mM stock or DMSO vehicle control). For the live mode assay, the Tracer (1 μl/well; final concentration 80 nM in 20 μl reaction) and 18.8 μl/well of cell suspension was used; for the lytic mode, except of the Tracer (1 μl/well, final concentration 30 nM in 20 μl reaction) and the cell suspension (16.8 μl/well), additionally 2 μl/well of digitonin was added. Next, the cell/Tracer mix (19.8 μl/well) was added onto the pre-plated tested compounds (100x concentrated, 0.2 μl/well; 11-point 2-fold dilution; Echo 555 acoustic dispenser, Labcyte). In the live mode assay, the plate was incubated at 37°C, 5% CO_2_ for 2 hours, and then at room temperature for 15 minutes to cool down. In the lytic mode assay, there was no incubation. The complete NanoBRET reagent was prepared according to the manufacturer’s recommendations, appropriately for live and lytic mode, and added to the tested wells (10 μl/well). The plate was incubated for 2 minutes at room temperature and the luminescence and fluorescence readings followed using CLARIOstar Multimode Plate Reader (donor emission RLU – 450nm; acceptor emission RFU – 610nm).

#### Protein Structure Analysis

A dataset of protein structures from the Protein Data Bank (17 structures) and our own structures (43 structures) were used. The structures were retrieved using the BioPython library.^[50]^ To ensure consistency, only structures with resolved residues 136, 137, 138, 163, 164, 165, 166, 167, 168, 169, 240, 241, 242, 243, 244, 245, 246, 250, 275, 276, 277, 278, 279 were retained. In cases where multiple chains were present, the first chain encountered was selected for further analysis. Four critical loops in the GID4 protein (L1 (136-138), L2 (163-169), L3 (240-250), and L4 (275-279)), previously described by Dong *et al.* were used for detailed analysis. Further analysis was performed using a distance matrix based on the alpha carbon (CA) atoms.^[21]^ The distance matrix was standardized using a standard scaler, and PCA was applied to extract the principal components to explore the structural variability among the GID4 protein structures.^[51]^ From the PCA results, two key distances, d0 (T165-S250) and d1 (H244-E279), were extracted as representative measures of structural variability as the major components of PC0 and PC1. Subsequently, K-means clustering was used for clustering; the results were visualized using the seaborn and matplotlib libraries.^[52,53]^ Visualization of the protein-ligand complexes was prepared by PyMol (https://github.com/schrodinger/pymol-open-source).

#### Calculation of PPI Probability and Pocket Volume

We assessed the ligase’s capability to form molecular glue-based ternary complexes using two predictors: pocket volume and PPI probability, predicted as surface patches with per-point PPI likelihoods. The probable PPI was predicted using the PESTO web tool (https://pesto.epfl.ch/) and saved as a PDB file.^[54]^ Binding pocket parameters (volume, druggability, *etc*.) were identified with the fpocket.^[55]^ The *vert.pqr files representing these pockets were converted into PDB format. Pocket definitions and known ligands were combined into a single PDB file. To estimate the PPIs around the binding pocket, we considered only exposed residues. Solvent accessible surface area (calculated using the freeSASA tool) provided information about such exposed residues.^[56]^ A custom Python script was employed to compute the mean PPI within a 9Å radius around the binding pocket, considering residues with an SASA value > 25Å² as exposed. The integrated data were plotted in PPI/volume coordinates.

## ACCESSION CODES

All atomic coordinates and crystallographic results of GID4/small molecule complexes: GID4/1 (PDB: 9QDX), GID4/9 (PDB: 9QDY), GID4/14 (PDB: 9QDZ), GID4/18 (PDB: 9QZG), GID4/21 (PDB: 9QZI) and GID4/33 (PDB: 9QZH) were deposited to Protein Data Bank and are accessible from their websites (i.e. https://www.rcsb.org/).

## Supporting information

Supporting information_Chemical Synthesis

## ACKNOWLEDGEMENTS

The project nr POIR.01.01.01-00-0931/19 “Development of an integrated technology platform in the field of targeted protein degradation and its implementation to the pharmaceutical market” was co-financed by the European Regional Development Fund. The authors would like to acknowledge all members of the CT Team who contributed to this work. We would like to thank all the staff at: Deutsches Elektronen-Synchrotron DESY (Hamburg, Germany) and Swiss Light Source (Villigen, Switzerland) for their support during data collection. The authors would like to extend their gratitude to Donald Coppen for the meticulous proofreading and valuable suggestions and Przemysław Glaza for helpful comments regarding biophysical assays.

## Authors Contributions

Conceptualization of the manuscript and writing: KD, DK, MWP, BW, DS, ZP; Review and edition of the manuscript: MB, TD, MJW, SC; Chemistry conceptualization: SC, AM, ZP, BW, MJW; Chemical synthesis: AK, AM, WW; Protein purification: KP, MS, IHW, DG, KJ; Crystallography: MWP, KG-M, MK, ANS, JW; Cell-based assays: KD, EK, MC-B, TT; Modelling: DS; Biophysical assays: JA, SF, AP; Project coordination: IM, DK; Research supervised by: SC, MB, MJW

## ABBREVIATIONS

TPD: targeted protein degradation
SBDD: Structure-based drug design
UPS: ubiquitin-proteasome system
CTLH: C-terminal to LisH complex
GID4: Glucose-Induced Degradation protein 4 homolog
hGID4: human GID4
HMGCS1: 3-hydroxy-3-methylglutaryl (HMG)-coenzyme A (CoA) synthase 1
FBS: Fragment-Based Screening
DSF: Differential Scanning Fluorimetry
SPR: surface plasmon resonance
c: concentration
PDB: Protein Data Bank
PCA: principal component analysis
LMW: Low Molecular Weight
PPI: protein-protein interface

## ASSISTIVE WRITING TECHNOLOGIES

We used assistive writing technologies such as Grammarly and OpenAI’s ChatGPT during manuscript preparation. These supplementary tools acted as editors, not as drivers of content creation. The listed authors thoroughly reviewed, revised, and selectively implemented suggested edits to ensure accuracy, consistency, and clarity. The responsibility for the paper’s content and quality remains with the authors.

