## Supporting information for "Ligand-induced Conformational Plasticity of the CTLH E3 Ligase Receptor GID4"

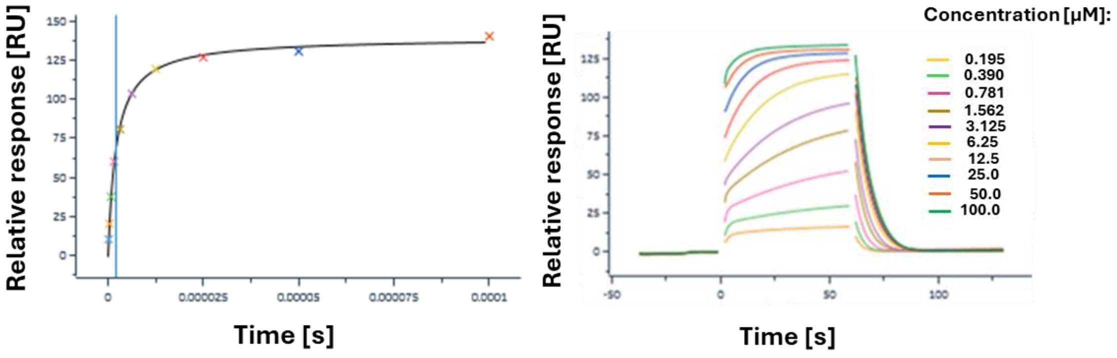


**Figure S1. Confirmation of activity of recombinant GID4 protein with degron peptide PGLWKS.** The $K_D$ was determined by using steady-state affinity binding model ($K_D$=2.1 µM).

**Table S1. X-ray data collection and model refinement statistics.**

| | GID4 + compound 1 | GID4 + compound 9 | GID4 + compound 14 | GID4 + compound 18 | GID4 + compound 21 | GID4 + compound 33 |
|---|---|---|---|---|---|---|
| **Data collection** | SLS | PETRA | PETRA | PETRA | PETRA | PETRA |
| Wavelength, Å | 1.000040 | 1.03318 | 1.0332 | 1.0332 | 1.0333 | 1.0332 |
| Space group | P 1 21 1 | P 1 21 1 | P 1 21 1 | P 21 21 21 | P 21 21 21 | P 1 21 1 |
| Cell dimensions a, b, c (Å) | 38.107 40.568 51.585 | 38.065 40.274 54.394 | 38.090 40.390 52.870 | 40.255 41.752 103.205 | 40.273 41.930 103.135 | 80.0 44.46 103.1 |
| α, β, γ (°) | 90.00 107.943 90.00 | 90.00 110.38 90.00 | 90.00 108.691 90.00 | 90.000 90.000 90.000 | 90.000 90.000 90.000 | 90.0 112.8 90.0 |
| Resolution, Å | 2.26 | 1.79 | 1.8 | 1.90 | 2.0 | 2.16 |
| Resolution Range, Å | 49.08-2.26 | 40.27-1.78 | 36.08-1.80 | 38.73-1.82 | 41.93-2.05 | 47.52-2.16 |
| $R_{meas}$ | 0.170 (0.970) | 0.554 (15.437) | 0.076 (0.709) | 0.475 (8.027) | 0.217 (2.427) | 0.171 (1.570) |
| *I/б* | 5.0 (1.1) | 3.89 (0.1) | 11.9 (2.2) | 6.1 (0.8) | 7.7 (1.2) | 7.6 (1.1) |
| Completeness, % | 99.7 (97.2) | 96.3 (77.6) | 97.3 (94.6) | 97.7 (92.1) | 99.8 (99.9) | |
| Reflections total/unique | 7112/631 | 14211/653 | 53988/14079 | 134697 / 16702 | 89687 / 12366 | 140534 / 35845 |
| Multiplicity | 1.9 | 4.3 | 3.8 | 8.1 | 7.3 | 3.9 |
| **Refinement** | | | | | | |
| $R_{work/free}$ | 0.21/0.26 | 0.257/0.32 | 0.18/0.212 | 0.28/ 0.29 | 0.20/0.26 | 0.282/0.33 |
| Residues/Atoms | | | | | | |

| | | | | | | |
|---|---|---|---|---|---|---|
| Protein | 36.1/2588 | 29.0/2514 | 30.1/2575 | 34.7/2553 | 39.1/2551 | Chain A: 33.5/2531<br>Chain B: 39.9/2544<br>Chain C: 38.0/2474<br>Chain D: 31.9/2483 |
| Ligand | 32.7 / 32 | 32.5 / 42 | 31.0 /62 | 38.7 / 66 | 77.8 / 79 | Chain C: 41.6/ 69<br>Chain D: 35.3/ 69 |
| Water | 27.4/ 12 | 23.4/ 15 | 34.8 / 48 | 25.9 / 8 | 36.1 / 19 | |
| R.M.S. deviations | | | | | | |
| Bond lengths, Å | 0.0065 | 0.0146 | 0.0100 | 0.0106 | 0.0147 | 0.0116 |
| Bond angles, ° | 1.601 | 2.373 | 1.679 | 1.905 | 2.348 | 2.234 |
| **PDB ID code** | 9QDX | 9QDY | 9QDZ | 9QZG | 9QZI | 9QZH |

Values in parentheses are for the highest resolution shell.

| | | | |
|---|---|---|---|
| A | | | |
| PDB: 6CD9 | GID4/Compound 1 | GID4/Compound 9 | GID4/Compound 14 |
| Degron peptide, aa:<br>PSRW | 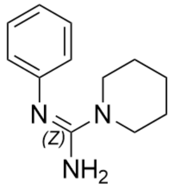 | 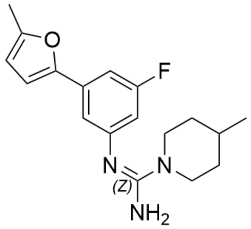 | 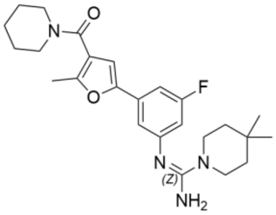 |
| B | | | |
| 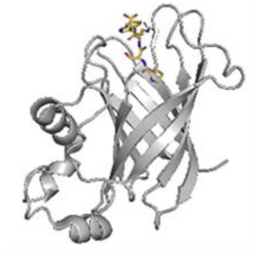 | 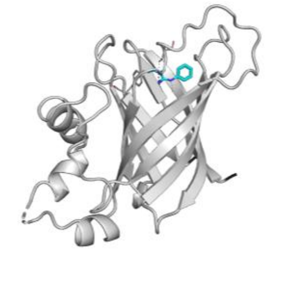 | 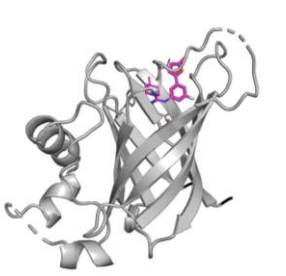 | 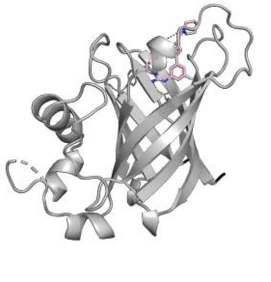 |
| C | | | |
| 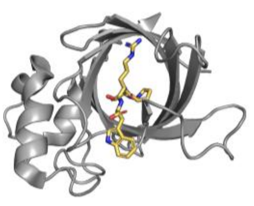 | 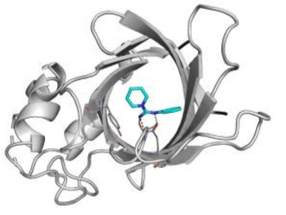 | 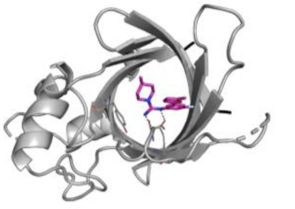 | 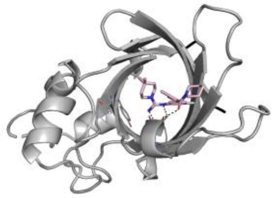 |
| D | | | |
| 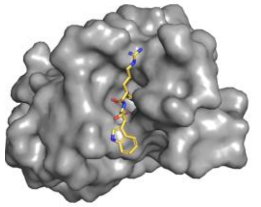 | 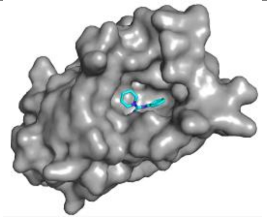 | 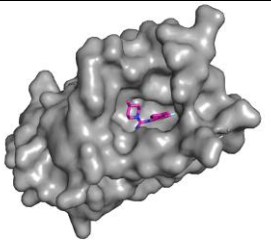 | 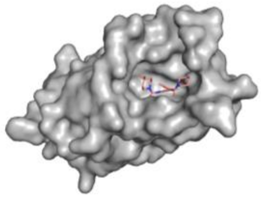 |
| E | | | |

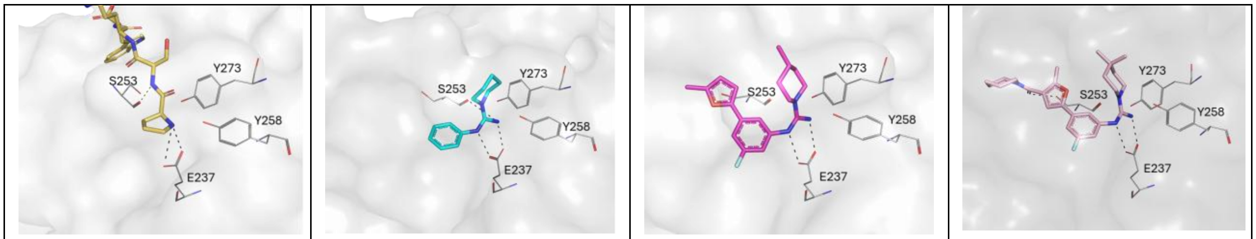


F

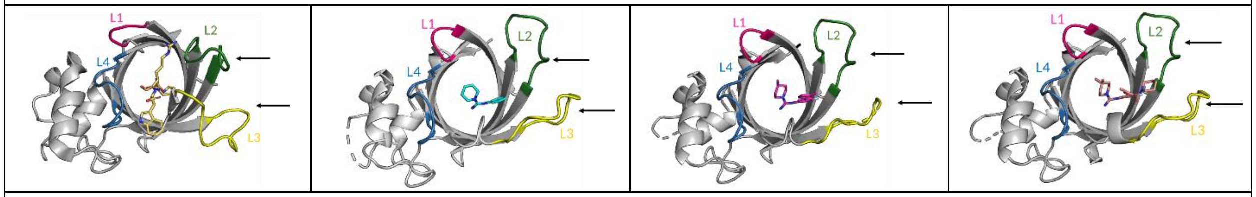


G

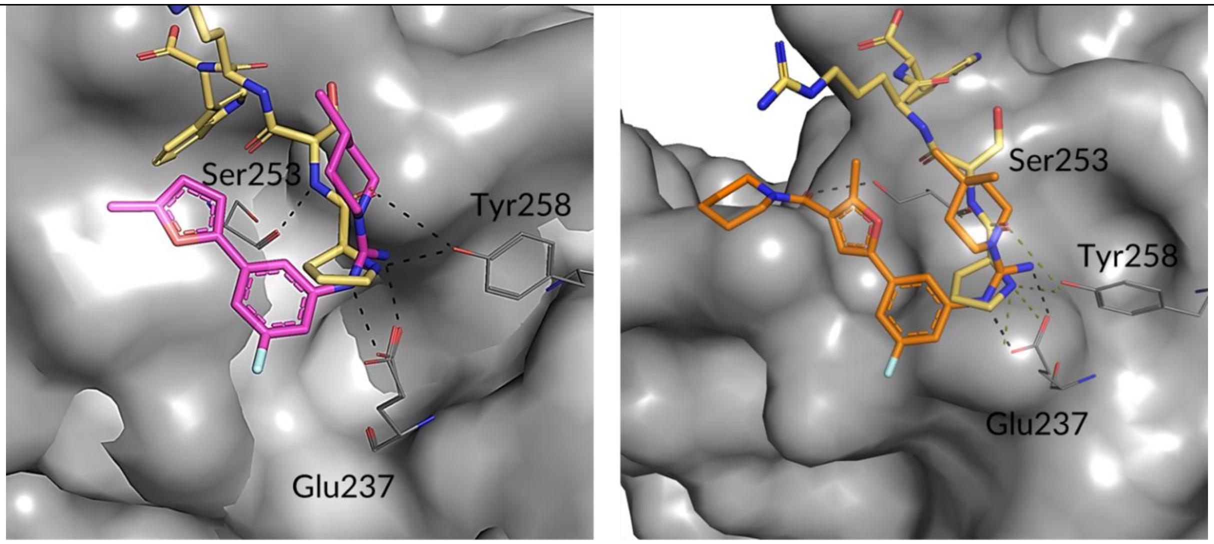


PDB: 6CD9 and Compound 9

PDB: 6CD9 and Compound 14

H

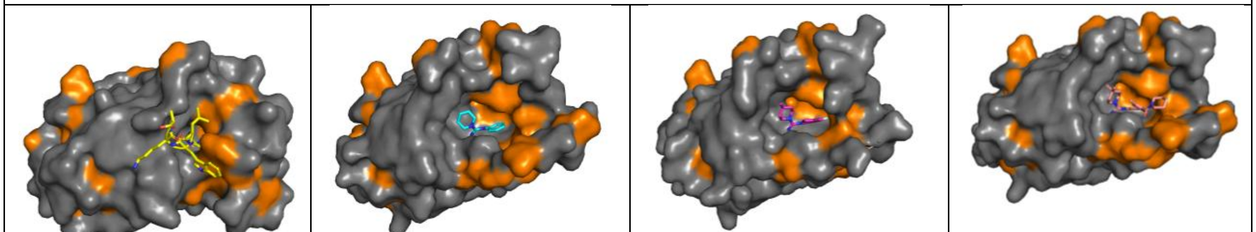

I

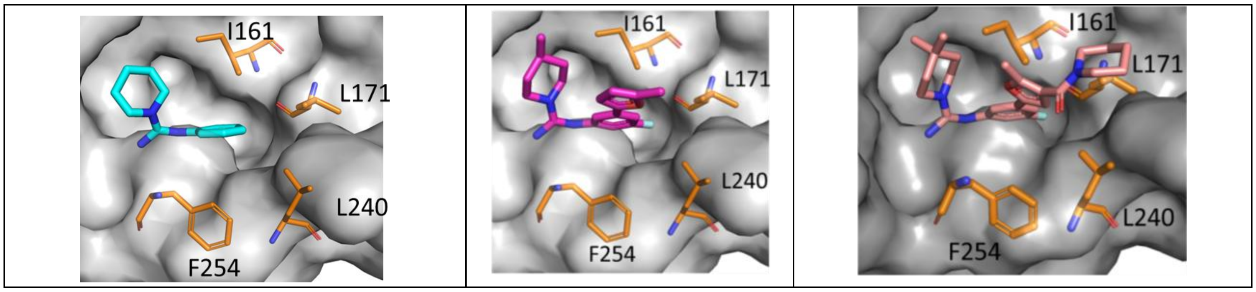


**Figure S2. Co-crystal structures of GID4 with ligands: compound 1 (turquoise), compound 9 (magenta), and compound 14 (salmon) in comparison to published structure of GID4 with a tetrapeptide PSRW; PDB: 6CD9 (straw). A)** 2D structures of co-crystallized compounds, starting from the left: degron peptide amino acid composition, compound 1, compound 9, and compound 14. **B)** Side view of the crystal structure of the GID4 barrel, shown as cartoon representation with a ligand bound in a binding pocket on top of the barrel; starting from the left: degron peptide amino acid composition, compound 1, compound 9, and compound 14. **C)** Top view on the binding pocket of GID4 as cartoon representation with the ligand present; starting from the left: degron peptide amino acid composition, compound 1, compound 9, and compound 14. **D)** Surface view of the GID4 binding pocket with ligands bound; starting from the left: degron peptide amino acid composition, compound 1, compound 9, and compound 14. **E)** Close view of the key interactions between GID4 and each hit fragment; starting from the left: degron peptide amino acid composition, compound 1, compound 9, and compound 14. **F)** Comparison of positions of the GID4 hairpin loops (L1 in magenta, L2 in green, L3 in yellow, and L4 in blue), in three hit fragment structures and in degron-peptide bound form. Arrows direct towards loops L2 and L3, which differ the most between the peptide-bound form and a small molecule ligand-bound form. **G)** Superposition of crystal structures of GID4 in complex with peptide PSRW (PDB: 6CD9; straw) and compound 9 (magenta) or compound 14 (salmon). The key interacting residues are labeled. **H)** Hydrophobic amino acids of GID4 protein (orange) form a degron peptide and a ligand binding pocket; starting from the left: degron peptide amino acid composition, compound 1, compound 9, and compound 14. **I)** Hydrophobic residues (orange) are involved in the binding of the phenyl and furan moiety of ligands; starting with left: compound 1, compound 9, and compound 14.

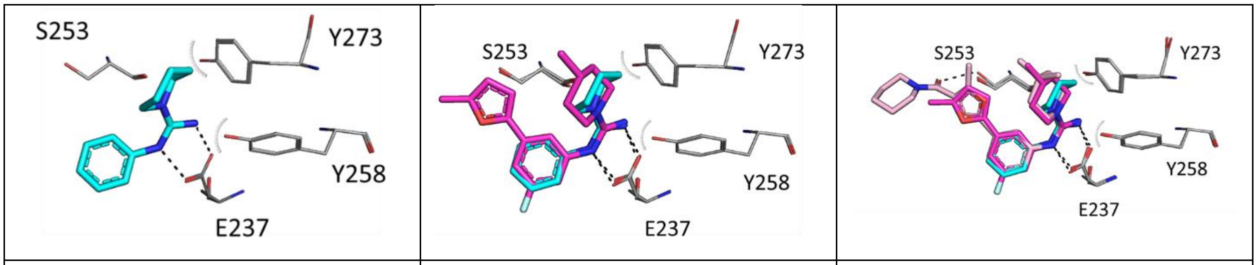


| Compound 1 | Compound 9 | Compound 14 |
|---|---|---|

**Figure S3. GID4 ligand evolution - from compound 1, via compound 9 to compound 14.**

**Table S2. Parameters of the GID4-BRD4 ternary complexes as well as BRD4 and GID4 intracellular target engagement mediated by the GID4-based bifunctional degraders.**

| Bifunctional degrader | KD binary GID4 | KD binary BRD4 | KD ternary | Cooperati-vity α | t ½ [s] | HTRF $EC_{50}$ | BRD4 NanoBRET $IC_{50}$ [μM] (live) (n=2, ±s.d.) | BRD4 NanoBRET $IC_{50}$ [μM] (lytic) (n=2, ±s.d.) | GID4 intracellular target engagement $\Delta T_{agg}$ [°C] (n=2, ±s.d.) |
|---|---|---|---|---|---|---|---|---|---|
| 24 | 0.32 | 0.013 | 1.58 | <1 | 9.2 | 0.06 | - | - | -1.7 ± 1.2 |
| 25 | 0.50 | 0.020 | 2.51 | <1 | 10.6 | 0.10 | - | - | -1.4 ± 0.3 |
| 26 | 0.63 | inactive | inactive | - | - | inactive | - | - | -2 ± 0.6 |
| 27 | - | 0.008 | - | - | - | 0.05 | 0.36 ± 0.01 | $(5.25 \pm 0.035) \times 10^{-2}$ | -0.8 ± 1.8 |
| 28 | - | 0.016 | - | - | - | 0.08 | - | - | -2.6 ± 3.7 |
| 29 | - | 0.006 | - | - | - | 0.05 | 1.74 ± 0.38 | 0.056 ± 0.007 | 0.1 ± 0.4 |
| 30 | - | 0.006 | 0.06 | <1 | 28.8 | 0.08 | 2.63 ± 0.3 | 0.083 ±0.01 | -0.3 ± 1.0 |

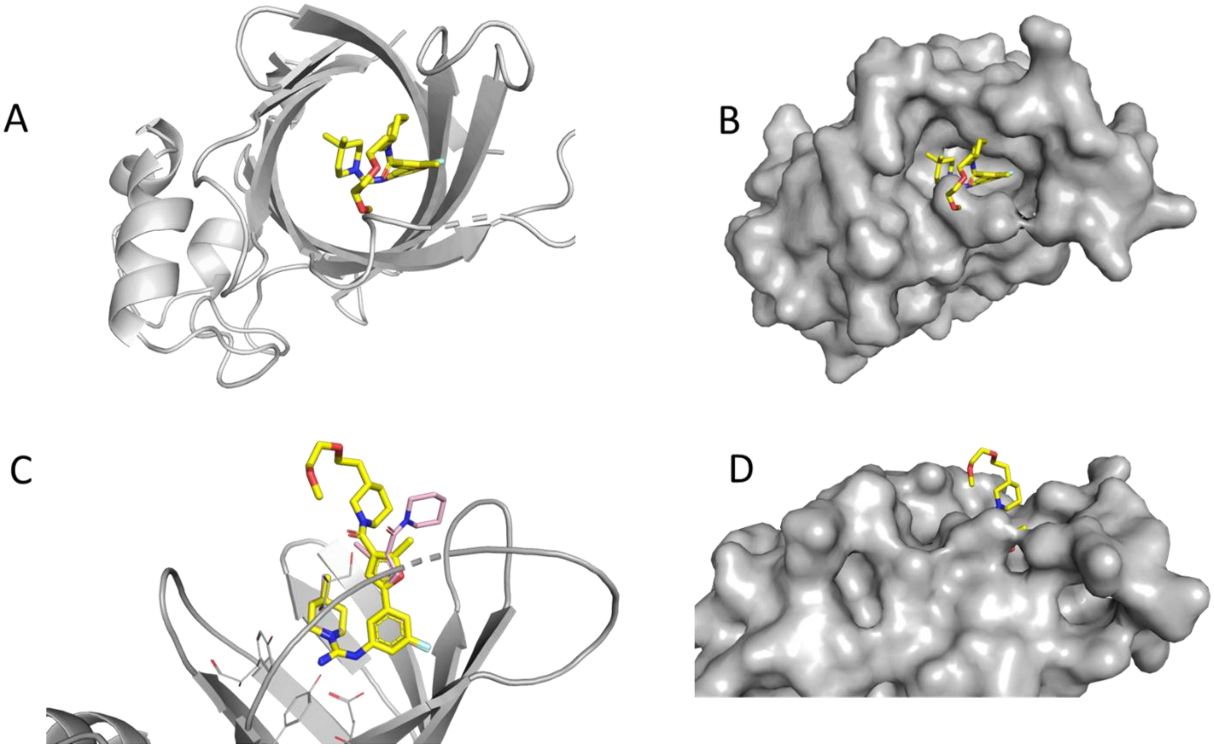


**Figure S4**. **The crystal structure of GID4 in complex with compound 21. A)** Top view of the GID4 binding pocket with compound 21 (yellow) bound in the degron binding pocket, GID4 represented as cartoon (grey). **B)** The surface representation of GID4 with compound 21 (yellow) bound in the GID4 binding pocket. **C)** Comparison of compound 21 (yellow) with compound 14 (salmon). Their furan moieties superpose well. Ligand parts beyond furan moieties were not visible in electron density as they were mobile and the point towards solvent and outside of the binding pocket. **D)** Piperidine ring moieties point in different directions in compound 21 and compound 14; however, their exact positions were not determined, so they should be considered approximate.

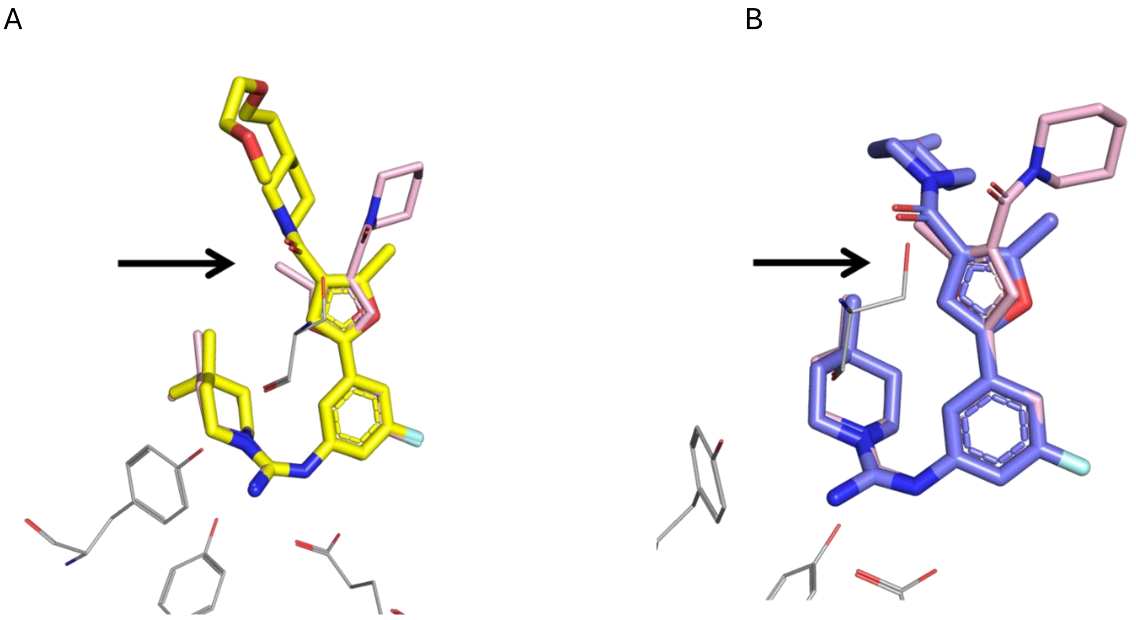


**Figure S5**. **Comparison of compound 21 (yellow) A) or compound 18 (blue) B) with compound 14 (pink).** All compounds' furan moieties superpose well. The ligand portions beyond the furan moieties were not visible in electron density because they were mobile, pointing towards the solvent and outside the binding pocket. The piperidine ring moieties point in different directions in compound 21 and compound 18 compared to compound 14, but their exact positions were not determined, so they should be considered approximate. From the region marked by the black arrow towards the top, the compound position is not defined.

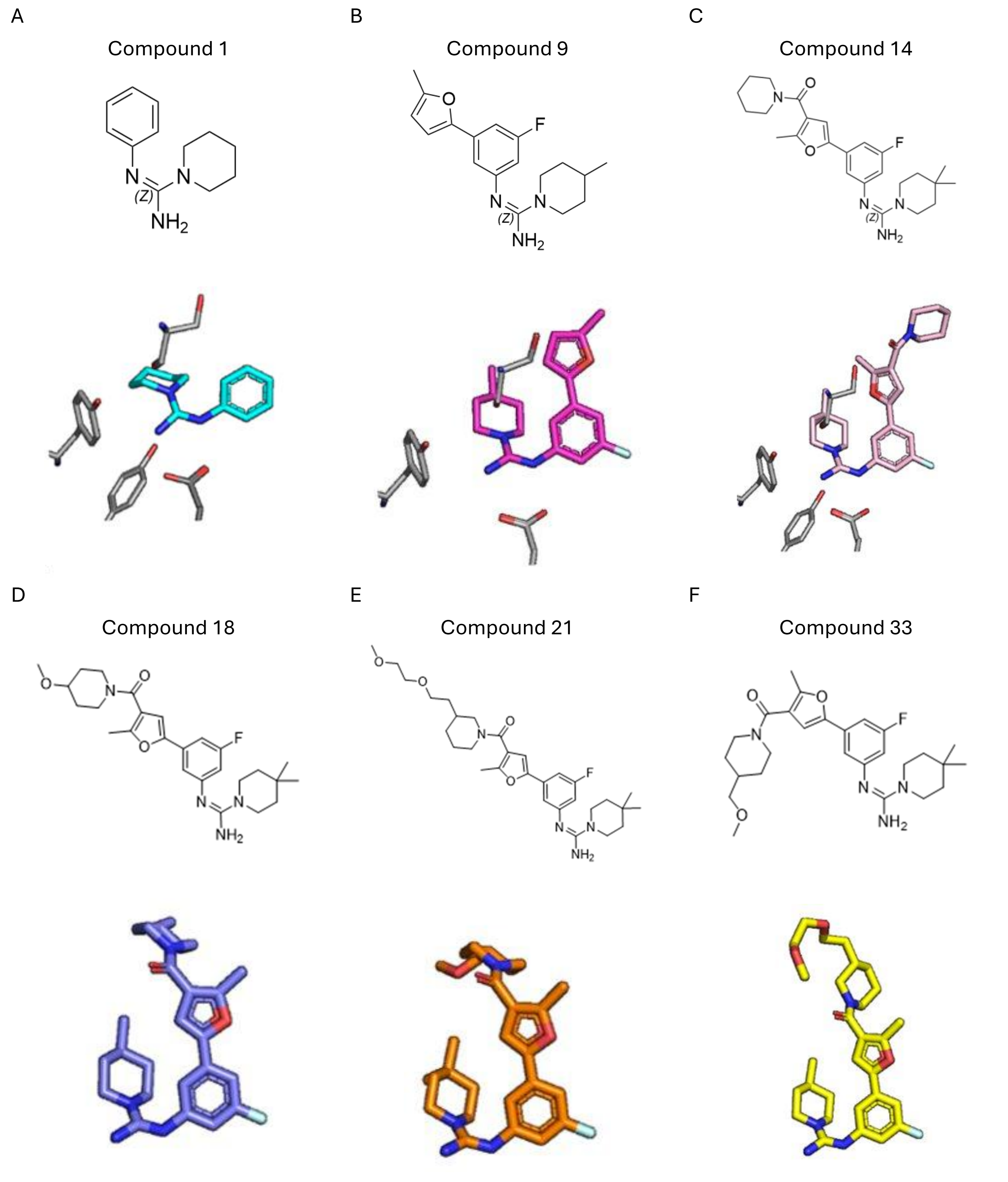


**Figure S6**. **Comparison of GID4 compounds superposed as bound in the binding pocket. A)** Compound 1 (turquoise); **B)** Compound 9 (magenta); **C)** Compound 14 (salmon); **D)** Compound 18 (blue); **E)** Compound 33 (orange); **F)** Compound 21 (yellow)**.** All compounds' furan moieties superimpose well. Ligand portions beyond the furan moieties were not visible in electron density because they were mobile and pointed towards the solvent and outside the binding pocket. The piperidine ring moieties point in different directions in compound 21 F), compound 33 E) and compound 18 D) in comparison to compound 14 (C) but their exact positions were not determined, so they should be considered approximate.

A B

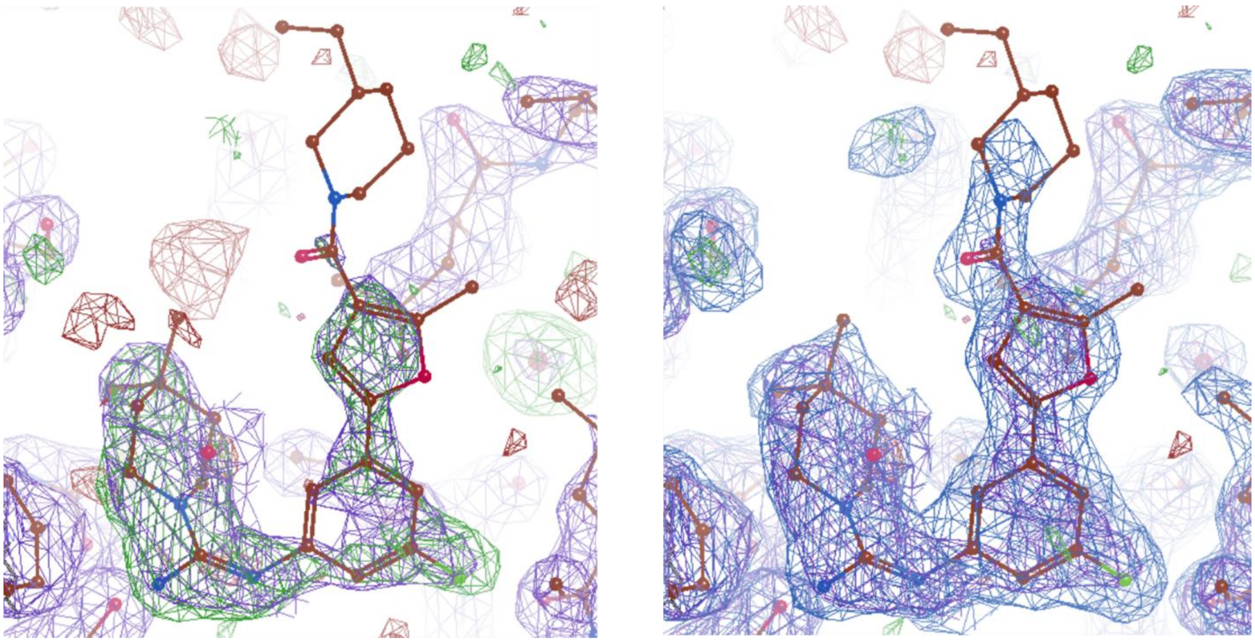

**Figure S7. Electron density map of compound 21. A)** Omit the electron density map before building compound 21, which shows the region of the binding pocket, where the compound is bound (green); 2 Å, 2Fo-F, σ=1. **B)** The electron density map (blue) after building compound 21. 2 Å, 2Fo-F, σ=1

A B

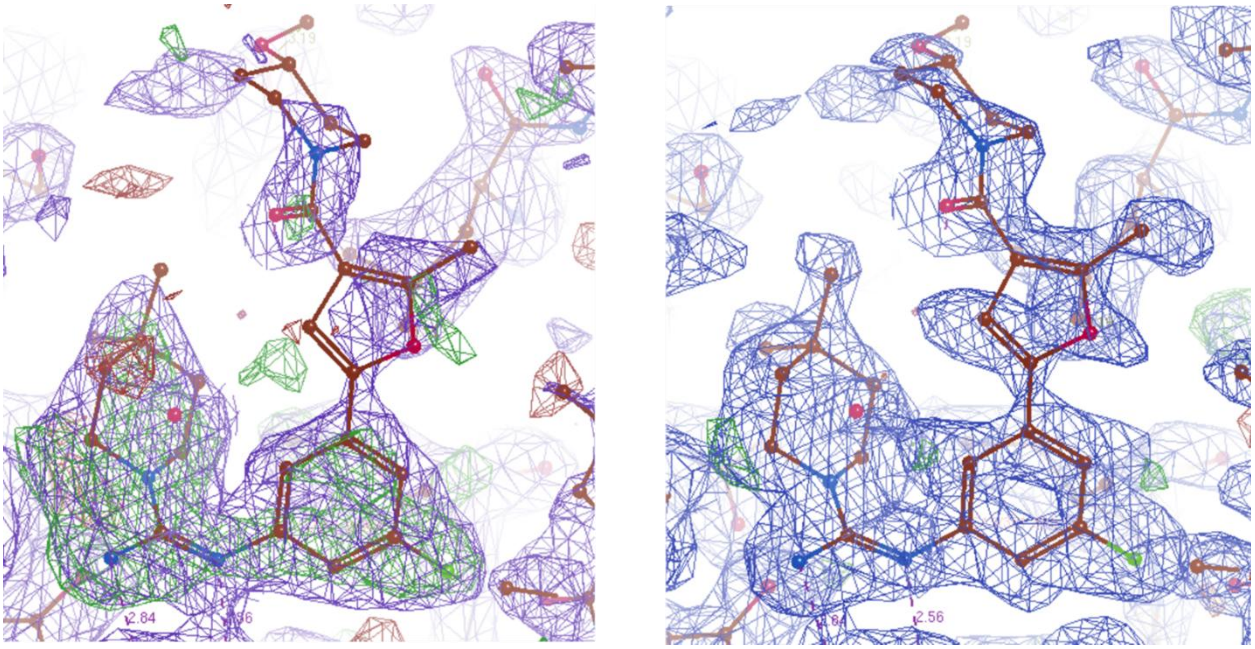


**Figure S8. Electron density map for compound 18. A)** Omit the electron density map before building compound 18, which shows the region of the binding pocket, where the compound is bound (green); 1.9 Å, 2Fo-F, σ=1. **B)** Electron density map (blue) after building compound 21. 1.9 Å, 2Fo-F, σ=1.

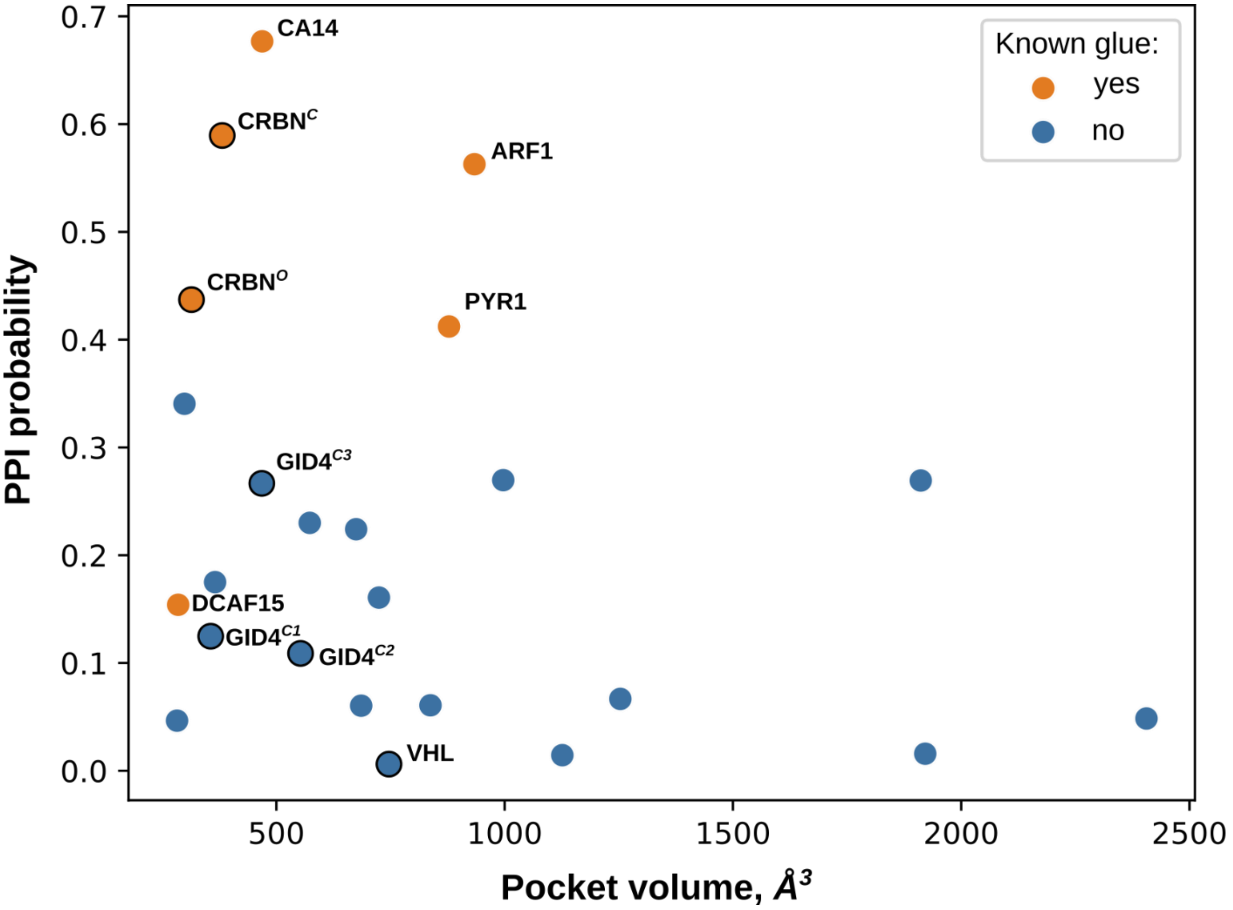


**Figure S9. Visualization of the probability of protein-protein interface (PPI probability) around the binding pocket and the binding pocket volume.** The color represents molecular glue interaction: orange indicates protein with known molecular glues (summarized in Rui et al.), while blue represents proteins with no confirmed molecular glues.[1] **The typical TBD ligases are labeled: VHL, CRBN (closed structure CRBN$^{C}$** and open structure CRBN$^{O}$), alongside GID4 structures (GID4$^{C1}$ from cluster C1, GID4$^{C2}$ from cluster C2 and GID4$^{C3}$ from cluster C3). The higher PPI values of ligases like CRBN suggest a propensity for interaction with molecular glues, while ligases with lower PPI values, such as VHL, are less conducive to such interactions.

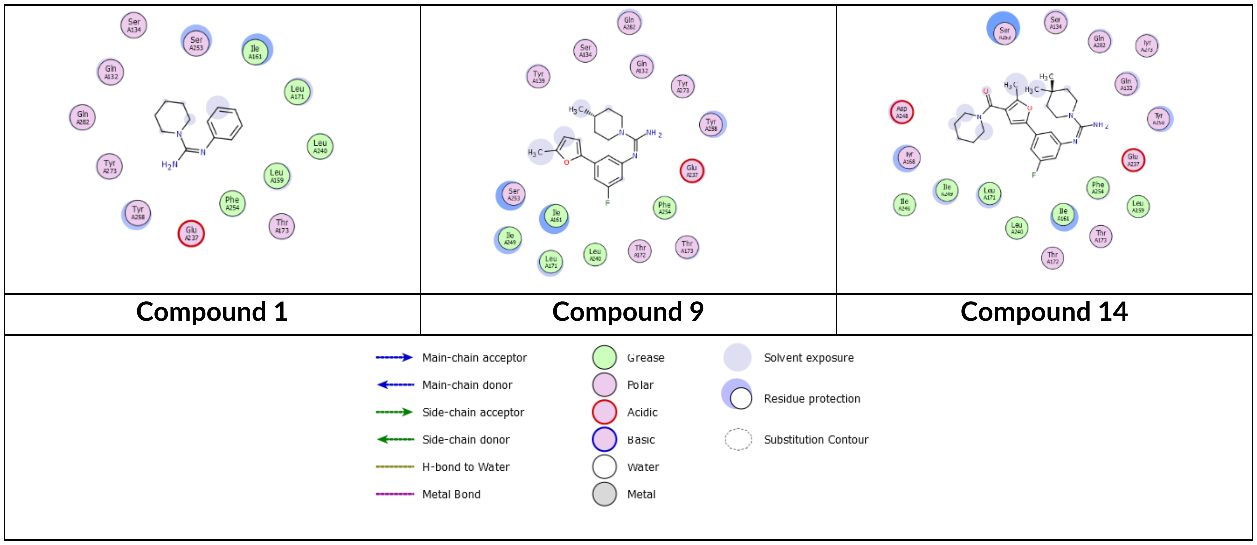


**Figure S10. FLEV for compounds 1, 9, and 14 generated using Coot. (FLEV = Flatland Ligand Environment View).** [2,3]

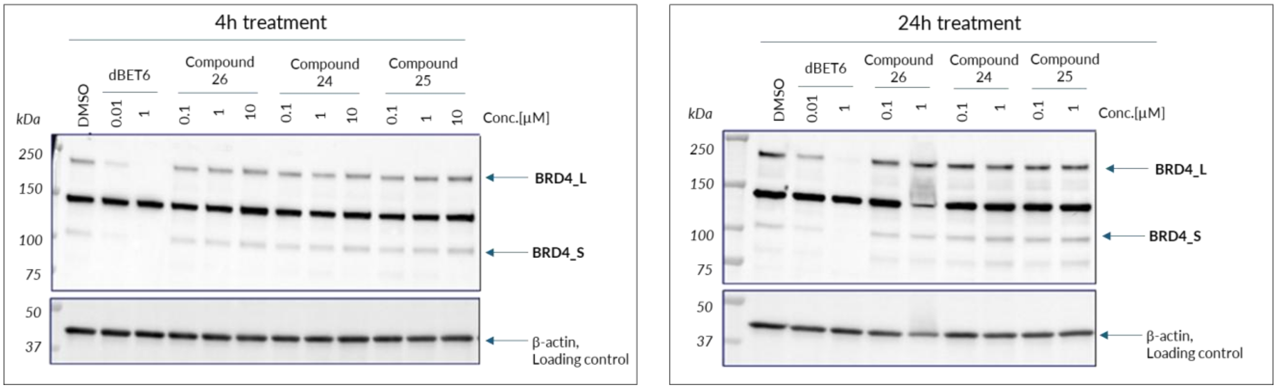


**Figure S11. BRD4 degradation assay in HEK293 cell line.** HEK293 cells were treated with selected GID4-BRD4 bifunctional compounds and the positive control degrader (dBET6) for 4 and 24 hours. After incubation, cells were collected by trypsinization, and whole cell lysates were isolated using RIPA lysis buffer with protease inhibitors. 25 µg of the cell lysate was resolved on an SDS-PAGE gel and transferred onto a nitrocellulose membrane to visualize BRD4 protein (the long isoform BRD4_L and the short isoform BRD4_S), as well as the loading control, using appropriate antibodies (anti-BRD4, cat. 128874, Abcam; anti-β-actin-HRP, cat. ab20272, Abcam). Data are representative of two independent biological replicates.

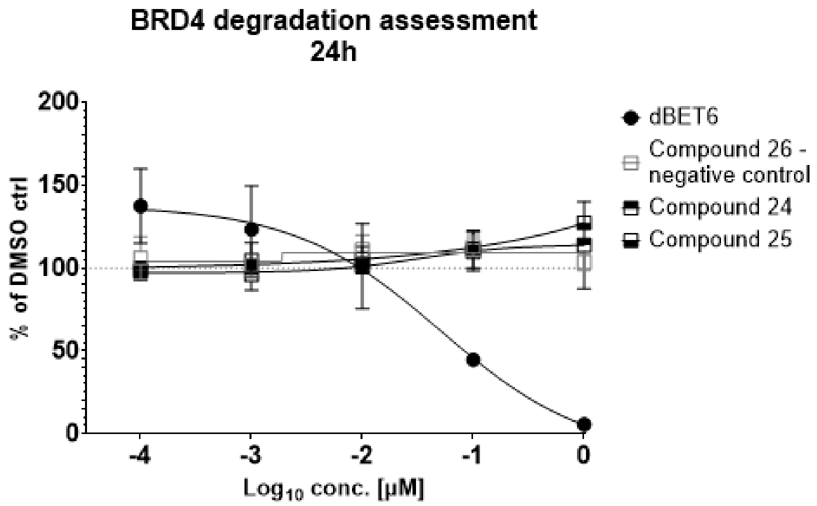


**Figure S12. HiBiT-BRD4 degradation assay in HEK293 cell line.** HiBiT-BRD4 KI HEK293 (LgBiT) cells (cat. CS302312, Promega) were treated with selected GID4-BRD4 bifunctional compounds and the positive control degrader, dBET6 for 24 hours (compounds concentration range: 0.1nM - 1 μM). After this time, luminescence levels corresponding to BRD4 protein expression were measured according to the manufacturer's recommendations using the Nano-Glo HiBiT Lytic Assay (cat. N3050, Promega) and compared to the DMSO control sample. Representative data from a single experiment performed in duplicate.

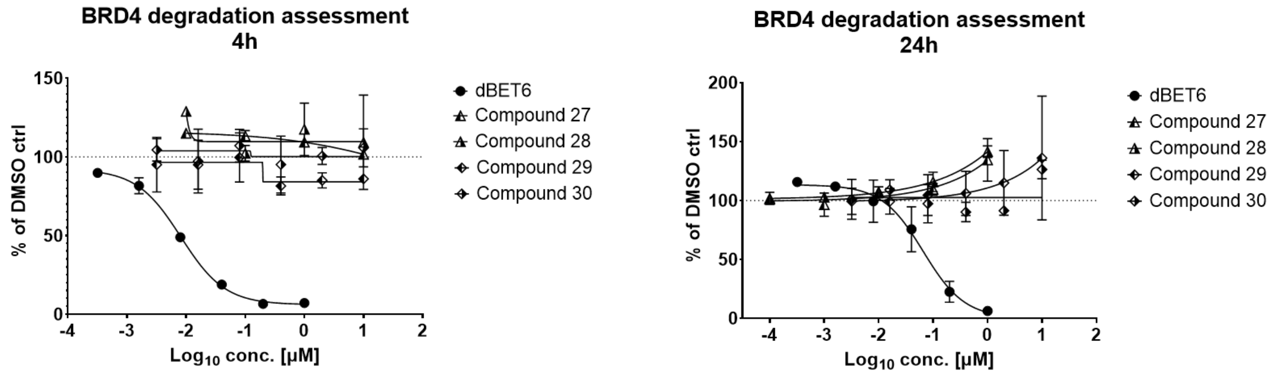


**Figure S13. HiBiT-BRD4 degradation assay in HEK293 cell line.** HiBiT-BRD4 KI HEK293 (LgBiT) cells (cat. CS302312, Promega) were treated with selected GID4-BRD4 bifunctional compounds and the positive control degrader, dBET6, for 4 and 24 hours (concentration range: 1 or 3 nM – 10 μM - tested compounds; 0.3 nM – 1 μM - dBET6). After this time, luminescence levels corresponding to BRD4 protein expression were measured according to the manufacturer's recommendations using the Nano-Glo HiBiT Lytic Assay (cat. N3050, Promega) and compared to the DMSO control sample. Representative data from a single experiment performed in duplicate.

**Table S3. Summary of GID4 compounds described in this study.**

**(* - crystal structure not available, cluster is unknown)**

| Compound | Structure | PDB ID | Cluster |
|---|---|---|---|
| 1 | 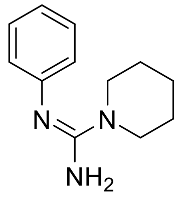 | 9QDX | C3 |
| 2 | 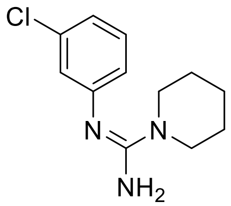 | n/a | C1 |
| 3 | 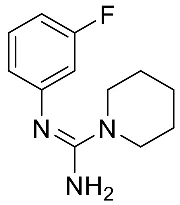 | n/a | * |
| 4 | 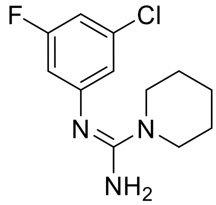 | n/a | * |
| 5 | 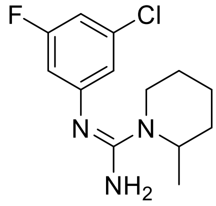 | n/a | * |
| 6 | 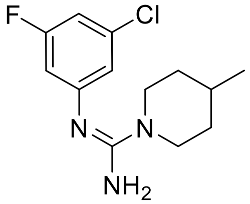 | n/a | * |

| | | | |
|---|---|---|---|
| 7 | 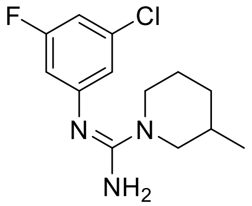 | n/a | * |
| 8 | 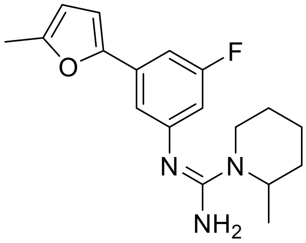 | n/a | * |
| 9 | 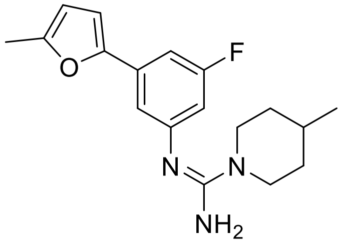 | 9QDY | C3 |
| 10 | 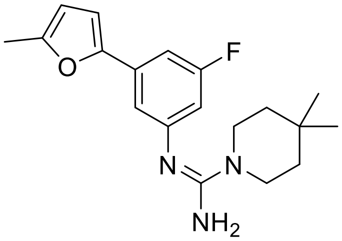 | n/a | * |
| 11 | 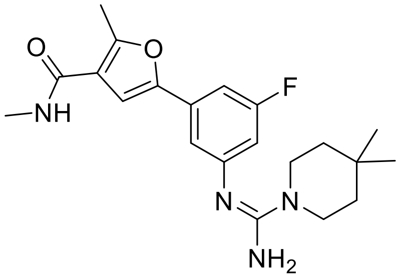 | n/a | * |
| 12 | 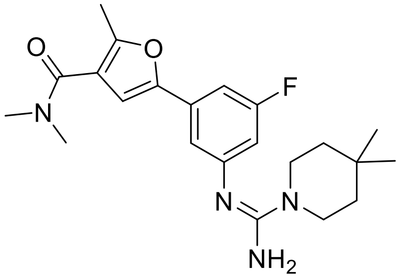 | n/a | * |

| | | | |
|---|---|---|---|
| 13 | 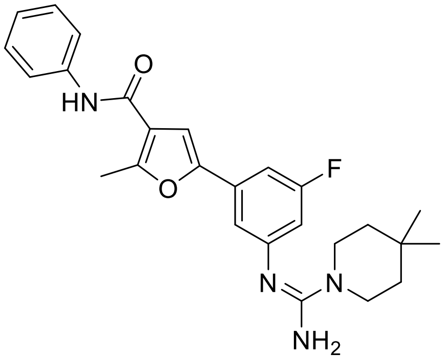 | n/a | C3 |
| 14 | 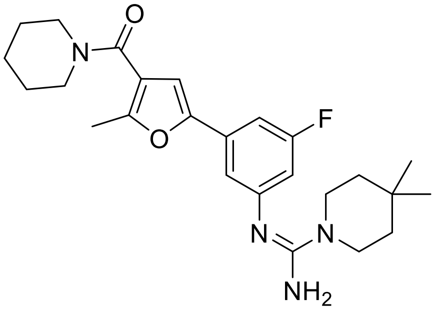 | 9QDZ | C3 |
| 15 | 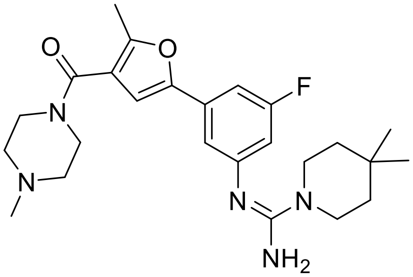 | n/a | C2 |
| 16 | 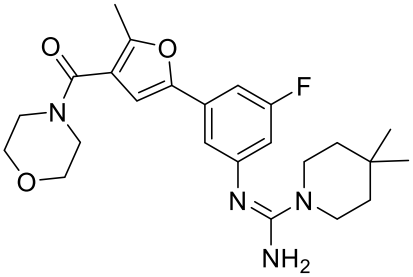 | n/a | * |
| 17 | 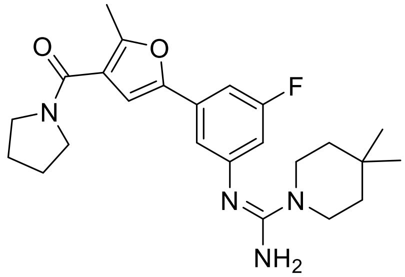 | n/a | * |

| | | | |
|---|---|---|---|
| 18 | 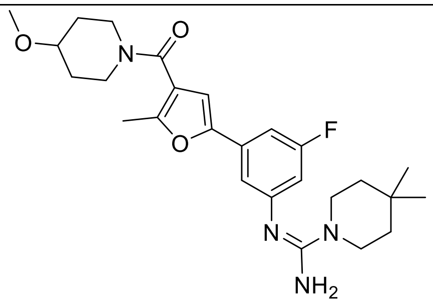 | 9QZG | C2 |
| 19 | 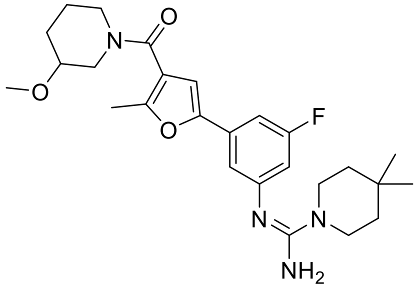 | n/a | * |
| 20 | 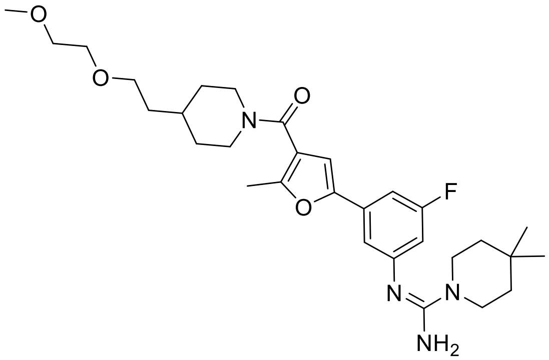 | n/a | * |
| 21 | 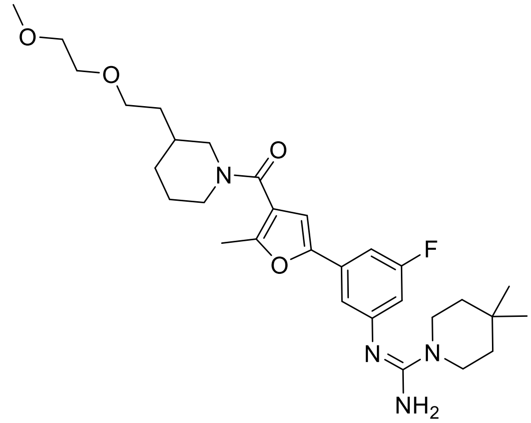 | 9QZI | C2 |

| | | | |
|---|---|---|---|
| 22 | 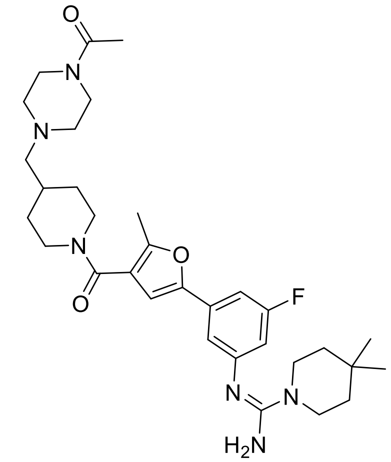 | n/a | * |
| 23 | 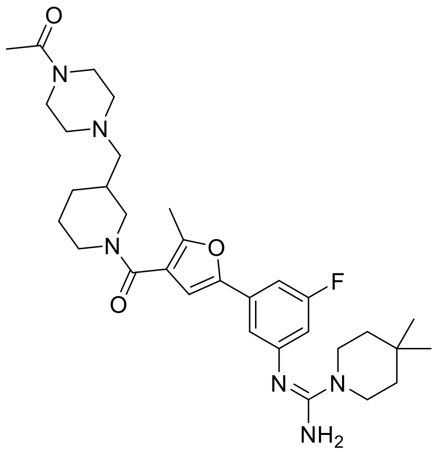 | n/a | * |
| 24 | 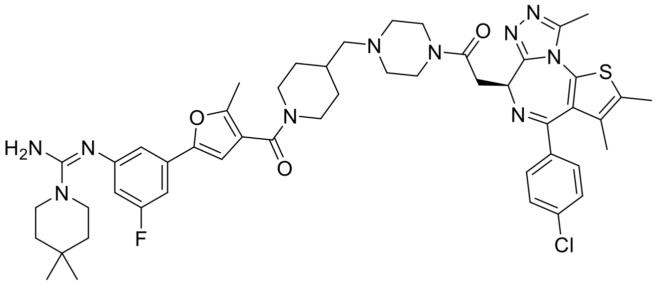 | n/a | * |

| | | | |
|---|---|---|---|
| 25 | 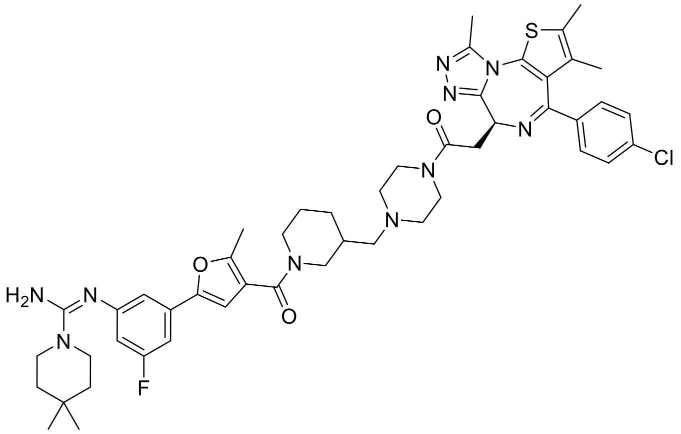 | n/a | * |
| 26 | 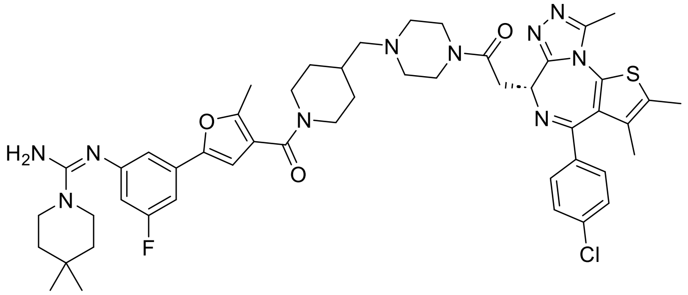 | n/a | * |
| 27 | 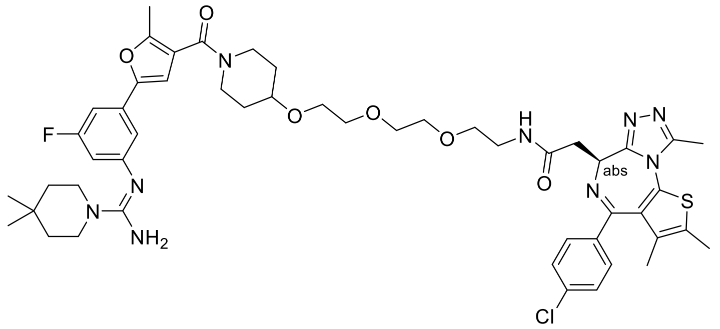 | n/a | * |
| 28 | 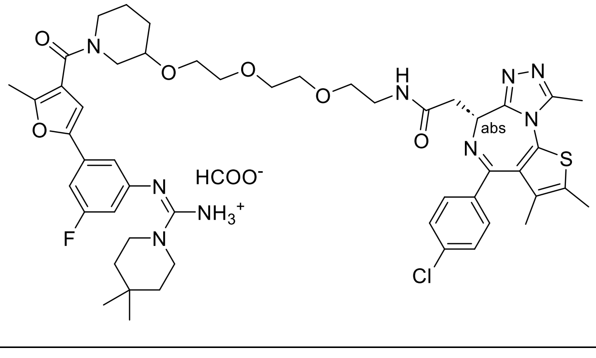 | n/a | * |

| | | | |
|---|---|---|---|
| 29 | 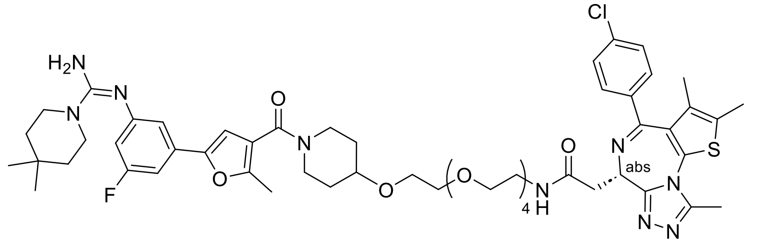 | n/a | * |
| 30 | 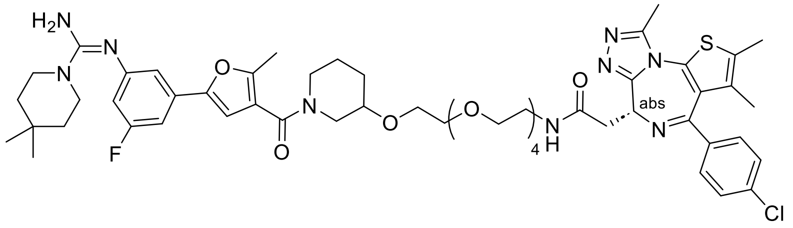 | n/a | * |
| 31 | 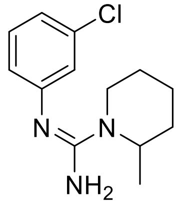 | n/a | C3 |
| 32 | 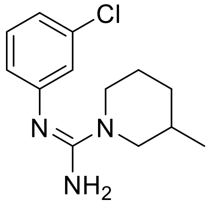 | n/a | C1 |
| 33 | 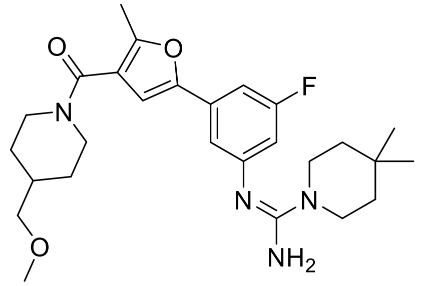 | 9QZH | C2 |
