## Supporting information_Chemical Synthesis for "Ligand-induced Conformational Plasticity of the CTLH E3 Ligase Receptor GID4"

1. **Synthetic procedures:**

**Synthesis of common intermediate Ia**

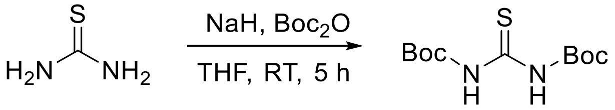


**Ia**

To a stirred suspension of NaH (60% dispersion in mineral oil, 4.729 g, 118.234 mmol, 4.5 eq.) in anhydrous THF (600 mL) at 0 °C was added thiourea (2.0 g, 26.274 mmol, 1.0 eq.) portion-wise. The resulting slurry was allowed to warm to room temperature and stirred for 15 min. The mixture was re-cooled to 0 °C before the dropwise addition of di-tert-butyl dicarbonate (12.616 g, 57.803 mmol, 2.2 eq.). Upon completion of the addition, the reaction was stirred at room temperature for 5 h. The reaction was quenched by the cautious addition of saturated aqueous sodium bicarbonate and water. The aqueous phase was extracted with ethyl acetate and the combined organic layers were dried over anhydrous sodium sulfate, filtered and concentrated under reduced pressure. The resulting residue was further dried *in vacuo* to afford N,N'-bis(tert-butoxycarbonyl)thiourea **Ia** (7.2 g, 99.2% yield) as a beige solid. The product was utilized in the subsequent synthetic step without further purification.

LCMS: No ionization

**Synthesis of common intermediates Ib and Ic**

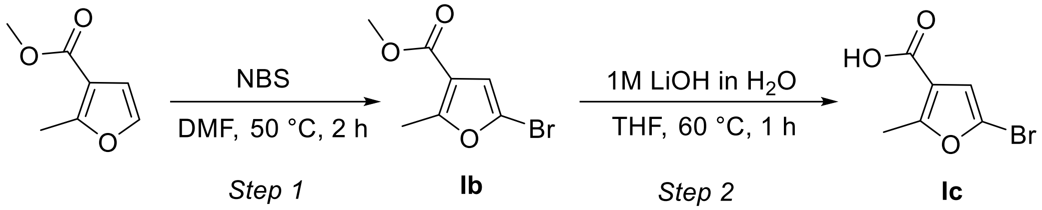


**Step 1:**

To a solution of methyl 2-methylfuran-3-carboxylate (10 g, 71.36 mmol, 1.0 eq.) in DMF (90 mL) at 0°C was added NBS (13.97 g, 78.49 mmol, 1.1 equiv.). The mixture was stirred at 0°C for 20 min and then heated at 50 °C for 2 h. Upon completion, the reaction mixture was diluted with water and extracted with ethyl acetate. The combined organic layers were washed with brine, dried over anhydrous sodium sulfate, filtered and concentrated under reduced pressure. The resulting residue was purified by silica gel column chromatography (EtOAc/Hex) to afford methyl 5-bromo-2-methylfuran-3-carboxylate **Ib** (14 g, 89% yield).

$^{1}$H NMR (400 MHz, chloroform-d) δ 6.55 (s, 1H), 3.81 (s, 3H), 2.56 (s, 3H).

**Step 2:**

To a solution of methyl 5-bromo-2-methylfuran-3-carboxylate (2.8 g, 12.783 mmol, 1.0 eq.) in THF (15.0 mL), was added an aqueous solution of LiOH (1M, 63.9 mL, 63.918 mmol, 5.0 eq.). The reaction mixture was stirred at 60 °C for 1 h. After cooling to room temperature, the solution was acidified to pH 1-2 by the addition of 1M HCl. The resulting mixture was extracted with DCM. The combined organic layers were dried over anhydrous sodium sulfate, filtered and concentrated under reduced pressure to afford 5-bromo-2-methylfuran-3-carboxylic acid (2.0 g, 76.3% yield) as a brown solid.

LCMS (ESI) $[M+H]^{+}$= no ionization

$^{1}$H NMR (400 MHz, DMSO-$d_6$) δ 12.88 (s, 1H), 6.70 (s, 1H), 2.49 (s, 3H).

**Synthesis of common intermediate Id**

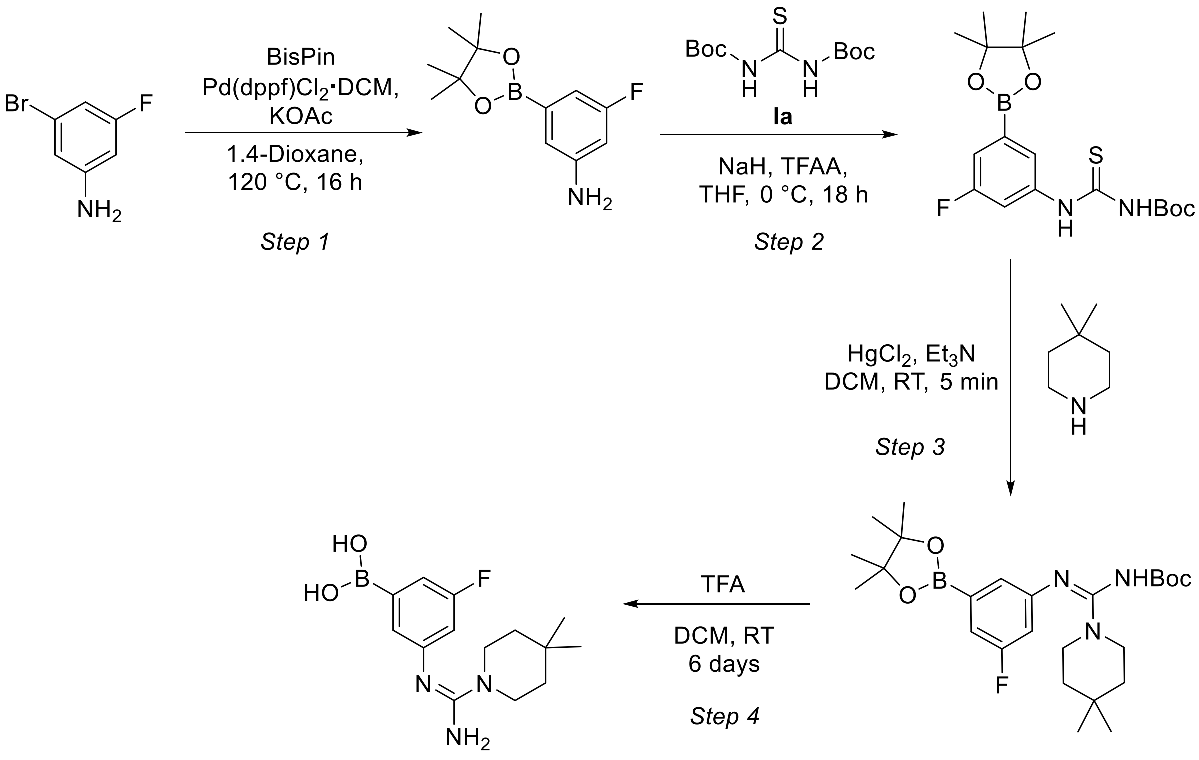


**Id**

**Step 1:**

To a stirred solution of 3-bromo-5-fluoroaniline (5.0 g, 26.32 mmol, 1.0 eq.) and Bis-Pin (8.02 g, 31.58 mmol, 1.2 eq.) in 1,4-dioxane (100 mL), was added KOAc (7.74 g, 78.95 mmol, 3.0 eq.). The reaction mixture was degassed with argon for 10 min, followed by the addition of Pd(dppf)$Cl_2$•DCM (0.96 g, 1.32 mmol, 0.05 eq.). The resulting mixture was heated at 120 °C for 16 h. Upon completion, the reaction was cooled to room temperature, diluted with water and extracted with ethyl acetate. The combined organic layers were washed with brine, dried over anhydrous sodium sulfate, filtered and concentrated under reduced pressure. The crude residue was purified by silica gel column chromatography to afford 3-fluoro-5-(4,4,5,5-tetramethyl-1,3,2-dioxaborolan-2-yl) aniline (4.5 g, 72% yield).

LCMS (ESI) $[M+H]^+$= 238.1

**Step 2:**

To a stirred solution of N,N'-(tert-butoxycarbonyl)thiourea **Ia** (3.08 g, 11.14 mmol, 1.1 eq.) in anhydrous THF (15 mL) at 0 °C was added NaH (60% dispersion in mineral oil, 0.49 g, 12.15 mmol,

1.2 eq.) portion-wise. After stirring for 30 min, trifluoroacetic anhydride (1.6 mL, 11.135 mmol, 1.1 eq.) was added dropwise and the resulting solution was stirred at 0 °C for an additional 30 min. 3-fluoro-5-(4,4,5,5-tetramethyl-1,3,2-dioxaborolan-2-yl) aniline (1.32 g, 5.57 mmol, 1.5 eq.) was then added and the reaction mixture was allowed to warm to room temperature and stirred for 16 h. Upon completion, the mixture was diluted with ice-cold water and extracted with ethyl acetate. The combined organic layers were dried over anhydrous sodium sulfate, filtered and concentrated under reduced pressure. The resulting residue was further dried *in vacuo* to afford tert-butyl N-{[3-fluoro-5-(4,4,5,5-tetramethyl-1,3,2-dioxaborolan-2-yl)phenyl]carbamothioyl}carbamate (3.4 g, 85% yield) as a white solid.

LCMS (ESI) $[M+H]^+$= 397.3

**Step 3:**

To a stirred solution of tert-butyl N-{[3-fluoro-5-(4,4,5,5-tetramethyl-1,3,2-dioxaborolan-2-yl)phenyl]carbamothioyl}carbamate (3.40 g, 8.58 mmol, 1.0 eq.) in DCM (15.0 mL) were added mercury chloride (2.56 g, 9.44 mmol, 1.1 eq.), 4,4-dimethylpiperidine hydrochloride (1.41 g, 9.44 mmol, 1.1 eq.) and triethylamine (6.0 mL, 42.90 mmol, 5.0 eq.). The reaction mixture was stirred at room temperature for 5 min. The resulting suspension was filtered through Celite and the filter cake was washed thoroughly with DCM. The filtrate was concentrated under reduced pressure to afford tert-butyl N-[(Z)-(4,4-dimethylpiperidin-1-yl)({[3-fluoro-5-(4,4,5,5-tetramethyl-1,3,2-dioxaborolan-2-yl)phenyl]imino})methyl]carbamate (3.7 g, 91% yield), which was used in the next step without further purification.

LCMS (ESI) $[M+H]^+$= 476.2

**Step 4:**

To a stirred solution of tert-butyl N-[(Z)-(4,4-dimethylpiperidin-1-yl)({[3-fluoro-5-(4,4,5,5-tetramethyl-1,3,2-dioxaborolan-2-yl)phenyl]imino})methyl]carbamate (3.70 g, 7.78 mmol, 1.0 eq.) in DCM (10.0 mL) was added TFA (2.0 mL, 26.12 mmol, 3.4 eq.). The reaction mixture was stirred at room temperature for 6 days. Upon completion, the mixture was concentrated under reduced pressure. The resulting residue was diluted with water and subsequently freeze dried to afford (Z)-(3-((amino(4,4-dimethylpiperidin-1-yl)methylene)amino)-5-fluorophenyl)boronic acid **Id** (2.0 g, 87.7% yield) as a light beige solid that was used in the next step without further purification.

LCMS (ESI) $[M+H]^+$= 294.2

**Synthesis of common intermediate Ie**

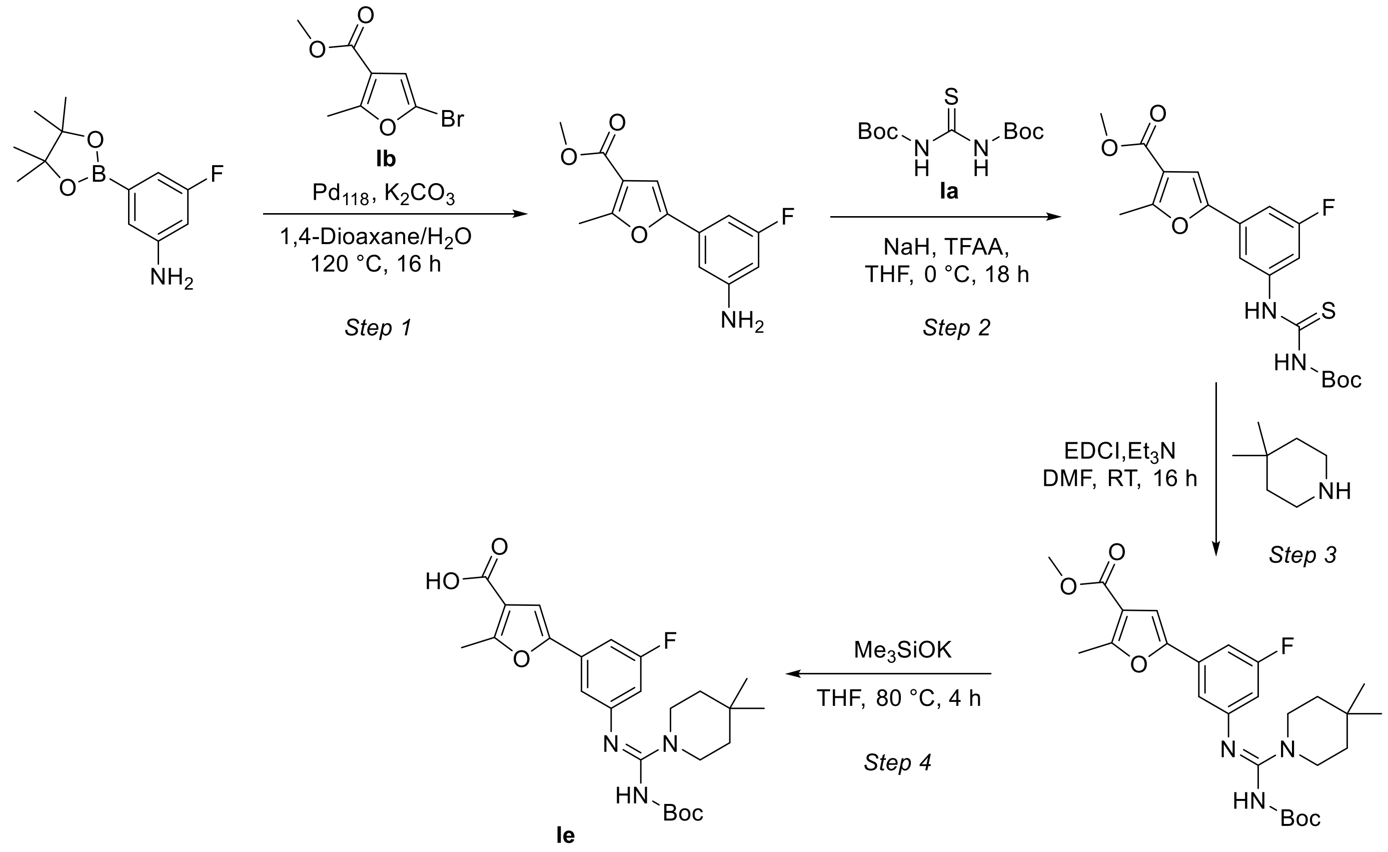


**Step 1:**

To a stirred solution of 3-fluoro-5-(4,4,5,5-tetramethyl-1,3,2-dioxaborolan-2-yl)aniline (1.5 g, 6.33 mmol, 1.0 eq.) and methyl 5-bromo-2-methylfuran-3-carboxylate (2.08 g, 9.49 mmol, 1.5 eq.) in a mixture of 1,4-dioxane/$H_2O$ (10:1, 36 mL) was added potassium carbonate (2.6 g, 18.98 mmol, 3.0 eq.). The reaction mixture was degassed with argon for 15 min, followed by the addition of $Pd_{118}$ (0.49 g, 0.63 mmol, 0.1 equiv.). The resulting mixture was heated to 100 °C and stirred overnight (16 h). Upon completion, the solvent was removed under reduced pressure. The residue was diluted with water and extracted with ethyl acetate. The combined organic layers were washed with brine, dried over anhydrous sodium sulfate, filtered and concentrated under reduced pressure. The crude product was purified by silica gel column chromatography (EtOAc/Hex) to afford the methyl 5-(3-amino-5-fluorophenyl)-2-methylfuran-3-carboxylate (0.65 g, 41% yield).
LCMS (ESI) $[M+H]^+$= 250.1

**Step 2:**

To a stirred solution of N,N'-(tert-Butoxycarbonyl)thiourea (0.87 g, 3.15 mmol, 1.0 eq.) in anhydrous THF (15 mL) at 0 °C was added NaH (0.19 g, 4.72 mmol, 1.5 eq.) portion-wise. The mixture was stirred for 1 h, followed by the addition of trifluoroacetic anhydride (0.48 ml, 3.46 mmol, 1.1 eq.). The resulting solution was stirred at 0 °C for an additional 1 h. Methyl 5-(3-amino-5-fluorophenyl)-2-methylfuran-3-carboxylate (0.86 g, 1.1 mmol, 1.1 eq.) was then added and the reaction was stirred for 16 h at room temperature. Upon completion, the mixture was diluted with water and extracted with ethyl acetate. The combined organic layers were washed with brine, dried over anhydrous sodium sulfate, filtered and concentrated under reduced pressure to afford methyl 5-(3-(3-(tert-butoxycarbonyl)thioureido)-5-fluorophenyl)-2-methylfuran-3-carboxylate (quantitative yield), which was used in the subsequent step without further purification.

LCMS (ESI) $[M+H]^+$= 408.9

**Step 3:**

To a stirred solution of methyl 5-(3-(3-(tert-butoxycarbonyl)thioureido)-5-fluorophenyl)-2-methylfuran-3-carboxylate (0.75 g, 1.84 mmol, 1.0 eq.) in anhydrous DMF (15 mL) at 0 °C were added EDCI (0.53g, 2.76 mmol, 1.5 eq.) and $Et_3N$ (0.77 mL, 5.52 mmol, 3.0 eq.). After stirring for 15 min, 4,4-dimethylpiperidine (0.28 mL, 2.2 mmol, 1.2 eq.) was added. The reaction mixture was allowed to warm to room temperature and stirred for 16 h. Upon completion, the mixture was diluted with water and extracted with ethyl acetate. The combined organic layers were washed with brine, dried over anhydrous sodium sulfate, filtered and concentrated under reduced pressure. The crude residue was purified by silica gel column chromatography (EtOAc/Hex) to afford methyl (Z)-5-(3-((((tert-butoxycarbonyl)amino)(4,4-dimethylpiperidin-1-yl)methylene)amino)-5-fluorophenyl)-2-methylfuran-3-carboxylate (0.225 g, 25% yield).

LCMS (ESI) $[M+H]^+$= 488.2

**Step 4:**

To a stirred solution of methyl (Z)-5-(3-((((tert-butoxycarbonyl)amino)(4,4-dimethylpiperidin-1-yl)methylene)amino)-5-fluorophenyl)-2-methylfuran-3-carboxylate (1.3 g, 2.66 mmol, 1.0 eq.) in anhydrous THF (15 mL) was added potassium trimethylsilanolate (0.68 g, 5.34 mmol, 2.0 eq.). The reaction mixture was stirred at 80 °C for 4 h. Upon cooling to room temperature, the solvent was removed under reduced pressure. The resulting residue was acidified with a saturated aqueous solution of $NaHSO_4$ and extracted with 10% MeOH in DCM. The combined organic layers were washed with brine, dried over anhydrous sodium sulfate, filtered and concentrated under reduced pressure. The crude material was triturated with pentane to afford (Z)-5-(3-((((tert-butoxycarbonyl)amino)(4,4-dimethylpiperidin-1-yl) methylene)amino)-5-fluorophenyl)-2-methylfuran-3-carboxylic acid **Ie** (1.2 g, 94% yield) as a light yellow solid.

LCMS (ESI) $[M+H]^+$= 474.2

$^1$H NMR (400 MHz, DMSO-$d_6$) δ 12.56 (s, 1H), 9.04 (s, broad, 1H), 7.14 – 7.05 (m, 2H), 7.03 – 6.85 (m, 1H), 6.61 – 6.37 (m, 1H), 3.47 – 3.41 (m, 4H), 2.58 (s, 3H), 1.39 – 1.33 (m, 4H), 1.21 (s, 9H), 0.97 (s, 6H).

**Synthesis of Compound 2**

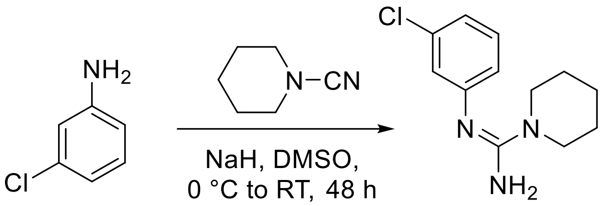


**2**

To a stirred suspension of NaH (60% dispersion in mineral oil, 85 mg, 3.54 mmol, 1.5 eq.) in anhydrous DMSO (3.0 mL) at 0 °C was added 3-chloroaniline (300 mg, 2.36 mmol, 1.0 eq.) in DMSO (1.0 mL) at 0 °C. The reaction mixture was allowed to warm to room temperature and stirred for 45 min. The mixture was re-cooled to 0 °C before the addition of piperidine-1-carbonitrile (311 mg, 2.83 mmol, 1.2 eq.). The resulting mixture was stirred at room temperature for 48 h. Upon completion, the reaction was quenched with ice-cold water and extracted with MTBE. The combined organic layers were washed with cold water and brine, dried over anhydrous sodium sulfate, filtered and concentrated under reduced pressure. The crude residue was triturated with mixture of hexanes and diethyl ether to afford (Z)-N'-(3-chlorophenyl)piperidine-1-carboximidamide (16.5 mg, 3% yield) as a dark brown solid.

$^1$H NMR (400 MHz, DMSO-$d_6$) δ 7.16 (t, *J* = 7.9 Hz, 1H), 6.81 (d, *J* = 7.8 Hz, 1H), 6.70 (m, 1H), 6.65 (d, *J* = 8.0 Hz, 1H), 5.43 (s, 2H), 3.33 – 3.28 (m, 4H), 1.59 – 1.45 (m, 6H).

LCMS (ESI) $[M+H]^+$= 237.97

**Synthesis of Compound 3**

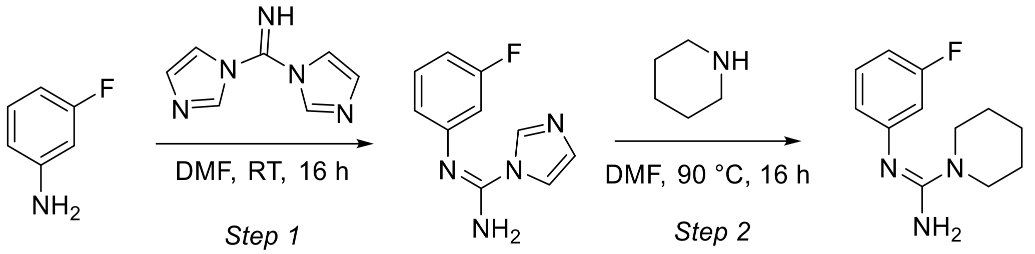


**3**

**Step 1:**

To a stirred solution of di(1H-imidazol-1-yl) methamine (200.0 mg, 1.24 mmol, 1.0 eq.) in anhydrous DMF (3.0. mL) was added 3-fluoroaniline (0.132 mL,1.37 mmol, 1.1 eq.). The reaction mixture was stirred for 16 h at room temperature. Upon completion, the reaction was diluted with water and extracted with ethyl acetate. The combined organic layers were washed with brine, dried over anhydrous sodium sulfate, filtered and concentrated under reduced pressure. The crude residue was purified by silica gel column chromatography (MeOH/DCM) to afford (Z)-N'-(3-fluorophenyl)-1H-imidazole-1-carboximidamide (210 mg, 85 % yield).

LCMS (ESI) $[M+H]^+$= 205.2

**Step 2:**

To a stirred solution of N-(3-fluorophenyl)-1H-imidazole-1-carboximidamide (200.0 mg, 0.98 mmol, 1.0 eq.) in anhydrous DMF (3.0 mL) was added piperidine (0.106 mL, 1.08 mmol, 1.1 eq.). The reaction mixture was heated at 90 °C and stirred for 16 h. Upon completion, the mixture was cooled to room temperature, diluted with water and extracted with ethyl acetate. The combined organic layers were washed with brine, dried over anhydrous sodium sulfate, filtered and concentrated under reduced pressure. The crude residue was purified by prep-HPLC (aq. $NH_4HCO_3$/ACN) to afford (Z)-N'-(3-fluorophenyl)piperidine-1-carboximidamide (75 mg, 35% yield) as a white powder.

$^1$H NMR (500 MHz, DMSO-$d_6$) δ7.16 (td, *J* = 8.1, 7.3 Hz, 1H), 6.61 – 6.55 (m, 1H), 6.52 (ddd, *J* = 8.0, 2.0, 0.9 Hz, 1H), 6.45 (dt, *J* = 11.6, 2.3 Hz, 1H), 5.32 (s, 1H), 5.31 (s, 2H), 3.32 – 3.29 (m, 4H), 1.56 – 1.46 (m, 6H).

LCMS: $[M+H]^+$= 222.2

**General procedure A for the synthesis of Compounds 4 to 7**

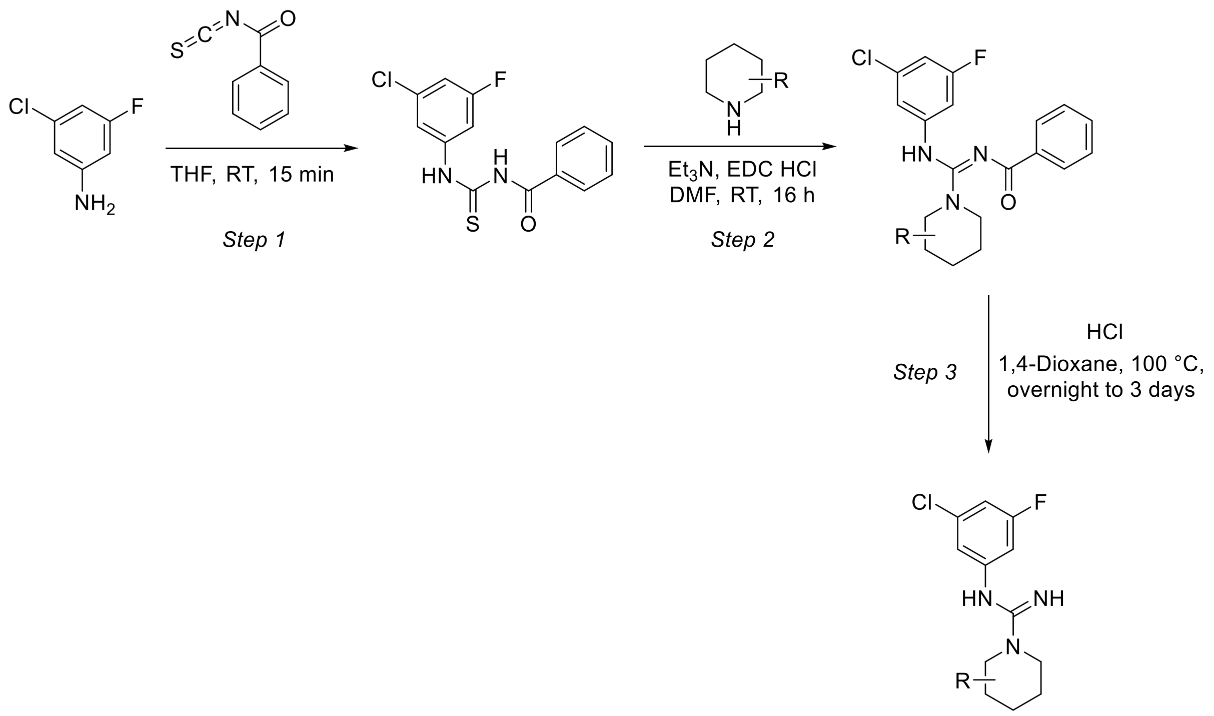


**Step 1**:

To a stirred solution of 3-chloro-5-fluoroaniline (1.0 g, 6.87 mmol, 1.0 eq.) in anhydrous THF (10.0 mL) was added benzoyl isothiocyanate (0.927 mL, 6.87 mmol, 1.0 eq.). The resulting mixture was stirred at room temperature for 15 min. Upon completion, the reaction mixture was diluted with ethyl acetate and washed with brine. The organic layers were dried over anhydrous sodium sulfate, filtered and concentrated under reduced pressure to afford N-((3-chloro-5 fluorophenyl)carbamothioyl)benzamide (2.08 g, 98.1% yield) as a yellow solid. The crude product was utilized in the subsequent step without further purification.

LCMS: [M+H]= 308.8

**Step 2:**

To a stirred solution of N-((3-chloro-5 fluorophenyl)carbamothioyl)benzamide (0.259 - 0.972 mmol, 1.0 eq.) in anhydrous DMF (2.0- 5.0 mL) were added corresponding piperidine (1.2 eq.), triethylamine (3.0 eq.) and EDC · HCl (1.5 eq.) were added. The resulting mixture was stirred at room temperature overnight. Upon completion, the mixture was diluted with water and extracted with DCM. The organic layers were separated, dried over anhydrous sodium sulfate, filtered and concentrated under reduced pressure to afford the corresponding product. The crude material was utilized in the subsequent step without further purification.

**<u>Step 3:</u>**

To a stirring solution of the corresponding *N*-benzoyl-guanidine substrate (0.219 - 0.778 mmol) in 1,4-dioxane (2.0 mL) was added concentrated hydrochloric acid (35% in water, 2.0 mL). The reaction mixture was heated at 100 °C and stirred for a period ranging from 16 h to 3 days. Upon completion, the mixture was concentrated under reduced pressure and the resulting residue purified by reversed-phase silica gel column chromatography (ACN/water + 0.1% FA) to afford desired product.

**Compound 4**

(Z)-N'-(3-chloro-5-fluorophenyl)piperidine-1-carboximidamide, formate salt (78.0 mg, 39.2% yield), colorless oil

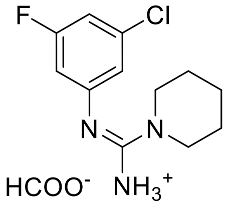


$^{1}$H NMR (500 MHz, methanol-d4) δ 7.14 – 7.12 (m, 1H), 7.10 (dt, 1H, *J* = 8.4, 2.1 Hz, 1H), 6.99 (dt, *J* = 9.8, 2.1 Hz, 1H), 3.59 – 3.53 (m, 4H), 1.79 – 1.70 (m, 6H).

LCMS: $[M+H]^{+}$= 256.0

**Compound 5**

(Z)-N'-(3-chloro-5-fluorophenyl)-2-methylpiperidine-1-carboximidamide, formate salt (128.0 mg, 63% yield), colorless oil.

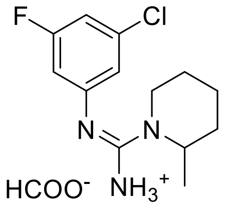

$^1$H NMR (500 MHz, methanol-d4) δ 7.14 – 7.13 (m, *J* = 2.6, 1.8 Hz, 1H), 7.13 – 7.09 (m, 1H), 7.00 (dt, *J* = 9.7, 2.1 Hz, 1H), 4.34 – 4.27 (m, 1H), 3.78 – 3.72 (m, 1H), 3.68 – 3.62 (m, 1H), 1.86 – 1.57 (m, 6H), 1.38 (d, *J* = 6.9 Hz, 3H).

LCMS: $[M+H]^+$= 270.0

**Compound 6**

(Z)-N'-(3-chloro-5-fluorophenyl)-4-methylpiperidine-1-carboximidamide, formate salt (15 mg, 25.4 % yield), white solid

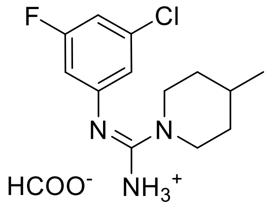


$^1$H NMR (500 MHz, methanol-d4) δ 7.14 – 7.08 (m, 2H), 6.99(dt, *J* = 9.7, 2.1 Hz, 1H)), 3.99 – 3.89 (m, 2H), 3.22 – 3.14 (m, 2H), 1.85 – 1.79 (m, 2H), 1.85 – 1.72 (m, 3H), 1.35-1.26 (m, 2H), 1.03 (d, *J* = 6.5 Hz, 3H).

LCMS: $[M+H]^+$= 270.3

**Compound 7**

(Z)-N'-(3-chloro-5-fluorophenyl)-3-methylpiperidine-1-carboximidamide, formate salt (78.0 mg, 37% yield), colorless oil

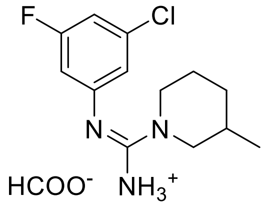


$^1$H NMR (500 MHz, methanol-d4) δ 7.13 – 7.12 (m, 1H), 7.10 (dt, *J* = 8.5, 2.1 Hz, 1H), 6.99 (dt, *J* = 9.8, 2.1 Hz, 1H), 3.92 – 3.80 (m, 2H), 3.14 (ddd, *J* = 13.5, 12.1, 3.0 Hz, 1H), 2.82 (dd, *J* = 13.3, 11.1 Hz, 1H), 1.96 – 1.89 (m, 1H), 1.87 – 1.73 (m, 2H), 1.72 – 1.61 (m, 1H), 1.28 (dd, *J* = 11.1, 4.0 Hz, 1H), 0.98 (d, *J* = 6.6 Hz, 3H).

LCMS: $[M+H]^+$= 270.0

**General procedure B for the synthesis of Compounds 8 to 10**

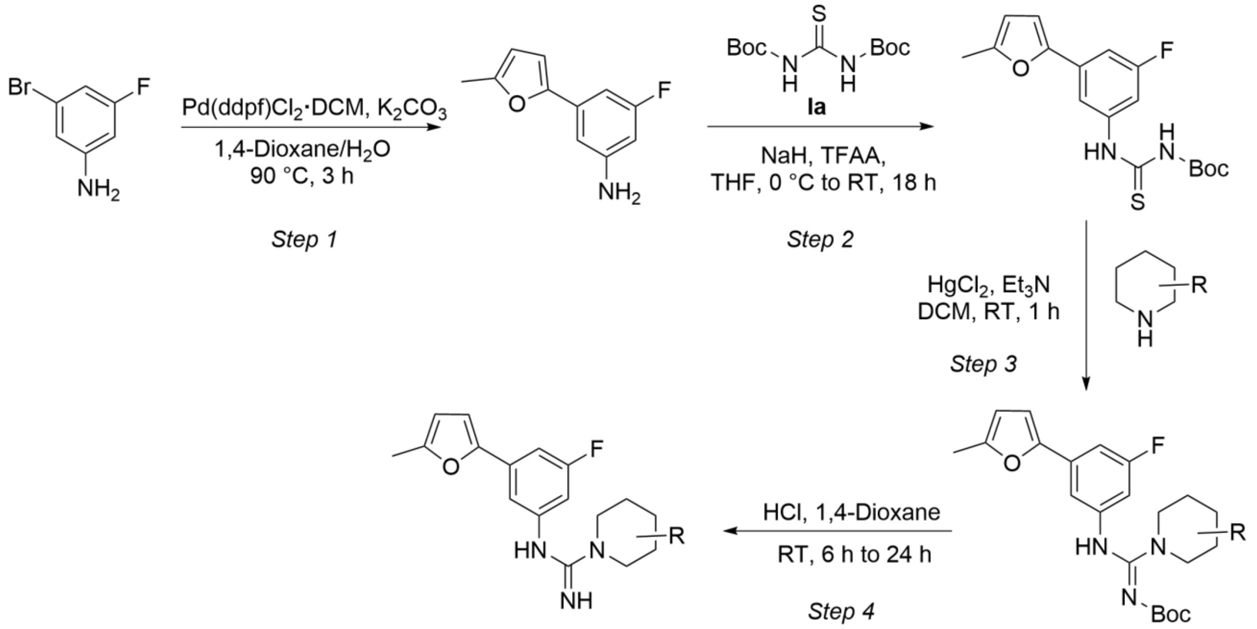


**Step 1:**

To a stirred solution of 3-bromo-5-fluoroaniline (400.0 mg, 2.11 mmol, 1.0 eq.), 4,4,5,5-tetramethyl-2-(5-methylfuran-2-yl)-1,3,2-dioxaborolane (525.6 mg, 2.53 mmol, 1.2 eq.) in a mixture of 1,4-dioxane (5.0 mL) and water (2.0 mL) was added potassium carbonate (727.3 mg, 5.26 mmol, 2.5 eq.). The mixture was degassed, followed by the addition of Pd(dppf)$Cl_2$•DCM (171.9 mg, 0.21 mmol, 0.1 eq.). The reaction mixture was heated at 90 °C and stirred for 3 h. Upon completion, the resulting solution was filtered through a pad of Celite and a filter cake was washed with ethyl acetate and water. The organic layers were separated and concentrated under reduced pressure to afford 3-fluoro-5-(5-methylfuran-2-yl)aniline (360 mg, 89% yield) as a brown oil. The crude product was utilized in the next step without further purification.

LCMS: $[M+H]^+$= 192.1

**Step 2:**

To a stirred solution of NaH (60% dispersion in mineral oil, 78.2 mg, 1.95 mmol, 1.2 eq.) in anhydrous THF (25.0 mL) at 0 °C was added N,N'-bis(tert-butoxycarbonyl)thiourea (450.0 mg, 1.63 mmol, 1.0 eq.). The resulting mixture was stirred for 30 min, followed by the addition of trifluoroacetic anhydride (0.253 mL, 1.79 mmol, 1.1 eq.). The reaction was maintained at 0 °C for an additional 30 min. 3-fluoro-5-(5-methylfuran-2-yl)aniline (342.5 mg, 1.79 mmol, 1.1 eq.) was then added and reaction mixture was stirred overnight at room temperature. The resulting mixture was quenched with water and extracted with ethyl acetate. The organic layers were separated, concentrated under reduced pressure and the residue was purified by silica gel column chromatography (EtOAc/*iso*Hex) to afford tert-butyl N-{[3-fluoro-5-(5-methylfuran-2-yl)phenyl]carbamothioyl} carbamate (550.0 mg, 96.4% yield) as a beige solid.

LCMS: $[M+H]^+$= 352.0

**Step 3:**

To a stirred solution of tert-butyl N-{[3-fluoro-5-(5-methylfuran-2-yl)phenyl]carbamothioyl}carbamate (50.0 mg, 142.69 mmol, 1.0 eq.) in DCM (2.0 mL) were added corresponding piperidine (1.2 eq.), mercury chloride (1.1 eq.) and triethylamine (5.0 eq.). the reaction mixture was stirred at room temperature for 1 h. Upon completion, the resulting suspension was filtered through a pad of Celite and washed thoroughly with DCM. The filtrate was concentrated under reduced pressure to afford crude product, which was utilized in the next step without further purification.

**Step 4:**

To a stirred solution of the corresponding *N*-Boc-guanidine substrate (0.132 – 0.409 mmol, 1.0 eq.) in 1,4-dioxane (2.0 mL) was added concentrated HCl (36% in water, approx. 83.0 - 88. eq.) The reaction mixture was stirred at room temperature for a period ranging from 6 to 24 h. Upon completion, the resulting solution was concentrated under reduced pressure and residue purified by prepHPLC (ACN/$H_2O$ + 0.1% FA) to afford the desired product.

**Compound 8**

(Z)-N'-(3-fluoro-5-(5-methylfuran-2-yl)phenyl)-2-methylpiperidine-1-carboximidamide formate (20 mg, 47.9% yield), beige oil

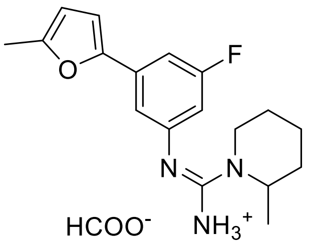


$^1$H NMR (500 MHz, methanol-d4) δ 7.32 (t, *J* = 1.6 Hz, 1H), 7.27 (ddd, *J* = 9.8, 2.3, 1.4 Hz, 1H), 6.86 (dt, *J* = 9.6, 2.1 Hz, 1H), 6.77 (d, *J* = 3.3 Hz, 1H), 6.16 – 6.14 (m, 1H), 4.36 – 4.292 (m, 1H),

3.78 – 3.73 (m, 1H), 3.35 (s, 1H), 2.36 (s, 3H), 1.88 – 1.78 (m, 3H), 1.75 – 1.68 (m, 2H), 1.66 – 1.59 (m, 1H), 1.39 (d, *J* = 6.9 Hz, 3H).

LCMS: $[M+H]^+$= 316.2

**Compound 9**

(Z)-N'-(3-fluoro-5-(5-methylfuran-2-yl)phenyl)-4-methylpiperidine-1-carboximidamide formate (9.0 mg, 7.0% yield), yellow solid

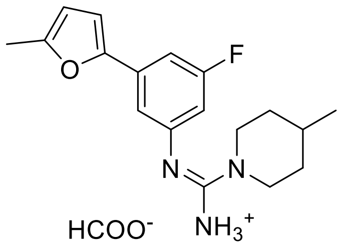


$^1$H NMR (500 MHz, methanol-d4) δ 7.32 (t, *J* = 1.7 Hz, 1H), 7.27 (ddd, *J* = 9.8, 2.4, 1.4 Hz, 1H), 6.86 (dt, *J* = 9.6, 2.2 Hz, 1H), 6.77 (d, *J* = 3.2 Hz, 1H), 6.18 – 6.12 (m, 1H), 4.00 – 3.92 (m, 2H), 3.22 – 3.15 (m, 2H), 2.36 (s, 3H), 1.86 – 1.72 (m, 3H), 1.38 – 1.29 (m, 2H), 1.03 (d, *J* = 6.5 Hz, 3H).

LCMS: $[M+H]^+$= 316.2

<u>**Compound 10**</u>

(Z)-N'-(3-fluoro-5-(5-methylfuran-2-yl)phenyl)-4,4-dimethylpiperidine-1-carboximidamide formate (20.0 mg, 43.5% yield), yellow solid

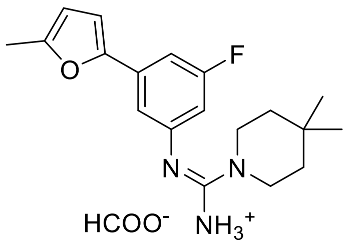


$^1$H NMR (500 MHz, methanol-d4) δ 7.33 (t, *J* = 1.6 Hz, 1H), 7.27 (ddd, *J* = 9.8, 2.3, 1.4 Hz, 1H), 6.86 (dt, *J* = 9.6, 2.2 Hz, 1H), 6.77 (d, *J* = 3.3 Hz, 1H), 6.17 – 6.14 (m, 1H), 3.60 – 3.55 (m, 4H), 2.36 (s, 3H), 1.58 – 1.53 (m, 4H), 1.07 (s, 6H)

LCMS: $[M+H]^+$= 329.4

### Synthesis of Compound 11

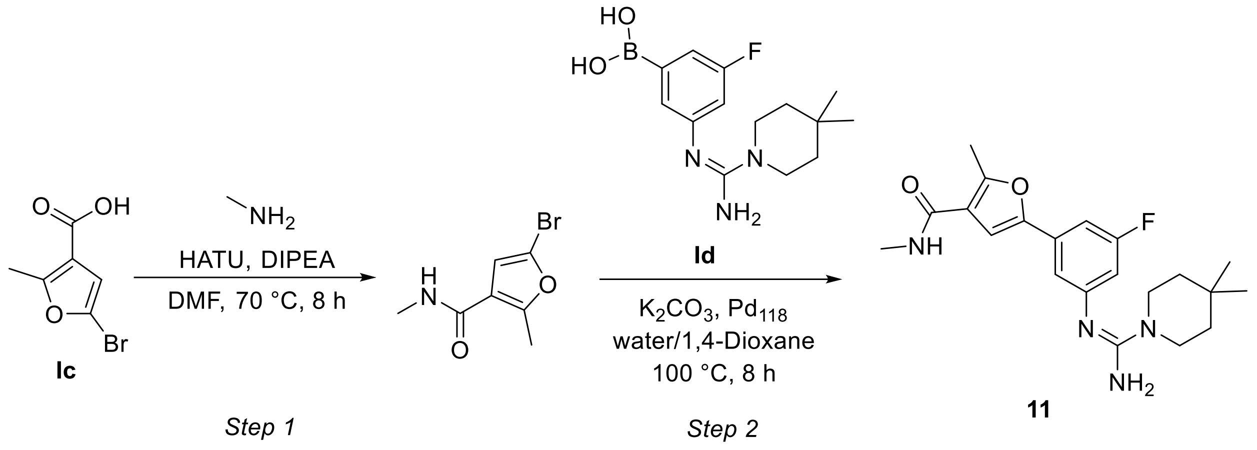


**Step 1:**

To a stirred solution of 5-bromo-2-methylfuran-3-carboxylic acid **Ic** (500 mg, 2.44 mmol, 1.0 eq.) in anhydrous DMF (5 mL) was added HATU (1.39 g, 3.64 mmol, 1.5 eq.) followed by DIPEA (1.2 mL, 7.32 mmol, 3.0 eq.). The resulting mixture was stirred at room temperature for 30 min. Subsequently, methyl amine (0.576 mL, 5.17 mmol, 2.1 eq.) was added. The reaction mixture was heated at 70 °C and stirred for 8 h. Upon completion, the mixture was quenched with ice-cold water and extracted with ethyl acetate. The combined organic layers were dried over anhydrous sodium sulfate, filtered and concentrated under reduced pressure. The crude residue was purified by silica gel column chromatography (EtOAc/Hex) to afford 5-bromo-N,2-dimethylfuran-3-carboxamide (260 mg, 49% yield).

LCMS (ESI) $[M+H]^+$= 218.1

**Step 2:**

To a stirred solution of 5-bromo-N,2-dimethylfuran-3-carboxamide (60 mg, 0.28 mmol, 1.0 eq.) and (Z)-(3-((amino(4,4-dimethylpiperidin-1-yl)methylene)amino)-5-fluorophenyl)boronic acid **Id** (108 mg, 0.33 mmol, 1.2 eq.) in a mixture of 1,4-dioxane and $H_2O$ (2:1, 3 mL) was added potassium carbonate (114 mg, 0.825 mmol, 3.0 eq.). The mixture was degassed with argon for 10 min. Subsequently, $Pd_{118}$ (18 mg,0.028 mmol, 0.1 eq.) was added. Reaction vessel was sealed and heated at 120 °C for 8 h. Upon completion, the reaction mixture was cooled to room temperature and filtered through Celite. The filtrate was concentrated under reduced pressure and the resulting crude residue was purified by prepHPLC (aq. $NH_4HCO_3$/ACN) to afford (Z)-5-(3-((amino(4,4-

dimethylpiperidin-1-yl)methylene)amino)-5-fluorophenyl)-N,2-dimethylfuran-3-carboxamide (10 mg, 9% yield) as an off white solid.

$^1$H NMR (400 MHz, methanol-d4) δ 7.08 – 7.00 (m, 2H), 6.99 – 6.95 (m, 1H), 6.63 – 6.58 (m, , 1H), 3.49 – 3.40 (m, 4H), 2.86 (s, 3H), 2.60 (s, 3H), 1.50 – 1.41 (m, 4H), 1.02 (s, 6H).

LCMS (ESI) $[M+H]^+$= 387.2

**Synthesis of Compound 12**

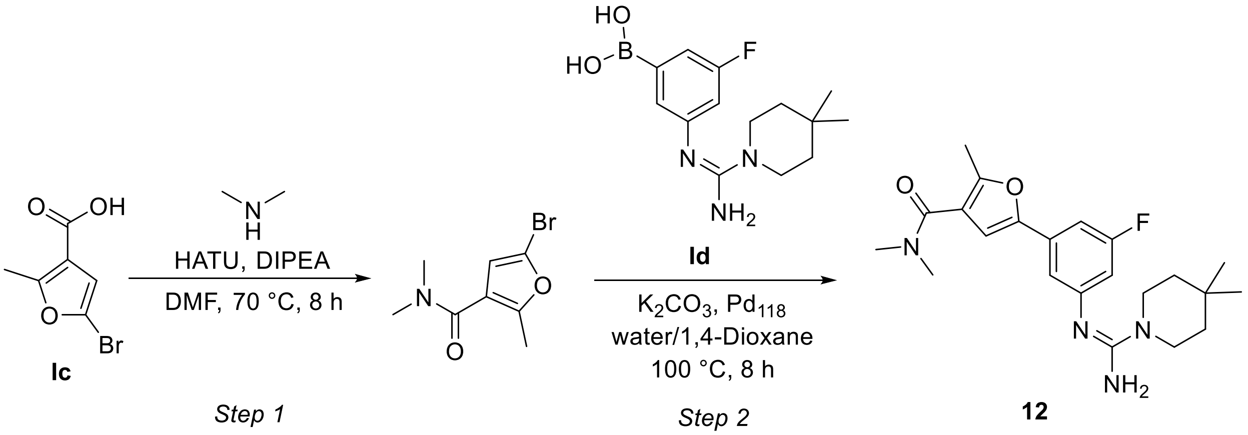


**Step 1:**

To a stirred solution of 5-bromo-2-methylfuran-3-carboxylic acid (300 mg, 1.47 mmol, 1.0 eq.) in DMF (4 mL), HATU (843 mg, 2.21 mmol, 1.5 eq.) was added followed by the addition of DIPEA (0.77 mL, 4.43 mmol, 3.0 eq.). The resulting mixture was stirred at room temperature for 30 min. Upon completion, dimethyl amine (0.298 mL, 4.43 mmol, 3.0 eq.) was added and the reaction was stirred at 70 °C for 8 h. Upon completion, the reaction mixture was quenched with ice-cold water and extracted with ethyl acetate, dried over anhydrous sodium sulfate, concentrated and purified by column chromatography (EtOAc/Hex) to afford 5-bromo-N,N,2-trimethylfuran-3-carboxamide (200 mg, 58% yield).

LCMS (ESI) $[M+H]^+$= 232.2

**Step 2:**

A stirred solution of 5-bromo-N,N,2-trimethylfuran-3-carboxamide (200 mg, 0.86 mmol, 1.0 eq.) and (Z)-(3-((amino(4,4-dimethylpiperidin-1-yl)methylene)amino)-5-fluorophenyl)boronic acid **Id** (311 mg, 0.95 mmol, 1.1 eq.) in 1,4-dioxane/$H_2O$ (2:1, 5 mL) was degassed with argon for 15 min.

Potassium carbonate (358 mg, 2.58 mmol, 3.0 eq.) was then added followed by the addition of $Pd_{118}$ (56 mg,0.086 mmol, 0.1 eq.) and the reaction was stirred at 100 °C for 8 h in a sealed tube. Upon completion, the reaction mixture was filtered through Celite, the filtrate concentrated under reduced pressure and crude residue purified by prepHPLC (aq. $NH_4HCO_3$/ACN) to give (Z)-5-(3-((amino(4,4-dimethylpiperidin-1-yl)methylene)amino)-5-fluorophenyl)-N,N,2-trimethylfuran-3-carboxamide (40 mg, 11% yield) as an off white solid.

LCMS (ESI) $[M+H]^+$= 401.3

$^1$H NMR (400 MHz, methanol-d4) δ 7.10 – 7.02 (m, 2H), 6.85 (s, 1H), 6.65 – 6.58 (m, 1H), 3.48 – 3.41 (m, 4H), 3.12 (s, 3H), 3.07 (s, 3H), 2.40 (s, 3H), 1.49 – 1.41 (m, 4H), 1.02 (s, 6H).

**Synthesis of Compound 13**

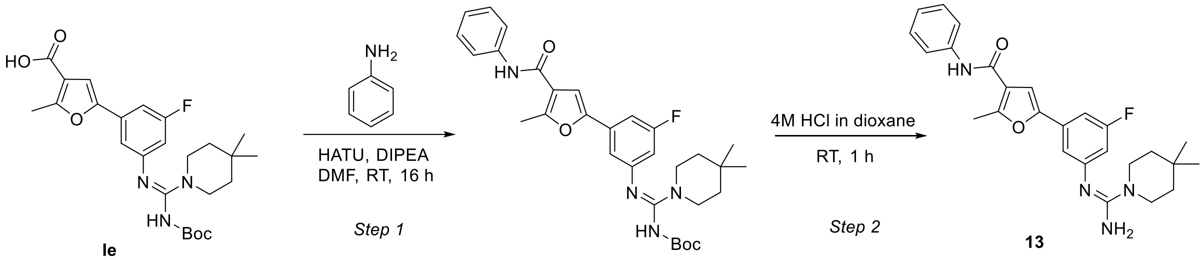


**Step 1:**

To a stirred solution of (Z)-5-(3-((((tert-butoxycarbonyl)amino)(4,4-dimethylpiperidin-1-yl)methylene)amino)-5-fluorophenyl)-2-methylfuran-3-carboxylic acid **Ie** (100 mg, 0.21 mmol, 1.0 eq.) in anhydrous DMF (3 mL) at 0 °C were added HATU (121 mg, 0.32 mmol, 1.5 eq.) and DIPEA (0.11 mL, 0.634 mmol, 3.0 eq.). After stirring for 15 min, aniline (0.02 mL, 0.23 mmol, 1.2 eq.) was added. The reaction mixture was allowed to warm to room temperature and stirred for 16 h. Upon completion, the reaction mixture was diluted with water and extracted with ethyl acetate. The combined organic layers were washed with brine, dried over anhydrous sodium sulfate, filtered and concentrated under reduced pressure. The resulting crude residue was purified by silica gel column chromatography (EtOAc/Hex) to afford tert-butyl (Z)-((4,4-dimethylpiperidin-1-yl)((3-fluoro-5-(5-methyl-4-(phenylcarbamoyl)furan-2-yl)phenyl)imino)methyl)carbamate (107 mg, 92% yield). LCMS (ESI) $[M+H]^+$= 549.3

**Step 2:**

To tert-butyl (Z)-((4,4-dimethylpiperidin-1-yl)((3-fluoro-5-(5-methyl-4-(phenylcarbamoyl)furan-2-yl)phenyl)imino)methyl)carbamate (107 mg, 0.18 mmol, 1.0 eq.) at 0°C was added 4M HCl in dioxane (1 mL). The reaction mixture was allowed to warm to room temperature and stirred for 1 h. Upon completion, the reaction mixture was concentrated under reduced pressure. The resulting crude residue was purified by prepHPLC (aq. $NH_4HCO_3$/ACN) to afford (Z)-5-(3-((amino(4,4-dimethylpiperidin-1-yl)methylene)amino)-5-fluorophenyl)-2-methyl-N-phenylfuran-3-carboxamide (15 mg, 18% yield) as a white powder.

$^1$H NMR (400 MHz, DMSO-$d_6$) δ 9.72 (s, 1H), 7.73 (d, *J* = 8.0 Hz, 2H), 7.48 (s, 1H), 7.34 (t, *J* = 7.7 Hz, 2H), 7.09 (t, *J* = 7.4 Hz, 1H), 6.86 – 6.81 (m, *J* = 7.5 Hz, 2H), 6.42 (d, *J* = 11.1 Hz, 1H), 5.52 (s, 2H), 3.39 – 3.34 (m, 4H), 2.62 (s, 3H), 1.35 – 1.28 (m, 4H), 0.96 (s, 6H).

LCMS (ESI) $[M+H]^+$= 449.5

**Synthesis of Compound 14**

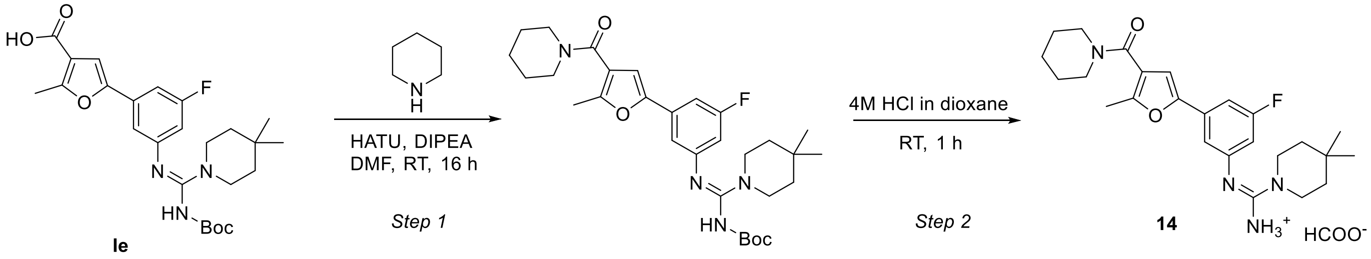


**Step 1:**

To a stirred solution of (Z)-5-(3-((((tert-butoxycarbonyl)amino)(4,4-dimethylpiperidin-1-yl)methylene)amino)-5-fluorophenyl)-2-methylfuran-3-carboxylic acid **Ie** (150 mg, 0.32 mmol, 1.0 eq.) in anhydrous DMF (3 mL) at 0 °C were added HATU (180 mg, 0.47 mmol, 1.5 eq.) and DIPEA (0.17 mL, 0.95 mmol, 3.0 eq.). After 5 min, piperidine (0.04 mL, 0.35 mmol, 1.1 eq.) was added, and the reaction stirred at room temperature for 16 h. Upon completion, the reaction mixture was diluted with water and extracted with ethyl acetate. The combined organic layers were washed with brine, dried over anhydrous sodium sulfate, filtered and concentrated under reduced pressure. The resulting crude residue was purified by a column chromatography (EtOAc/Hex) to afford (Z)-N'-(3-fluoro-5-(5-methyl-4-(piperidine-1-carbonyl)furan-2-yl)phenyl)-4,4-dimethylpiperidine-1-carboximidamide (120 mg, 72% yield).

LCMS (ESI) $[M+H]^+$= 540.9

**Step 2:**

Tert-butyl (Z)-((4,4-dimethylpiperidin-1-yl)((3-fluoro-5-(5-methyl-4-(piperidine-1- carbonyl)furan-2-yl)phenyl)imino)methyl)carbamate (120 mg, 0.23 mmol, 1.0 eq.) was dissolved in 4M HCl in dioxane (1.0 mL) and the resulting mixture was stirred at room temperature for 3 h. Upon completion, the resulting mixture was concentrated and purified by prepHPLC (ACN/water + 0.1% FA) to afford (Z)-N'-(3-fluoro-5-(5-methyl-4-(piperidine-1- carbonyl)furan-2-yl)phenyl)-4,4-dimethylpiperidine-1-carboximidamide formate (25 mg, 25% yield) as a white powder.

$^1$H NMR (500 MHz, methanol-d4) δ 7.38 (t, *J* = 1.7 Hz, 1H), 7.34 (ddd, *J* = 9.6, 2.4, 1.4 Hz, 1H), 6.94 (dt, *J* = 9.6, 2.1 Hz, 1H), 6.91 (s, 1H), 3.74-3.62 (m, 2H), 3.62–3.50 (m, 6H), 2.41 (s, 3H), 1.77 – 1.70 (m, 2H), 1.69-1.58 (m, 4H), 1.58 – 1.52 (m, 4H), 1.08 (s, 6H).

LCMS (ESI) $[M+H]^+$= 441.2

**Synthesis of Compound 15**

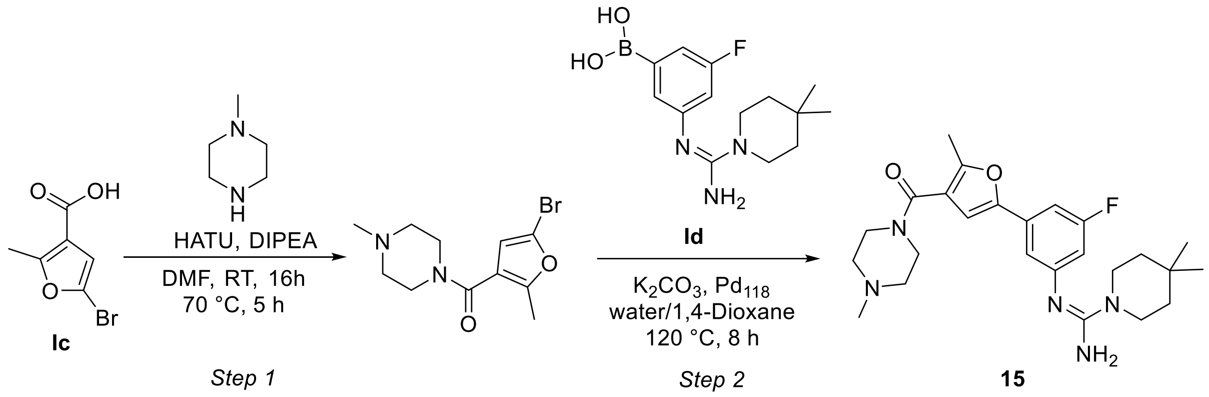


**Step 1:**

To a stirred solution of 5-bromo-2-methylfuran-3-carboxylic acid **Ic** (350 mg, 1.72 mmol, 1.0 eq.) in anhydrous DMF (4 mL) were added HATU (984 mg, 2.58 mmol, 1.5 eq.) and DIPEA (0.9 mL, 5.16 mmol, 3.0 eq.). The reaction mixture was stirred at room temperature for 30 min. 1-methylpiperazine (0.576 mL, 5.17 mmol, 3.0 eq.) was then added, and the reaction was allowed to stir at room temperature for an additional 16 h. To drive the reaction to completion, the mixture was heated at 70°C and stirred for a further 5 h. Upon completion, the reaction was quenched with ice-cold water and extracted with ethyl acetate. The combined organic layers were dried over anhydrous sodium sulfate, filtered and concentrated under reduced pressure. The resulting crude residue was purified by silica gel column chromatography (EtOAc/Hex) to afford (5-bromo-2-methylfuran-3-yl)(4-methylpiperazin-1-yl)methanone (260 mg, 52% yield).

LCMS (ESI) $[M+H]^{+}$= 286.9

**Step 2:**

A stirred solution of (5-bromo-2-methylfuran-3-yl)(4-methylpiperazin-1-yl)methanone (120 mg, 0.418 mmol, 1.0 eq.) and (Z)-(3-((amino(4,4-dimethylpiperidin-1-yl)methylene)amino)-5-fluorophenyl)boronic acid **Id** (151 mg, 0.46 mmol, 1.1 eq.) in 1,4- Dioxane/$H_2O$ (2:1, 3 mL) was degassed with argon for 10 min. Potassium carbonate (173 mg, 1.25 mmol, 3.0 eq.) and $Pd_{118}$ (27 mg, 0.041 mmol) were then added, and the reaction was stirred at 120 °C for 8 h in a sealed tube. Upon completion, the reaction mixture was filtered through Celite, concentrated and purified by prepHPLC (aq. $NH_4HCO_3$/ACN) to afford (Z)-N'-(3-fluoro-5-(5-methyl-4-(4-methylpiperazine-1-carbonyl)furan-2-yl)phenyl)-4,4-dimethylpiperidine-1-carboximidamide (38 mg, 19% yield) as a light brown solid.

LCMS (ESI) $[M+H]^{+}$= 456.3

$^{1}$H NMR (400 MHz, DMSO-$d_6$) δ 6.98 (s, 1H), 6.96 – 6.91 (m, 1H), 6.88 – 6.85 (m, 1H), 6.41 (d, *J* = 11.0 Hz, 1H), 5.54 (s, 2H), 3.62 – 3.44 (m, 4H), 3.38 – 3.33 (m, 4H), 2.36 (s, 3H), 2.35 – 2.29 (m, 4H), 2.20 (s, 3H), 1.34 – 1.28 (m, 4H), 0.95 (s, 6H).

**Synthesis of Compound 16**

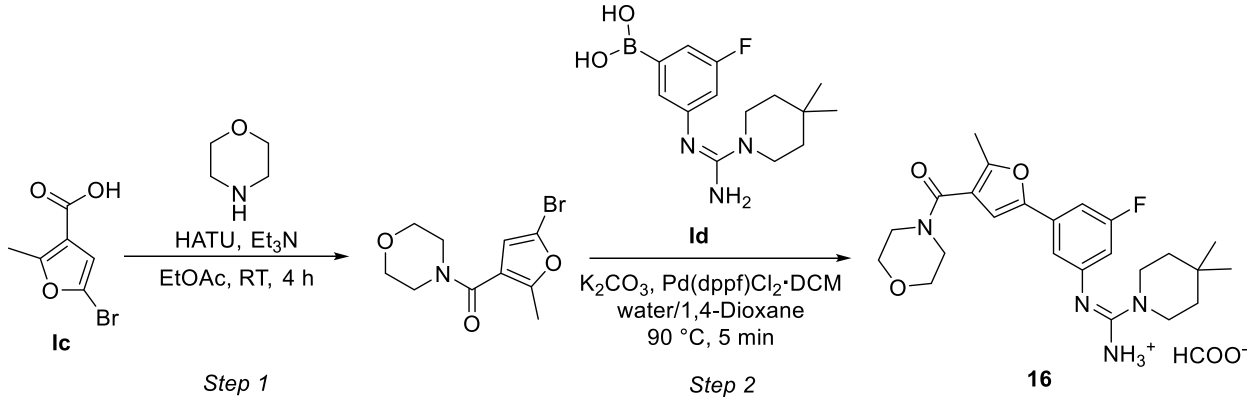

**Step 1:**

To a stirring solution of 5-bromo-2-methylfuran-3-carboxylic acid **Ic** (50.0 mg, 0.24 mmol, 1.0 eq.) and HATU (111.3 mg, 0.29 mmol, 1.2 eq.) in EtOAc (2.0 mL) were added morpholine (0.026 mL, 0.29 mmol, 1.2 eq.) and triethylamine (0.170 mL, 1.22 mmol, 5.0 eq.). The reaction mixture was stirred at room temperature for 4 h. Upon completion, the resulting solution was diluted with $NaHCO_3$ and extracted with ethyl acetate. The organic layers were dried over anhydrous sodium sulfate, filtered and concentrated under reduced pressure to afford 4-(5-bromo-2-methylfuran-3-carbonyl)morpholine (50 mg, 75% yield). The crude product was utilized in the next step without further purification.

LCMS (ESI) $[M+H]^+$= 275.9

**Step 2:**

To the stirred solution of 4-(5-bromo-2-methylfuran-3-carbonyl)morpholine (45.0 mg, 0.16 mmol, 1.0 eq.) in a mixture of 1,4-dioxane/$H_2O$ (2:1, 3.0 mL) were added (Z)-(3-((amino(4,4-dimethylpiperidin-1-yl)methylene)amino)-5-fluorophenyl)boronic acid **Ic** (57.8 mg, 0.20 mmol, 1.2 eq.), Pd(dppf)$Cl_2$•DCM (13.4 mg, 0.016 mmol, 0.1 eq.) and potassium carbonate (45.4 mg, 0.33 mmol, 2.0 eq.). The reaction mixture was heated at 90 °C and stirred for 5 min. The resulting mixture was filtered through Celite and purified by reversed-phase silica gel column chromatography (ACN/water + 0.1% FA) to afford (Z)-N'-(3-fluoro-5-(5-methyl-4-(morpholine-4-carbonyl)furan-2-yl)phenyl)-4,4-dimethylpiperidine-1-carboximidamide formate (65.0 mg, 0.147 mmol, 89.5%) as a beige gum.

LCMS (ESI) $[M+H]^+$= 443.2

$^1$H NMR (500 MHz, methanol-d4) δ 8.40 (s, 1H), 7.38 (t, *J* = 1.7 Hz, 1H), 7.35 (ddd, *J* = 9.5, 2.4, 1.4 Hz, 1H), 6.97 – 6.91 (m, 2H), 3.78 – 3.62 (m, 8H), 3.60 – 3.56 (m, 4H), 2.44 (s, 3H), 1.59 – 1.52 (m, 4H), 1.08 (s, 6H).

### Synthesis of Compound 17

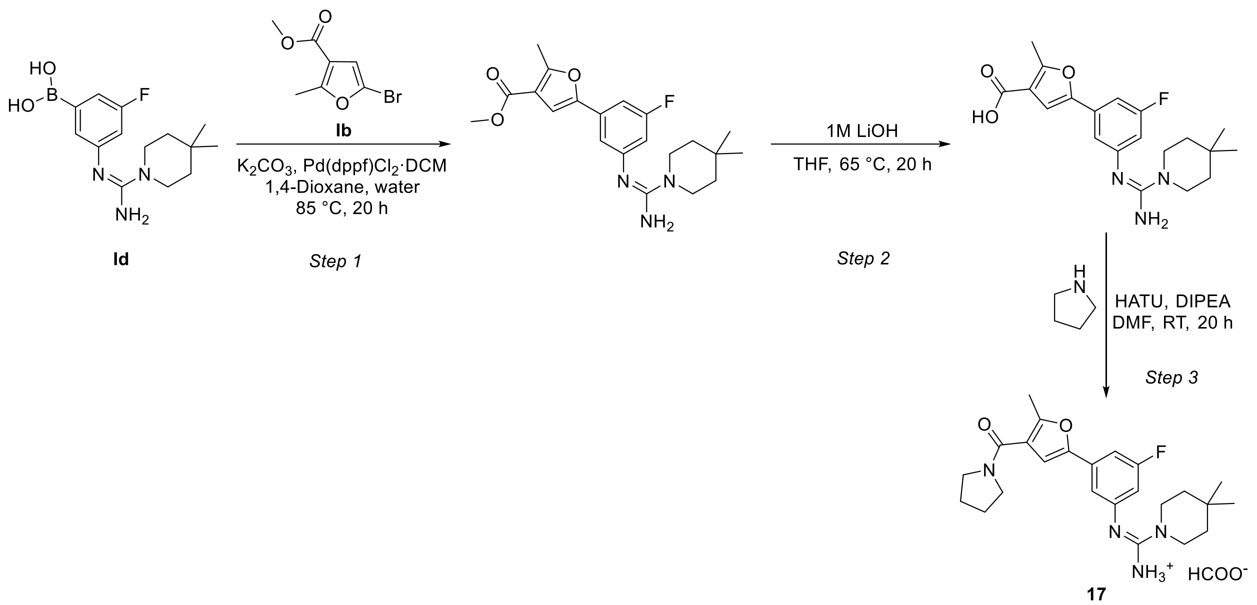


**Step 1:**

To a stirred solution of methyl 5-bromo-2-methylfuran-3-carboxylate **Ib** (10.0 mg, 0.046 mmol, 1.0 eq.) and (Z)-(3-((amino(4,4-dimethylpiperidin-1-yl)methylene)amino)-5-fluorophenyl)boronic acid **Id** (20.1 mg, 0.068 mmol, 1.5 eq.) in 1,4-dioxane/water (9:1, 1.0 mL) were added potassium carbonate (12.6 mg, 0.091 mmol, 2.0 eq.) and Pd(dppf)$Cl_2$·DCM (3.7 mg, 0.005 mmol, 0.1 eq.). The reaction mixture was heated at 85 °C and stirred for 20 h. Upon completion, the resulting mixture was filtered through Celite, concentrated under reduced pressure and purified by reversed-phase silica gel column chromatography (ACN/water + 0.1% FA) to afford methyl (Z)-5-(3-((amino(4,4-dimethylpiperidin-1-yl)methylene)amino)-5-fluorophenyl)-2-methylfuran-3-carboxylate (10.0 mg, 56.5% yield).

LCMS (ESI) $[M+H]^+$= 388.4

**Step 2:**

To a stirred solution of methyl (Z)-5-(3-((amino(4,4-dimethylpiperidin-1-yl)methylene)amino)-5-fluorophenyl)-2-methylfuran-3-carboxylate (72.0 mg, 0.186 mmol, 1.0 eq.) in THF (2.0 mL) was added 1M LiOH (10.0 mL). The reaction mixture was heated at 65 °C and stirred for 20 h. Upon completion, the solution was acidified to pH 1 with 1M HCl and extracted with ethyl acetate. The combined organic layers were dried over magnesium sulfate, filtrated and concentrated to anhydrousness to afford 5-{3-[(Z)-[amino(4,4-dimethylpiperidin-1-yl)methylidene]amino]-5-fluorophenyl}-2-methylfuran-3-carboxylic acid (30.0 mg, 43.2% yield).

LCMS (ESI) $[M+H]^+$=372.1

**Step 3:**

To a stirred solution of 5-{3-[(Z)-[amino(4,4-dimethylpiperidin-1-yl)methylidene]amino]-5-fluorophenyl}-2-methylfuran-3-carboxylic acid (15.0 mg, 0.040 mmol, 1.0 eq.) in anhydrous DMF (0.5 mL) were added pyrrolidine (0.004 mL, 0.044 mmol, 1.1 eq.) and DIPEA (0.01 mL, 0.060 mmol, 1.5 eq.). The reaction mixture was stirred at room temperature for 10 min. HATU (18.3 mg, 0.048 mmol, 1.2 eq.) was then added and the reaction was stirred at room temperature for an additional 20 h. Upon completion, the solution was concentrated under reduced pressure and purified by prepHPLC (ACN/water + 0.1% FA) to afford (Z)-N'-(3-fluoro-5-(5-methyl-4-(pyrrolidine-1-carbonyl)furan-2-yl)phenyl)-4,4-dimethylpiperidine-1-carboximidamide formate (5.5 mg, 28.6% yield, FA salt) as a white solid.

LCMS (ESI) $[M+H]^+$=427.2

$^1$H NMR (500 MHz, DMSO-$d_6$) δ 8.29 (s, 1H), 7.18 (s, 1H), 7.00 (dt, *J* = 10.0, 1.9 Hz, 1H), 6.94 (t, *J* = 1.6 Hz, 1H), 6.48 (dt, *J* = 11.0, 2.1 Hz, 1H), 5.96 (s, 2H), 3.58 – 3.44 (m, 8H), 2.44 (s, 3H), 1.89 – 1.79 (m, 4H), 1.37 – 1.29 (m, 4H), 0.96 (s, 6H).

**Synthesis of Compound 18**

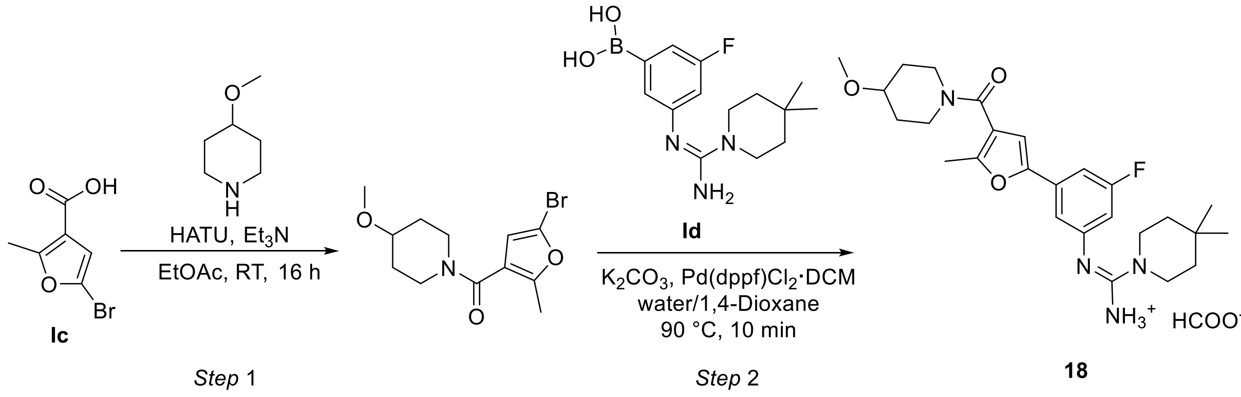


**Step 1:**

To a stirred solution of 4-methoxypiperidine (33.7 mg, 0.29 mmol, 1.2 eq.) in ethyl acetate (2.0 mL) were added 5-bromo-2-methylfuran-3-carboxylic acid **Ic** (50.0 mg, 0.24 mmol, 1.0 eq.), HATU (111.3 mg, 0.29 mmol, 1.2 eq.) and triethylamine (0.170 mL, 1.22 mmol, 5.0 eq.). The reaction mixture was stirred at room temperature overnight. The resulting solution was diluted with saturated solution of sodium bicarbonate and extracted with ethyl acetate. The combined organic layers were dried over anhydrous sodium sulfate, filtrated and concentrated under reduced pressure to afford 1-(5-bromo-2-methylfuran-3-carbonyl)-4-methoxypiperidine (60 mg, 81.4% yield). The product was utilized in the next step without further purification.

LCMS (ESI) $[M+H]^+$= 303.9

**Step 2:**

1-(5-bromo-2-methylfuran-3-carbonyl)-4-methoxypiperidine (60.0 mg, 0.20 mmol, 1.0 eq.) and (Z)-(3-((amino(4,4-dimethylpiperidin-1-yl)methylene)amino)-5-fluorophenyl)boronic acid **Id** (64.0 mg, 0.22 mmol, 1.1 eq.) were dissolved in 1,4-dioxane/water (2:1, 3.0 mL). The reaction vessel was evacuated and backfilled with argon (3x) and then Pd(dppf)$Cl_2$·DCM (16.2 mg, 0.020 mmol, 0.1 eq.) and potassium carbonate (54.9 mg, 0.40 mmol, 2.0 eq.) were added. The reaction mixture was heated at 90 °C and stirred for 10 min. Upon completion, the resulting solution was filtered through Celite, concentrated under reduced pressure and purified by prepHPLC (ACN/water + 0.1% FA) to afford (Z)-N'-{3-fluoro-5-[4-(4-methoxypiperidine-1-carbonyl)-5-methylfuran-2-yl]phenyl}-4,4-dimethylpiperidine-1-carboximidamide formate (6.0 mg, 6.4%) as a grey solid.

LCMS (ESI) $[M+H]^+$= 471.2

$^1$H NMR (500 MHz, methanol-d4) δ 7.38 (t, *J* = 1.7 Hz, 1H), 7.35 (ddd, *J* = 9.6, 2.4, 1.4 Hz, 1H), 6.96 – 6.90 (m, 2H), 4.04 – 3.89 (m, 1H), 3.84 – 3.70 (m, 1H), 3.61 – 3.56 (m, 4H), 3.56 – 3.51 (m, 1H), 3.49 – 3.41 (m, 2H), 3.38 (s, 3H), 2.42 (s, 3H), 1.99 – 1.88 (m, 2H), 1.65 – 1.57 (m, 2H), 1.57 – 1.53 (m, 4H), 1.08 (s, 6H).

### Synthesis of Compound 19

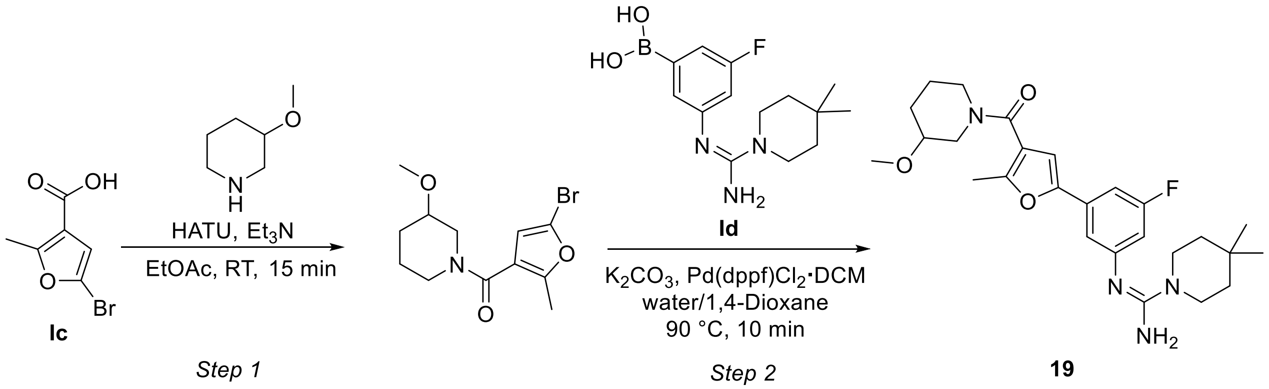


**Step 1:**

To a stirred solution of 3-methoxypiperidine (racemic mixture, 56.3 mg, 0.49 mmol, 2.0 eq.) and 5-bromo-2-methylfuran-3-carboxylic acid **Ic** (50.0 mg, 0.24 mmol, 1.0 eq.) in ethyl acetate (2.0 mL) were added HATU (111.3 mg, 0.29 mmol, 1.2 eq.) and triethylamine (0.102 mL, 0.73 mmol, 3.0 eq.). The reaction mixture was stirred at room temperature for 15 min. Upon completion, the resulting solution was diluted with saturated solution of sodium bicarbonate and extracted with ethyl acetate. The combined organic layers were dried over anhydrous sodium sulfate, filtrated and concentrated under reduced pressure to afford 1-(5-bromo-2-methylfuran-3-carbonyl)-3-methoxypiperidine (50 mg, 67.8% yield). The crude product was utilized in the next step without further purification.

LCMS (ESI) $[M+H]^+$= 303.9

**Step 2:**

(Z)-(3-((amino(4,4-dimethylpiperidin-1-yl)methylene)amino)-5-fluorophenyl)boronic acid **Id** (53.4 mg, 0.182 mmol, 1.1 eq.) and 1-(5-bromo-2-methylfuran-3-carbonyl)-3-methoxypiperidine (50.0 mg, 0.165 mmol, 1.0 eq.) were dissolved in 1,4-dioxane/water (2:1, 3.0 mL). The reaction vessel was evacuated and backfilled with argon (3x) and then potassium carbonate (45.7 mg, 0.33 mmol, 2.0 eq.) and Pd(dppf)$Cl_2$·DCM (13.5 mg, 0.017 mmol, 0.1 eq.) were added. The resulting mixture was stirred at 90 °C for 10 min. Upon completion, the mixture was filtered through Celite, concentrated under reduced pressure and purified by reversed-phase silica gel column chromatography (ACN/water + 0.1 % FA) to afford (Z)-N'-{3-fluoro-5-[4-(3-methoxypiperidine-1-carbonyl)-5-methylfuran-2-yl]phenyl}-4,4-dimethylpiperidine-1-carboximidamide (12.0 mg, 15.4% yield) as a beige solid.

LCMS (ESI) $[M+H]^{+}$= 471.2

$^{1}H$ NMR (500 MHz, DMSO-$d_6$, 353K) δ 7.32 – 7.23 (m, 2H), 7.01 (s, 1H), 6.88 (dt, *J* = 10.2, 2.1 Hz, 1H), 3.75 – 3.68 (m, 1H), 3.54 – 3.49 (m, 4H), 3.49 – 3.45 (m, 2H), 3.41 (dd, *J* = 13.1, 6.6 Hz, 1H), 3.32 – 3.27 (m, 1H), 3.25 (s, 3H), 2.38 (s, 3H), 1.93 – 1.85 (m, 1H), 1.78 – 1.70 (m, 1H), 1.66 – 1.59 (m, 1H), 1.50 – 1.39 (m, 5H), 1.02 (s, 6H).

**Synthesis of Compound 20**

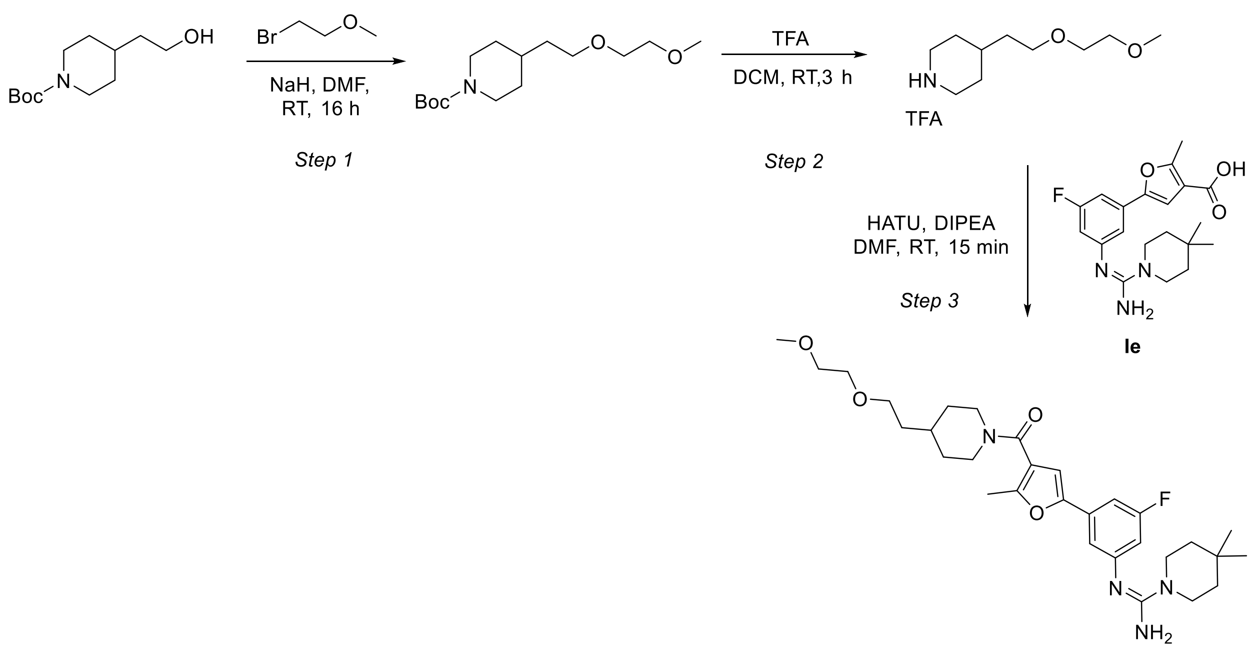


20

**Step 1:**

To a stirred solution of tert-butyl 4-(2-hydroxyethyl)piperidine-1-carboxylate (2.0 g, 8.73 mmol, 1.0 eq.) in anhydrous DMF (5.0 mL) at room temperature was added NaH (60% dispersion in mineral oil, 530 mg, 13.1 mmol, 1.5 eq.) After stirring for 15 min, a solution of 1-bromo-2-methoxyethane (1.8 g, 13.1 mmol, 1.5 eq.) in anhydrous DMF (2.0 mL) was added dropwise. The reaction mixture was stirred at room temperature for 16 h. Upon completion, the mixture was quenched with ice-cold water and extracted with ethyl acetate. The organic layers were washed with brine solution, dried over anhydrous anhydrous sodium sulfate, filtrated and concentrated under reduced pressure. The crude product was purified by silica gel column chromatography

(EtOAc/Hex) to afford tert-butyl 4-(2-(2-methoxyethoxy)ethyl)piperidine-1-carboxylate (2.0 g, 80% yield).

LCMS (ESI) $[M+H]^+$= 287.2

**Step 2:**

A mixture of tert-butyl 4-(2-(2-methoxyethoxy)ethyl)piperidine-1-carboxylate (500 mg, 1.74 mmol, 1.0 eq.) and TFA (2.5 mL) in DCM (2.5 mL) was stirred at room temperature for 3 h. Upon completion, the resulting solution was concentrated to anhydrousness. The crude product was triturated with diethyl ether and dried to afford 4-(2-(2-methoxyethoxy)ethyl)piperidine (350 mg, 66% yield, TFA salt). The crude product was utilized in the next step without further purification.

LCMS (ESI) $[M+H]^+$= 187.16

**Step 3:**

To a stirred solution of 5-{3-[(Z)-[amino(4,4-dimethylpiperidin-1-yl)methylidene]amino]-5-fluorophenyl}-2-methylfuran-3-carboxylic acid **Ie** (15.0 mg, 0.04 mmol, 1.0 eq.) and 4-[2-(2-methoxyethoxy)ethyl]piperidine (9.0 mg, 0.05 mmol, 1.2 eq.) in anhydrous DMF (1.0 mL) under an argon atmosphere were added HATU (22.9 mg, 0.060 mmol, 1.5 eq.) and DIPEA (0.035 mL, 0.201 mmol, 5.0 eq.). The reaction mixture was stirred at room temperature for 15 min. The resulting solution was filtered through a syringe filter and concentrated under reduced pressure. The resulting crude residue was purified by prepHPLC (ACN/water + 0.1% FA) to afford (Z)-N'-[3-fluoro-5-(4-{4-[2-(2-methoxyethoxy)ethyl]piperidine-1-carbonyl}-5-methylfuran-2-yl)phenyl]-4,4-dimethylpiperidine-1-carboximidamide (8.5 mg, 39.0% yield) as a light pink solid.

LCMS (ESI) $[M+H]^+$= 543.2

$^1$H NMR (500 MHz, DMSO-$d_6$) δ 7.21 – 7.15 (m, 1H), 7.11 (s, 1H), 7.04 (s, 1H), 6.73 – 6.67 (m, 1H), 4.51 –4.21 (m, 1H), 3.92 – 3.67 (m, 1H), 3.48 – 3.44 (m, 4H), 3.43 – 3.40 (m, 6H), 3.24 (s, 3H), 3.12 – 2.94 (m, 1H), 2.84 – 2.65 (m, 1H), 2.36 (s, 3H), 1.74 – 1.62 (m, 3H), 1.47 (q, *J* = 6.6 Hz, 2H), 1.41 – 1.34 (m, 4H), 1.14 – 1.05 (m, 2H), 0.97 (s, 6H).

**Synthesis of Compound 21**

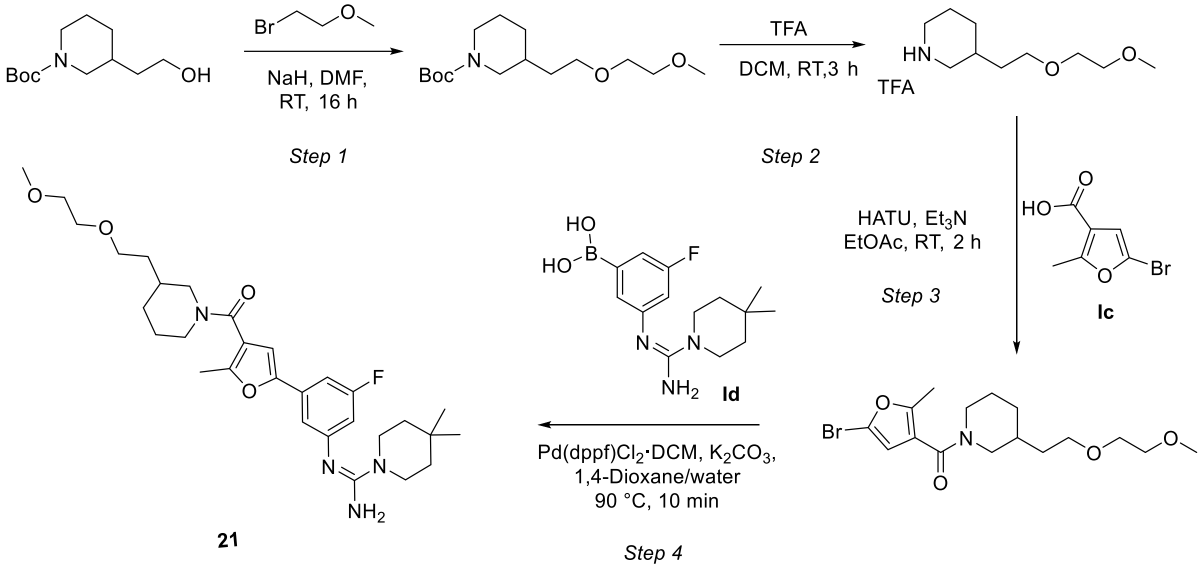


**Step 1:**

To a stirred solution of tert-butyl 3-(2-hydroxyethyl)piperidine-1-carboxylate (racemic mixture, 1.0 g, 4.37 mmol, 1.0 eq.) in anhydrous DMF (5.0 mL) was added NaH (60% dispersion in mineral oil, 0.191 g, 4.80 mmol, 1.1 eq.) The reaction mixture was stirred at room temperature for 15 min, followed by the dropwise addition of 1-bromo-2-methoxyethane (0.723 g, 5.24 mmol, 1.2 eq.) in DMF (2.5 mL). Stirring was continued at room temperature for 16 h. Upon completion, the reaction mixture was quenched with ice-cold water and extracted with ethyl acetate. The organic layers were washed with brine, dried over anhydrous anhydrous sodium sulfate, filtrated and concentrated under reduced pressure. The crude product was purified by silica gel column chromatography (EtOAc/Hex) to afford tert-butyl 3-(2-(2-methoxyethoxy)ethyl)piperidine-1-carboxylate (1.5 g, 67% yield).

LCMS (ESI) $[M+H]^+$= 287.2

**Step 2:**

To a stirring solution of tert-butyl 3-(2-(2-methoxyethoxy)ethyl)piperidine-1-carboxylate (370 mg, 1.287 mmol, 1.0 eq.) in DCM (2.0 mL) was added TFA (2.0 mL). The reaction mixture was stirred at room temperature for 3 h. Upon completion, the resulting solution was concentrated under reduced pressure. The resulting residue was triturated with diethyl ether and dried *in vacuo* to afford 3-(2-(2-methoxyethoxy)ethyl)piperidine (120 mg, 31% yield, TFA salt). The crude product was utilized in the next step without further purification. LCMS (ESI) $[M+H]^+$= 187.16

**Step 3**

To a stirring solution of of 3-[2-(2-methoxyethoxy)ethyl]piperidine (50.2 mg, 0.27 mmol, 1.1 eq.) and 5-bromo-2-methylfuran-3-carboxylic acid **Ic** (50.0 mg, 0.24 mmol, 1.0 eq.) in ethyl acetate (2.0 mL) were added HATU (102.0 mg, 0.268 mmol, 1.1 eq.) and triethylamine (0.102 mL, 0.732 mmol, 3.0 eq.). The reaction mixture was stirred at room temperature for 2 h. Upon completion, the resulting mixture was diluted with brine and extracted with ethyl acetate. The organic layers were dried over magnesium sulfate, filtered and concentrated under reduced pressure. The resulting crude residue was purified by reversed-phase silica gel column chromatography (ACN/water + 0.1% FA) to afford 1-(5-bromo-2-methylfuran-3-carbonyl)-3-[2-(2-methoxyethoxy)ethyl]piperidine (60.0 mg, 65.7% yield) as an off-white solid.

LCMS (ESI) $[M+H]^+$= 375.8

**Step 4:**

1-(5-bromo-2-methylfuran-3-carbonyl)-3-[2-(2-methoxyethoxy)ethyl]piperidine (60.0 mg, 0.16 mmol, 1.0 eq.) and (Z)-(3-((amino(4,4-dimethylpiperidin-1-yl)methylene)amino)-5-fluorophenyl)boronic acid **Id** (51.7 mg, 0.18 mmol, 1.1 eq.) were dissolved in 1,4-dioxane/water (2:1, 3.0 mL). The reaction vessel was evacuated and backfilled with argon (3x) and then Pd(dppf)$Cl_2$·DCM (13.1 mg, 0.016 mmol, 0.1 eq.) and potassium carbonate (44.3 mg, 0.32 mmol, 2.0 eq.) were added. The reaction mixture was heated at 90 °C and stirred for 10 min. Upon completion, the mixture was filtered through Celite, concentrated under reduced pressue and purified by reversed-phase silica gel column chromatography (ACN/water + 0.1 % FA) to afford (Z)-N'-[3-fluoro-5-(4-{3-[2-(2-methoxyethoxy)ethyl]piperidine-1-carbonyl}-5-methylfuran-2-yl)phenyl]-4,4-dimethylpiperidine-1-carboximidamide (15.0 mg, 17.2% yield) as an off-white solid.

LCMS (ESI) $[M+H]^+$= 543.25 m/z

$^1$H NMR (500 MHz, DMSO-$d_6$) δ 7.28 – 7.20 (m, 2H), 6.99 – 6.96 (m,1H), 6.84 (dt, *J* = 10.3, 2.1 Hz, 1H), 4.14 – 3.89 (m, 2H), 3.53 – 3.48 (m, 5H), 3.48 – 3.44 (m, 3H), 3.44 – 3.40 (m, 2H), 3.28 – 3.23 (m, 3H), 2.95 – 2.87 (m, 1H), 2.76 (dd, *J* = 13.0, 10.0 Hz, 1H), 2.40 – 2.34 (m, 3H), 1.89 – 1.81 (m, 1H), 1.73 – 1.67 (m, 1H), 1.66 – 1.57*J* (m, 1H), 1.51 (q, *J* = 6.5 Hz, 1H), 1.48 – 1.38 (m, 5H), 1.30 – 1.22 (m, 1H), 1.20 – 1.08 (m, 1H), 1.01 (s, 6H).

### Synthesis of Compound 22

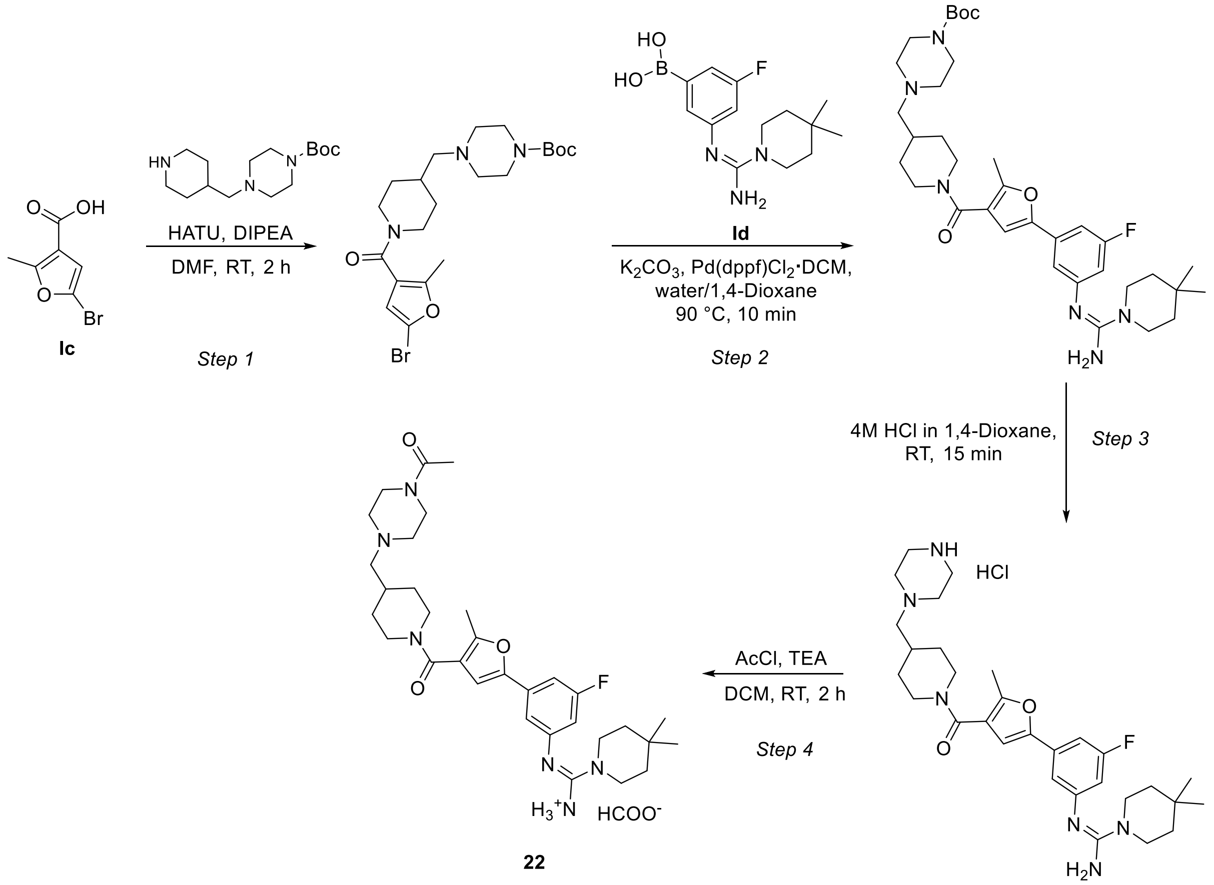

**Step 1:**

To a mixture of tert-butyl 4-[(piperidin-4-yl)methyl]piperazine-1-carboxylate (38.0 mg, 0.13 mmol, 1.1 eq.), 5-bromo-2-methylfuran-3-carboxylic acid **Ic** (25.0 mg, 0.12 mmol, 1.0 eq.) and HATU (51.0 mg, 0.13 mmol, 1.1 eq.) in ethyl acetate (2.0 mL) was added DIPEA (0.170 mL, 1.219 mmol, 5.0 eq.). The reaction mixture was stirred at room temperature for 2 h. The resulting solution was diluted with sodium bicarbonate and extracted with ethyl acetate. The combined organic layers were dried over anhydrous sodium sulfate, filtrated and concentrated to anhydrousness to afford tert-butyl 4-{[1-(5-bromo-2-methylfuran-3-carbonyl)piperidin-4-yl]methyl}piperazine-1-carboxylate (40 mg, 70% yield). The crude product was utilized in the next step without further purification.

LCMS (ESI) $[M+H]^+$= 472.1

**Step 2:**

The a mixture of tert-butyl 4-{[1-(5-bromo-2-methylfuran-3-carbonyl)piperidin-4-yl]methyl}piperazine-1-carboxylate (100.0 mg, 0.21 mmol, 1.0 eq.), (Z)-(3-((amino(4,4-dimethylpiperidin-1-yl)methylene)amino)-5-fluorophenyl)boronic acid **Id** (68.6 mg, 0.23 mmol, 1.1 eq.), Pd(dppf)$Cl_2$·DCM (8.7 mg, 0.011 mmol, 0.05 eq.) and potassium carbonate (58.8 mg, 0.43 mmol, 2.0 eq.) was dissolved in a mixture of 1,4-dioxane/$H_2O$ (2:1, 7.5 mL). The reaction mixture was heated at 90 °C and stirred for 10 min. Upon completion, the mixture was filtered through a pad of Celite, concentrated under reduced pressure and purified by prepHPLC (ACN/water + 0.1% FA) to afford tert-butyl 4-{[1-(5-{3-[(Z)-[amino(4,4-dimethylpiperidin-1-yl)methylidene]amino]-5-fluorophenyl}-2-methylfuran-3-carbonyl)piperidin-4-yl]methyl}piperazine-1-carboxylate (41.5 mg, 30.6% yield) as a beige solid.

LCMS (ESI) $[M+H]^+$= 639.3

$^1H$ NMR (500 MHz, DMSO-$d_6$, 353K) δ 6.95 (ddd, *J* = 9.8, 2.4, 1.4 Hz, 1H), 6.92 (t, *J* = 1.7 Hz, 1H), 6.86 (s, 1H), 6.47 (dt, *J* = 11.0, 2.1 Hz, 1H), 3.42 – 3.38 (m, 4H), 3.34 – 3.29 (m, 4H), 2.93 (t, *J* = 12.5 Hz, 4H), 2.36 (s, 3H), 2.35 – 2.31 (m, 4H), 2.20 (d, *J* = 6.9 Hz, 2H), 1.86 – 1.73 (m, 3H), 1.41 (s, 9H), 1.39 – 1.35 (m, 4H), 1.17 – 1.05 (m, 2H), 0.98 (s, 6H).

**Step 3:**

Tert-butyl 4-{[1-(5-{3-[(Z)-[amino(4,4-dimethylpiperidin-1-yl)methylidene]amino]-5-fluorophenyl}-2-methylfuran-3-carbonyl)piperidin-4-yl]methyl}piperazine-1-carboxylate (42.3 mg, 0.066 mmol, 1.0 eq.) was dissolved in 4M HCl in dioxane (1.0 mL, 28.8 mmol) with the addition of water (3 drops). The reaction mixture was stirred at room temperature for 15 min. Upon completion, the resulting solution was concentrated to anhydrousness to afford (Z)-N'-[3-fluoro-5-(5-methyl-4-{4-[(piperazin-1-yl)methyl]piperidine-1-carbonyl}furan-2-yl)phenyl]-4,4-dimethylpiperidine-1-carboximidamide hydrochloride (38.0 mg, 99.8% yield, HCl salt), which was used in the subsequent step without further purification.

LCMS (ESI) $[M+H]^+$= 539.4

**Step 4:**

To a stirred solution of (Z)-N'-[3-fluoro-5-(5-methyl-4-{4-[(piperazin-1-yl)methyl]piperidine-1-carbonyl}furan-2-yl)phenyl]-4,4-dimethylpiperidine-1-carboximidamide hydrochloride (20.0 mg, 0.035 mmol, 1.0 eq.) in anhydrous DCM (1.0 mL) was added triethylamine (0.007 mL, 0.052 mmol, 1.5 eq.). The reaction mixture was cooled to 0°C and stirred for 5 min, followed by the addition of acetyl chloride (0.004 mL, 0.052 mmol, 1.5 eq.). The reaction mixture was allowed to warm to room temperature and stirred for 2 h under an argon atmosphere. The resulting solution was concentrated under reduced pressure and purified by prepHPLC (ACN/water + 0.1% FA) to afford (Z)-N'-[3-(4-{4-[(4-acetylpiperazin-1-yl)methyl]piperidine-1-carbonyl}-5-methylfuran-2-yl)-5-fluorophenyl]-4,4-dimethylpiperidine-1-carboximidamide formate (10.9 mg, 54.0% yield,) as a white solid.

$^1$H NMR (500 MHz, DMSO-$d_6$) δ 8.24 (s, 1H), 7.06 – 7.02 (m, 1H), 7.01 – 6.97 (m, 2H), 6.61 (dt, *J* = 10.8, 2.3 Hz, 1H), 4.32 – 4.05 (m, 2H), 3.23 – 2.81 (m, 10H), 2.37 – 2.32 (m, 2H), 2.30 – 2.25 (m, 2H), 2.18 (d, *J* = 7.1 Hz, 2H), 2.15 (s, 3H), 1.97 (s, 3H), 1.88 – 1.77 (m, 3H), 1.36 – 1.31 (m, 4H), 1.16 – 1.05 (m, 2H), 0.96 (s, 6H)

LCMS (ESI) $[M+H]^+$= 581.4

**Synthesis of Compound 23**

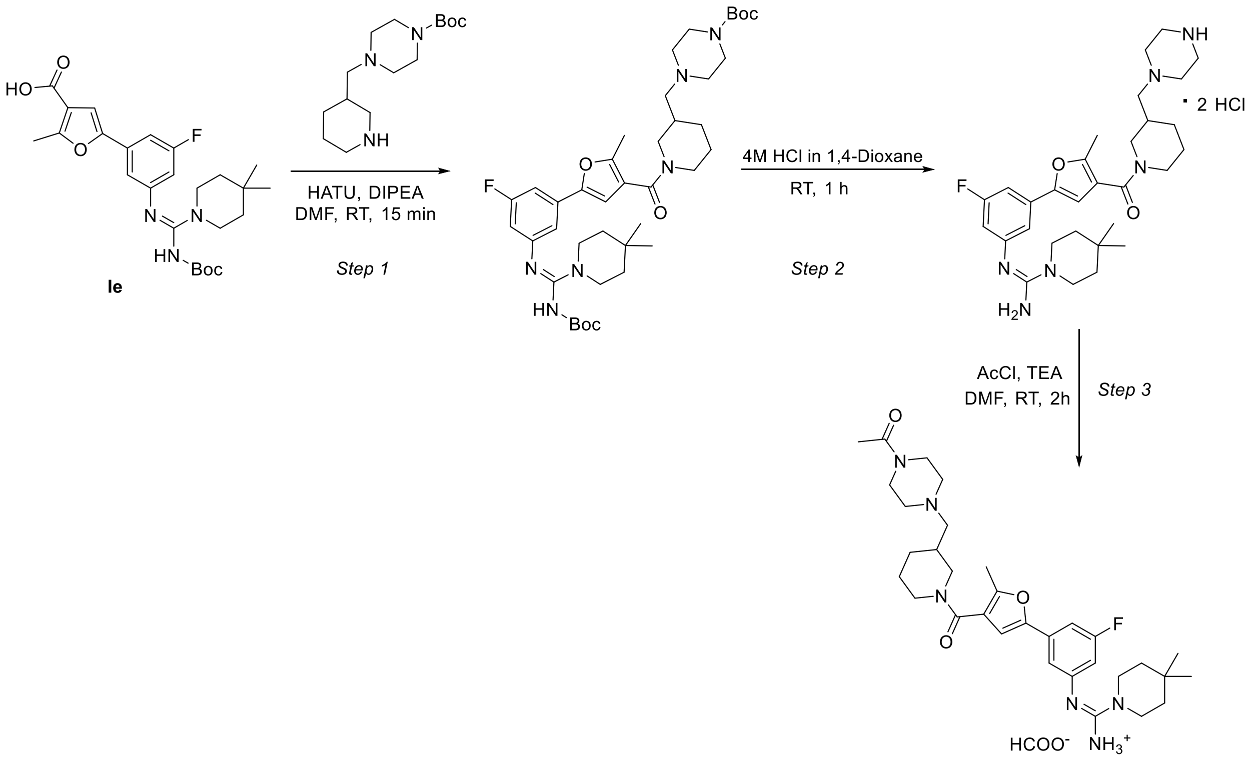


23

**Step 1:**

To a stirred solution of 5-{3-[(Z)-({[(tert-butoxy)carbonyl]amino}(4,4-dimethylpiperidin-1-yl)methylidene)amino]-5-fluorophenyl}-2-methylfuran-3-carboxylic acid **Ie** (40.0 mg, 0.084 mmol,

1.0 eq.) and tert-butyl 4-[(piperidin-3-yl)methyl]piperazine-1-carboxylate (racemic mixture, 28.7 mg, 0.10 mmol, 1.2 eq.) in anhydrous DMF (1.6 mL) under an argon atmosphere was added HATU (48.2 mg, 0.13 mmol, 1.5 eq.). After stirring for 15 min, DIPEA (0.074 mL, 0.42 mmol, 5.0 eq.) was added and the reaction mixture was stirred at room temperature for an additional 15 min. The solution was concentrated under reduced pressure and purified by prepHPLC (ACN/water + 0.1% FA) to afford tert-butyl 4-{[1-(5-{3-[(Z)-({[(tert-butoxy)carbonyl]amino}(4,4-dimethylpiperidin-1-yl)methylidene)amino]-5-fluorophenyl}-2-methylfuran-3-carbonyl)piperidin-3-yl]methyl}piperazine-1-carboxylate (40.0 mg, 64.1% yield) as a beige solid.

LCMS (ESI) $[M+H]^+$= 739.4

**Step 2:**

Tert-butyl 4-{[1-(5-{3-[(Z)-({[(tert-butoxy)carbonyl]amino}(4,4-dimethylpiperidin-1-yl)methylidene)amino]-5-fluorophenyl}-2-methylfuran-3-carbonyl)piperidin-3-yl]methyl}piperazine-1-carboxylate (40.0 mg, 0.05 mmol, 1.0 eq.) was dissolved in 4M HCl in dioxane (1.5 mL, 0.043 mol, 798.0 eq.). The reaction mixture was stirred at room temperature for 1 h. Upon completion, the solvent was removed under reduced pressure and the residue was freeze-dried to afford (Z)-N'-[3-fluoro-5-(5-methyl-4-{3-[(piperazin-1-yl)methyl]piperidine-1-carbonyl}furan-2-yl)phenyl]-4,4-dimethylpiperidine-1-carboximidamide dihydrochloride (33.0 mg, 99.7% yield),which was used in the subsequent step without further purification.

LCMS (ESI) $[M+H]^+$= 539.5

**Step 3:**

To a stirred solution of (Z)-N'-[3-fluoro-5-(5-methyl-4-{3-[(piperazin-1-yl)methyl]piperidine-1-carbonyl}furan-2-yl)phenyl]-4,4-dimethylpiperidine-1-carboximidamide dihydrochloride (15.0 mg, 0.025 mmol, 1.0 eq.) in anhydrous DMF (1.0 mL), triethylamine (0.005 mL, 0.037 mmol, 1.5 eq.) was added and the reaction mixture cooled to 0 °C. After 5 min, acetyl chloride (0.003 mL, 0.037 mmol, 1.5 eq.) was added and the reaction mixture was allowed to stir at room temperature for 2 h under argon atmosphere. The resulting solution was concentrated and purified by prepHPLC (ACN/water + 0.1% FA) resulting in (Z)-N'-[3-(4-{3-[(4-acetylpiperazin-1-yl)methyl]piperidine-1-carbonyl}-5-methylfuran-2-yl)-5-fluorophenyl]-4,4-dimethylpiperidine-1-carboximidamide formate (10.0 mg, 65.1% yield) as a white solid.

$^1$H NMR (500 MHz, DMSO-$d_6$, 353K) δ 8.22 (s, 2H), 6.92 – 6.89 (m, 1H), 6.89 – 6.86 (m, 2H), 6.43 (dt, *J* = 11.1, 2.2 Hz, 1H), 4.11 – 4.04 (m, 1H), 4.02 – 3.93 (m, 1H), 3.40 – 3.37 (m, 4H), 3.33 – 3.31

(m, 4H), 2.99 – 2.98– 3.00 (m, 1H), 2.75 (dd, *J* = 13.2, 9.6 Hz, 1H), 2.38 – 2.33 (m, 5H), 2.28 – 2.22 (m, 2H), 2.20 – 2.12 (m, 2H), 1.93 (s, 3H), 1.83 – 1.74 (m, 2H), 1.71 – 1.65 (m, 1H), 1.49 – 1.42 (m, 1H), 1.37 – 1.34 (m, 4H), 1.27 – 1.21 (m, 1H), 1.00 (s, 6H).
LCMS (ESI) $[M+H]^+$= 581.4

**Synthesis of Compound 24**

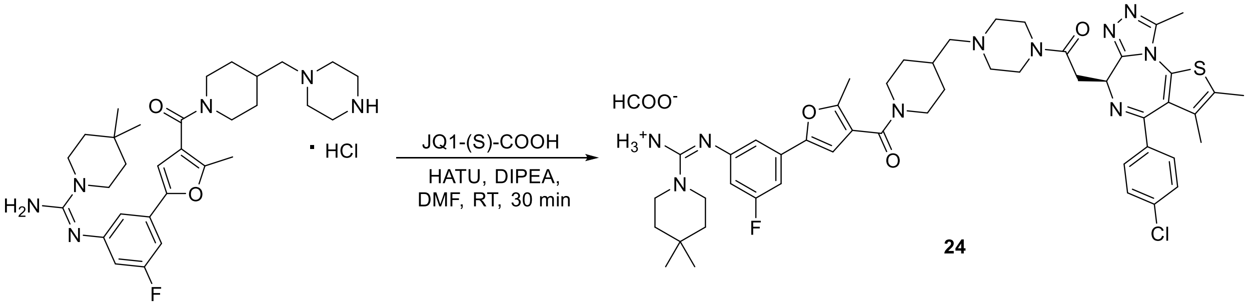


To a stirred solution of (Z)-N'-[3-fluoro-5-(5-methyl-4-{4-[(piperazin-1-yl)methyl]piperidine-1-carbonyl}furan-2-yl)phenyl]-4,4-dimethylpiperidine-1-carboximidamide hydrochloride (17.0 mg, 0.030 mmol, 1.0 eq.) and 2-[(9S)-7-(4-chlorophenyl)-4,5,13-trimethyl-3-thia-1,8,11,12-tetraazatricyclo[8.3.0.0$^{2}$,$^{6}$]trideca-2(6),4,7,10,12-pentaen-9-yl]acetic acid (13.0 mg, 0.032 mmol, 1.1 eq.) in anhydrous DMF (0.850 mL) was added HATU (16.9 mg, 0.044 mmol, 1.5 eq.) under an argon atmosphere. After stirring for 15 min, DIPEA (0.026 mL, 0.15 mmol, 5.0 eq.) was added and the mixture was stirred at room temperature for an additional 30 min. The resulting solution was concentrated under reduced pressure and purified by prepHPLC (ACN/water + 0.1% FA) resulting in (Z)-N'-[3-(4-{4-[(4-{2-[(9R)-7-(4-chlorophenyl)-4,5,13-trimethyl-3-thia-1,8,11,12-tetraazatricyclo[8.3.0.0$^{2}$,$^{6}$]trideca-2(6),4,7,10,12-pentaen-9-yl]acetyl}piperazin-1-yl)methyl]piperidine-1-carbonyl}-5-methylfuran-2-yl)-5-fluorophenyl]-4,4-dimethylpiperidine-1-carboximidamide formate (9.7 mg, 35.5% yield) as a white solid.
LCMS (ESI) $[M+H]^+$= 921.4
$^{1}$H NMR (500 MHz, DMSO-$d_6$, 353K) δ 8.14 (s, 1H), 7.47 – 7.44 (m, 4H), 7.01 (dt, *J* = 9.7, 2.0 Hz, 1H), 6.99 – 6.97 (m, 1H), 6.90 (s, 1H), 6.54 (dt, *J* = 10.8, 2.2 Hz, 1H), 4.60 (t, *J* = 6.7 Hz, 1H), 4.19 - 4.05 (m, 2H), 3.65 – 3.58 (m, 3H), 3.44 – 3.39 (m, 5H), 2.95 – 2.88 (m, 4H), 2.60 (s, 3H), 2.44 – 2.42 (m, 5H), 2.39 – 2.36 (m, 5H), 2.27 (d, *J* = 6.9 Hz, 2H), 1.86-1.78 (m, 3H), 1.66 (s, 3H), 1.40 – 1.36 (m, 4H), 1.16 – 1.09 (m, 2H), 0.99 (s, 6H).

**Synthesis of Compound 25**

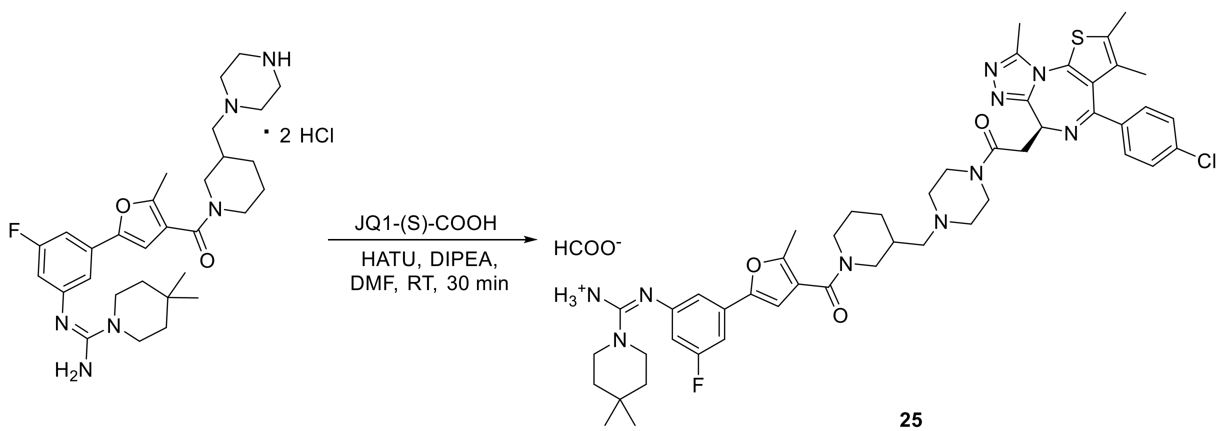


To a stirred solution of (Z)-N'-[3-fluoro-5-(5-methyl-4-{3-[(piperazin-1-yl)methyl]piperidine-1-carbonyl}furan-2-yl)phenyl]-4,4-dimethylpiperidine-1-carboximidamide dihydrochloride (18.0 mg, 0.029 mmol, 1.0 eq.) and _2-[(9S)-7-(4-chlorophenyl)-4,5,13-trimethyl-3-thia-1,8,11,12-tetraazatricyclo[8.3.0.0$^{2}$,$^{6}$]trideca-2(6),4,7,10,12-pentaen-9-yl]acetic acid (13.0 mg, 0.032 mmol, 1.1 eq.) in anhydrous DMF (1.0 mL) was added HATU (16.8 mg, 0.044 mmol, 1.5 eq.) under an argon atmosphere. After stirring for 15 min, DIPEA (0.026 mL, 0.147 mmol, 5.0 eq.) was added and the mixture was stirred at room temperature for an additional 30 min. The resulting solution was concentrated under reduced pressure and purified by prepHPLC (ACN/water + 0.1% FA) to afford (Z)-N'-[3-(4-{3-[(4-{2-[(9S)-7-(4-chlorophenyl)-4,5,13-trimethyl-3-thia-1,8,11,12-tetraazatricyclo[8.3.0.0$^{2}$,$^{6}$]trideca-2(6),4,7,10,12-pentaen-9-yl]acetyl}piperazin-1-yl)methyl]piperidine-1-carbonyl}-5-methylfuran-2-yl)-5-fluorophenyl]-4,4-dimethylpiperidine-1-carboximidamide formate (16.0 mg, 59.0% yield) as a white solid.

LCMS (ESI) $[M+H]^+$= 921.4

$^1$H NMR (500 MHz, DMSO-$d_6$, 353K) δ 8.16 (s, 1H), 7.47 – 7.42 (m, 4H), 7.01 – 6.92 (m, 2H), 6.91 (s, 1H), 6.51 – 6.44 (m, 1H), 4.58 (t, *J* = 6.7 Hz, 1H), 4.16 – 4.05 (m, 1H), 4.04 – 3.92 (m, 1H), 3.59 – 3.44 (m, 5H), 3.40 – 3.36 (m, 5H), 2.81 – 2.75 (m, 2H), 2.60 (s, 3H), 2.44 – 2.40 (m, 4H), 2.38 (s, 3H), 2.37 – 2.35 (m, 1H), 2.28 – 2.14 (m, 3H), 1.86 – 1.77 (m, 2H), 1.73 – 1.67 (m, 1H), 1.66 (s, 3H), 1.50 – 1.43 (m, 1H), 1.36 – 1.32 (m, 4H), 1.29 – 1.21 (m, 2H), 0.96 (d, *J* = 1.4 Hz, 6H).

**Synthesis of Compound 26**

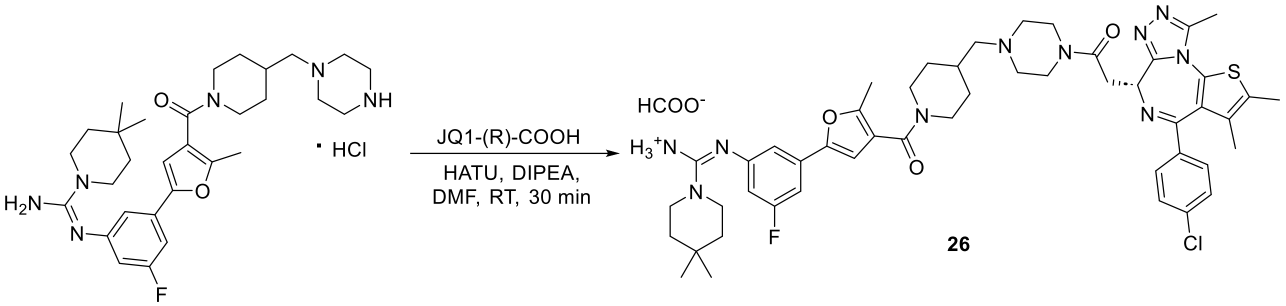


26

To a stirred solution of (Z)-N'-[3-fluoro-5-(5-methyl-4-{4-[(piperazin-1-yl)methyl]piperidine-1-carbonyl}furan-2-yl)phenyl]-4,4-dimethylpiperidine-1-carboximidamide hydrochloride (20.0 mg, 0.035 mmol, 1.0 eq.) and 2-[(9R)-7-(4-chlorophenyl)-4,5,13-trimethyl-3-thia-1,8,11,12-tetraazatricyclo[8.3.0.0$^{2}$,$^{6}$]trideca-2(6),4,7,10,12-pentaen-9-yl]acetic acid (13.9 mg, 0.035 mmol, 1.0 eq.) in anhydrous DMF (1.0 mL) was added HATU (19.8 mg, 0.052 mmol, 1.5 eq.) under an argon atmosphere. After stirring for 15 min, DIPEA (0.030 mL, 0.174 mmol, 5.0 eq.) was added and the mixture was stirred at room temperature for an additional 30 min. The resulting solution was concentrated under reduced pressure and purified by prepHPLC (ACN/water + 0.1% FA) to afford (Z)-N'-[3-(4-{4-[(4-{2-[(9R)-7-(4-chlorophenyl)-4,5,13-trimethyl-3-thia-1,8,11,12-tetraazatricyclo[8.3.0.0$^{2}$,$^{6}$]trideca-2(6),4,7,10,12-pentaen-9-yl]acetyl}piperazin-1-yl)methyl]piperidine-1-carbonyl}-5-methylfuran-2-yl)-5-fluorophenyl]-4,4-dimethylpiperidine-1-carboximidamide formate (9.7 mg, 30.3%) as a white solid.

LCMS (ESI) $[M+H]^+$= 921.5

$^1$H NMR (500 MHz, DMSO-$d_6$, 353K) δ 8.21 (s, 1H), 7.47 – 7.42 (m, 4H), 6.93 – 6.89 (m, 2H), 6.87 (s, 1H), 6.47 (dt, *J* = 11.1, 2.2 Hz, 1H), 4.60 (t, *J* = 6.7 Hz, 1H), 4.25 – 4.19 (m, 2H), 3.62 – 3.56 (m, 3H), 3.43 (d, *J* = 6.6 Hz, 1H), 3.41 – 3.37 (m, 4H), 3.01 – 2.99 (m, 4H), 2.60 (s, 3H), 2.48 – 2.44 (m, 2H), 2.44 – 2.36 (m, 5H), 2.27 (d, *J* = 6.9 Hz, 2H), 2.18 (s, 3H), 1.95 – 1.83 (m, 3H), 1.66 (d, *J*= 0.8 Hz 3H), 1.37 – 1.33 (m, 4H), 1.24 – 1.13 (m, 2H), 0.99 (s, 6H)

### Synthesis of Compound 27

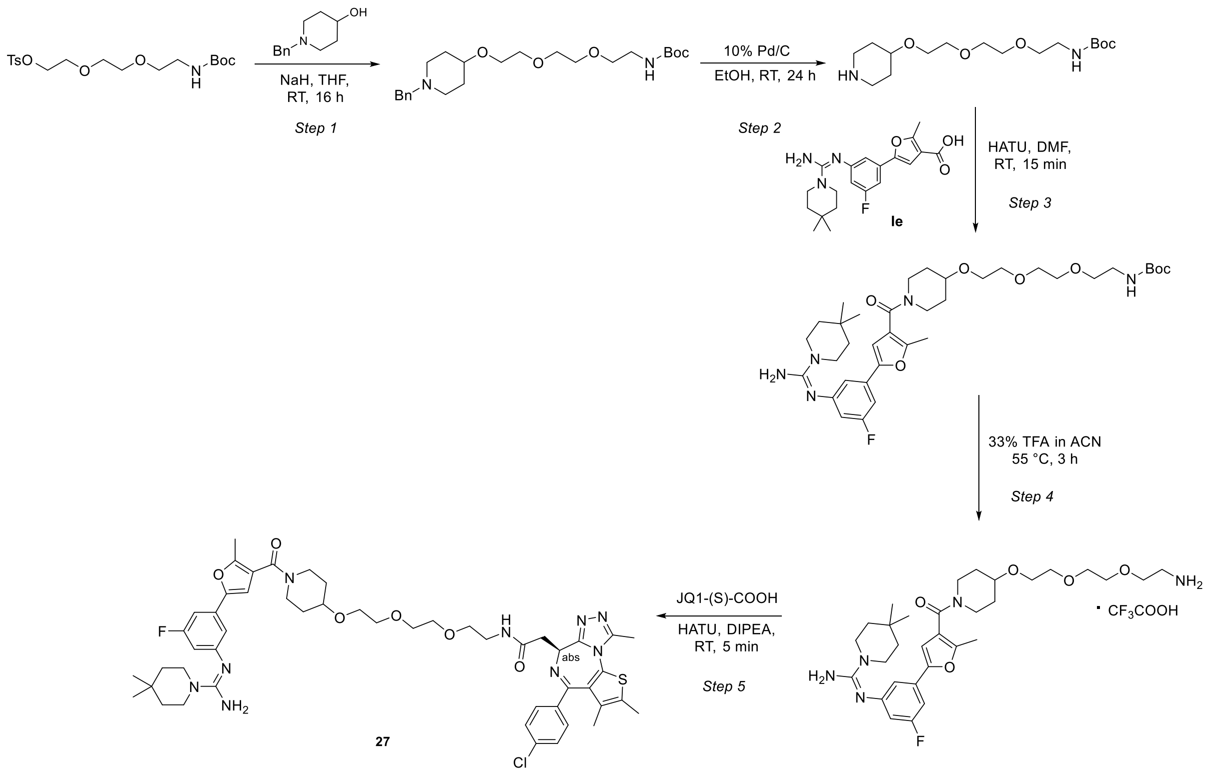

**Step 1:**

To a stirred solution of 1-benzylpiperidin-4-ol (61.6 mg, 0.32 mmol, 1.3 eq.) in anhydrous THF (2.0 mL) under an argon atmosphere at 0°C was added NaH (60% dispersion in mineral oil, 49.6 mg, 1.24 mmol, 5.0 eq.). The mixture was stirred for 15 min, followed by the dropwise addition of 2,2-dimethyl-4-oxo-3,8,11-trioxa-5-azatridecan-13-yl 4-methylbenzenesulfonate (100.0 mg, 0.25 mmol, 1.0 eq.) in anhydrous THF (1.0 mL). The reaction mixture was allowed to warm to room temperature and stirred overnight. The resulting mixture was quenched with ice-cold water and extracted with EtOAc. The combined organic layers were concentrated under reduced pressure and the crude residue was purified by reversed-phase silica gel column chromatography (ACN/water + 0.1% FA) to afford tert-butyl N-[2-(2-{2-[(1-benzylpiperidin-4-yl)oxy]ethoxy}ethoxy)ethyl]carbamate (42.0 mg, 40.1% yield) as a colorless oil.

LCMS (ESI) $[M+H]^+$= 423.2

**Step 2:**

To a stirred solution of tert-butyl N-[2-(2-{2-[(1-benzylpiperidin-4-yl)oxy]ethoxy}ethoxy)ethyl]carbamate (15.0 mg, 0.028 mmol, 1.0 eq.) in EtOH (1.0 mL) was added

10% Pd/C (6.8 mg, 0.006 mmol, 0.226 eq.). The flask was purged with hydrogen and the reaction was stirred under hydrogen atmosphere (balloon) at room temperature for 24 h. Upon completion, the mixture was filtered through a syringe filter and filter was washed with EtOH. The combined filtrate was concentrated under reduced pressure and dried *in vacuo* to afford tert-butyl N-(2-{2-[2-(piperidin-4-yloxy)ethoxy]ethoxy}ethyl)carbamate (10.1 mg, 85.6% yield) as a colorless oil.

LCMS (ESI) $[M+H]^+$= 333.2

**Step 3:**

To a stirred solution of 5-{3-[(Z)-[amino(4,4-dimethylpiperidin-1-yl)methylidene]amino]-5-fluorophenyl}-2-methylfuran-3-carboxylic acid **Ie** (17.0 mg, 0.046 mmol, 1.0 eq.) and tert-butyl N-(2-{2-[2-(piperidin-4-yloxy)ethoxy]ethoxy}ethyl)carbamate (18.2 mg, 0.055 mmol, 1.2 eq.) in anhydrous DMF (1.0 mL) under an argon atmosphere was added HATU (26.0 mg, 0.068 mmol, 1.5 eq.). After stirring for 15 min, DIPEA (0.040 mL, 0.228 mmol, 5.0 eq.) was added and the reaction mxiture was stirred at room temperature for an additional 15 min. Upon completion, the mixture was concentrated under reduced pressure and the crude residue was purified by prepHPLC (ACN/water + 0.1% FA) to afford tert-butyl (Z)-(2-(2-(2-((1-(5-(3-((amino(4,4-dimethylpiperidin-1-yl)methylene)amino)-5-fluorophenyl)-2-methylfuran-3-carbonyl)piperidin-4-yl)oxy)ethoxy)ethoxy)ethyl)carbamate (18.9 mg, 60.4% yield) as a beige solid.

LCMS (ESI) $[M+H]^+$=688.8

**Step 4:**

Tert-butyl (Z)-(2-(2-(2-((1-(5-(3-((amino(4,4-dimethylpiperidin-1-yl)methylene)amino)-5-fluorophenyl)-2-methylfuran-3-carbonyl)piperidin-4-yl)oxy)ethoxy)ethoxy)ethyl)carbamate (12.1 mg, 0.018 mmol, 1.0 eq.) was dissolved in 33% TFA in ACN (1.0 mL, 138.5 eq.) and the reaction mixture was stirred at 55 °C for 3 h. Upon completion, the mixture was concentrated under reduced prsssure and dried *in vacuo* to afford (Z)-N'-{3-[4-(4-{2-[2-(2-aminoethoxy)ethoxy]ethoxy}piperidine-1-carbonyl)-5-methylfuran-2-yl]-5-fluorophenyl}-4,4-dimethylpiperidine-1-carboximidamide; trifluoracetic acid (10.0 mg, quantitative yield) as a yellow oil.

LCMS (ESI) $[M+H]^+$= 588.4

**Step 5:**

To a stirred solution of (Z)-N'-{3-[4-(4-{2-[2-(2-aminoethoxy)ethoxy]ethoxy}piperidine-1-carbonyl)-5-methylfuran-2-yl]-5-fluorophenyl}-4,4-dimethylpiperidine-1-carboximidamide (20.0 mg, 0.034 mmol, 1.0 eq.) and 2-[(9S)-7-(4-chlorophenyl)-4,5,13-trimethyl-3-thia-1,8,11,12-tetraazatricyclo[8.3.0.0$^{2,6}$]trideca-2(6),4,7,10,12-pentaen-9-yl]acetic acid (16.4 mg, 0.041 mmol, 1.2 eq.) in anhydrous DMF (1.0 mL) under an argon atmosphere was added HATU (19.4 mg, 0.051 mmol, 1.5 eq.). After stirring for 15 min, DIPEA (0.030 mL, 0.170 mmol, 5.0 eq.) was added and the reaction was stirred at room temperature for 5 min. Upon completion, the mixture was concentrated under reduced pressure and the crude residue was purified by prepHPLC (ACN/water + 0.1% FA) to afford N-{2-[2-(2-{[1-(5-{3-[(Z)-[amino(4,4-dimethylpiperidin-1-yl)methylidene]amino]-5-fluorophenyl}-2-methylfuran-3-carbonyl)piperidin-4-yl]oxy}ethoxy)ethoxy]ethyl}-2-[(9S)-7-(4-chlorophenyl)-4,5,13-trimethyl-3-thia-1,8,11,12-tetraazatricyclo[8.3.0.0$^{2,6}$]trideca-2(6),4,7,10,12-pentaen-9-yl]acetamide (8.0 mg, 24.2% yield) as a pale pink solid.

LCMS (ESI) $[M+H]^+$= 970.4

$^1$H NMR (500 MHz, DMSO-$d_6$, 353K) δ 7.92 – 7.87 (m, 1H), 7.45 – 7.43 (m, 4H), 7.00 – 6.97 (m, 1H), 6.97 – 6.95 (m, 1H), 6.90 (s, 1H), 6.52 (d, *J* = 10.8 Hz, 1H), 4.53 (t, *J* = 6.9 Hz, 1H), 3.80 – 3.74 (m, 2H), 3.59 – 3.55 (m, 10H), 3.50 (t, *J* = 6.0 Hz, 2H), 3.42 – 3.39 (m, 4H), 3.29 – 3.24 (m, 5H), 2.60 (s, 3H), 2.41 (s, 3H), 2.36 (s, 3H), 1.88 – 1.81 (m, 2H), 1.66 – 1.64 (m, 3H), 1.48 (dtd, *J* = 12.6, 8.4, 3.9 Hz, 2H), 1.39 – 1.35 (m, 4H), 0.98 (s, 6H).

### Synthesis of Compound 28

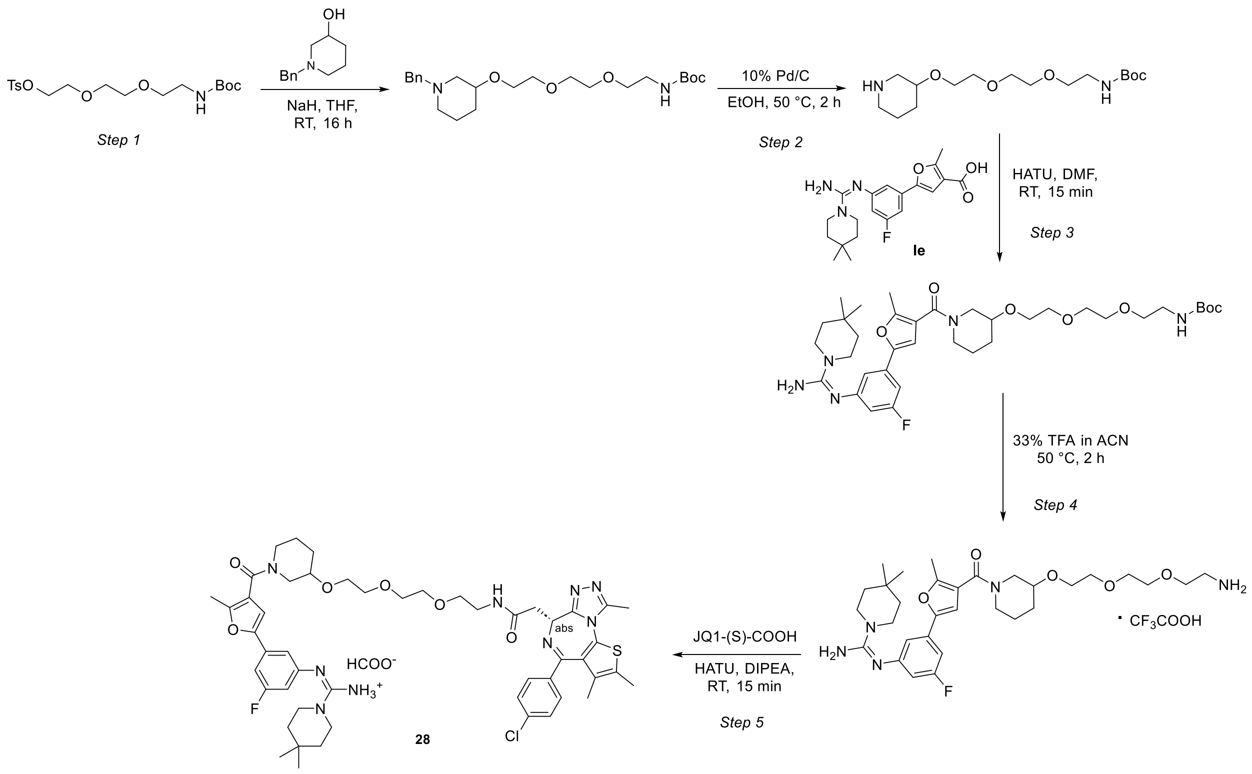


### Step 1:

To a stirred solution of 1-benzylpiperidin-3-ol (92.4 mg, 0.48 mmol, 1.3 eq.) in anhydrous THF (4.0 mL) under an argon atmosphere at 0°C was added NaH (60% dispersion in mineral oil, 74.3 mg, 1.86 mmol, 5.0 eq.). The reaction mixture was stirred for 15 min, followed by the dropwise addition of 2,2-dimethyl-4-oxo-3,8,11-trioxa-5-azatridecan-13-yl 4-methylbenzenesulfonate (150.0 mg, 0.37 mmol, 1.0 eq.) in anhydrous THF (1.0 mL). The reaction mixture was allowed to warm to room temperature and stirred overnight. The mixture was quenched with ice-cold water and extracted with EtOAc. The combined organic layers were concentrated under reduced pressure and the crude residue was purified by reversed-phase silica gel column chromatography (ACN/water + 0.1% FA) to afford tert-butyl N-[2-(2-{2-[(1-benzylpiperidin-3-yl)oxy]ethoxy}ethoxy)ethyl]carbamate (76.0 mg, 48.4% yield) as a colorless oil.

LCMS (ESI) $[M+H]^+$= 423.2

**Step 2:**

To a stirred solution of tert-butyl N-[2-(2-{2-[(1-benzylpiperidin-3-yl)oxy]ethoxy}ethoxy)ethyl]carbamate (22.0 mg, 0.052 mmol, 1.0 eq.) in EtOH (2.0 mL) was added 10% Pd/C (16.6 mg, 0.016 mmol, 0.3 eq.). The flask was purged with hydrogen and the reaction mixture was stirred under hydrogen atmosphere (balloon) at 50 °C for 2 h. Upon completion, the mixture was then filtered through syringe filter and the filter was washed with EtOH. The combined filtrate was concentrated under reduced pressure and dried *in vacuo* to afford tert-butyl N-(2-{2-[2-(piperidin-3-yloxy)ethoxy]ethoxy}ethyl)carbamate (15.0 mg, 86.7% yield) as a colorless oil.

LCMS (ESI) $[M+H]^+$= 333.2

**Step 3:**

To a stirred solution of 5-{3-[(Z)-[amino(4,4-dimethylpiperidin-1-yl)methylidene]amino]-5-fluorophenyl}-2-methylfuran-3-carboxylic acid **Ie** (12.0 mg, 0.032 mmol, 1.0 eq.) and tert-butyl N-(2-{2-[2-(piperidin-3-yloxy)ethoxy]ethoxy}ethyl)carbamate (11.2 mg, 0.034 mmol, 1.05 eq.) in anhydrous DMF (1.0 mL) under an argon atmosphere was added HATU (18.3 mg, 0.048 mmol, 1.5 eq.). After stirring for 15 min, DIPEA (0.028 mL, 0.161 mmol, 5.0 eq.) was added and the reaction was stirred at room temperature for an additional 15 min. Upon completion, the mixture was concentrated under reduced pressure and the crude residue was purified by prepHPLC (ACN/water + 0.1% FA) to afford tert-butyl (Z)-(2-(2-(2-((1-(5-(3-((amino(4,4-dimethylpiperidin-1-yl)methylene)amino)-5-fluorophenyl)-2-methylfuran-3-carbonyl)piperidin-3-yl)oxy)ethoxy)ethoxy)ethyl)carbamate (11 mg, 50.0% yield) as a white solid.

LCMS (ESI) $[M+H]^+$=688.4

**Step 4:**

Tert-butyl (Z)-(2-(2-(2-((1-(5-(3-((amino(4,4-dimethylpiperidin-1-yl)methylene)amino)-5-fluorophenyl)-2-methylfuran-3-carbonyl)piperidin-3-yl)oxy)ethoxy)ethoxy)ethyl)carbamate (11.0 mg, 0.016 mmol, 1.0 eq.) was dissolved in 33% TFA in ACN (1.0 mL, 133.07 eq.) and the reaction was stirred at 50 °C for 2 h. Upon completion, the mixture was concentrated under reduced pressure and dried *in vacuo* to afford (Z)-N'-(3-(4-(3-(2-(2-(2-aminoethoxy)ethoxy)ethoxy)piperidine-1-carbonyl)-5-methylfuran-2-yl)-5-fluorophenyl)-4,4-dimethylpiperidine-1-carboximidamide; trifluoroacetic acid (10.0 mg, quantitative yield) as a yellow oil.

LCMS (ESI) $[M+H]^+$= 588.3

**Step 5:**

To a stirred solution of (Z)-N'-(3-(4-(3-(2-(2-(2-aminoethoxy)ethoxy)ethoxy)piperidine-1-carbonyl)-5-methylfuran-2-yl)-5-fluorophenyl)-4,4-dimethylpiperidine-1-carboximidamide (10.0 mg, 0.017 mmol, 1.0 eq.) and (S)-2-(4-(4-chlorophenyl)-2,3,9-trimethyl-6H-thieno[3,2-f][1,2,4]triazolo[4,3-a][1,4]diazepin-6-yl)acetic acid (8.2 mg, 0.020 mmol, 1.2 eq.) in anhydrous DMF (1.0 mL) under an argon atmosphere was added HATU (9.7 mg, 0.026 mmol, 1.5 eq.). After stirring for 15 min, DIPEA (0.015 mL, 0.085 mmol, 5.0 eq.) was added and the reaction was stirred at room temperature for an additional 15 min. The resulting mixture was concentrated under reduced pressure and crude residue was purified by prepHPLC (ACN/water + 0.1% FA) to afford N-(2-(2-(2-((1-(5-(3-(((Z)-amino(4,4-dimethylpiperidin-1-yl)methylene)amino)-5-fluorophenyl)-2-methylfuran-3-carbonyl)piperidin-3-yl)oxy)ethoxy)ethoxy)ethyl)-2-((S)-4-(4-chlorophenyl)-2,3,9-trimethyl-6H-thieno[3,2-f][1,2,4]triazolo[4,3-a][1,4]diazepin-6-yl)acetamide formate (2.8 mg, 17.0% yield) as a white solid.

LCMS (ESI) $[M+H]^+$= 970.3

$^1$H NMR (500 MHz, DMSO-$d_6$, 353K) δ 8.39 (s, 1H), 7.91 – 7.84 (m, 1H), 7.45 – 7.29 (m, 4H), 6.89 – 6.85 (m, 3H), 6.40 (dt, *J* = 11.1, 2.1 Hz, 1H), 4.52 (t, *J* = 6.9 Hz, 1H), 3.79 – 3.70 (m, 1H), 3.56 – 3.52 (m, 8H), 3.50 – 3.46 (m, 3H), 3.45 – 3.40 (m, 2H), 3.39 – 3.36 (m, 4H), 3.33 – 3.25 (m, 5H), 2.60 (s, 3H), 2.42 (s, 3H), 2.35 (s, 3H), 1.92 – 1.84 (m, 1H), 1.76 – 1.67 (m, 1H), 1.65 (s, 3H), 1.63 – 1.55 (m, 1H), 1.46 – 1.38 (m, 1H), 1.37 – 1.30 (m, 4H), 0.97 (s, 6H).

**Synthesis of Compound 29**

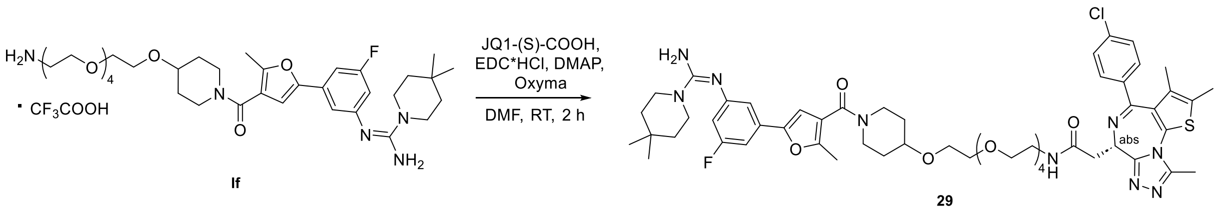


If

29

Compound (Z)-N'-(3-(4-(4-((14-amino-3,6,9,12-tetraoxatetradecyl)oxy)piperidine-1-carbonyl)-5-methylfuran-2-yl)-5-fluorophenyl)-4,4-dimethylpiperidine-1-carboximidamide; trifluoroacetic acid **If** was synthesized following the procedure for compound **27** starting from 2,2-dimethyl-4-oxo-3,8,11,14,17-pentaoxa-5-azanonadecan-19-yl 4-methylbenzenesulfonate (4 steps, overall yield: 9%).

**Final step:**

To a stirred solution of (S)-2-(4-(4-chlorophenyl)-2,3,9-trimethyl-6H-thieno[3,2-f][1,2,4]triazolo[4,3-a][1,4]diazepin-6-yl)acetic acid (10.2 mg, 0.025 mmol, 1.0 eq.) in anhydrous DMF (1.5 mL) were added EDC · HCl (14.6 mg, 0.076 mmol, 3.0 eq.), and Oxyma (10.8 mg, 0.076 mmol, 3.0 eq.). The mixture was stirred at room temperature for 15 min, followed by the addition of DMAP (9.3 mg, 0.076 mmol, 3.0 eq.) and (Z)-N'-(3-(4-(4-((14-amino-3,6,9,12-tetraoxatetradecyl)oxy)piperidine-1-carbonyl)-5-methylfuran-2-yl)-5-fluorophenyl)piperidine-1-carboximidamide; trifluoroacetic acid **If** (20.0 mg, 0.025 mmol, 1.0 eq.). The reaction mixture was stirred at room temperature for 2 h. Upon completion, the mixture was concentrated under reduced pressure and the crude residue was purified by prepHPLC (ACN/water + 0.1% FA) to afford N-(14-{[1-(5-{3-[(Z)-[amino(4,4-dimethylpiperidin-1-yl)methylidene]amino]-5-fluorophenyl}-2-methylfuran-3-carbonyl)piperidin-4-yl]oxy}-3,6,9,12-tetraoxatetradecan-1-yl)-2-[(9S)-7-(4-chlorophenyl)-4,5,13-trimethyl-3-thia-1,8,11,12-tetraazatricyclo[8.3.0.0$^{2}$,$^{6}$]trideca-2(6),4,7,10,12-pentaen-9-yl]acetamide (7.9 mg, 29.5% yield) as a white solid.

LCMS (ESI) $[M+H]^+$= 1058.3

$^1$H NMR (500 MHz, DMSO-$d_6$) δ 7.92 – 7.86 (m, 1H), 7.46 – 7.43 (m, 4H), , 6.95 – 6.91 (m, 1H), 6.91 – 6.89 (m, 1H), 6.88 (s, 1H), 6.44 (dt, *J* = 11.1, 2.3 Hz, 1H), 4.53 (t, *J* = 7.0 Hz, 1H), 3.81 – 3.73 (m, 2H), 3.60 – 3.53 (m, 17H), 3.49 (t, *J* = 5.9 Hz, 2H), 3.41 – 3.38 (m, 4H), 3.33– 3.25 (m, 6H), 2.60 (s, 3H), 2.42 (m, 3H), 2.36 (s, 3H), 1.87 – 1.81 (m, 2H), 1.67 – 1.64 (m, 3H), 1.52 – 1.44 (m, 2H), 1.37 – 1.34 (m, 4H), 0.98 (s, 6H).

**Synthesis of Compound 30**

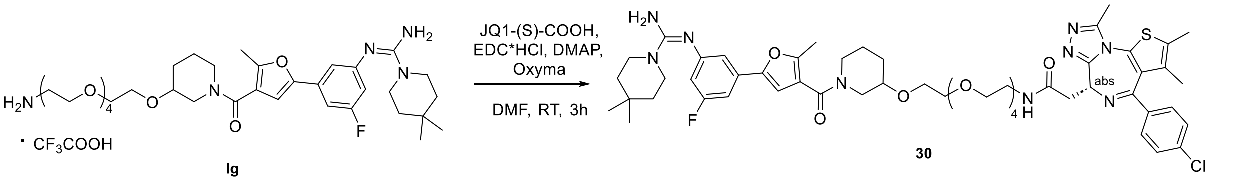


Compound (Z)-N'-(3-(4-(3-((14-amino-3,6,9,12-tetraoxatetradecyl)oxy)piperidine-1-carbonyl)-5-methylfuran-2-yl)-5-fluorophenyl)piperidine-1-carboximidamide; trifluoroacetic acid **Ig** was synthesized following the procedure for compound **29** starting from 2,2-dimethyl-4-oxo-3,8,11,14,17-pentaoxa-5-azanonadecan-19-yl 4-methylbenzenesulfonate (4 steps, overall yield: 18%).

**Final step:**

To a stirred solution of 2-[(9S)-7-(4-chlorophenyl)-4,5,13-trimethyl-3-thia-1,8,11,12-tetraazatricyclo[8.3.0.0$^{2}$,$^{6}$]trideca-2(6),4,7,10,12-pentaen-9-yl]acetic acid (12.0 mg, 0.030 mmol, 1.0 eq.) in anhydrous DMF (1.5 mL) were added EDC · HCl (17.2 mg, 0.090 mmol, 3.0 eq.) and Oxyma (12.8 mg, 0.090 mmol, 3.0 eq.). The mixture was stirred at room temperature for 15 min, followed by the addition of DMAP (11.0 mg, 0.090 mmol, 3.0 eq.) and (Z)-N'-[3-(4-{3-[(14-amino-3,6,9,12-tetraoxatetradecan-1-yl)oxy]piperidine-1-carbonyl}-5-methylfuran-2-yl)-5-fluorophenyl]-4,4-dimethylpiperidine-1-carboximidamide; trifluoroacetic acid **Ig** (28.4 mg, 0.036 mmol, 1.2 eq.). The reaction mixture was stirred at room temperature for 3 h. Upon completion, the solvent was removed under reduced pressure and the residue was purified by prepHPLC (ACN/water + 0.1% FA) to afford N-(14-{[1-(5-{3-[(Z)-[amino(4,4-dimethylpiperidin-1-yl)methylidene]amino]-5-fluorophenyl}-2-methylfuran-3-carbonyl)piperidin-3-yl]oxy}-3,6,9,12-tetraoxatetradecan-1-yl)-2-[(9S)-7-(4-chlorophenyl)-4,5,13-trimethyl-3-thia-1,8,11,12-tetraazatricyclo[8.3.0.0$^{2}$,$^{6}$]trideca-2(6),4,7,10,12-pentaen-9-yl]acetamide (7.5 mg, 23.6% yield) as a white solid.

LCMS (ESI) $[M+H]^{+}$= 1058.3

$^{1}$H NMR (500 MHz, DMSO-$d_6$, 353K) δ 7.93 – 7.85 (m, 1H), 7.46 – 7.43 (m, 4H), 6.93 – 6.88 (m, 2H), 6.87 (s, 1H), 6.44 (dt, *J* = 11.1, 2.1 Hz, 1H), 4.53 (t, *J* = 6.9 Hz, 1H), 3.78 – 3.71 (m, 1H), 3.56 – 3.51 (m, 17H), 3.49 – 3.47 (m, 2H), 3.44 – 3.40 (m, 2H), 3.40 – 3.37 (m, 4H), 3.32 – 3.25 (m, 5H), 2.60 (s, 3H), 2.43 – 2.41 (m, 3H), 2.36 (s, 3H), 1.92 – 1.84 (m, 1H), 1.76 – 1.69 (m, 1H), 1.65 (s, 3H), 1.62 – 1.56 (m, 1H), 1.45 – 1.39 (m, 1H), 1.37 – 1.34 (m, 4H), 0.97 (s, 6H).

**Synthesis of Compound 31**

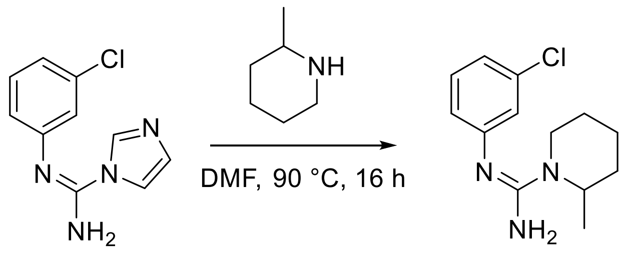


**Ih** **31**

Compound (Z)-N'-(3-chlorophenyl)-1H-imidazole-1-carboximidamide **Ih** was synthesized following the procedure for compound **3** (step 1) starting from 3-chloroaniline and di(1H-imidazol-1-yl)methanimine (yield: 65%).

**Final step:**

To a stirred solution of (Z)-N'-(3-chlorophenyl)-1H-imidazole-1-carboximidamide **(Ih)** (300 mg, 1.36 mmol) in anhydrous DMF (5.0. mL) was added 3-methylpiperidine (0.255 mL, 2.18 mmol, 1.6 eq.). The reaction mixture was heated at 90°C and stirred for 16 h. Upon completion, the reaction mixture was concentrated under reduced pressure to provide the crude residue, which was purified by prep-HPLC (aq. $NH_4HCO_3$/ACN) to afford (Z)-N'-(3-chlorophenyl)-2-methylpiperidine-1-carboximidamide (127 mg, 27%) as off white solid.

LCMS: $[M+H]^+$= 251.9

$^1$H NMR (400 MHz, DMSO-$d_6$) δ7.19 (t, *J* = 8.0, 1H), 6.89 – 6.82 (m, 1H), 6.76 – 6.71 (m, 1H), 6.70 – 6.65 (m, 1H), 4.31 – 4.25 (m, 1H), 3.73 – 3.67 (m, 1H), 2.87 – 2.76 (m, 1H), 1.63 – 1.44 (m, 5H), 1.36 – 1.29 (m, 1H), 1.08 (d, *J*= 6.8 Hz, 3H).

**Synthesis of Compound 32**

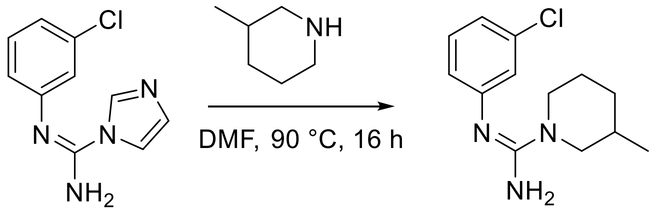


Ih    32

Compound (Z)-N'-(3-chlorophenyl)-1H-imidazole-1-carboximidamide **Ih** was synthesized following the procedure for compound **3** (step 1) starting from 3-chloroaniline and di(1H-imidazol-1-yl)methanimine (yield: 65%).

**Final step:**

To a stirred solution of (Z)-N'-(3-chlorophenyl)-1H-imidazole-1-carboximidamide **(Ih)** (400 mg, 1.81 mmol) in anhydrous DMF (5.0. mL) was added 2-methylpiperidine (0.256 mL, 2.18 mmol, 1.2 eq.). The reaction mixture was heated at 90°C and stirred for 16 h. Upon completion, the reaction mixture was diluted with water and extracted with ethyl acetate. The combined organic layers were washed with brine, dried over anhydrous sodium sulfate, filtered and concentrated under reduced pressure. The resulting crude residue was purified by prep-HPLC (aq. $NH_4HCO_3$/ACN) to afford (Z)-N'-(3-chlorophenyl)-3-methylpiperidine-1-carboximidamide (236 mg, 68%) as off white solid.

LCMS: $[M+H]^+$= 252.1

$^1H$ NMR (400 MHz, DMSO-$d_6$) δ7.17 (t, *J* = 8.0, 1H), 6.82 (d, *J* = 8.0, 1H), 6.72 – 6.62 (m, 2H), 3.90 – 3.85 (m, 1H), 2.69 – 2.57 (m, 1H), 2.37 – 2.27 (m, 1H), 1.76 – 1.69 (m, 1H), 1.61 – 1.33 (m, 4H), 1.09 – 0.99 (m, 1H), 0.83 (d, *J*= 6.6 Hz, 3H).

### Synthesis of Compound 33

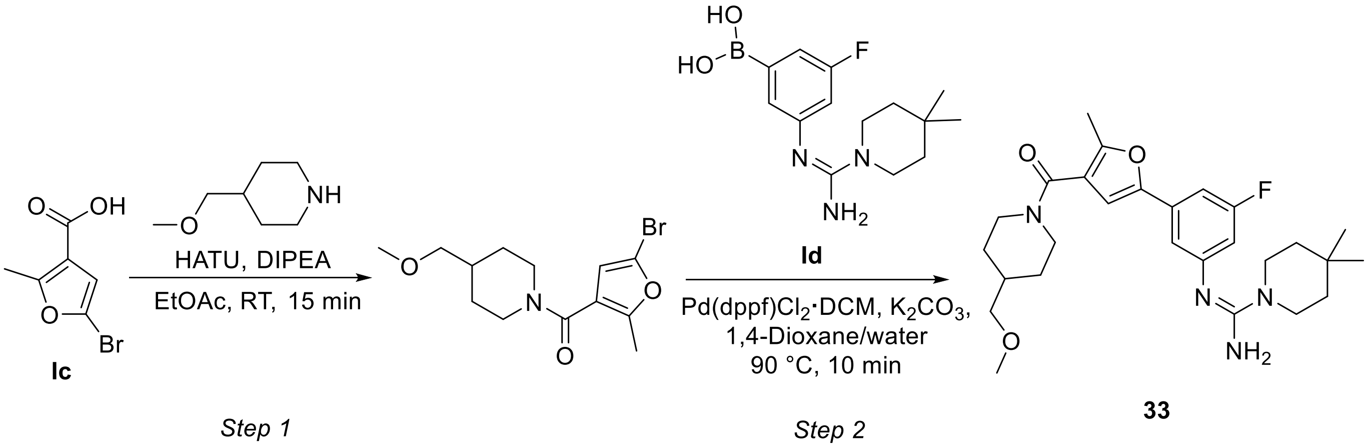


#### Step 1:

To a stirring solution of 5-bromo-2-methylfuran-3-carboxylic acid **Ic** (50.0 mg, 0.24 mmol, 1.0 eq.) and HATU (111.3 mg, 0.29 mmol, 1.2 eq.) in EtOAc (2.0 mL) were added 4-(methoxymethyl)piperidine (48.5 mg, 0.293 mmol, 1.2 eq.) and triethylamine (0.170 mL, 1.22 mmol, 5.0 eq.). The reaction mixture was stirred at room temperature for 15 min. Upon completion, the resulting solution was diluted with $NaHCO_3$ and extracted with ethyl acetate. The organic layers were dried over anhydrous sodium sulfate, filtered and concentrated under reduced pressure to afford (5-bromo-2-methylfuran-3-yl)(4-(methoxymethyl)piperidin-1-yl)methanone (70 mg, 91% yield). The crude product was utilized in the next step without further purification.

LCMS (ESI) $[M+H]^+$= 316.1

**Step 2:**

To the stirred solution of (5-bromo-2-methylfuran-3-yl)(4-(methoxymethyl)piperidin-1-yl)methanone (39.2 mg, 0.12 mmol, 1.0 eq.) in a mixture of 1,4-dioxane/$H_2O$ (2:1, 3.0 mL) were added (Z)-(3-((amino(4,4-dimethylpiperidin-1-yl)methylene)amino)-5-fluorophenyl)boronic acid **Ic** (40.0 mg, 0.14 mmol, 1.1 eq.), Pd(dppf)$Cl_2$•DCM (10.1 mg, 0.012 mmol, 0.1 eq.) and potassium carbonate (34.3 mg, 0.25 mmol, 2.0 eq.). The reaction mixture was heated at 90 °C and stirred for 10 min. The resulting mixture was filtered through Celite and purified by reversed-phase silica gel column chromatography (ACN/water + 0.1% FA) to afford (Z)-N'-(3-fluoro-5-(4-(4-(methoxymethyl)piperidine-1-carbonyl)-5-methylfuran-2-yl)phenyl)-4,4-dimethylpiperidine-1-carboximidamide formate (24.0 mg, 0.050 mmol, 39.9%) as a beige solid.

LCMS (ESI) $[M+H]^+$= 484.2

$^1$H NMR (500 MHz, methanol-d4) δ 8.40 (s, 1H), 7.38 (t, *J* = 1.7 Hz, 1H), 7.34 (ddd, *J* = 9.5, 2.3, 1.3 Hz, 1H), 6.94 (dt, *J* = 9.6, 2.2, 1H), 6.91 (s, 1H), 4.67 – 4.51 (m, 1H), 4.03 – 3.89 (m, 1H), 3.61 – 3.55 (m, 4H), 3.33 (s, 3H), 3.30 – 3.26 (m, 2H), 3.22 – 3.12 (m, 1H), 2.88 – 2.76 (m, 1H), 2.41 (s, 3H), 1.95 – 1.88 (m, 1H), 1.83 – 1.79 (m, 2H), 1.59 – 1.53 (m, 4H), 1.31 – 1.22 (m, 2H), 1.08 (s, 6H).

### 2. NMR spectra of synthesized compounds

$^{1}H$-NMR (400 MHz) of methyl 5-bromo-2-methylfuran-3-carboxylate (**Ib**) in $CDCl_3$

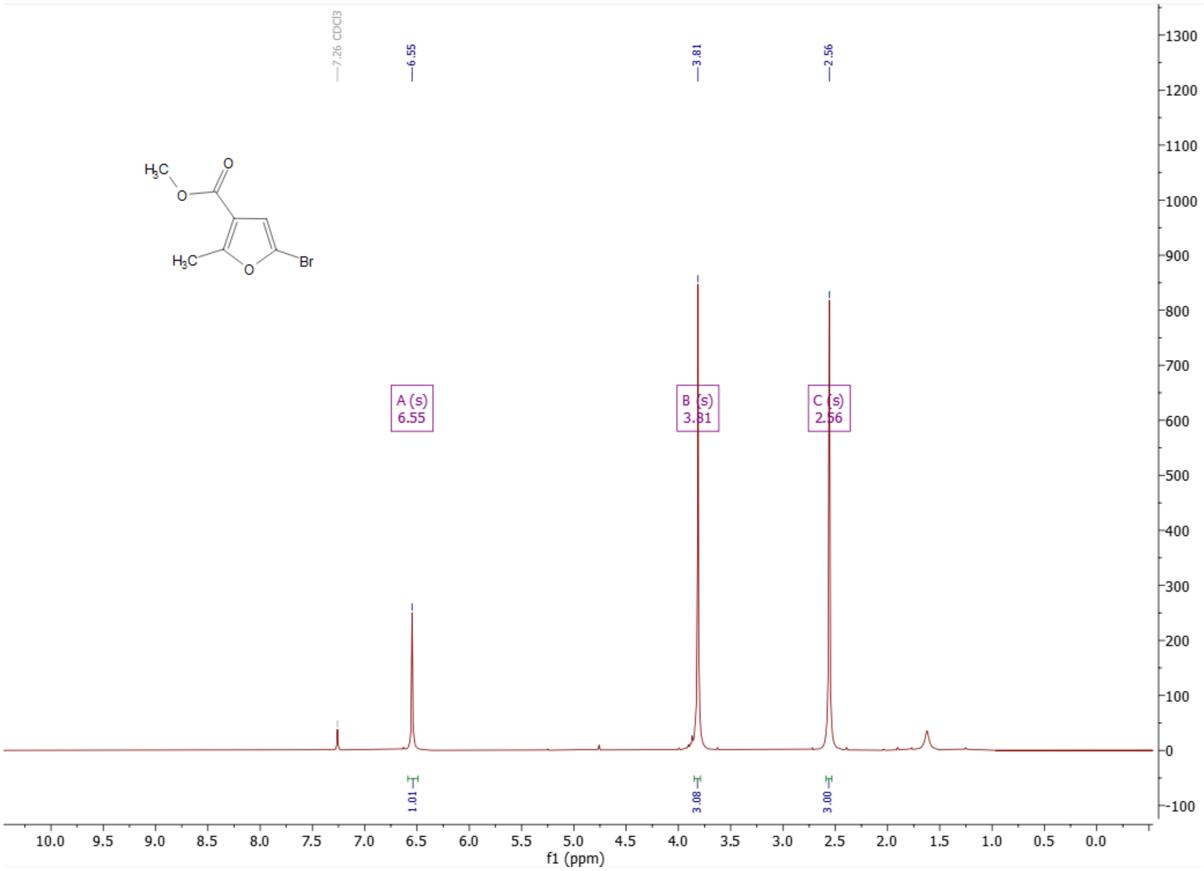

$^1$H-NMR (400 MHz) of 5-bromo-2-methylfuran-3-carboxylic acid (**Ic**) in DMSO-$d_6$

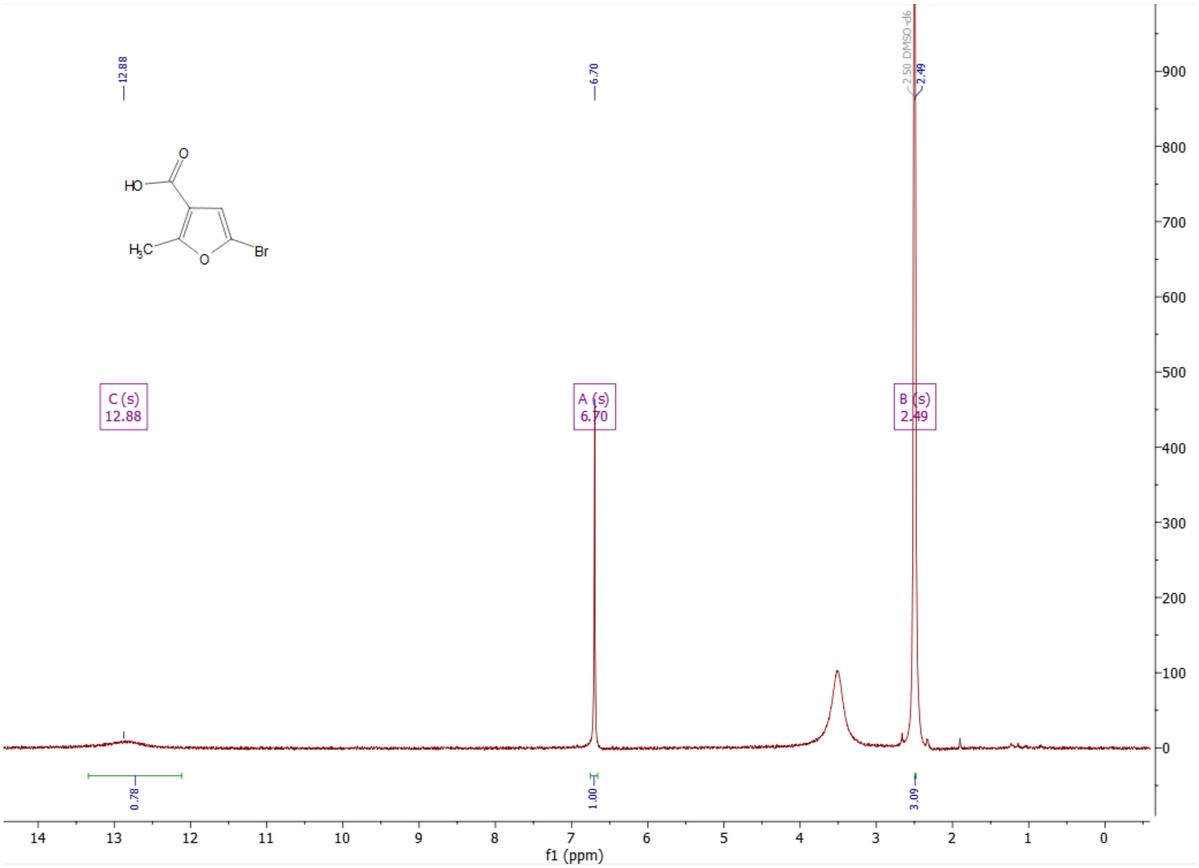

[1]H-NMR (400 MHz) of (Z)-5-(3-((((tert-butoxycarbonyl)amino)(4,4-dimethylpiperidin-1-yl) methylene)amino)-5-fluorophenyl)-2-methylfuran-3-carboxylic acid (**Ie**) in DMSO-$d_6$

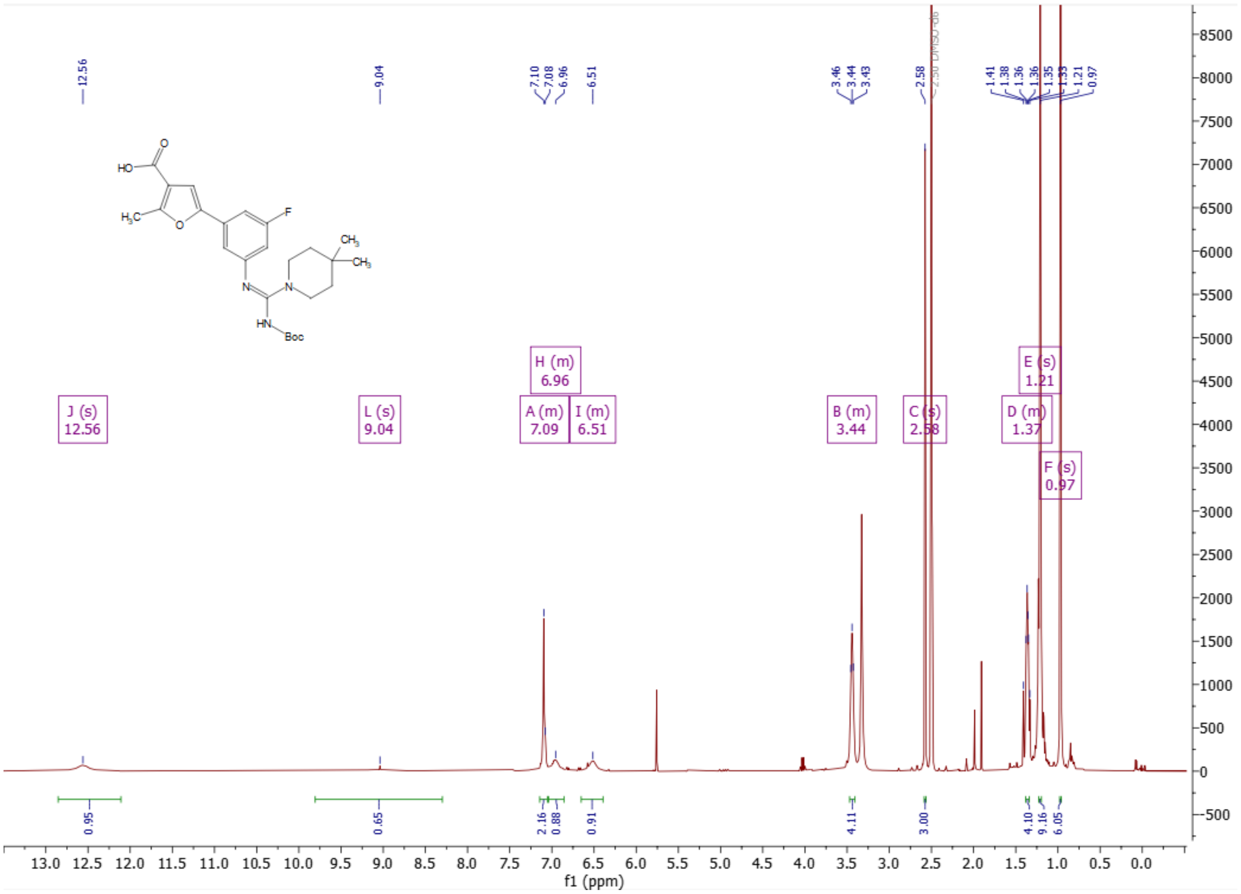

$^{1}H$-NMR (500 MHz) of (Z)-N'-phenylpiperidine-1-carboxímidamide (**1**) in DMSO-$d_6$

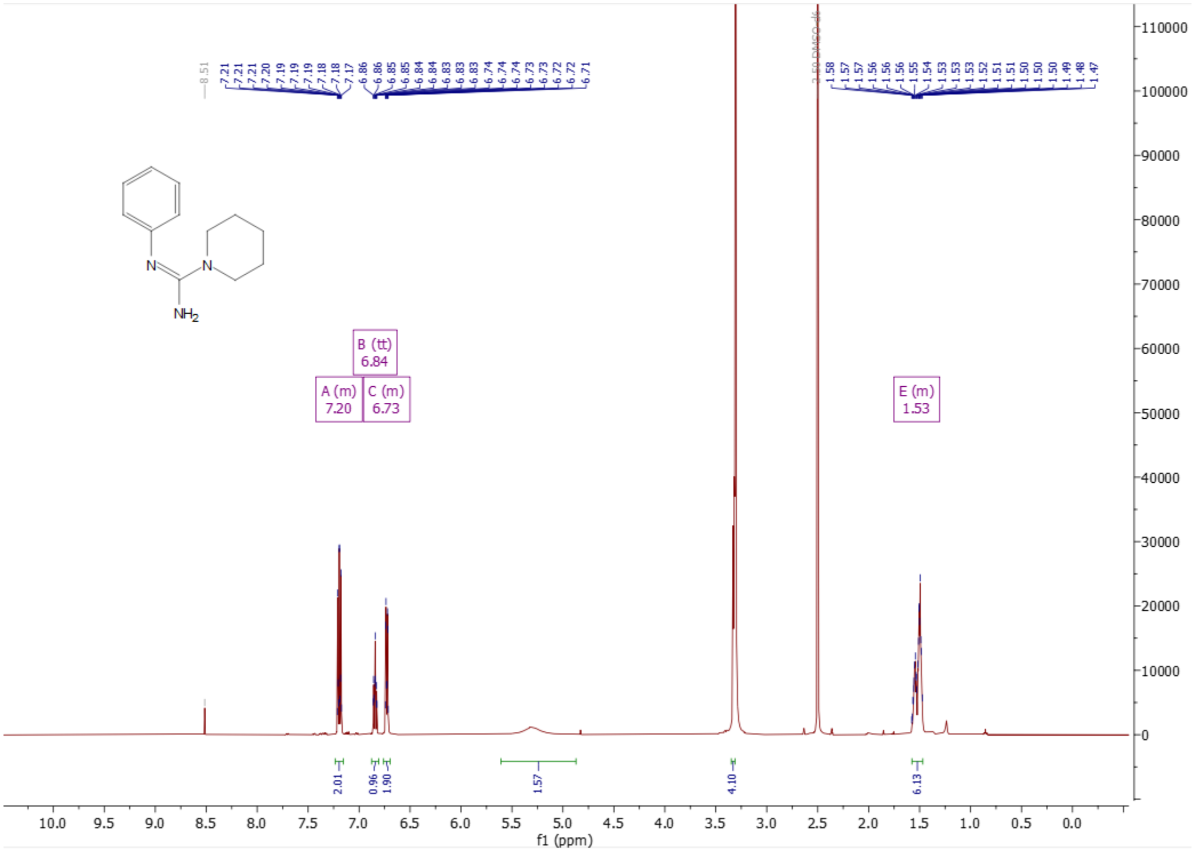

$^{1}$H-NMR (400 MHz) of (Z)-N'-(3-chlorophenyl)piperidine-1-carboximidamide (**2**) in DMSO-$d_6$

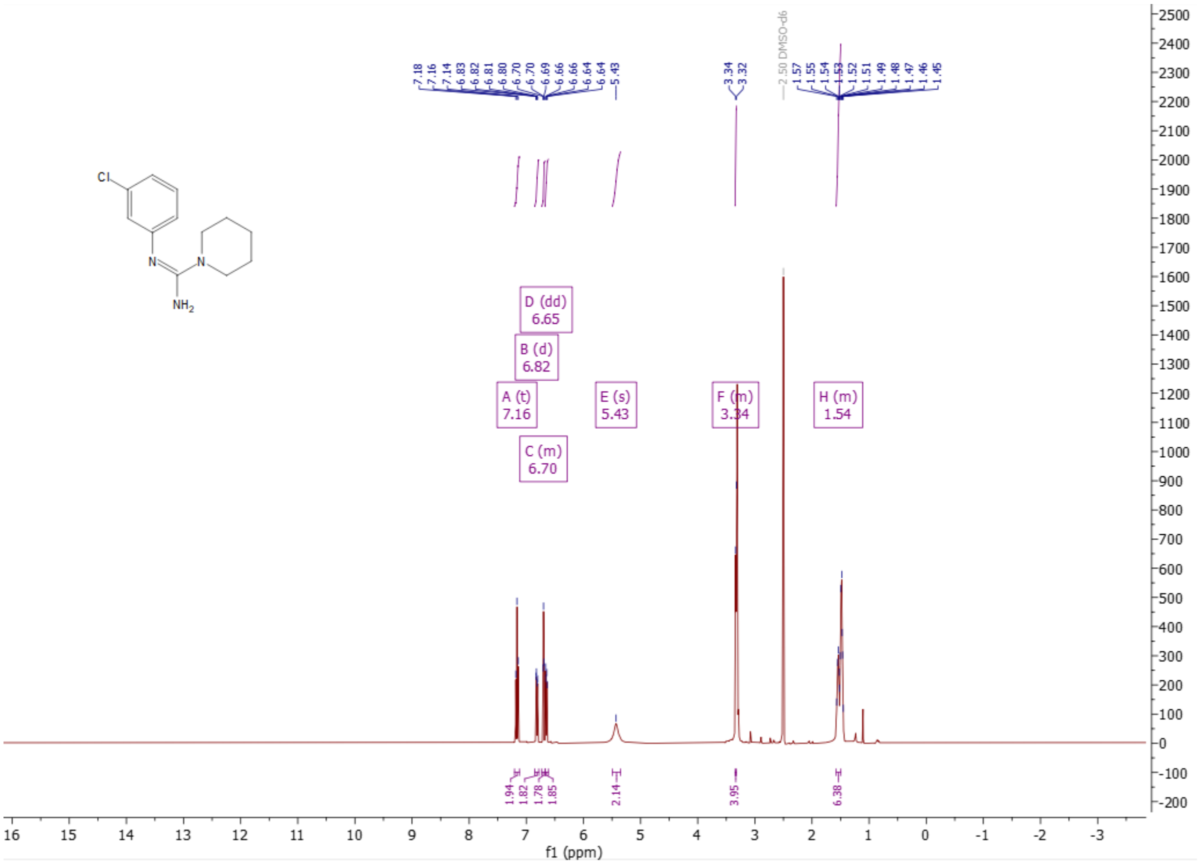

$^{1}$H-NMR (400 MHz) of (Z)-N'-(3-fluorophenyl)piperidine-1-carboximidamide (**3**) in DMSO-$d_6$

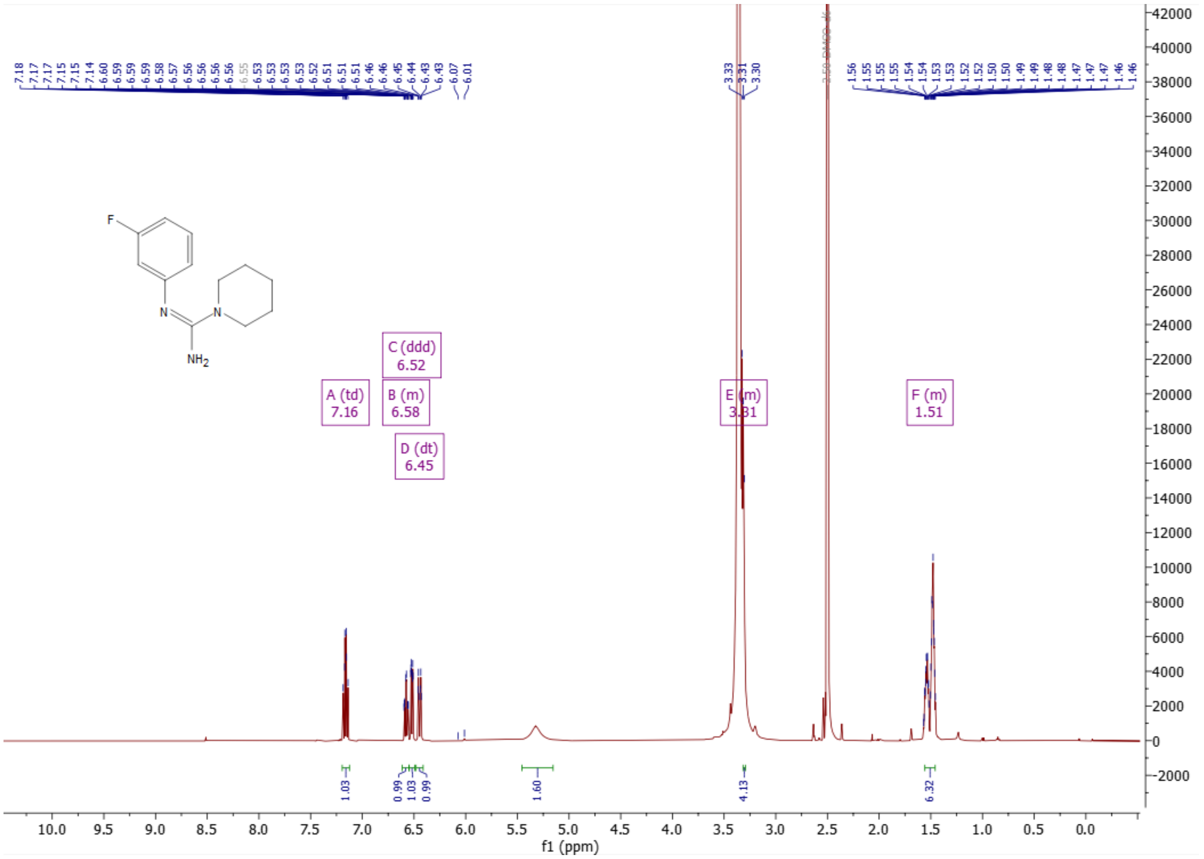

$^1$H-NMR (500 MHz) of N-(3-chloro-5-fluorophenyl)piperidine-1-carboximidamide (**4**) in $CD_3OD$

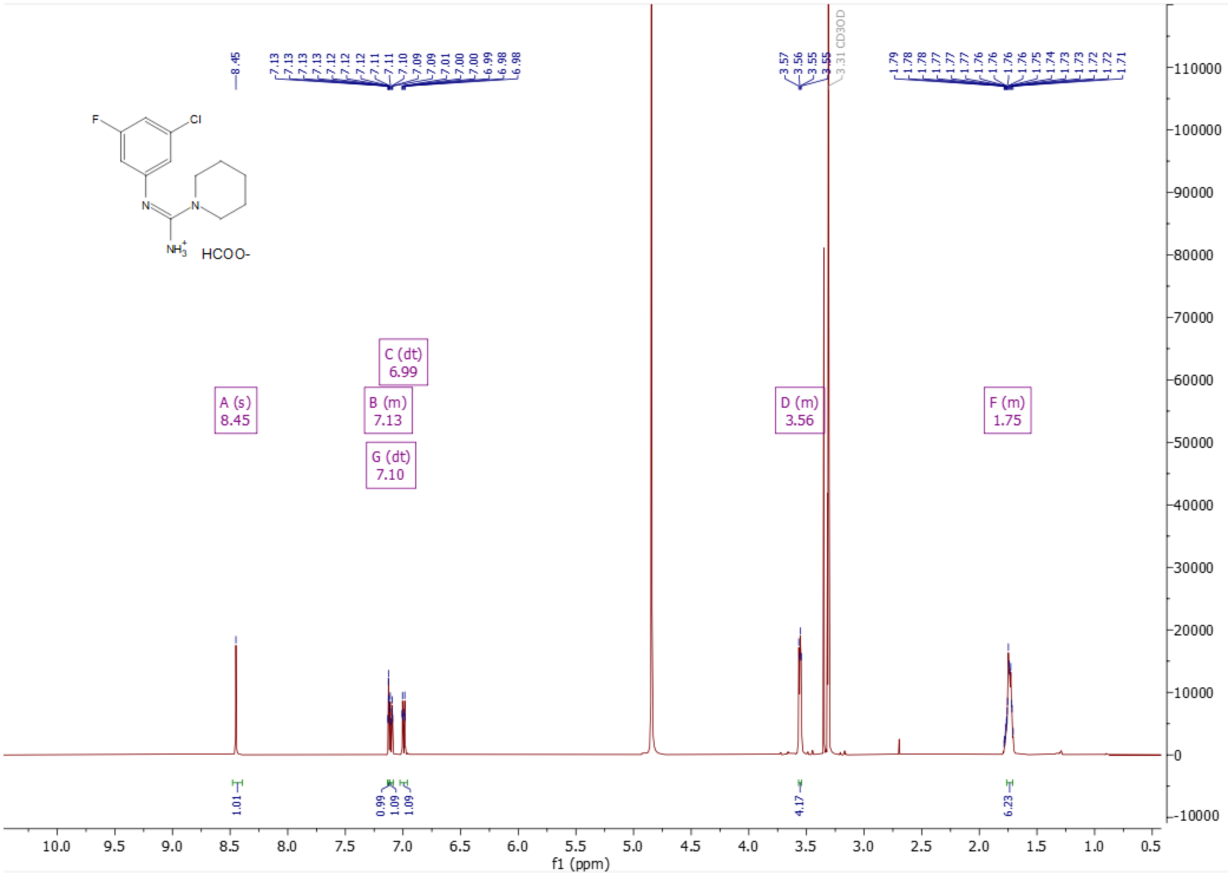

[1]H-NMR (500 MHz) of (N-(3-chloro-5-fluorophenyl)-2-methylpiperidine-1-carboximidamide (**5**) in $CD_3OD$

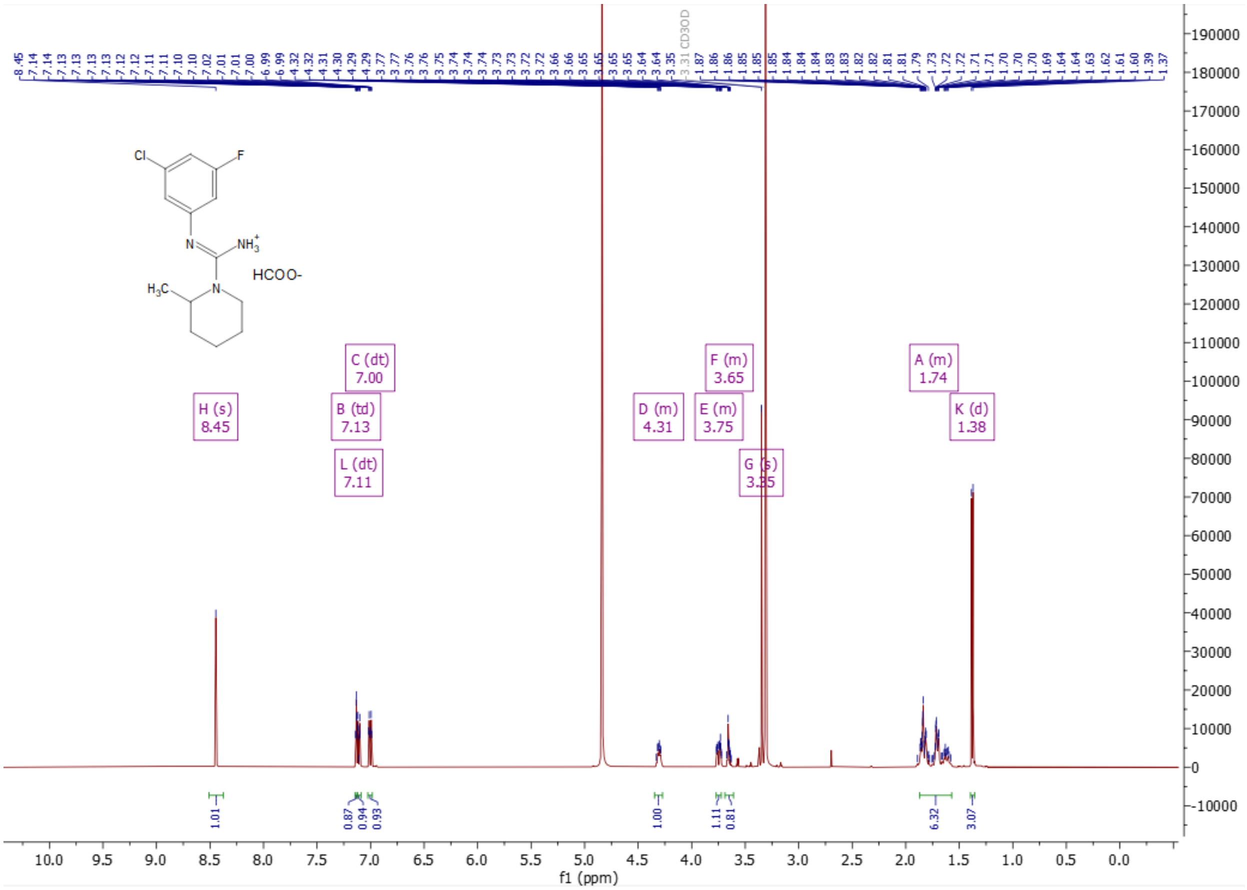

[1]H-NMR (500 MHz) of N-(3-chloro-5-fluorophenyl)-4-methylpiperidine-1-carboximidamide (**6**) in $CD_3OD$

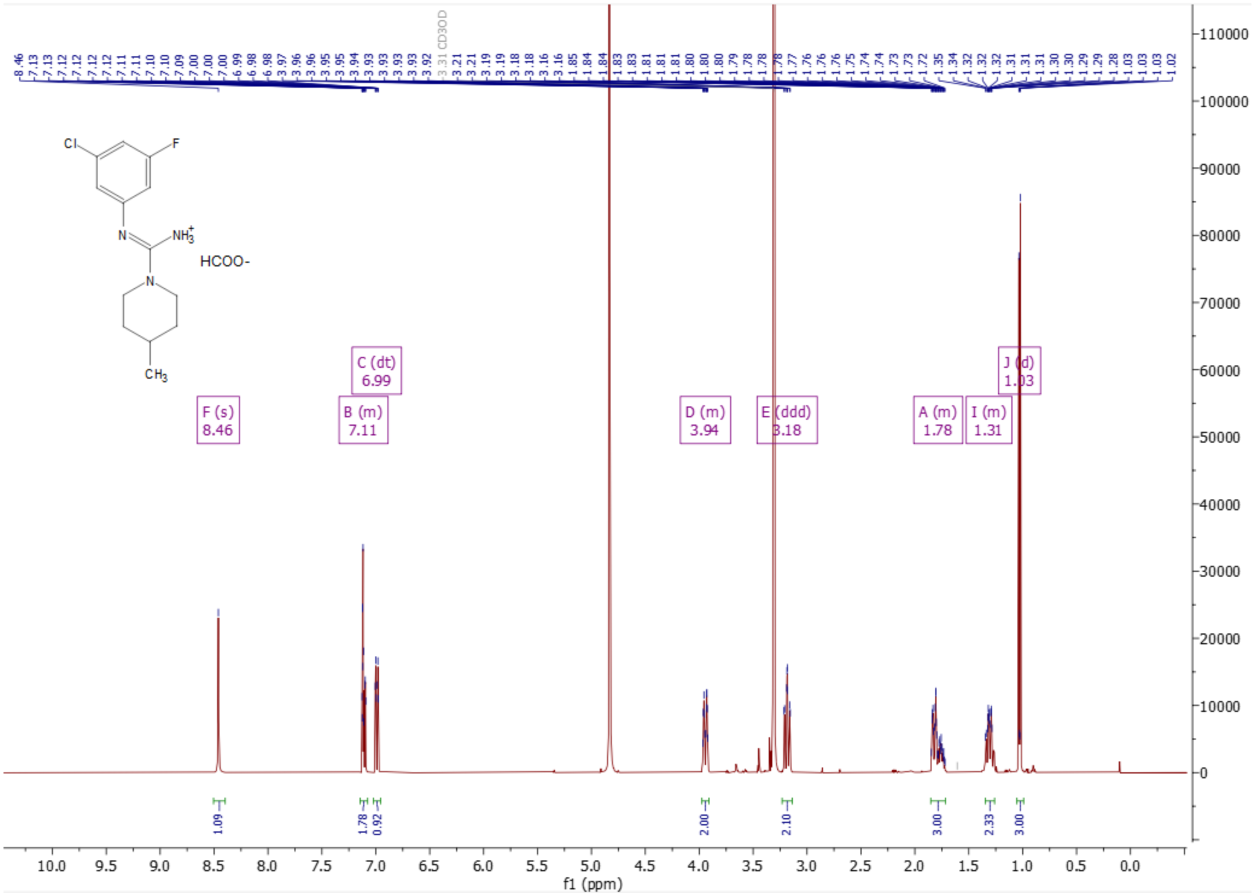

$^{1}$H-NMR (500 MHz) of N-(3-chloro-5-fluorophenyl)-3-methylpiperidine-1-carboximidamide (**7**) in $CD_3OD$

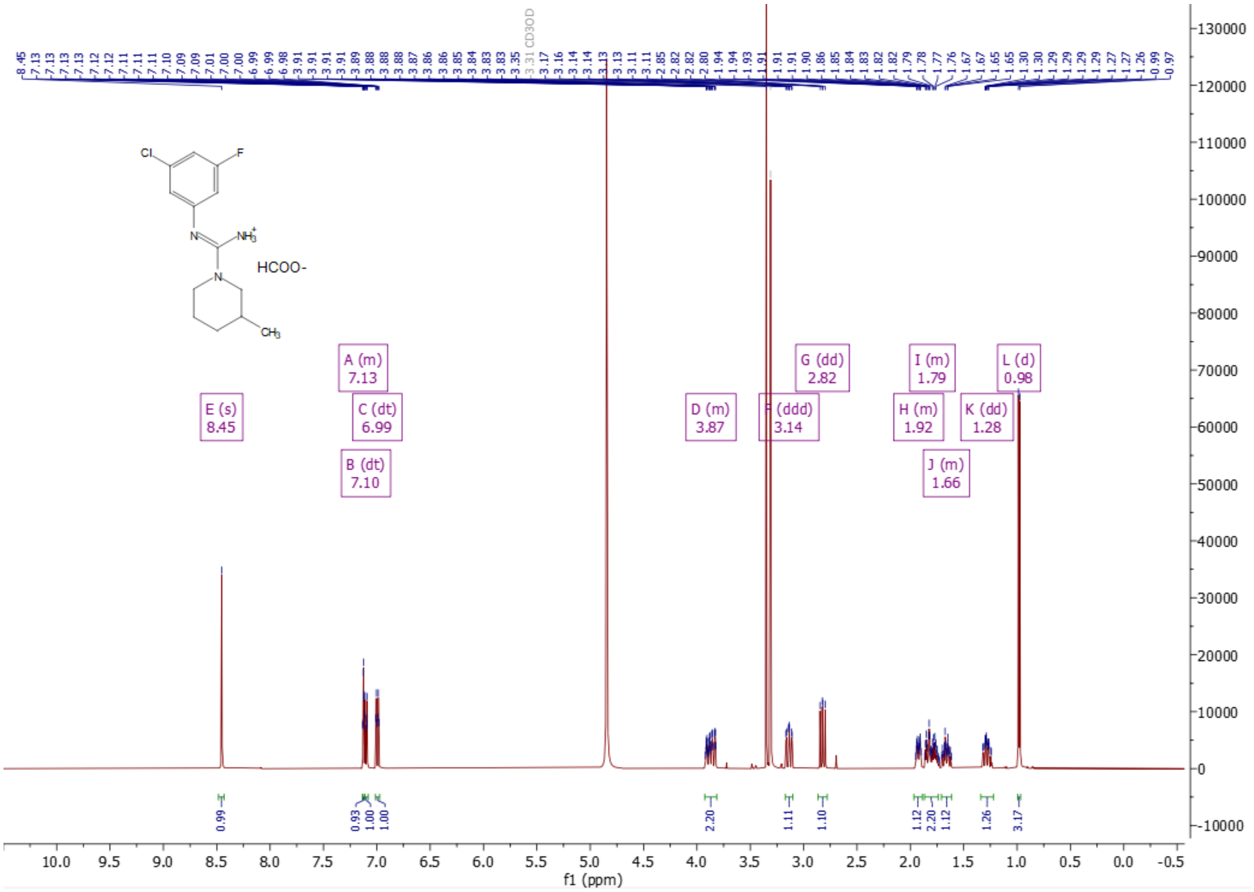

[1]H-NMR (500 MHz) of (Z)-N'-(3-fluoro-5-(5-methylfuran-2-yl)phenyl)-2-methylpiperidine-1-carboximidamide (**8**) in $CD_3OD$

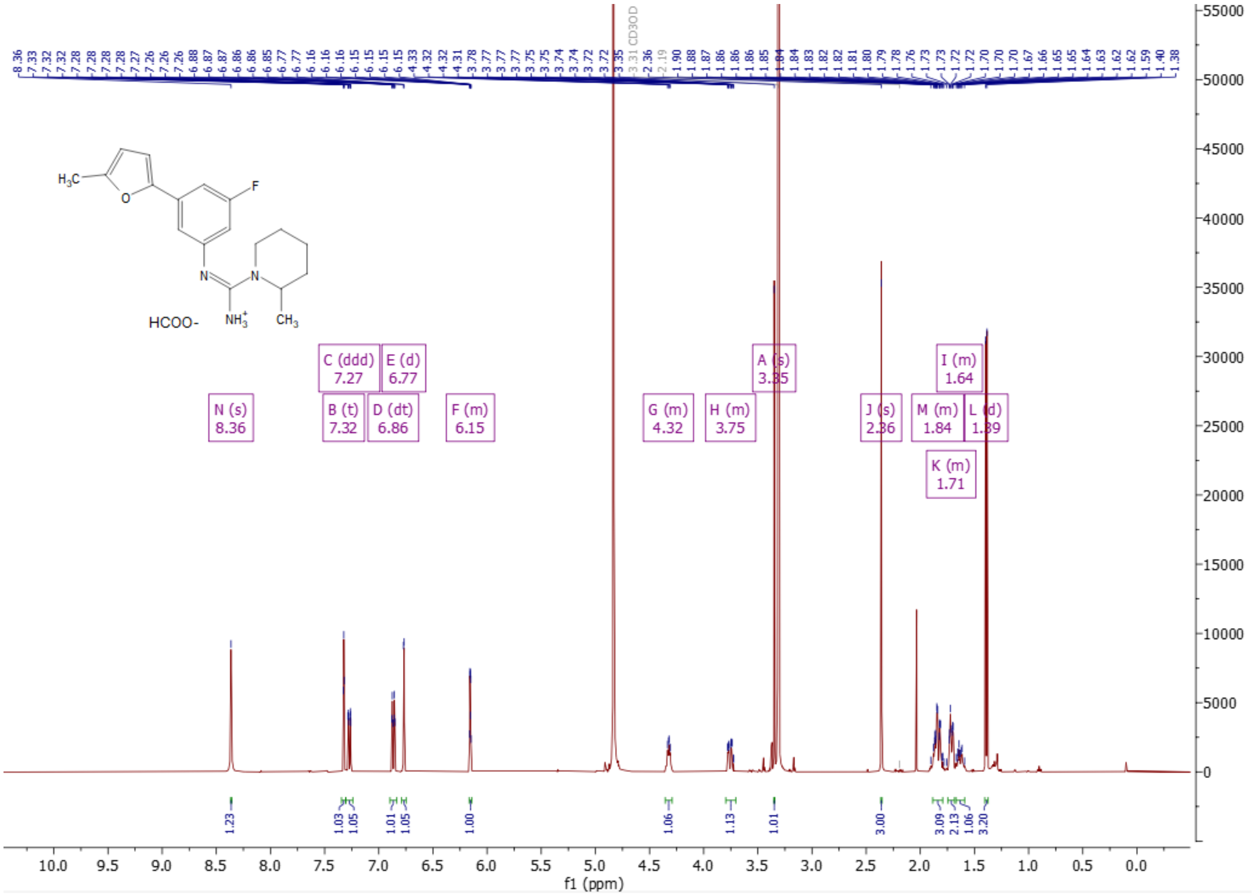

[1]H-NMR (500 MHz) of (Z)-N'-(3-fluoro-5-(5-methylfuran-2-yl)phenyl)-4-methylpiperidine-1-carboximidamide (**9**) in $CD_3OD$

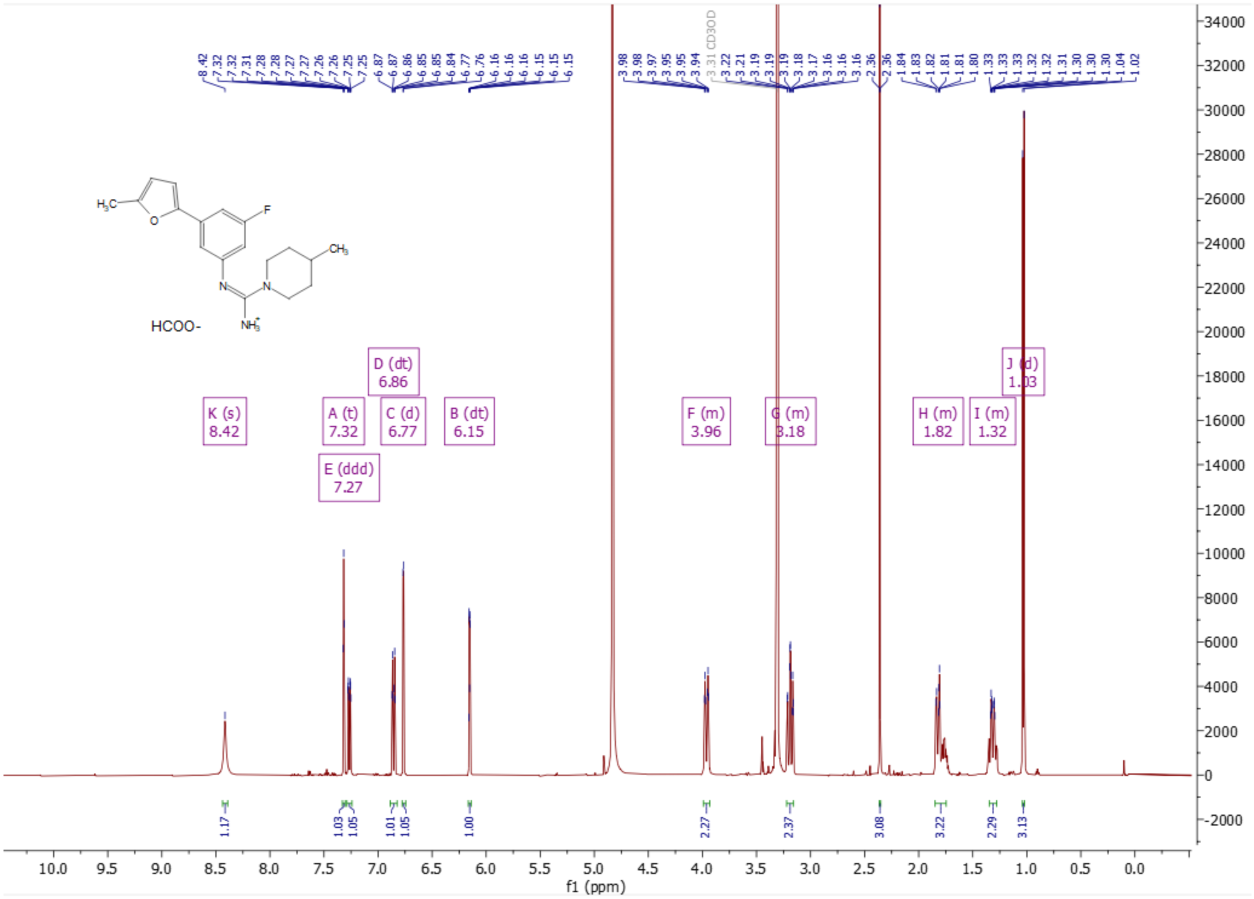

$^{1}H$-NMR (500 MHz) of (Z)-N'-(3-fluoro-5-(5-methylfuran-2-yl)phenyl)-4,4-dimethylpiperidine-1-carboxamidamide (**10**) in $CD_3OD$

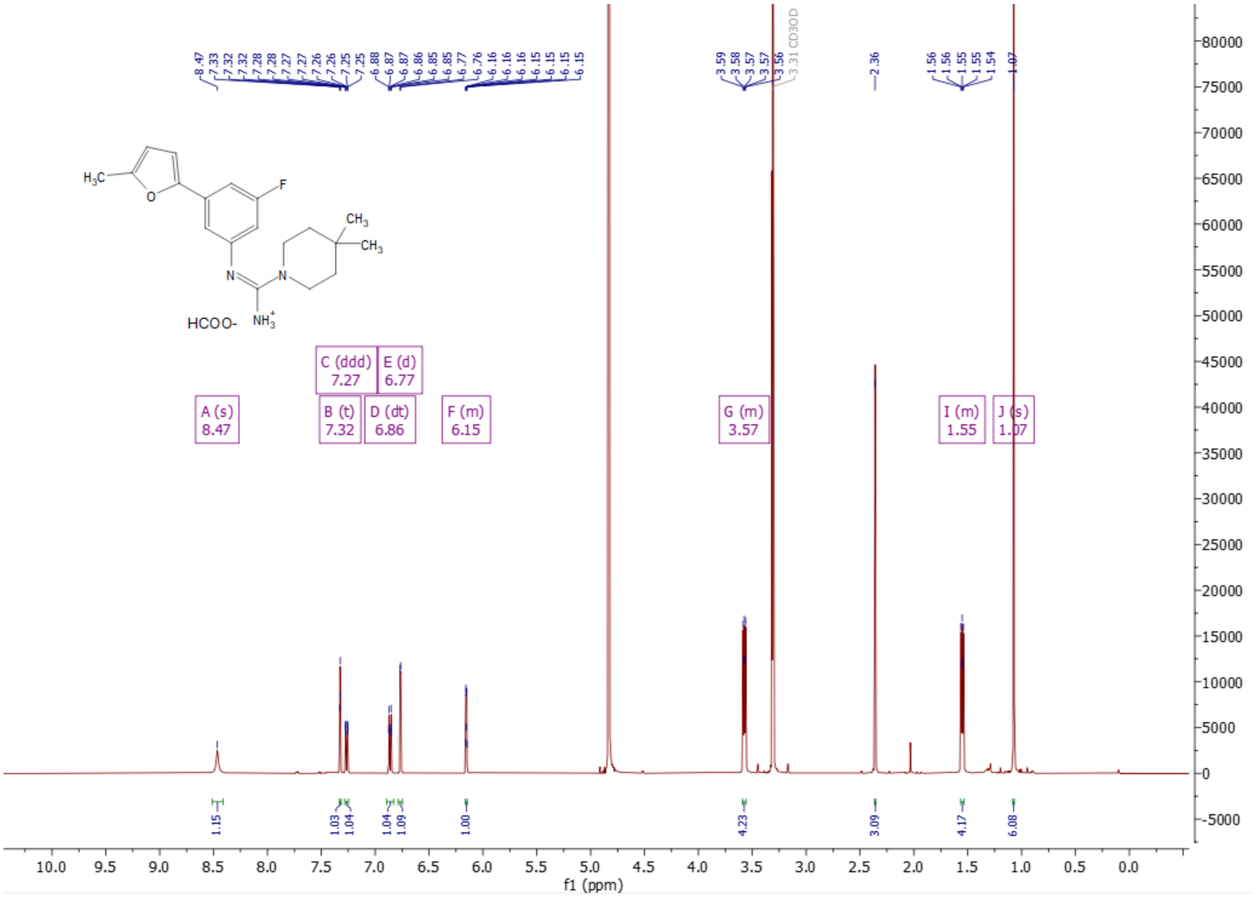

[1]H-NMR (500 MHz) of (Z)-5-(3-((amino(4,4-dimethylpiperidin-1-yl)methylene)amino)-5-fluorophenyl)-N,2-dimethylfuran-3-carboxamide (**11**) in $CD_3OD$

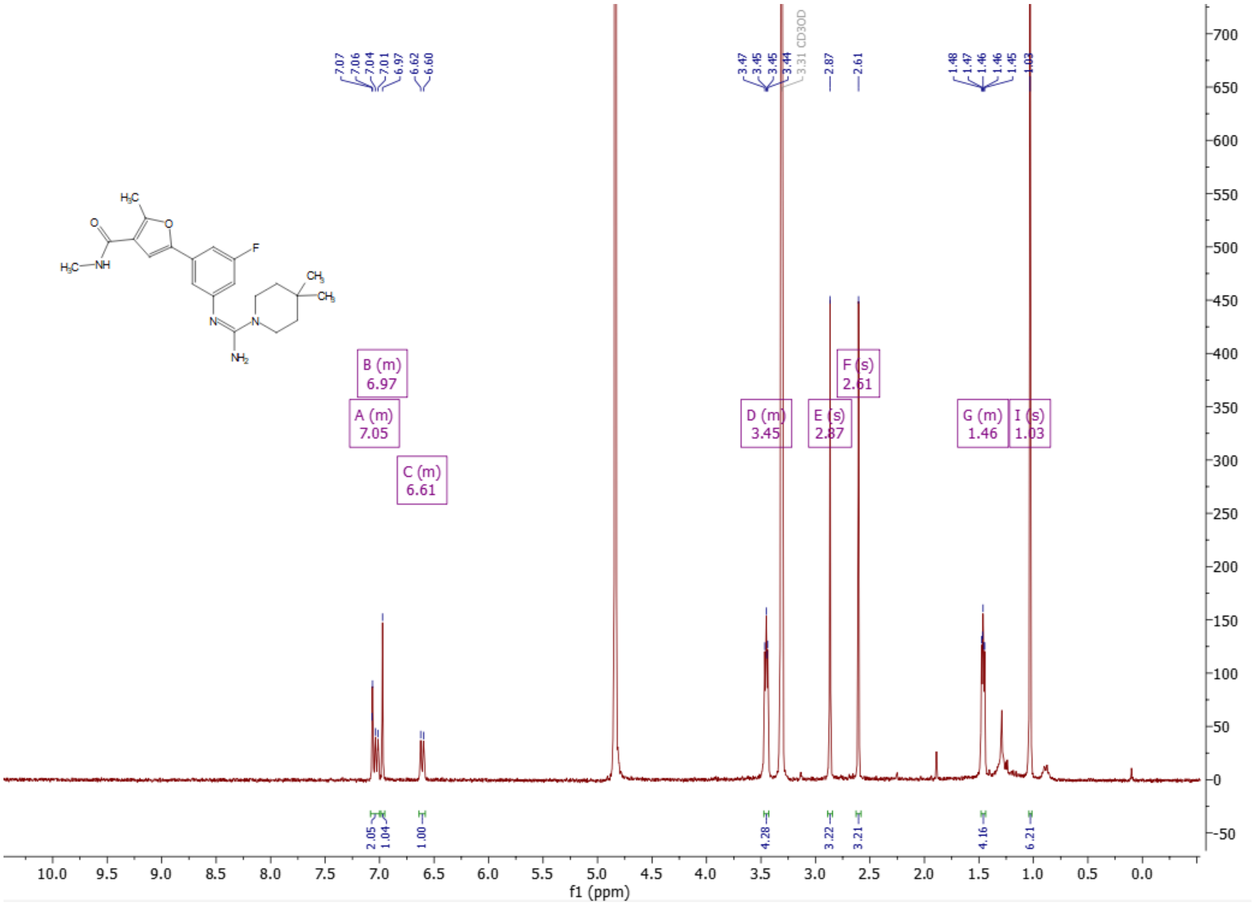

[1]H-NMR (500 MHz) of (Z)-5-(3-((amino(4,4-dimethylpiperidin-1-yl)methylene)amino)-5-fluorophenyl)-N,N,2-trimethylfuran-3-carboxamide (**12**) in $CD_3OD$

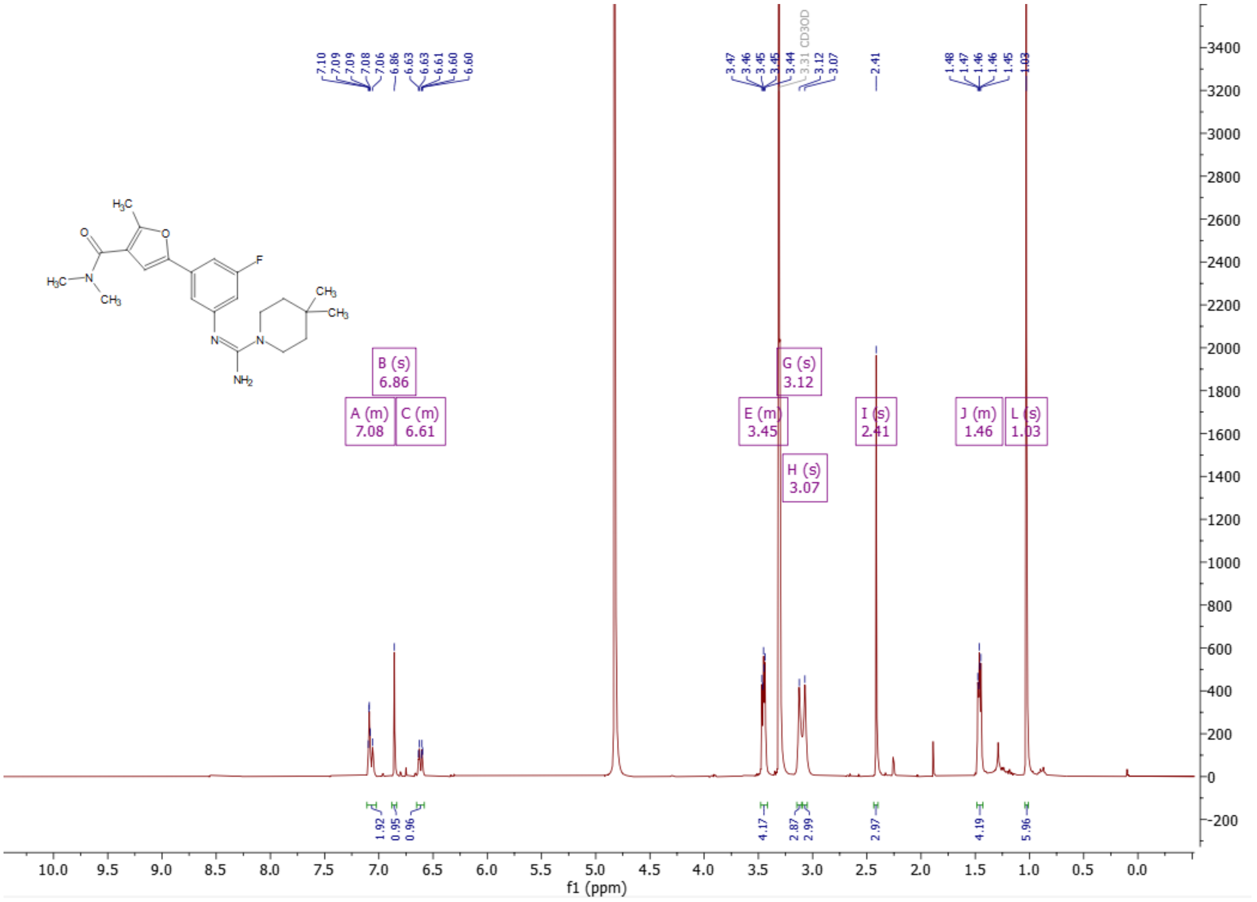

[1]H-NMR (500 MHz) of (Z)-5-(3-((amino(4,4-dimethylpiperidin-1-yl)methylene)amino)-5-fluorophenyl)-2-methyl-N-phenylfuran-3-carboxamide (**13**) in DMSO-$d_6$

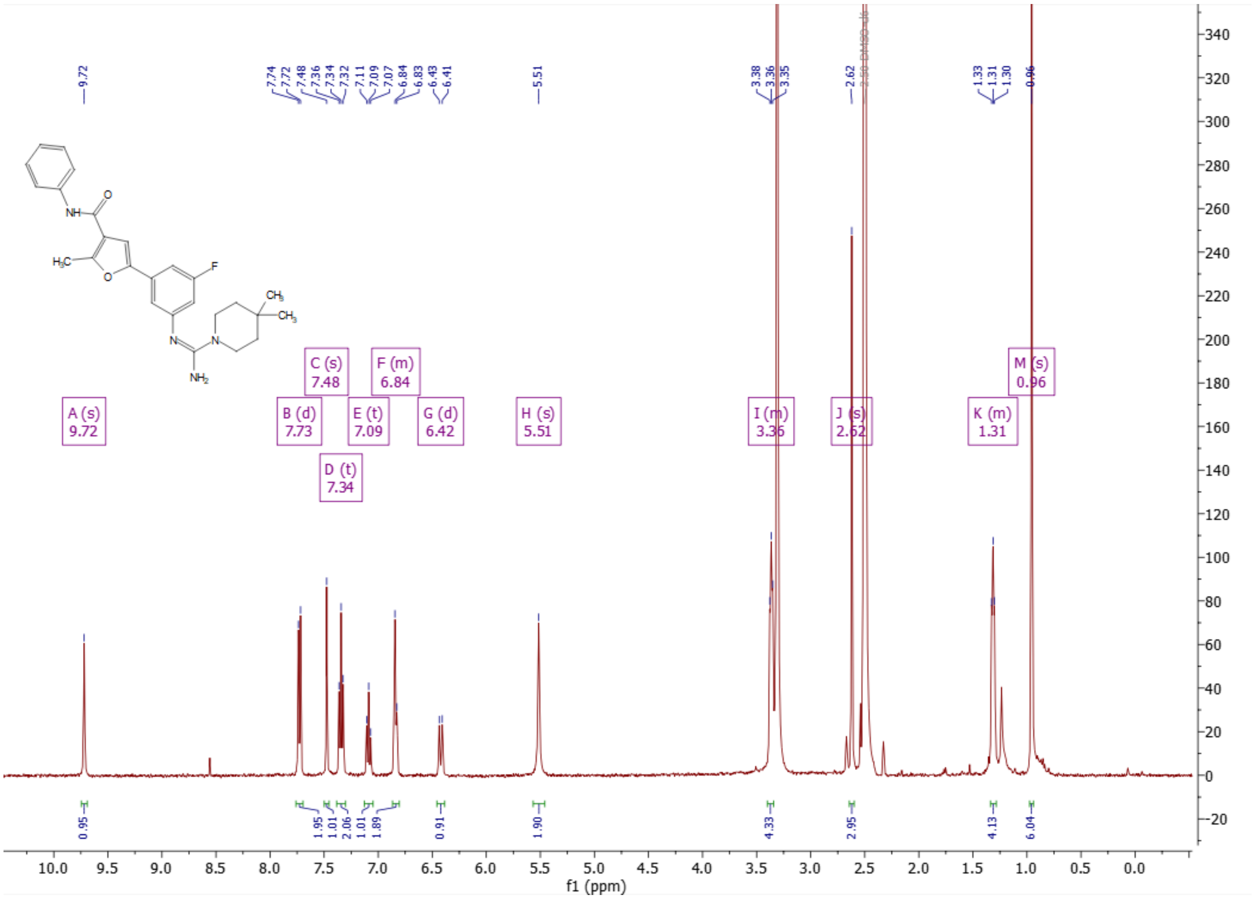

[1]H-NMR (500 MHz) of (Z)-N'-(3-fluoro-5-(5-methyl-4-(piperidine-1- carbonyl)furan-2-yl)phenyl)-4,4-dimethylpiperidine-1-carboximidamide formate (**14**) in $CD_3OD$

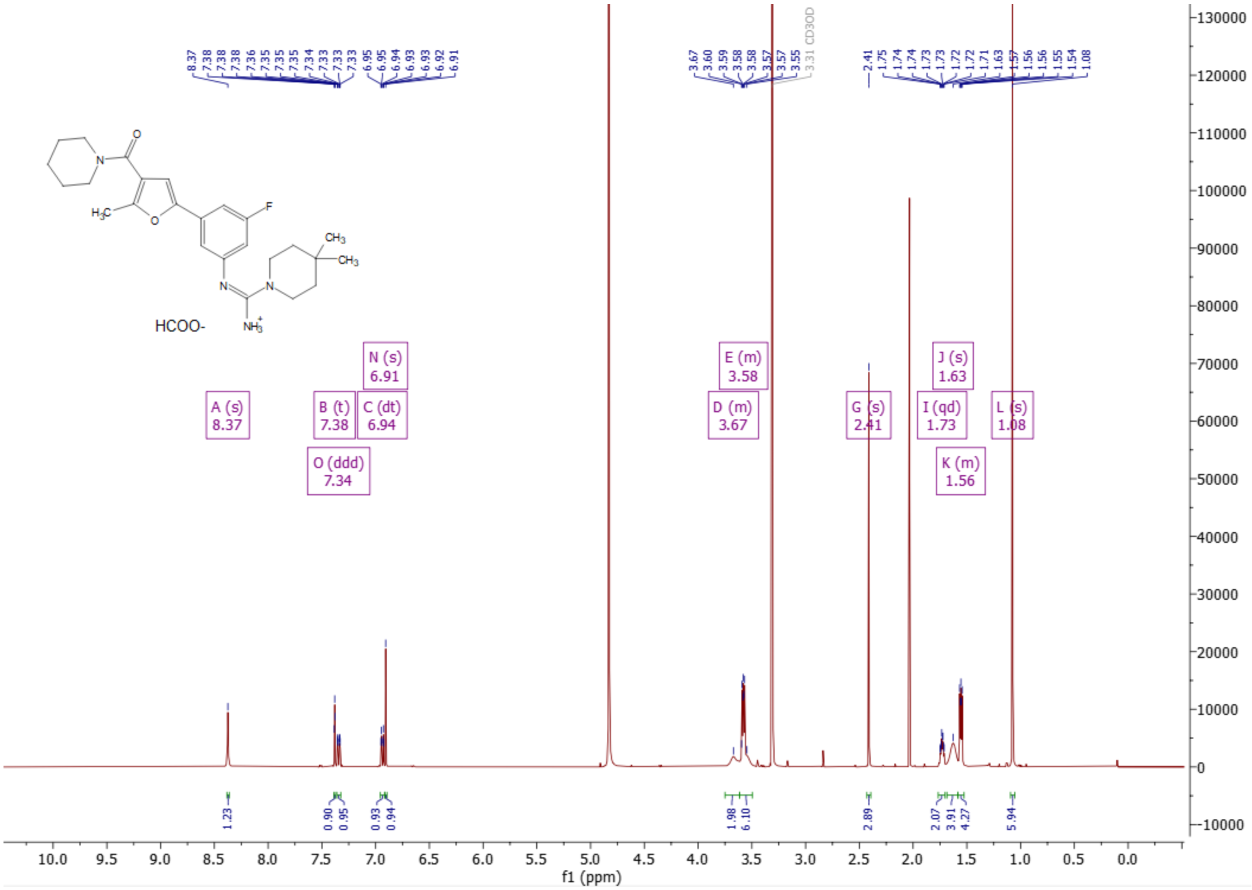

[1]H-NMR (400 MHz) of (Z)-N'-(3-fluoro-5-(5-methyl-4-(4-methylpiperazine-1-carbonyl)furan-2-yl)phenyl)-4,4-dimethylpiperidine-1-carboximidamide (**15**) in DMSO-$d_6$

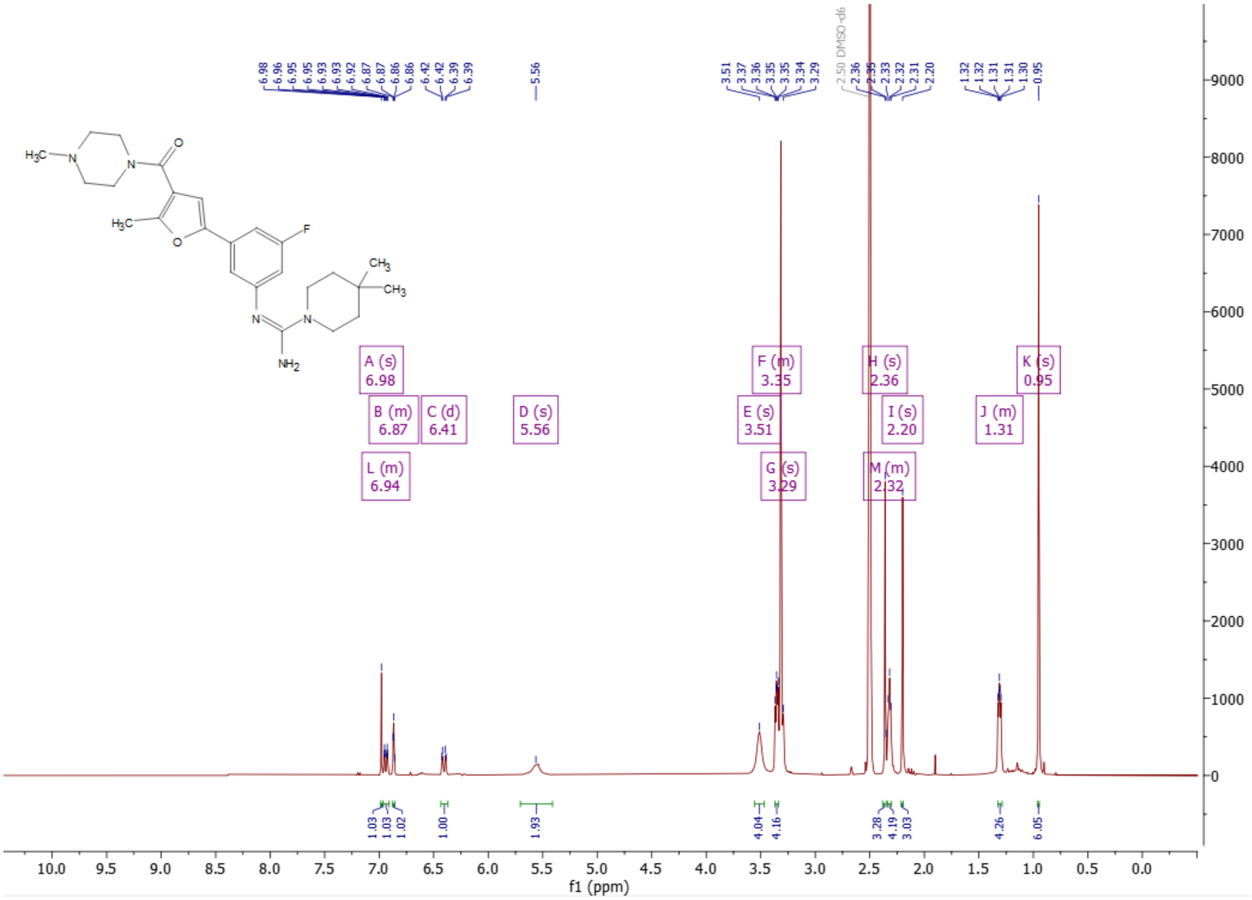

[1]H-NMR (500 MHz) of (Z)-N'-(3-fluoro-5-(5-methyl-4-(morpholine-4-carbonyl)furan-2-yl)phenyl)-4,4-dimethylpiperidine-1-carboximidamide formate (**16**) in $CD_3OD$

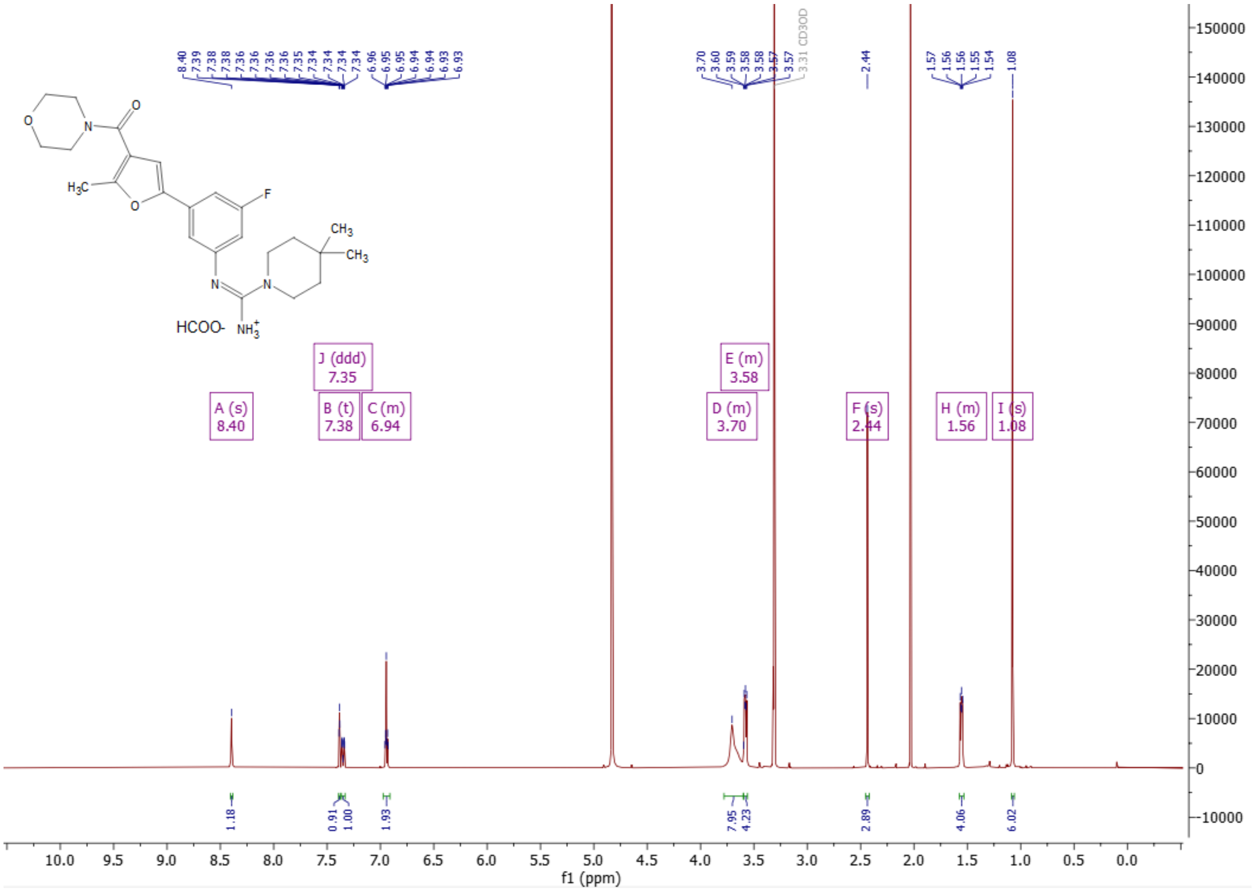

[1]H-NMR (500 MHz) of (Z)-N'-(3-fluoro-5-(5-methyl-4-(pyrrolidine-1-carbonyl)furan-2-yl)phenyl)-4,4-dimethylpiperidine-1-carboximidamide formate (**17**) in DMSO-$d_6$

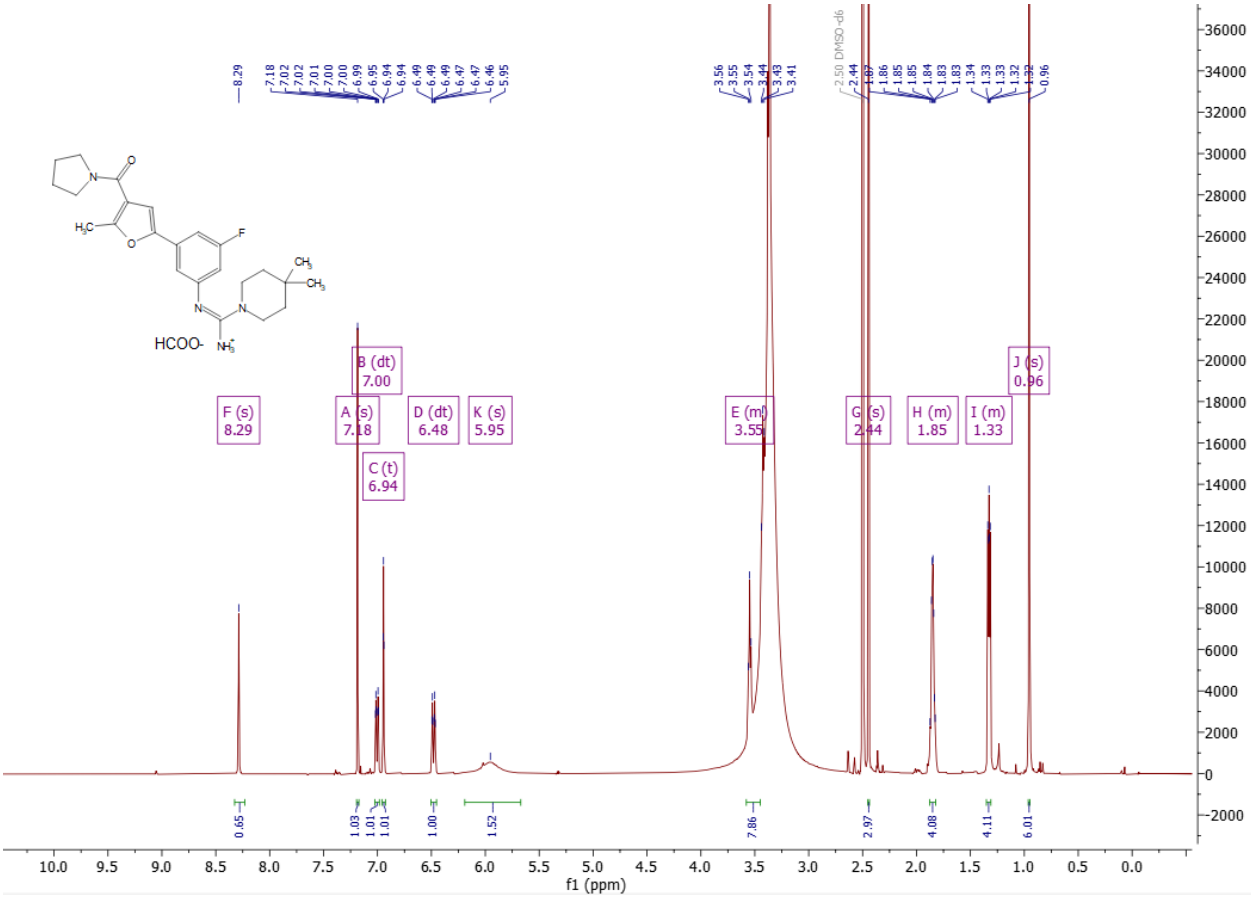

$^{1}$H-NMR (500 MHz) of (Z)-N'-{3-fluoro-5-[4-(4-methoxypiperidine-1-carbonyl)-5-methylfuran-2-yl]phenyl}-4,4-dimethylpiperidine-1-carboximidamide formate (**18**) in $CD_3OD$

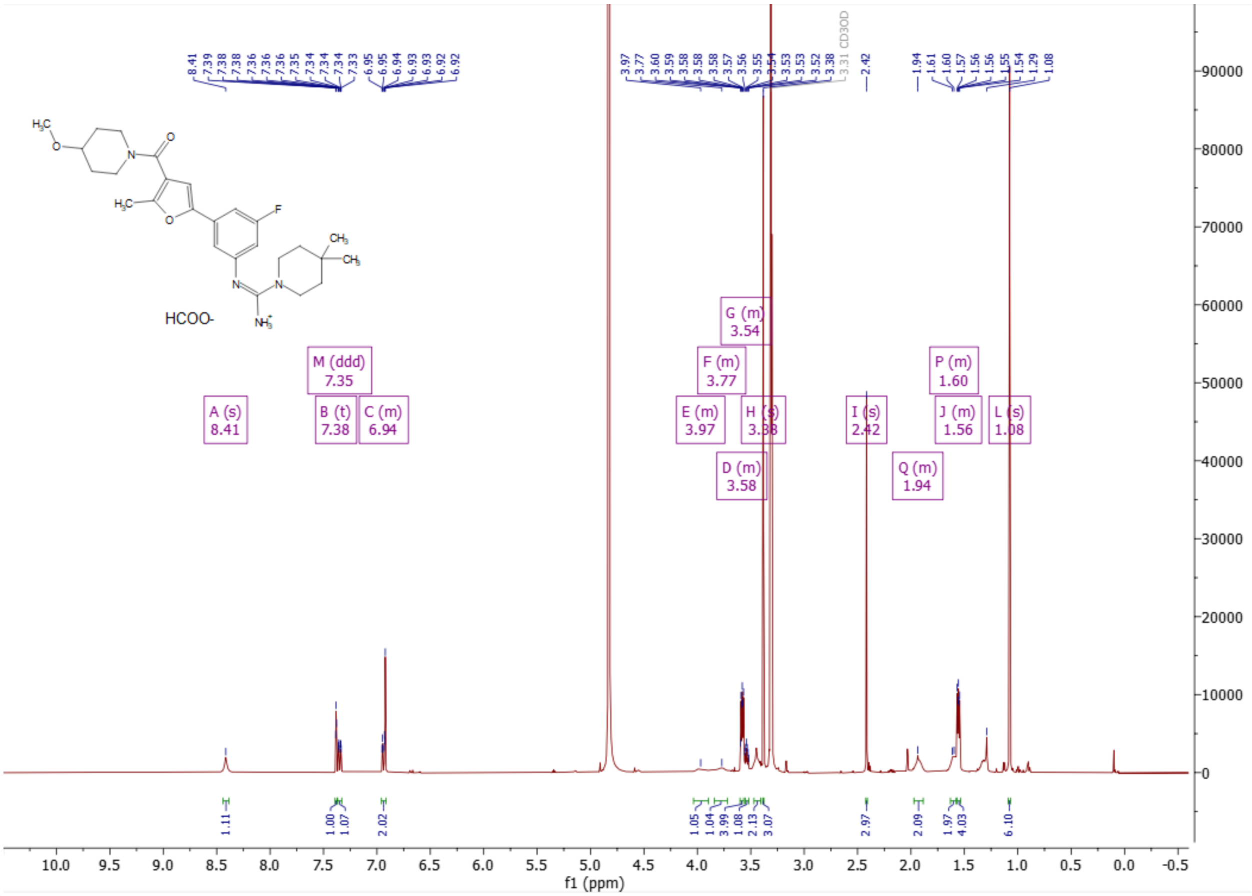

[1]H-NMR (500 MHz) of (Z)-N'-{3-fluoro-5-[4-(3-methoxypiperidine-1-carbonyl)-5-methylfuran-2-yl]phenyl}-4,4-dimethylpiperidine-1-carboximidamide (**19**) in DMSO-$d_6$ at 80°C

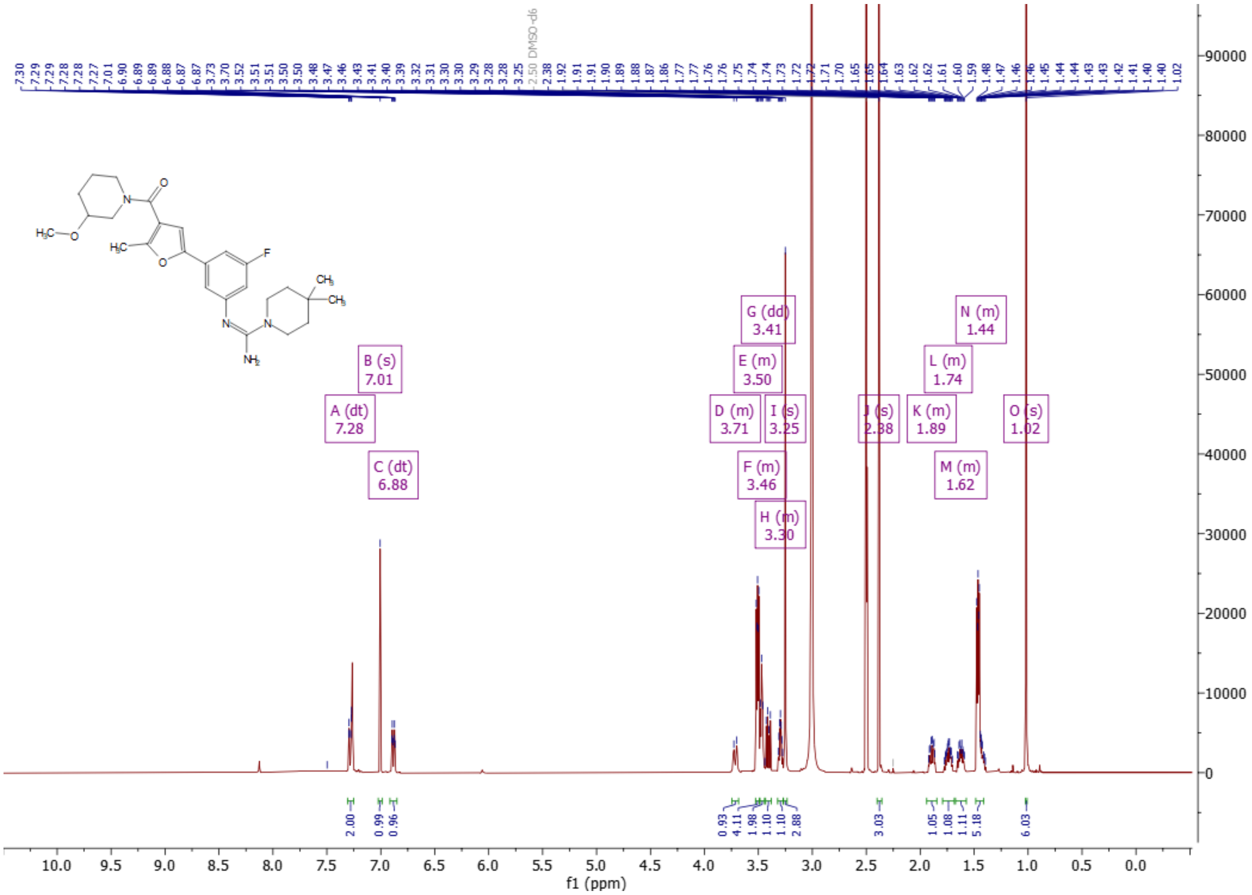

$^1$H-NMR (500 MHz) of (Z)-N'-[3-fluoro-5-(4-{4-[2-(2-methoxyethoxy)ethyl]piperidine-1-carbonyl}-5-methylfuran-2-yl)phenyl]-4,4-dimethylpiperidine-1-carboximidamide (**20**) in DMSO-$d_6$

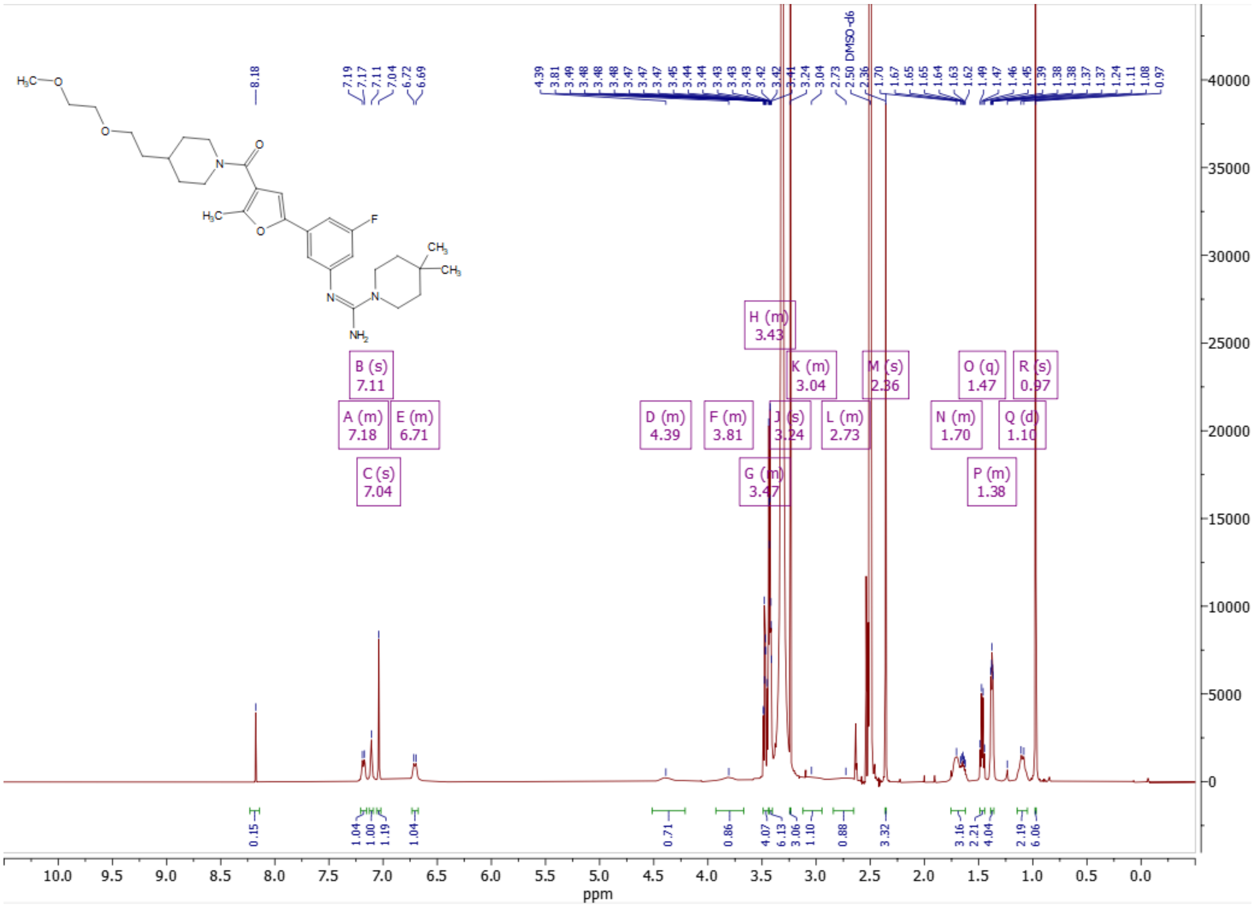

$^1$H-NMR (500 MHz) of (Z)-N'-[3-fluoro-5-(4-{3-[2-(2-methoxyethoxy)ethyl]piperidine-1-carbonyl}-5-methylfuran-2-yl)phenyl]-4,4-dimethylpiperidine-1-carboximidamide (**21**) in DMSO-$d_6$

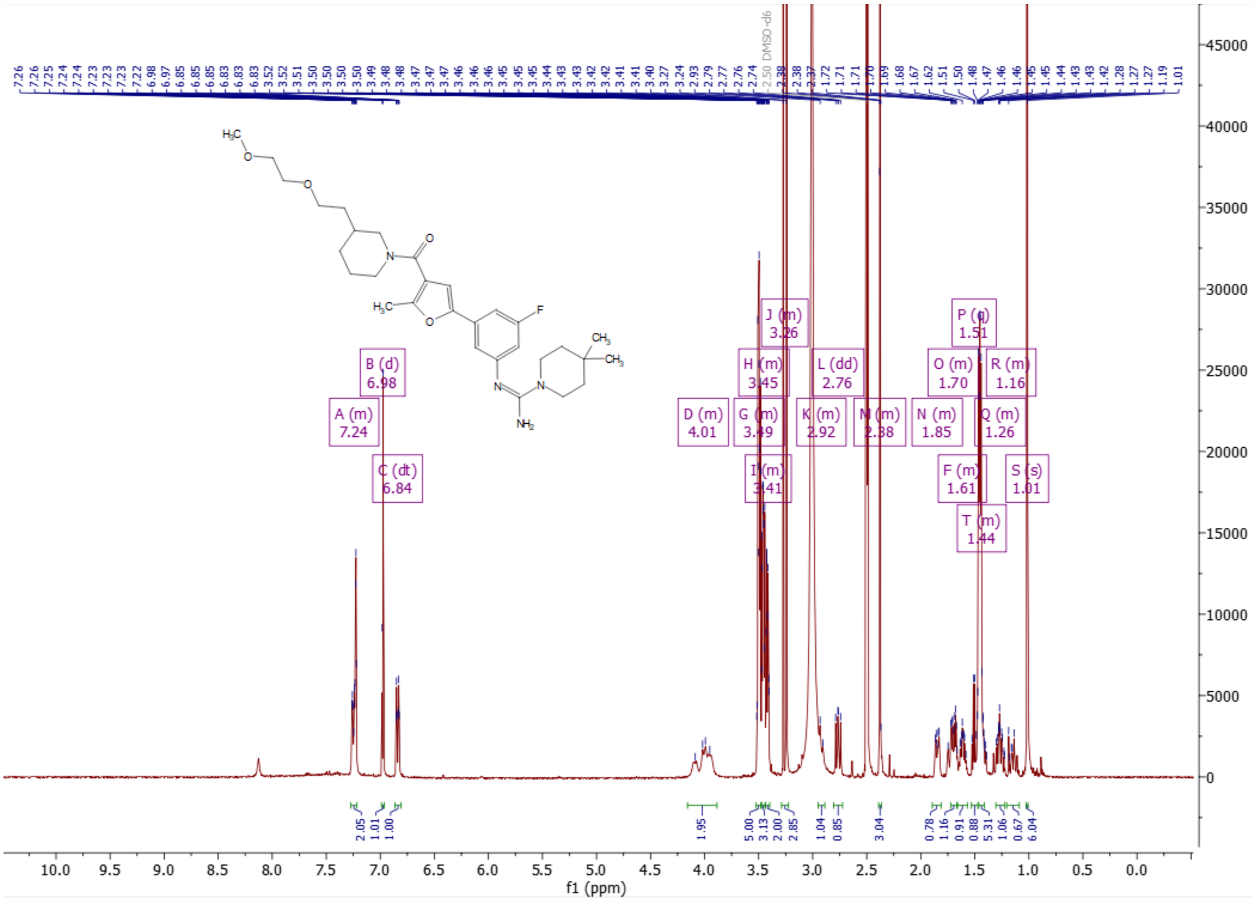

$^{1}$H-NMR (500 MHz) of (Z)-N'-[3-(4-{4-[(4-acetylpiperazin-1-yl)methyl]piperidine-1-carbonyl}-5-methylfuran-2-yl)-5-fluorophenyl]-4,4-dimethylpiperidine-1-carboximidamide formate (**22**) in DMSO-$d_6$

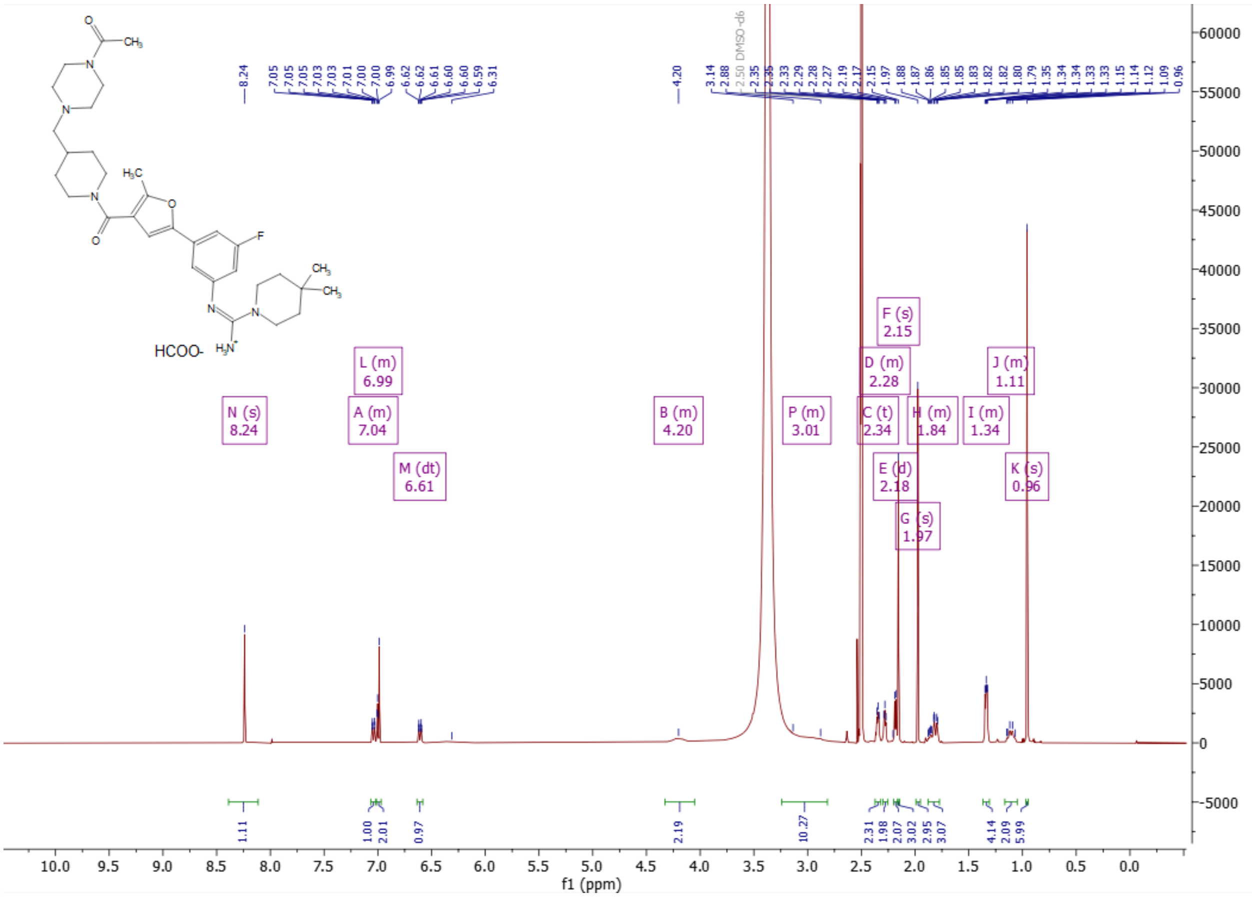

[1]H-NMR (500 MHz) of (Z)-N'-[3-(4-{3-[(4-acetylpiperazin-1-yl)methyl]piperidine-1-carbonyl}-5-methylfuran-2-yl)-5-fluorophenyl]-4,4-dimethylpiperidine-1-carboximidamide formate (**23**) in DMSO-$d_6$ at 80°C

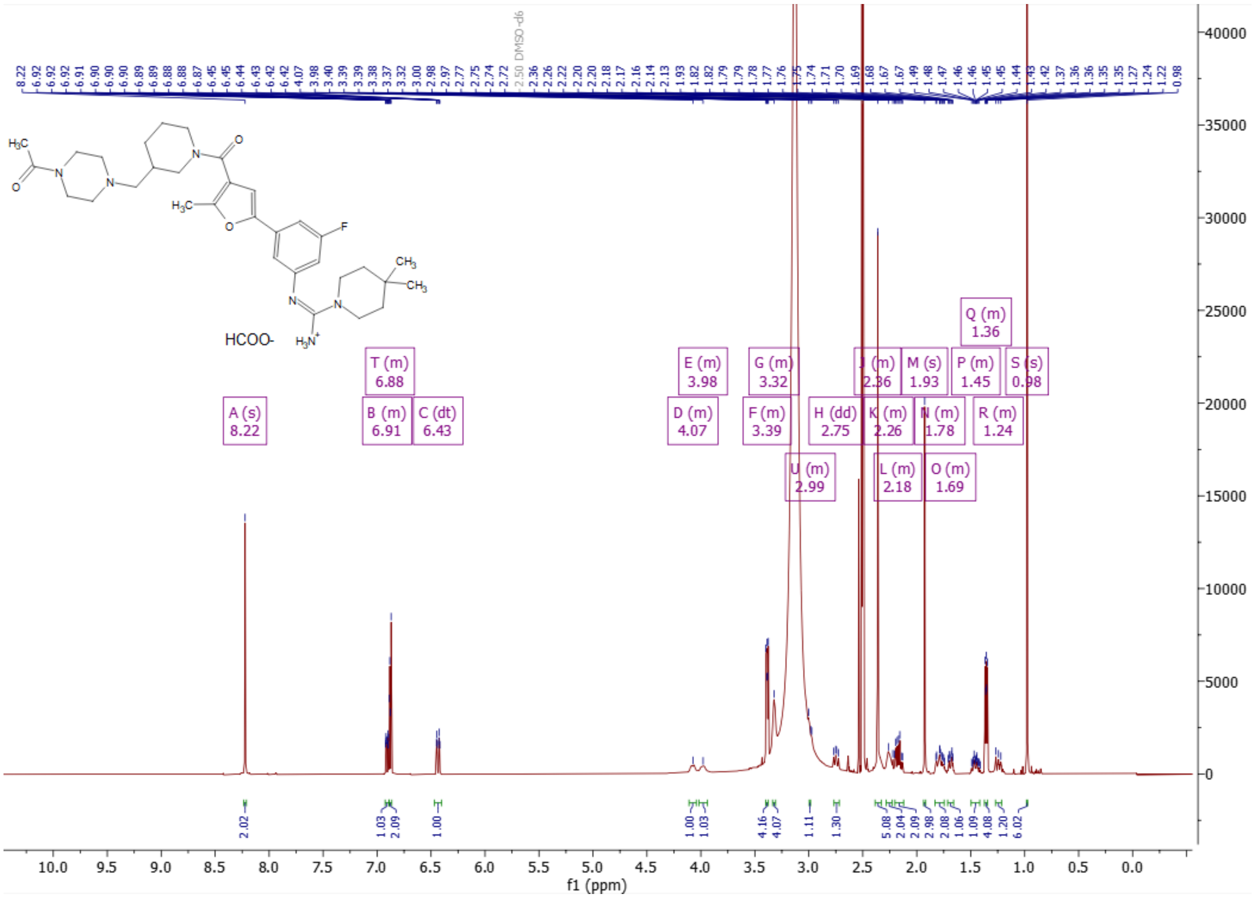

[1]H-NMR (500 MHz) of (Z)-N'-[3-(4-{4-[(4-{2-[(9R)-7-(4-chlorophenyl)-4,5,13-trimethyl-3-thia-1,8,11,12-tetraazatricyclo[8.3.0.0$^{2}$,$^{6}$]trideca-2(6),4,7,10,12-pentaen-9-yl]acetyl}piperazin-1-yl)methyl]piperidine-1-carbonyl}-5-methylfuran-2-yl)-5-fluorophenyl]-4,4-dimethylpiperidine-1-carboximidamide formate (**24**) in DMSO-$d_6$ at 80°C

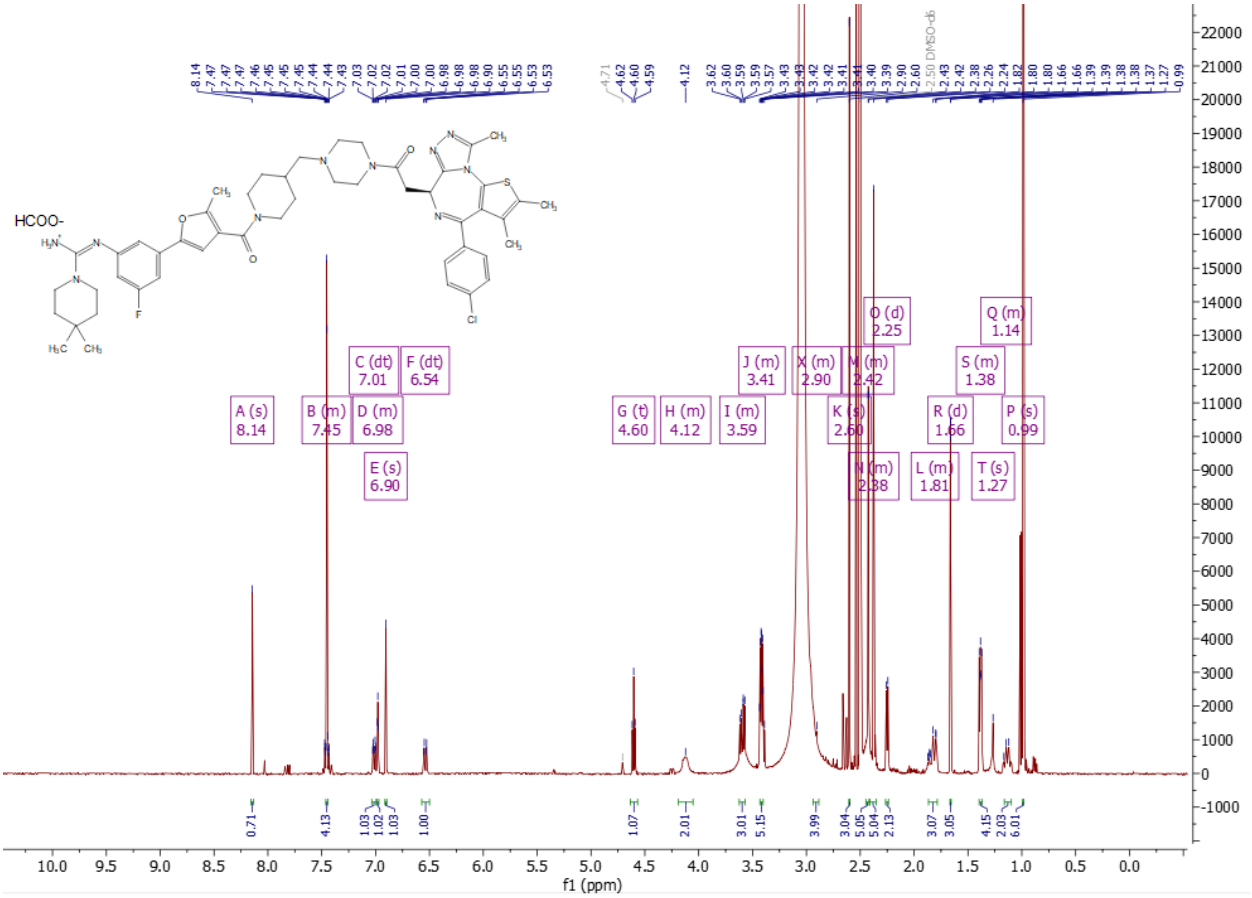

$^{1}$H-NMR (500 MHz) of (Z)-N'-[3-(4-{3-[(4-{2-[(9S)-7-(4-chlorophenyl)-4,5,13-trimethyl-3-thia-1,8,11,12-tetraazatricyclo[8.3.0.$0^{2,6}$]trideca-2(6),4,7,10,12-pentaen-9-yl]acetyl}piperazin-1-yl)methyl]piperidine-1-carbonyl}-5-methylfuran-2-yl)-5-fluorophenyl]-4,4-dimethylpiperidine-1-carboximidamide formate (**25**) in DMSO-$d_6$ at 80°C

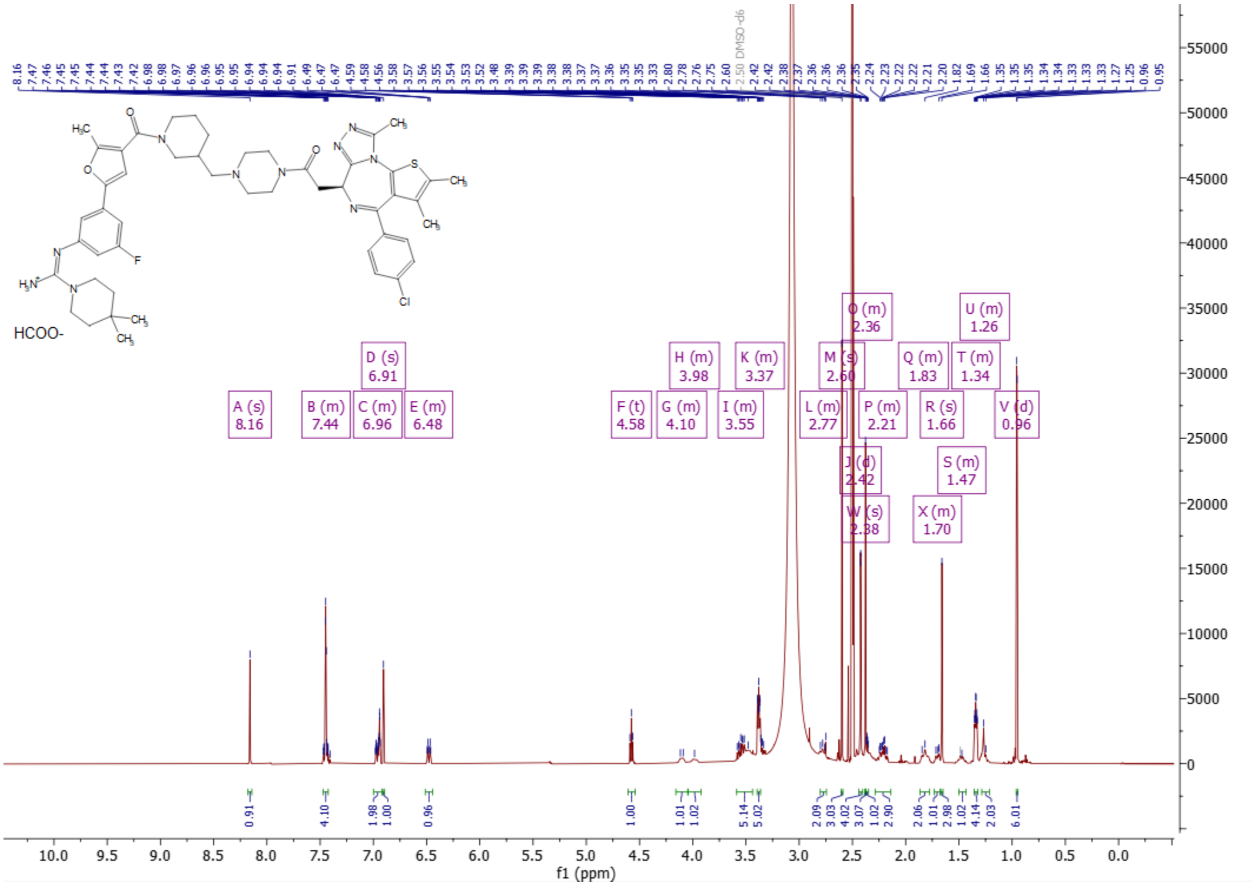

[1]H-NMR (500 MHz) of (Z)-N'-[3-(4-{4-[(4-{2-[(9R)-7-(4-chlorophenyl)-4,5,13-trimethyl-3-thia-1,8,11,12-tetraazatricyclo[8.3.0.0$^{2}$,$^{6}$]trideca-2(6),4,7,10,12-pentaen-9-yl]acetyl}piperazin-1-yl)methyl]piperidine-1-carbonyl}-5-methylfuran-2-yl)-5-fluorophenyl]-4,4-dimethylpiperidine-1-carboximidamide formate (**26**) in DMSO-$d_6$ at 80$^{o}$C

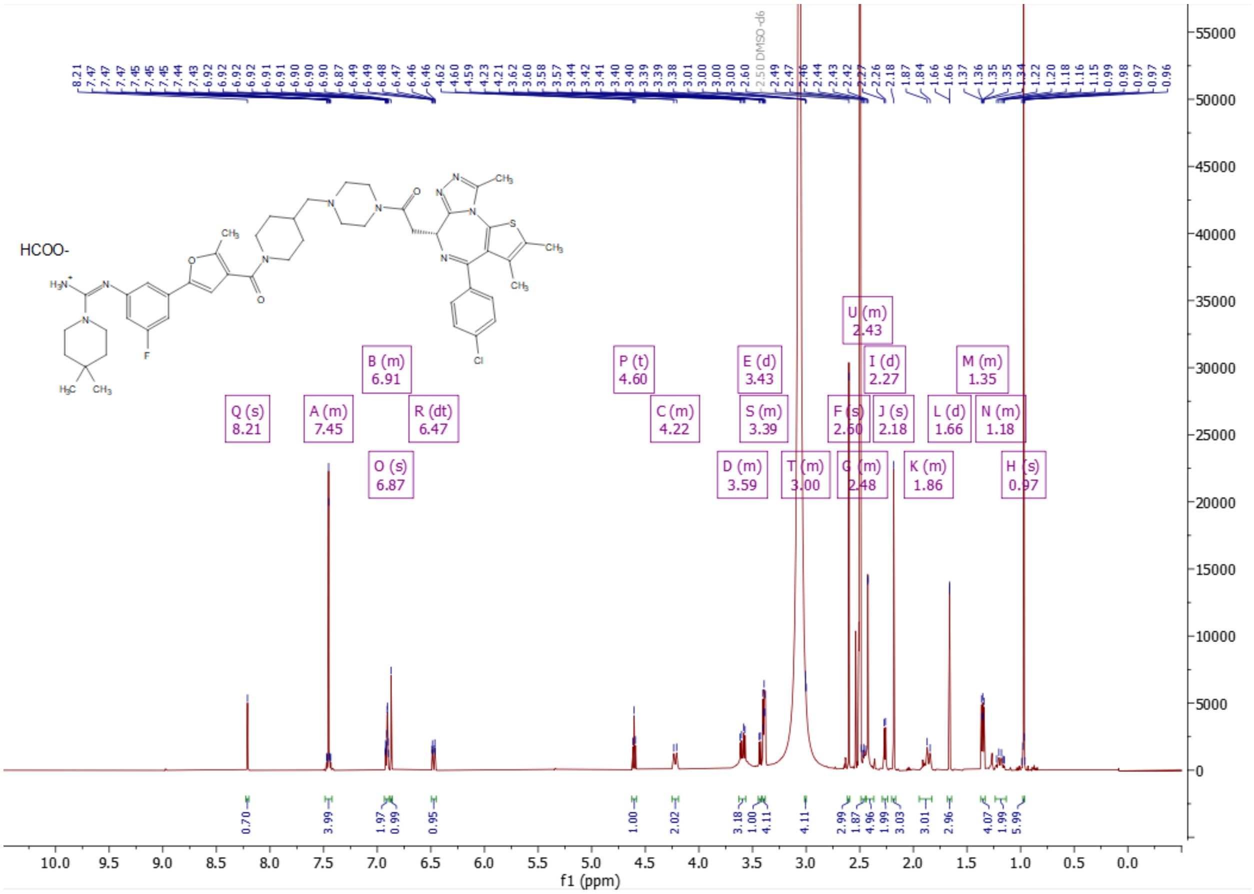

[1]H-NMR (500 MHz) of N-{2-[2-(2-{[1-(5-{3-[(Z)-[amino(4,4-dimethylpiperidin-1-yl)methylidene]amino]-5-fluorophenyl}-2-methylfuran-3-carbonyl)piperidin-4-yl]oxy}ethoxy)ethoxy]ethyl}-2-[(9S)-7-(4-chlorophenyl)-4,5,13-trimethyl-3-thia-1,8,11,12-tetraazatricyclo[8.3.0.0$^{2,6}$]trideca-2(6),4,7,10,12-pentaen-9-yl]acetamide (**27**) in DMSO-$d_6$ at 80°C

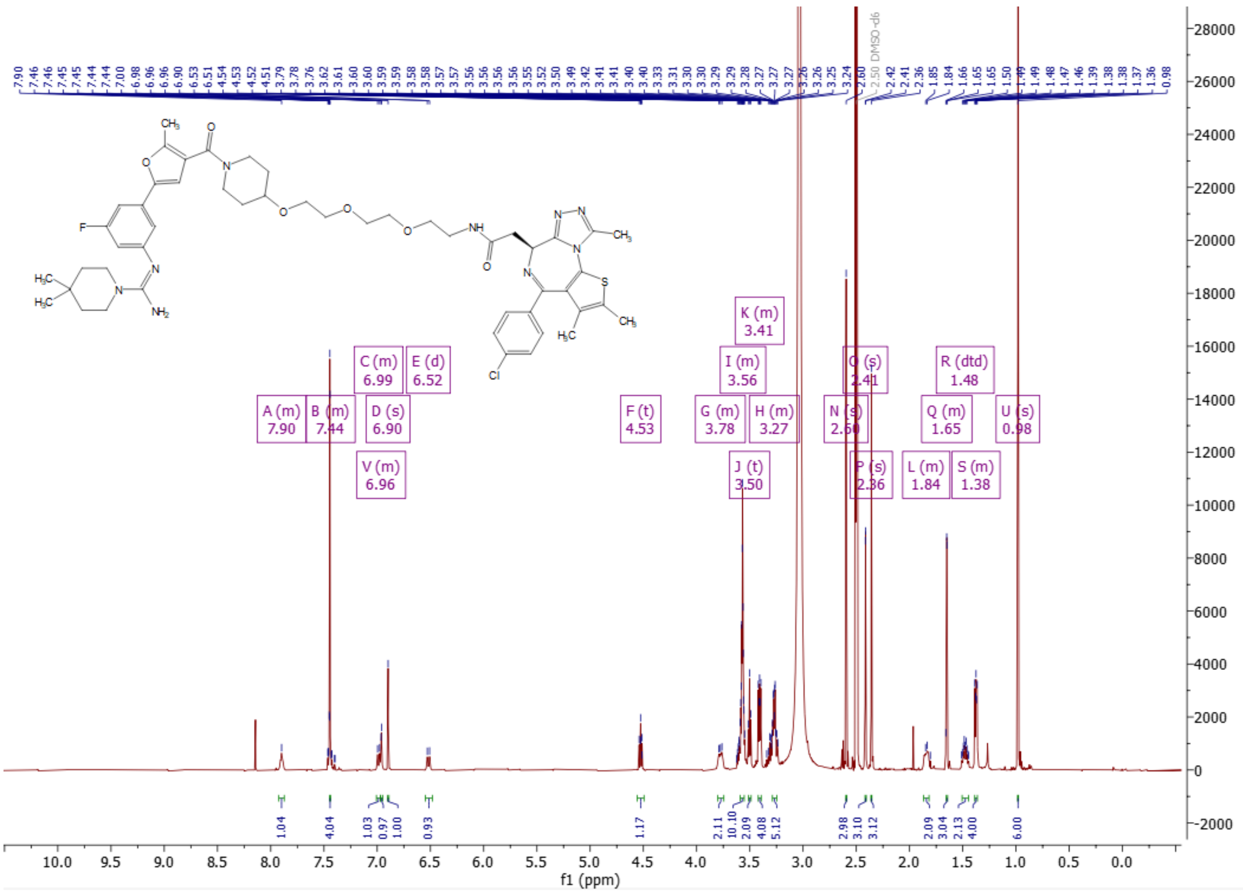

[1]H-NMR (500 MHz) of N-(2-(2-(2-((1-(5-(3-(((Z)-amino(4,4-dimethylpiperidin-1-yl)methylene)amino)-5-fluorophenyl)-2-methylfuran-3-carbonyl)piperidin-3-yl)oxy)ethoxy)ethoxy)ethyl)-2-((S)-4-(4-chlorophenyl)-2,3,9-trimethyl-6H-thieno[3,2-f][1,2,4]triazolo[4,3-a][1,4]diazepin-6-yl)acetamide formate (**28**) in DMSO-$d_6$ at 80°C

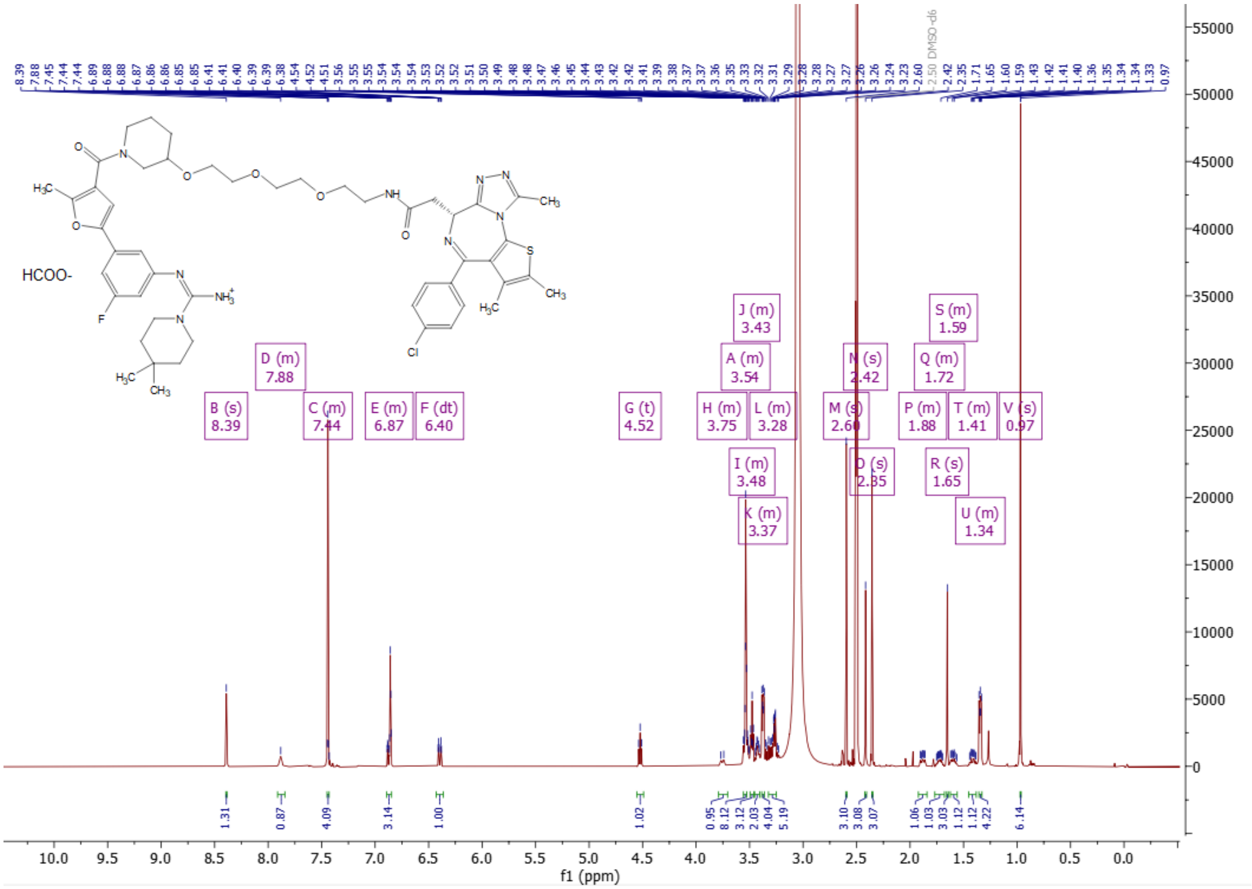

$^1$H-NMR (500 MHz) of N-(14-{[1-(5-{3-[(Z)-[amino(4,4-dimethylpiperidin-1-yl)methylidene]amino]-5-fluorophenyl}-2-methylfuran-3-carbonyl)piperidin-4-yl]oxy}-3,6,9,12-tetraoxatetradecan-1-yl)-2-[(9S)-7-(4-chlorophenyl)-4,5,13-trimethyl-3-thia-1,8,11,12-tetraazatricyclo[8.3.0.0$^{2,6}$]trideca-2(6),4,7,10,12-pentaen-9-yl]acetamide (**29**) in DMSO-$d_6$ at 80$^o$C

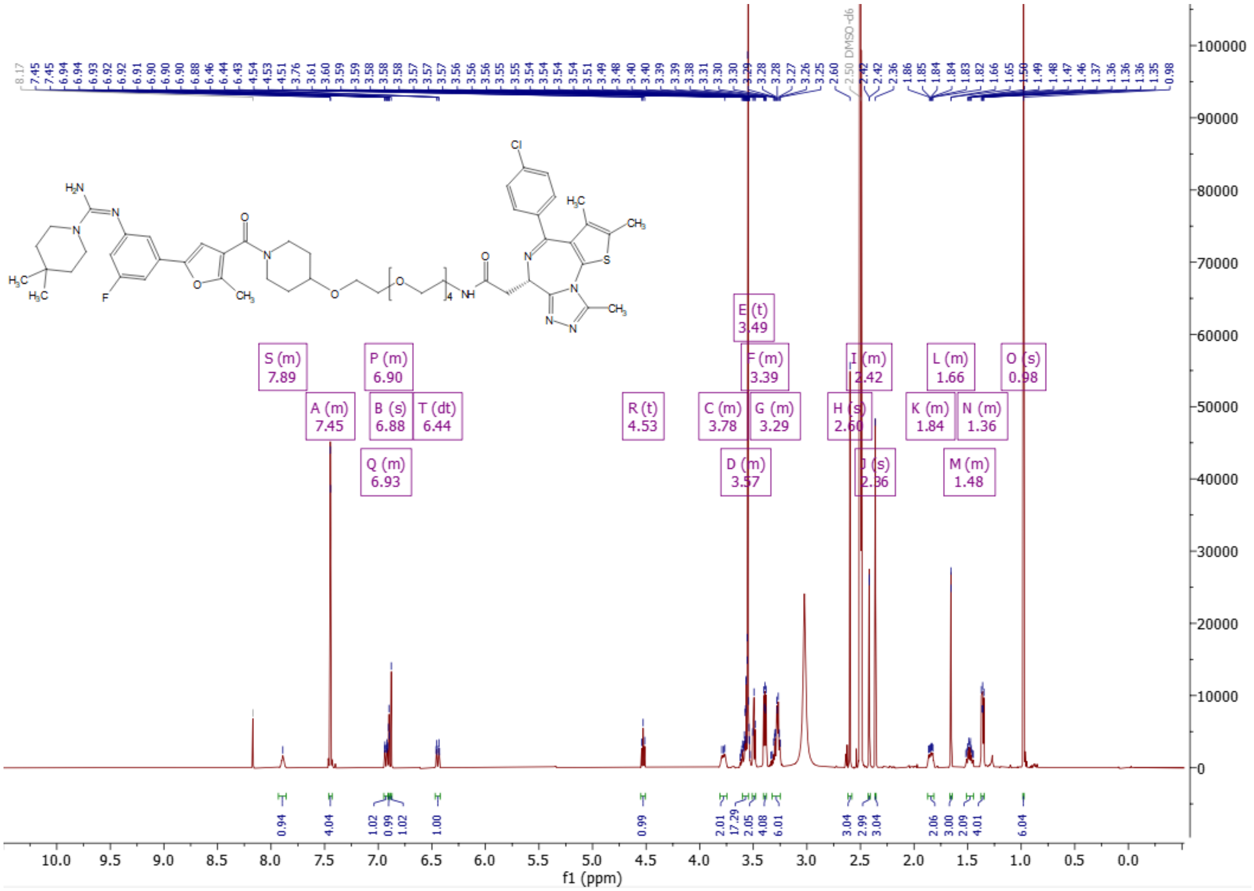

[1]H-NMR (500 MHz) of N-(14-{[1-(5-{3-[(Z)-[amino(4,4-dimethylpiperidin-1-yl)methylidene]amino]-5-fluorophenyl}-2-methylfuran-3-carbonyl)piperidin-3-yl]oxy}-3,6,9,12-tetraoxatetradecan-1-yl)-2-[(9S)-7-(4-chlorophenyl)-4,5,13-trimethyl-3-thia-1,8,11,12-tetraazatricyclo[8.3.0.0$^{2,6}$]trideca-2(6),4,7,10,12-pentaen-9-yl]acetamide (**30**) in DMSO-$d_6$ at 80°C

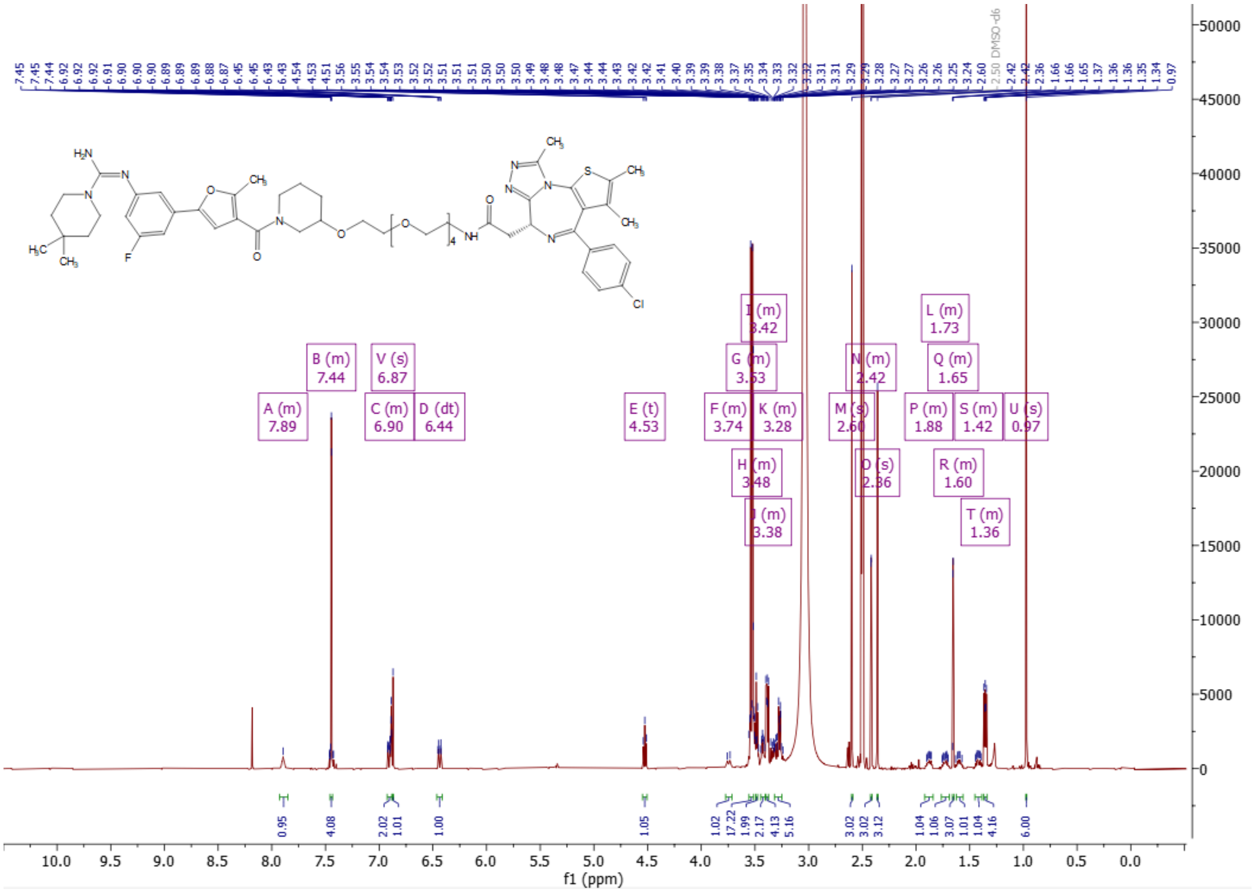

[1]H-NMR (400 MHz) of (Z)-N'-(3-chlorophenyl)-2-methylpiperidine-1-carboximidamide (**31**) in DMSO-d$_6$

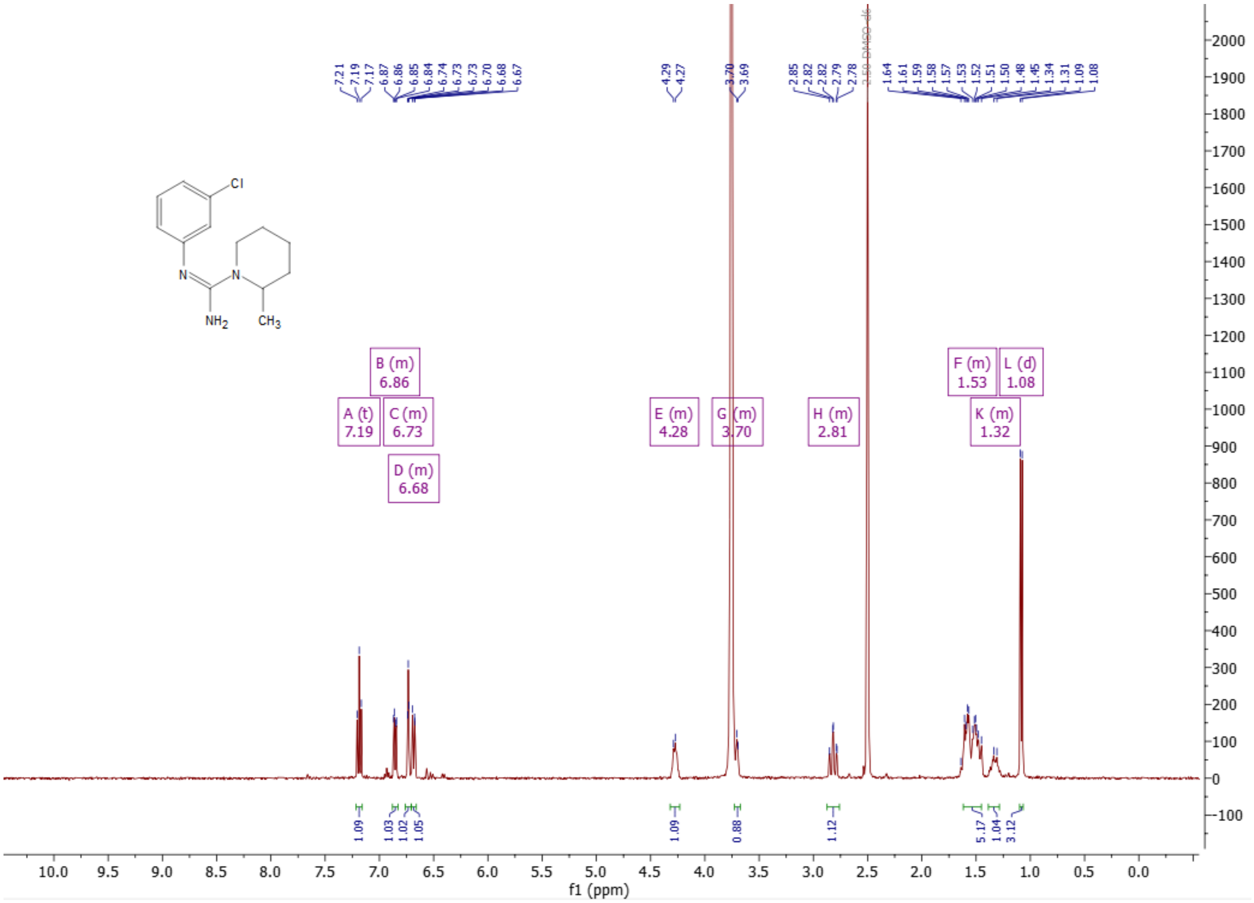

[1]H-NMR (400 MHz) of (Z)-N'-(3-chlorophenyl)-3-methylpiperidine-1-carboximidamide (**32**) in DMSO-$d_6$

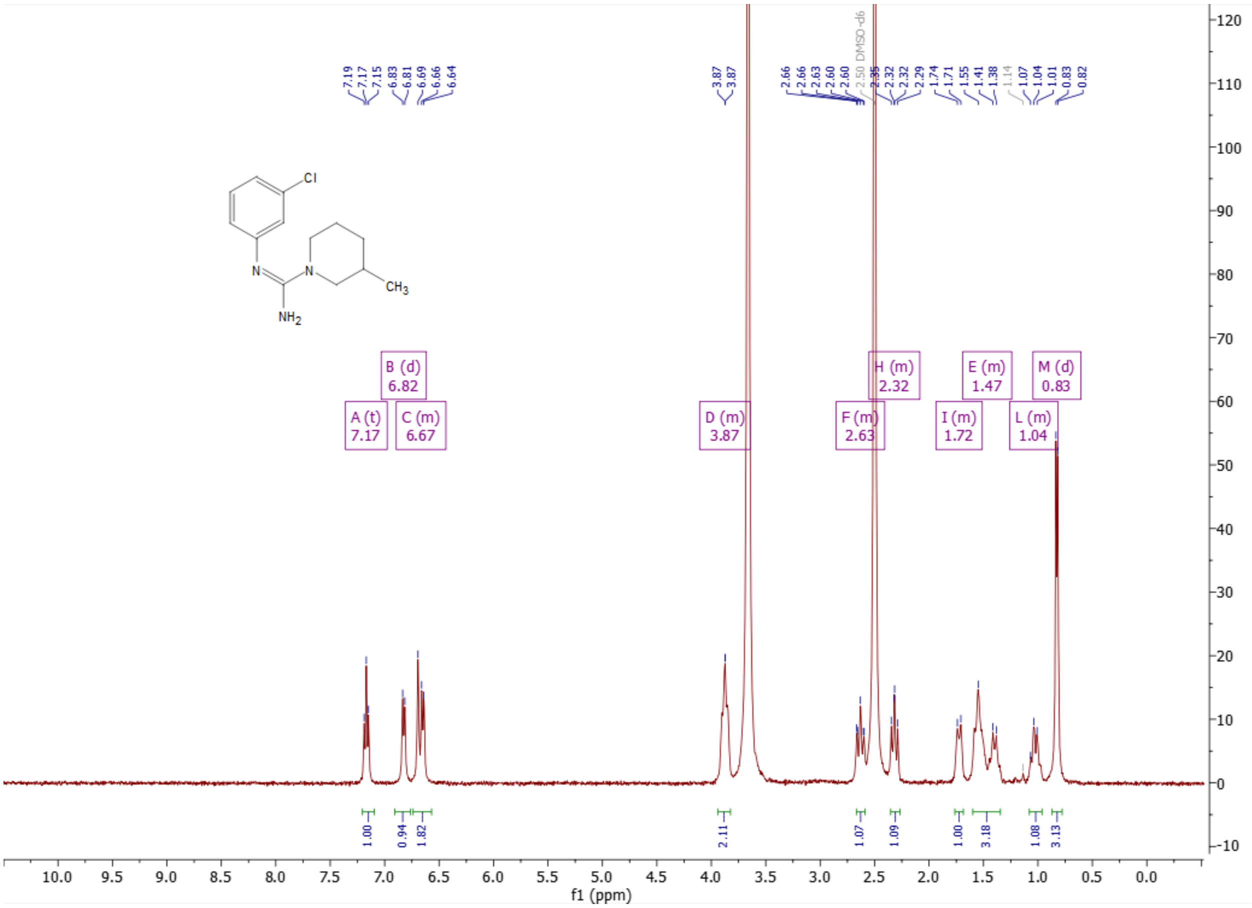

$^{1}$H-NMR (500 MHz) of (Z)-N'-(3-fluoro-5-(4-(4-(methoxymethyl)piperidine-1-carbonyl)-5-methylfuran-2-yl)phenyl)-4,4-dimethylpiperidine-1-carboximidamide formate (**33**) in $CD_3OD$

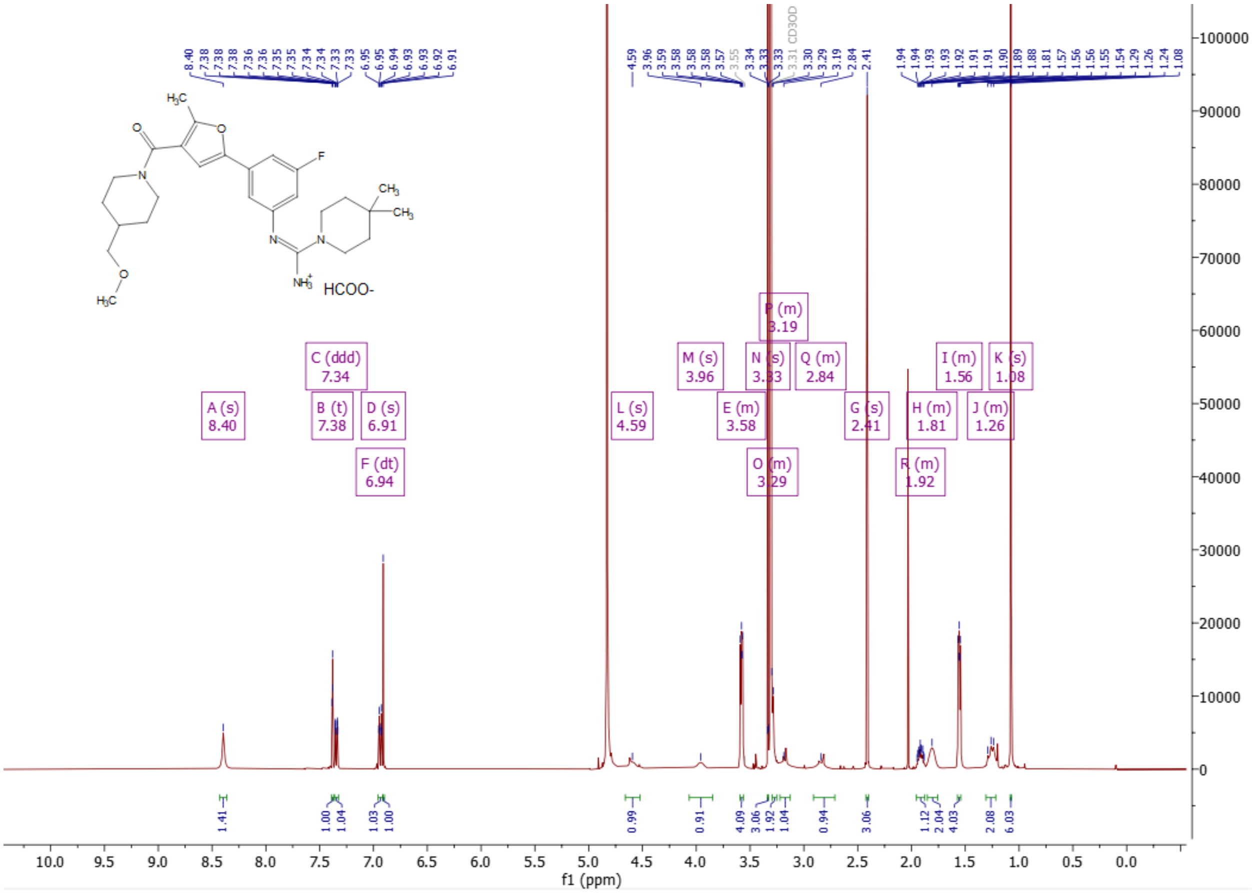
